# An unbiased template of the *Drosophila* brain and ventral nerve cord

**DOI:** 10.1101/376384

**Authors:** John A Bogovic, Hideo Otsuna, Larissa Heinrich, Masayoshi Ito, Jennifer Jeter, Geoffrey Meissner, Aljoscha Nern, Jennifer Colonell, Oz Malkesman, Kei Ito, Stephan Saalfeld

## Abstract

The fruit fly *Drosophila melanogaster* is an important model organism for neuroscience with a wide array of genetic tools that enable the mapping of individuals neurons and neural subtypes. Brain templates are essential for comparative biological studies because they enable analyzing many individuals in a common reference space. Several central brain templates exist for *Drosophila*, but every one is either biased, uses sub-optimal tissue preparation, is imaged at low resolution, or does not account for artifacts. No publicly available *Drosophila* ventral nerve cord template currently exists. In this work, we created high-resolution templates of the *Drosophila* brain and ventral nerve cord using the best-available technologies for imaging, artifact correction, stitching, and template construction using groupwise registration. We evaluated our central brain template against the four most competitive, publicly available brain templates and demonstrate that ours enables more accurate registration with fewer local deformations in shorter time.

## 1 Introduction and Related work

Canonical templates (or “atlases”) of stereotypical anatomy are vital in intersubject biological studies. In neuroscience, atlases of the central nervous system have become an important tool for cumulative and comparative studies. Neuroanatomical templates for humans (Talairach and Tournoux, 1988; Lancaster et al., 2000), mouse (Allen Institute for Brain Science, 2015; Lein et al., 2007), C. *elegans* (Long et al., 2009), and *Drosophila* (Jenett et al., 2012; Aso et al., 2014; Arganda-Carreras et al., 2018) have been valuable and influential for studies of anatomy, function, and behavior (Ashburner et al., 1998; Aso et al., 2014; Panser et al., 2016; Robie et al., 2017).

For a template space to be useful, it must be possible to reliably find spatial transformations between individual subjects and that template. The transformation must be capable of expressing the biological variability so that stereotypical anatomical features of many transformed subjects are well aligned in template space. This serves to normalize for “irrelevant” sources of variability while comparing across a population. Early templates used simple affine spatial transformations (Talairach and Tournoux, 1988), for which identifying a small number of stereotypical landmarks was sufficient. More recently, image registration has enabled the computation of more flexible (elastic, diffeomorphic, etc. Sotiras et al., 2013) transformations to be found between pairs of images. As a result, modern templates consist of a representative digital image of the anatomy and imaging modality of interest, which is used as the target (or “fixed” image) for image registration, which generates the spatial transformation (Maurer and Fitzpatrick, 1993).

Transforming individual subjects’ anatomy to a template space enables the comparison and analysis of a cohort of individuals in a common reference space. Typical registration approaches are either feature/landmark-based, as in Peng et al. (2011), or pixel based, using generic image registration software libraries, such as ANTs (Avants et al., 2009), CMTK (Rohlfing and Maurer, 2003), or elastix (Klein et al., 2010). If the template itself has anatomical labels superimposed, then it also enables an individual’s anatomy to be labeled via the spatial transformation to the template (Cabezas et al., 2011).

The fruit fly *Drosophila melanogaster* is an important model organism for neuroscience. The availability of powerful genetic tools that enable precise imaging and manipulation of specific neuronal populations (Luan et al., 2006; Pfeiffer et al., 2010; Tirian and Dickson, 2017) have made possible many important biological experiments. For example, the Fly-circuit database by Chiang et al. (2011) reconstructed approximately 16,000 neurons in *Drosophila* from light microscopy. Jenett et al. (2012) produced and imaged 6,650 GAL4 lines that have enabled the cataloging of a wide array of neurons in the *Drosophila* central nervous system. Among other advances, these resources enabled the creation of a spatial map of projection neurons for the lateral horn and mushroom body of *Drosophila* (Jefferis et al., 2007). Panser et al. (2016) generated a spatial clustering of the *Drosophila* brain into functional units on the basis of enhancer expression. Using over 2,000 GAL4 lines, Robie et al. (2017) created a whole brain map linking behavior to brain regions. Yu et al. (2013) explore *Drosophila* development, analyzing the connectivity of neuronal lineages by neuropil compartment.

Canonical templates and spatial alignment are an important aspect of the above studies that leverage genetic tools and imagery in *Drosophila*. Any given line labels a different sub-population of neurons and so it is necessary to perform comparisons in a canonical space. Many brain templates exist for *Drosophila*, a summary of which we give below.

Rein et al. (2002) generated a template starting with 28 individual brains, stained with nc82 (Kittel et al., 2006) and imaged at 0.6 × 0.6 × 1.1 µm resolution. The template brain was chosen as the individual with “the average volume for each substructure.” Registration was done by first estimating a global rigid, then estimating a per-structure rigid or similarity transformation which was interpolated over space.

Jenett et al. (2012) selected a representative confocal image of a female fly brain, imaged at 0.62 × 0.62 × 1.0 µm/px as a standard brain for their GAL4 driver line resource, and was later resampled in *z* to an isotropic resolution of 0.62 µm/px. We will refer to this template as JFRC 2010.

Aso et al. (2014) selected another single female brain for their work, called the JFRC 2013 template. It comprises five stitched tiles, imaged at 0.19 × 0.19 × 0.38 µm/px, then downsampled to an isotropic resolution of 0.38 × 0.38 × 0.38 µm/px. Since the JFRC 2010 and JFRC 2013 templates consist of a brain image of a single individual fly, we will call these “individual templates.”

Ito et al. (2014) introduced the “Ito half-brain” as their reference standard. It includes a rich set of compartment labels for neuropil boundaries and fiber bundles but automatic registration of whole brains to this standard is not straightforward.

The female, male, and unisex FCWB templates by Ostrovsky and Jefferis (2014) were generated from images manually selected from the FlyCircuit database (Chiang et al., 2011), imaged at 0.32 × 0.32 × 1.0 µm/px, using groupwise registration (Avants and Gee, 2004) with the CMTK registration software. Groupwise registration is the process of co-registering a set of images without specifying one particular image as the registration target and thereby avoids bias (see section Section 2.4 for a more detailed description). Seventeen brain samples were used for the female template and nine for the male template. The two gendered average brain templates were themselves registered and averaged with equal weight to create a unisex template.

Arganda-Carreras et al. (2018) recently used groupwise registration with the ANTs registration software to generate an improved unbiased *Drosophila* brain template from ten individual fly brains, imaged at 0.6 × 0.6 × 0.98 µm/px and labeled with the nc82 antibody. We call this the “Tefor” template. To measure registration performance, the authors compare overlap of anatomical labels of individuals after registration, and conclude that templates generated by groupwise registration outperform templates consisting of an individual brain image.

Given the abundance of brain templates for *Drosophila*, Manton et al. (2014) created “bridging transformations” that align many of these templates, including JFRC 2010, JFRC 2013, FCWB, and the Ito half-brain. Bridging transformations link previously disparate data-sets and thereby enable comparisons across all datasets in the space of any of these templates. The combination of neuronal database, image registration, and brain template has made neuron matching with NBLAST (Costa et al., 2016) an important technology. NBLAST has also recently found success matching neurons obtained from different, complementary data-sets (Schlegel et al., 2017), including the first complete electron microscopy (EM) volume of the female adult fly brain (FAFB) generated by Zheng et al. (2017). This image volume enables the complete tracing of every neuron and identification of synapses spanning the central brain in a single individual, and is potentially a very useful reference standard brain.

Meinertzhagen (2018) describes how neurons identified from LM will offer an important form of validation for neurons traced in EM, either manually or automatically. This capability depends on spatial alignment between the two modalities. A plugin for Fiji (Schindelin et al., 2012) called ELM (github.com/saalfeldlab/elm) is a custom wrapper of the BigWarp plugin (Bogovic et al., 2016), and was used to manually place landmark point correspondences between the LM template and generate a spatial transformation from the EM image space to the JFRC 2013 template space.

Using this registration, Zheng et al. (2017) showed that neurons traced in FAFB can be matched with neurons cataloged from light microscopy (LM), a capability that will enable researchers to simultaneously leverage the advantages of each modality: dense connectivity from EM, and cell type, neurotransmitter, gene expression, etc. information from LM (Schlegel et al., 2017).

While studies of the *Drosophila* ventral nerve cord (VNC) are numerous and ongoing (Prokop and Technau, 1991; Stockinger et al., 2005; Lacin and Truman, 2016; Ache et al., 2019), efforts in generating a standard anatomical coordinate system are somewhat lacking. Bö rner and Duch (2010) created a standard *Drosophila* VNC, but to our knowledge, the data are not publicly available; thus its impact is limited. Recently, Court et al. (2017) developed a standard nomenclature for the ventral nervous system.

In summary, all existing *Drosophila* brain templates lack certain desirable characteristics. Some consist of a individual samples and are therefore biased. Those that use groupwise registration to avoid bias use a small number of subjects, imaged at relatively low resolution and do not leverage new advances in tissue preparation, imaging, and artifact correction. No publicly available groupwise averaged ventral nerve cord template exists.

### 1.1 Contributions

We generated unbiased, symmetric, high-resolution, male, female, and unisex templates for the *Drosophila* brain and ventral nerve cord using the newest advances in tissue preparation, imaging, artifact correction, and image stitching. The images of individual samples that comprised the template were acquired at high resolution of 0.19 × 0.19 × 0.38 µm/px. We used groupwise image registration to ensure that the shape of the resulting template is not biased toward our choice of subject. Figure 1 shows the female, unisex, and male templates for the *Drosophila* central brain and ventral nerve cord. We call this set of templates “JRC 2018.”

**Figure 1:**
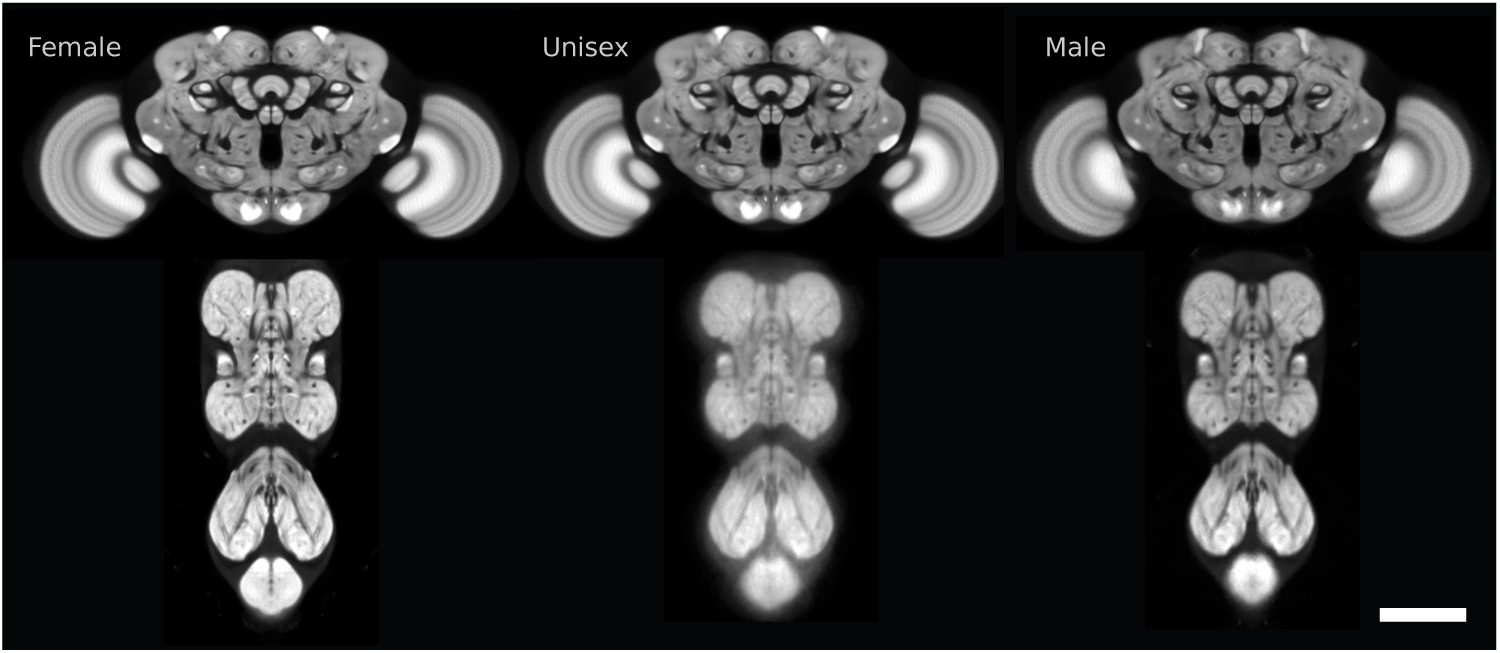
Slices of the six templates we created for female, unisex, and male *Drosophila* central brains and ventral nerve cords. Scale bar 100 µm.

We performed a thorough comparison using the four most competitive publicly available *Drosophila* brain templates, and three leading image registration software libraries, measuring registration quality, amount of deformation, and computational cost. Each template was evaluated using eight different choices of registration algorithms/parameter settings. We show that *Drosophila* brain samples register significantly *better*, *faster*, and with *less local deformation* to our JRC 2018 template than to any prior template, enabling more accurate comparison studies than were previously possible.

We also generated a new, automatic registration between the female full adult fly brain (FAFB) electron microscopy data-set of Zheng et al. (2017) using automated synapse predictions (Heinrich et al., 2018). This registration is more data-driven, less likely to suffer from variable error, and potentially better regularized than the existing manual registration generated with ELM (Zheng et al., 2017; Bogovic et al., 2016).

We developed software to create, apply, convert, and compare templates and transformations. Our software depends on the publicly available registration packages elastix, CMTK, and ANTs. Our specific contributions are:

1. Scripts and parameters for template construction (these are customized versions of scripts from ANTs).
2. Scripts and parameters for registration used in evaluation and analysis, including registration quality estimation (see Section 3.2.1).
3. Software for applying transformations to images, skeletons, and sets of point coordinates. Both command line utilities and Fiji (Schindelin et al., 2012) plugins are available.
4. A new compressed HDF5 based format (see Appendix A.8) for multiscale transformations and conversion tools between this new format (including quantization and downsampling) and transformations generated by the registration packages elastix, CMTK, and ANTs.

The templates and transformations can be found on-line at https://www.janelia.org/open-science/jrc-2018-brain-templates. Software and code supporting these resources are available at https://github.com/saalfeldlab/template-building.

## 2 Materials and Methods

### 2.1 Sample Preparation and Image Acquisition

#### 2.1.1 Template data

All samples were based on the transgene brp-SNAP (Kohl et al., 2014) and labeled with 2 µM Cy2 SNAP-tag ligand (Grimm et al., 2017). Samples were fixed for 55 minutes in 2% PFA at room temperature, SNAP-tag labeled for 15 minutes at room temperature, then fixed for 4 hours in 4% PFA at room temperature. Samples were dehydrated in ethanol, cleared in xylene, and mounted in DPX, as described at https://www.janelia.org/project-team/flylight/protocols (Aso et al., 2014; Nern et al., 2015). Samples were imaged unidirectionally on six Zeiss 710 LSM confocal microscopes with Plan Apo 63×/1.4 Oil DIC 420780/2-9900 lenses, at a resolution of 0.19 µm/px, in tiles of 1024 × 1024 px (192.6 × 192.6 µm), a pixel dwell time of 1.27 µm, a *z*-interval of 0.38 µm, and a pinhole of 1 AU to 488 nm. Brains were imaged in five overlapping tiles, VNCs in three. Scanning was controlled by Zeiss ZEN 2010 software and a custom MultiTime macro, as described by Jenett et al. (2012).

Tiles were corrected for lens-distortion and chromatic aberration (see Section 2.2) and stitched with Fiji’s (Schindelin et al., 2012) stitching plugin (Preibisch et al., 2009). Custom scripts were used to parallelize processing on the Janelia CPU cluster. For template construction, 62 central brains (36 female) and 75 ventral nerve cords (36 female) were acquired.

#### 2.1.2 Evaluation data

For evaluation, we chose 20 female flies imaged with both an nc82 channel and a channel in which neuronal membrane of a split GAL4 driver line were labeled with a myristoylated FLAG reporter, as described by Aso et al. (2014). We selected four split GAL4 lines that label neurons with broad arborization, that together, cover nearly the whole brain. Maximum intensity projections of the neuronal membrane channel for these lines are shown in Figure 2. Our testing cohort consisted of 20 individuals in total, summarized in Table 1. As a result, our evaluation includes measurements across the whole brain and does not focus on a particular subset of the anatomy.

**Figure 2:**
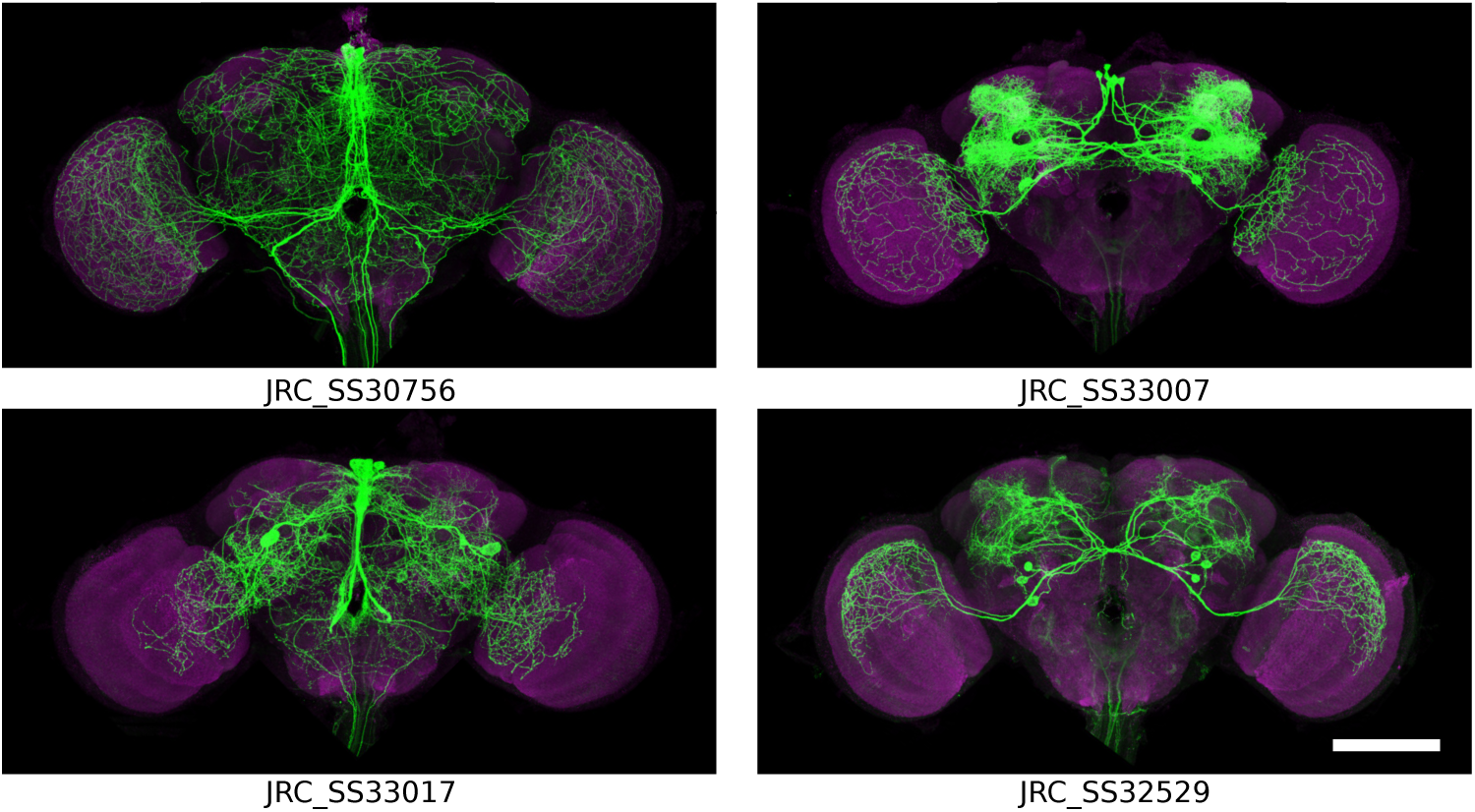
Maximum intensity projections of individuals from four of the GAL4 driver lines used to evaluate registration accuracy (magenta nc82, green GAL4). Notice the broad arborization spanning most of the central brain and optic lobes. Scale bar 100 µm.

**Table 1:** Split GAL4 driver lines chosen for evaluation with the count of the number of subjects per line (#).

| Line |  | # |
| --- | --- | --- |
| JRC_SS23204 | GMR_13F04-p65ADZp in attP40 and GMR_13B05-ZpGDBD in attP2 | 2 |
| JRC_SS23206 | GMR_28F06-p65ADZp in attP40 and GMR_13B05-ZpGDBD in attP2 | 2 |
| JRC_SS23208 | GMR_40F04-p65ADZp in attP40 and GMR_13B05-ZpGDBD in attP2 | 2 |
| JRC_SS30756 | GMR_15C12-p65ADZp in attP40 and GMR_64B11-ZpGDBD in attP2 | 3 |
| JRC_SS30777 | GMR_70D06-p65ADZp in attP40 and GMR_64B11-ZpGDBD in attP2 | 2 |
| JRC_SS33007 | GMR_54H04-p65ADZp in attP40 and GMR_10D10-ZpGDBD in attP2 | 3 |
| JRC_SS33017 | GMR_67A07-p65ADZp in attP40 and GMR_35G08-ZpGDBD in attP2 | 3 |
| JRC_SS32529 | GMR_21G11-p65ADZp in attP40 and GMR_78H08-ZpGDBD in attP2 | 3 |

### 2.2 Lens Distortion and Chromatic Aberration Correction

#### 2.2.1 Re-usable Calibration Slides

Measurements to establish and update the calibration models were made by imaging multicolor beads mounted, like the tissue samples, on #1.5H cover slips. The beads are 1 µm silica (Polysciences 24326-15), functionalized through reaction with tri-ethoxy-sliane-PEG biotin (Nanocs #PEG6-0023). The biotinylated surface was labeled with a mixture of dye-labeled streptavidin. The dyes were Alexa 488 (ThermoFisher S32354), cy3 (Jackson Immunoresearch 016-160-084), Alexa 594 (ThermoFisher S32356), and Atto647N (Sigma-Aldrich 94149-1MG). The beads were deposited out of tris-buffered saline onto plasma cleaned coverslips, dried, and mounted in DPX according to the same tissue mounting protocol described above.

#### 2.2.2 Distortion Correction

Image stack mosaics of 4 × 4 tiles with 50–60% overlap were taken using the same settings as described above. Image stacks were acquired in two passes, first at 488 nm and 594 nm, then at 488 nm, 561 nm, and 647 nm.

Channels were separated and max-intensity projections for each single channel tile were created with a custom Fiji script. Single channel 4 × 4 mosaics were imported as individual layers in TrakEM2 (Cardona et al., 2012). Channel mosaics were pre-stitched to account for stage shift. A non-linear lens-distortion correction model for each channel was estimated with a new extended version of the method by Kaynig et al. (2010) that we made available in TrakEM2. Lens-corrected channel mosaics were stitched and then all channels were globally aligned with an affine transformation model. The composition of the non-linear lens-correction model and the affine alignment transformation correct for both lens-deformation and chromatic aberration. These correction models were generated and exported, for all confocal microscopes (five Zeiss LSM 710’s and one Zeiss LSM 780). Finally, they were applied to individual 3D image stacks of *Drosophila* brain samples prior to stitching.

We recorded detailed video instructions to reproduce the calibration protocol and made them available on Youtube: https://www.youtube.com/watch?v=lPt-WQuniUs. Code can be found on-line at https://github.com/saalfeldlab/confocal-lens.

### 2.3 Neuron skeletonization

We computed skeletons from the neuron channel by first applying 3D direction-selective local-thresholding (DSLT) (Kawase et al., 2015) to each raw image tile. DLST convolves the image with multiple scaled and rotated cylindrical kernels. We used kernel radii of 2, 6, and 10 voxels. The maximum response was thresholded to give a neuron mask. This mask was transformed along with the tile during image stitching, and then skeletonized using the 3D Skeletonization plugin in Fiji (Schindelin et al., 2012).

### 2.4 Template construction

We constructed our template using groupwise registration, which seeks to find both an average shape and average intensity across all individuals in a cohort and is described in Avants and Gee (2004) and Avants et al. (2011) The script buildtemplateparallel.sh that is part of the ANTs library, implements this. Our modifications, described below, can be found on-line at https://github.com/saalfeldlab/template-building.

Groupwise registration begins with an initial template - often a single individual or the average of the unregistered cohort of images is selected (we chose the latter). Next, every individual image is registered to that initial template using a particular registration algorithm (transformation model, similarity measure, optimization scheme) and then transformed to the template space. Next, the set of transformed individual images are averaged and the transformations are averaged. This yields new mean intensities as well as a mean transformation from each subject to the current template estimate. Finally, the new mean intensity image is transformed through the inverse of the mean transformation to obtain a new template. This procedure of registration-averaging-transformation is iterated to transport the initial template toward the mean intensity and shape.

We made several changes seeking to reduce the amount of deformation possible during registration. First, we used the elastic transformation model rather than SyN diffeomorphic model (Avants et al., 2009). Second, we regularized the transform more strongly, the details of which can be found in our open source repository. We sought to generate a template with left-right symmetry. Similar to the approach in Allen Institute for Brain Science (2015), we left-right flipped every individual brain and VNC in our cohort and included both the original and flipped images in the groupwise registration procedure. This doubled both our effective image count and computational expense during groupwise registration.

We downsampled the raw and left-right flipped images to 0.76 µm/px isotropic resolution prior to running groupwise registration in order to reduce computational cost/runtime. We believe that this downsampling did not have an appreciable effect on the template construction. This is supported by experiments we performed showing that lower resolution templates can have similarly good performance as high resolution templates (see Appendix A.4). The final template was obtained by applying the transformation computed at low resolution to the original, high resolution images, resulting in a high resolution template. All templates were rendered at 0.19 µm/px isotropic resolution. This equals the *xy*-resolution of the original images, but has a higher *z*-resolution (by about a factor of 2). We chose this primarily for convenience, as the only downside is the additional storage space.

We applied this procedure to different subsets of our image data to produce female, male and unisex templates. For the central brain, we used 36 female individuals (72 images including left-right flips) for the female template, 26 male individuals (52 image with left-right flips) for the male template, and the union of both for the unisex brain template: 62 individuals (124 images with left-right flips). For the ventral nerve cord we used 36 female individuals (72 images including left-right flips) for the female template, 39 male individuals (78 image with left-right flips) for the male template, and the union of both for the unisex VNC template: 75 individuals (150 images with left-right flips).

### 2.5 Bridging registrations

Transformations between different template spaces have proven useful in neuroscience because they unify disparate sets of data that are each aligned to different templates (Manton et al., 2014). We computed forward and inverse bridging registrations between our template and the other templates we evaluated in this work: JFRC 2010, JFRC 2013, FCWB, and Tefor. We used the algorithm (“ANTs A”) that we found performs best overall (see Appendix A.2). In Figure 3, we show these templates in the space of JRC 2018 and vice versa. Qualitatively, these registrations appear to be about as accurate as those from individuals to templates. We used these transformations to normalize distances measured across templates (see Section 3.2). We also provide transformations between the unisex template and the male and female templates to facilitate inter-sex comparisons.

**Figure 3:**
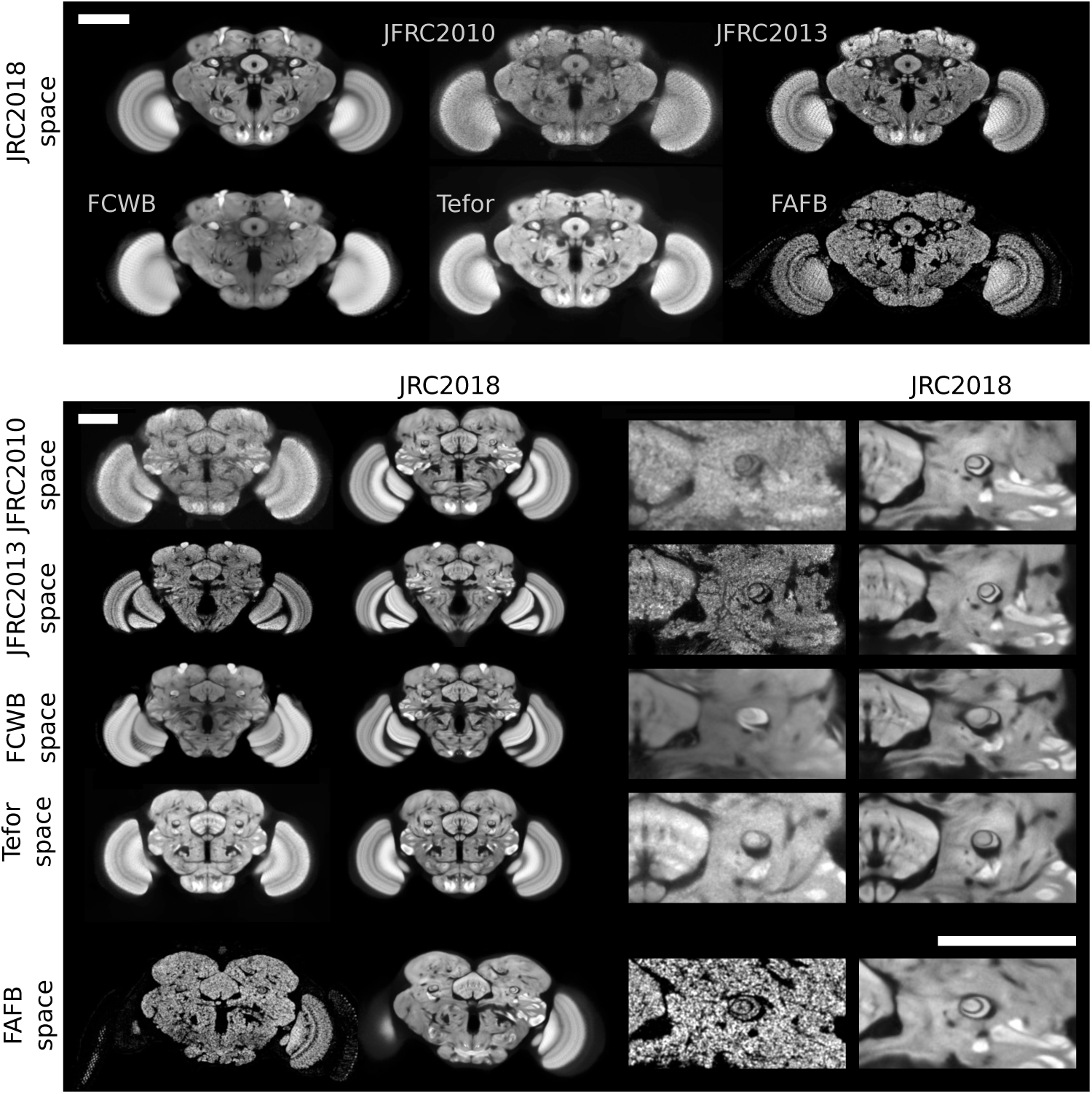
Visual comparison of *Drosophila* brain templates and bridging transformations. The top two rows show four existing templates registered to our JRC 2018 female template, as well as synaptic cleft predictions derived from the FAFB EM volume, transformed into the space of JRC 2018F. The middle four rows show JRC 2018F (second and fourth columns) registered to each of the three templates, along with a close-up around the fanshaped body and the pedunculus of the mushroom body. The bottom row shows JRC 2018F transformed into the space of FAFB. Scale bars 100 µm.

### 2.6 Registration with electron microscopy

We automatically aligned the FAFB EM volume and the JRC 2018 female template (see Figure 3). We rendered and blurred (with a 1.0 µm Gaussian kernel) the synapse cleft distance predictions generated by Heinrich et al. (2018) at low resolution (1.02 × 1.02 × 1.04 µm) so that the resulting image has an appearance similar to an nc82 or brp-SNAP labeled confocal image. We used ANTs A to find a transformation between the two images, the result of which produces a qualitatively accurate alignment.

In Appendix A.7, we discuss additional motivation and tradeoffs, including additional visualizations and a quantitative evaluation. In particular, we show that this transformation reproduces scientific results by Zheng et al. (2017). The accuracy of the manual registration between FAFB and JFRC 2013 by Zheng et al. (2017) suffers from variable error caused by the preference of the human annotator for specific brain regions and spurious details. Our automatic registration does not have this preference and is potentially better regularized.

## 3 Results

We compared the registration tools ANTs (Avants et al., 2009), CMTK (Rohlfing and Maurer, 2003), and elastix (Klein et al., 2010) using three sets of parameters for ANTs and CMTK and two sets of parameters for elastix, giving a total of eight different registration algorithms. We used images from 20 female flies for evaluation, with details described in Section 2.1.2. Since all testing subjects were female, we used the female JRC 2018 template for the experiments below, but will refer to it as simply “JRC2018F.”

### 3.1 Qualitative comparison

In Figure 3, we visually compare slices through the JFRC 2010 (Jenett et al., 2012), JFRC 2013 (Aso et al., 2014), FCWB (Ostrovsky and Jefferis, 2014), Tefor (Arganda-Carreras et al., 2018), and the JRC 2018F brain templates. Note the improved contrast and sharpness of anatomical structures in our JRC 2018F template relative to the others. We describe possible reasons for this in the Section 4.

Figure 4 shows *xz*-slices through the five templates and FAFB, where the *x*-axis is medial-lateral and the *z*-axis is anterior-posterior. Confocal imaging results in poorer *z*-than *xy*-resolution: 0.38 µm/px vs. 0.19 µm/px for our images. The brain templates vary notably in physical size and resolution (see Table 2). Specifically, the FCWB and JFRC 2010 templates are physically much smaller in the anterior-posterior direction. The affine part of the bridging transformations computed for Figure 3 indicate that the FCWB template is about 40% smaller than JRC 2018F, and JFRC 2010 is about 20% smaller.

**Figure 4:**
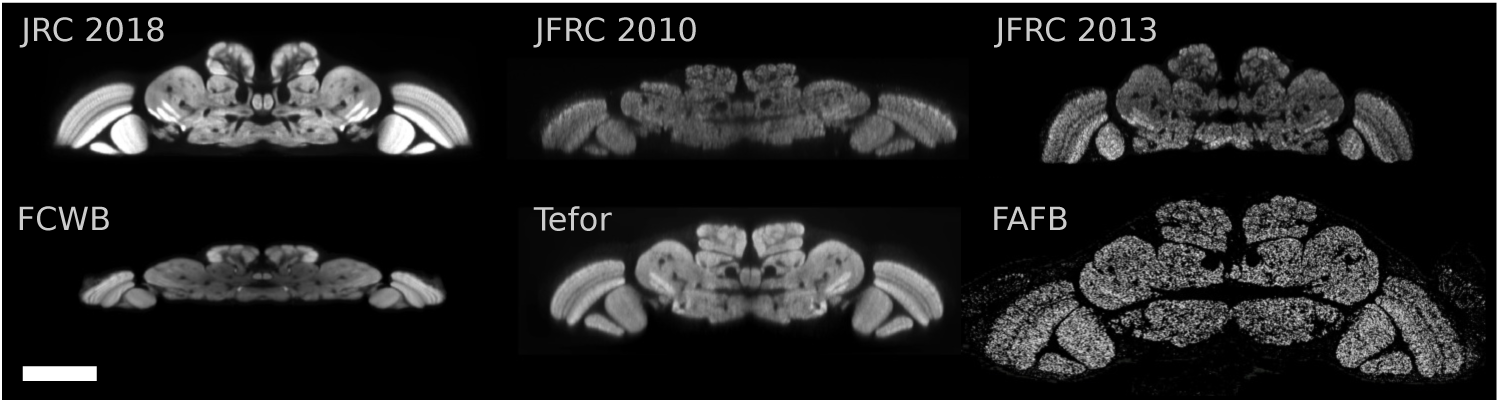
Horizontal (*xz*-slice) visualization of five brain templates and FAFB in physical coordinates. Note that the lower *z*-resolution is appreciable for individual templates (JFRC 2010 and JFRC 2013). Furthermore, observe the significant differences in physical sizes across these brain templates. Scale bar 100 µm.

**Table 2:** The resolutions of all templates in µm/pixel. Evaluation against our JRC 2018F template was performed at a resolution of 0.62 µm/px or less. We added the resolution of the FAFB synapse cloud for comparison.

|  | JRC 2018F | JFRC 2010 | JFRC 2013 | FCWB | Tefor | FAFB |
| --- | --- | --- | --- | --- | --- | --- |
| $xy$ | 0.19 | 0.62 | 0.62 | 0.32 | 0.61 | 0.004 |
| $z$ | 0.19 | 0.62 | 0.62 | 1.0 | 0.99 | 0.04 |

We make additional qualitative comparisons available in the supplementary material. Figure 8 in Appendix A.1 shows an overlay a registered image to each template, for one choice of registration algorithm. Appendix C contains much more detail, showing registration results for every template and algorithm.

### 3.2 Quantitative comparison

Measuring and evaluating the accuracy of image registration is notoriously challenging (Rohlfing, 2012). Common measures include image similarity, distance between landmark points, or overlap of semantic (anatomical) regions, each of which come with their own disadvantages. Image similarity can suffer from bias because the measurement may be the same quantity the registration algorithm is optimizing for (or a correlated quantity) and does not account for implausible deformation. Landmark points measure accuracy sparsely and are costly to generate (since they are usually manually placed). Region overlap is even more costly and only sensitive to differences along or near boundaries of those regions. We discuss this in more detail in Section 4.1.

#### 3.2.1 Registration accuracy measure

We measure registration accuracy with a näıve per-node skeleton distance between the same neuronal arbor for different individual flies after registration to the template and normalizing for template size. Normalizing for size and shape is necessary given the observations from Figure 4 that some templates are physically smaller than others. Transforming skeletons to a physically smaller space would artificially decrease the distance measure. Similarly, if shape differs, regions with locally dense distributions of arbors could be compressed, leading to smaller distances for many point samples. We compute this distance using the following procedure. Neurons are skeletonized using the methodology described in Section 2.3. Nc82 channels for two *Drosophila* individuals are independently registered to a template. The transformation found using the nc82 channel is then applied to the neuronal skeleton to bring them into template space. This is followed by an additional normalizing transformation that ensures all templates are at an equivalent scale and shape, that of JRC 2018F (see Section 2.5). The distance transform of both skeletons is computed and rendered at an isotropic resolution of 0.5 µm/px. For a given point (pixel) on one skeleton, the value of the distance transform of the other skeleton gives the näıve orthogonal distance between the skeletons. We report statistics of this skeleton-distance (in µm) both over the entire brain and split by compartment labels defined by JFRC 2010. Image processing was done using custom code based on ImgLib2 (Pietzsch et al., 2012). Statistics, analysis and visualization used pandas (McKinney, 2010) and matplotlib (Hunter, 2007). All evaluation code can be found on-line at https://github.com/saalfeldlab/template-building.

The skeleton distance is similar to landmark distance, but involves no manual human decision making or interaction since the structures of interest are directly specified by the anatomy. Our approach has the advantage of being fast and simple to compute as well as providing a distance for every point on the skeleton. It is limited in that it only measures distances perpendicular to the skeletons and assumes strict correspondence between the nearest points on two skeletons. It will therefore tend to underestimate errors. See Section 4.1 in Section 4 for more details on the benefits and limitations of this performance measure.

We measured the skeleton distance between all pairs of individuals belonging to the same line, though we grouped the first five lines in Table 1 (JRC SS23204, JRC SS23206, JRC SS23208, JRC SS30756, JRC SS30777) since they have very similar expression patterns. There exist 110 permutations of the 11 flies in the first line-group and 6 permutations each of the other three lines, making 128 pairwise comparisons in total. This distance is computed at every pixel for a given skeleton pair. When rendered at 0.5 µm resolution, each skeleton comprises about 250,000 pixels on average, yielding about 32 million points for which distance is estimated across all 128 skeleton pairs. This is repeated for each template and algorithm pair we evaluated.

We then compute several statistics of the distance across pairs of skeletons with a fixed registration algorithm and template. Table 3 shows the mean and standard deviation of the distance distribution for different templates and algorithms. JRC 2018F has the lowest mean values of distance (indicating best performance) both when averaging across algorithms, and when choosing the best algorithm for a given template. Figures 5 and 6 plot skeleton distance against a deformation measure (see Section 3.2.2) and CPU time, respectively. We omit p-values here because their magnitude is largely a reflection of our large sample size (all pairwise hypothesis tests are significant). Table 3 shows the mean and standard deviation of skeleton distances averaged across 128 pairs of flies (four driver lines). The second column gives the mean distance averaged also across eight choices of registration algorithm/parameters, while the third and fourth columns give average distances for the “best” two algorithms (see Table 5). The JRC 2018F template performs best according to this measure, independent of the registration algorithm used.

**Figure 5:**
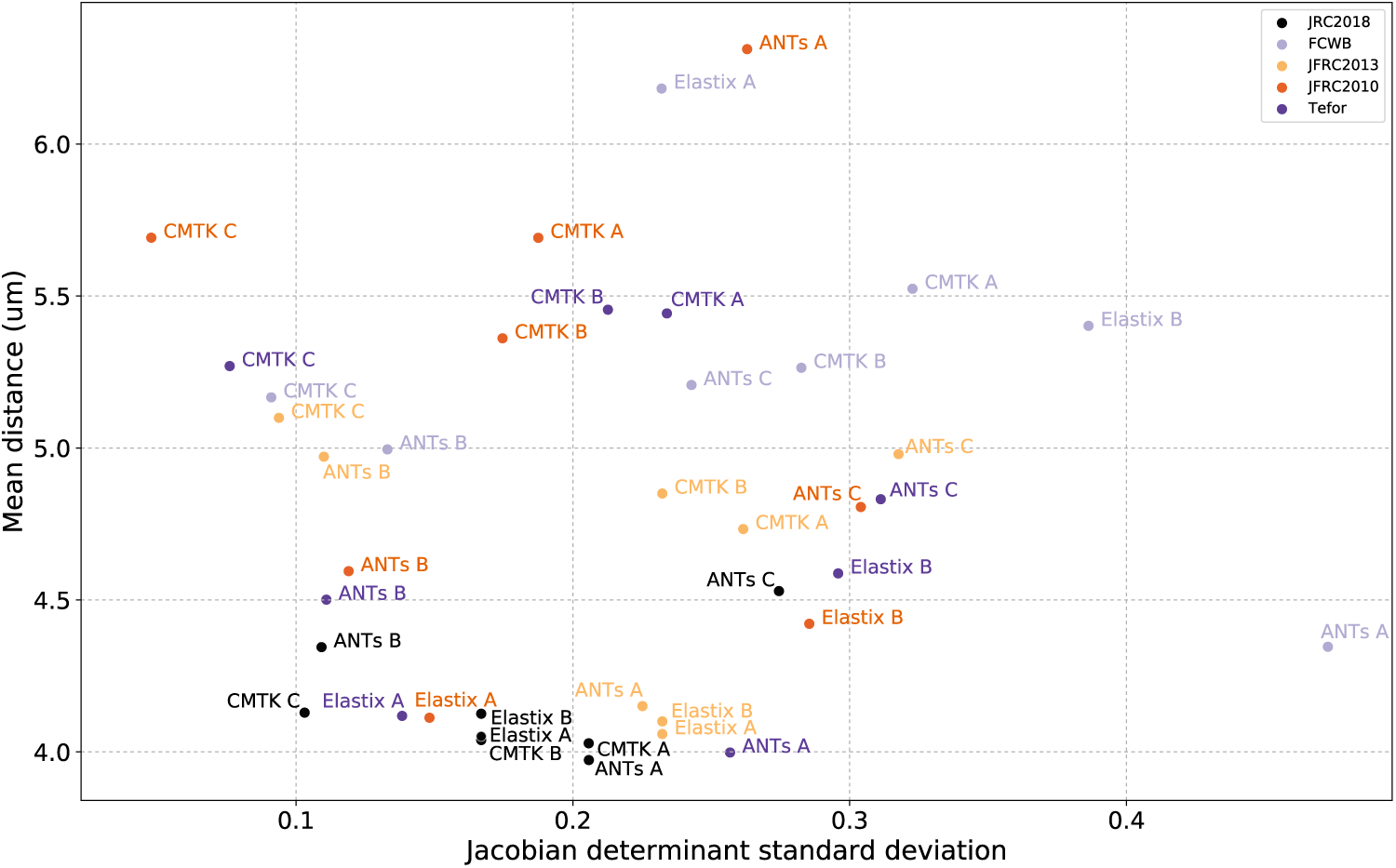
Scatterplot showing mean skeleton distance and standard deviation of the Jacobian determinant for all template-algorithm pairs.

**Figure 6:**
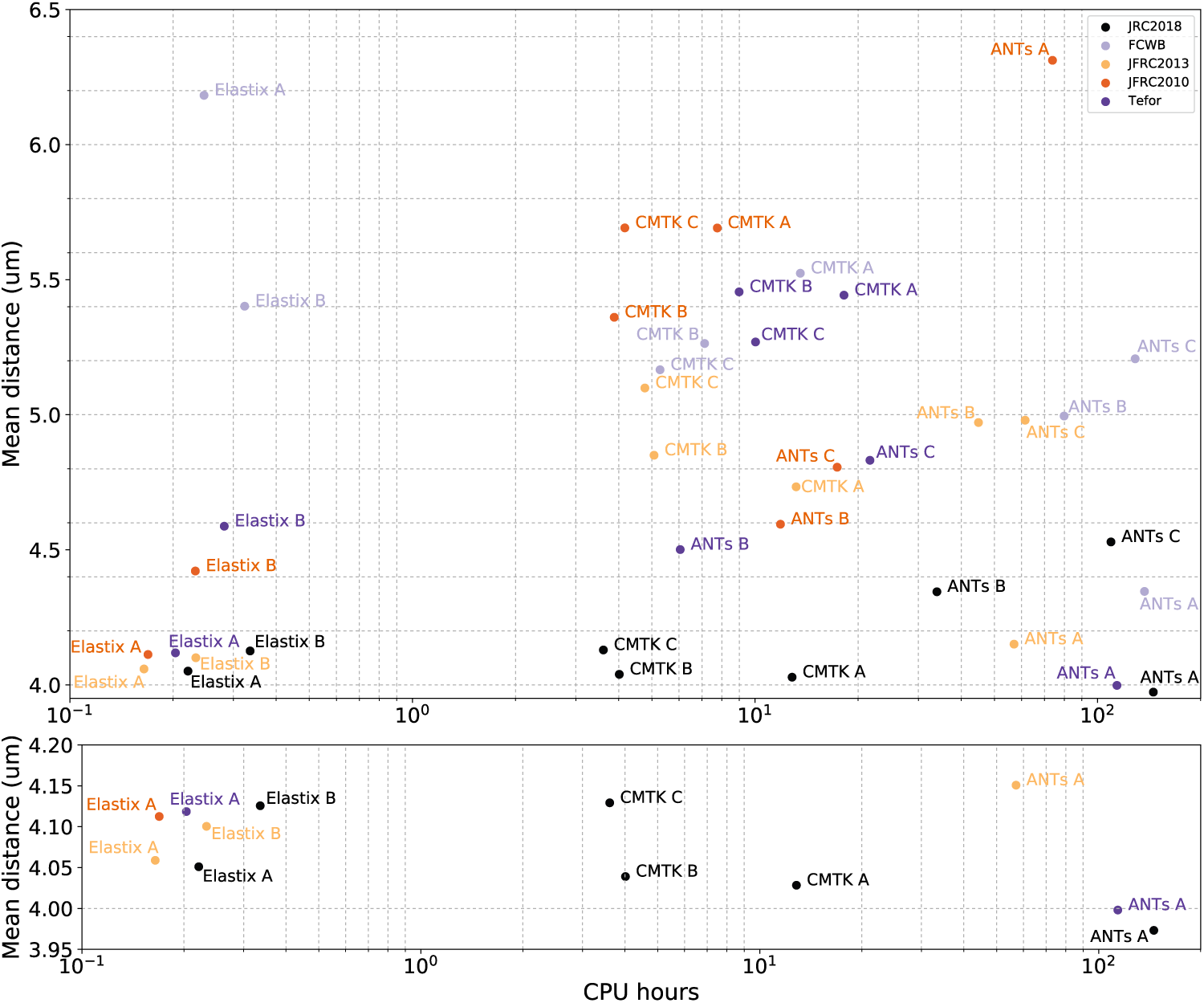
Scatterplot showing mean skeleton distance and the mean computation time in CPU-hours for all template-algorithm pairs (above), and the best performing pairs (below).

**Table 3:** Mean (Standard deviation) of skeleton distance by template, in µm. Templates are ordered by decreasing performance for each template’s best algorithm (given in the third column). Lower distances indicate better registration performance. The three rightmost columns show statistics of skeleton distance for all templates when fixing the algorithm, where we choose one set of parameters for each registration library.

| Template | Mean (standard deviation) skeleton distance ($\mu\text{m}$) | | | | |
| --- | --- | --- | --- | --- | --- |
|  | Best per template |  | Fixed algorithm |  |  |
|  |  | Algorithm | ANTs A | CMTK A | Elastix A |
| JRC2018F | 3.97 (3.65) | ANTs A | 3.97 (3.65) | 4.03 (3.70) | 4.05 (3.73) |
| Tefor | 4.00 (3.68) | ANTs A | 4.00 (3.68) | 5.44 (5.75) | 4.12 (3.69) |
| JFRC2013 | 4.06 (3.68) | Elastix A | 4.15 (3.84) | 4.73 (4.71) | 4.06 (3.68) |
| JFRC2010 | 4.11 (3.68) | Elastix A | 6.31 (6.20) | 5.69 (5.50) | 4.11 (3.68) |
| FCWB | 4.35 (3.97) | ANTs A | 4.35 (3.97) | 5.52 (4.97) | 6.18 (5.93) |

**Table 4:** Jacobian determinant standard deviation (JSD) and Hessian frobenius norm mean (HFM) where templates are sorted by descending JSD for a fixed algorithm (ANTs A). This algorithm has the capability of producing large deformations (i.e. it is not overly regularized), as by the large value of JSD for the FCWB template. The smallest average amount of deformation was obtained after registration to our template, JRC 2018F, followed closely by JFRC 2013.

| Template | JSD |  |  | HFM |  |  |
| --- | --- | --- | --- | --- | --- | --- |
|  | ANTs A | CMTK A | Elastix A | ANTs A | CMTK A | Elastix A |
| JFRC2010 | 0.26 | 0.19 | 0.18 | 0.032 | 0.014 | 0.0029 |
| JFRC2013 | 0.23 | 0.26 | 0.23 | 0.039 | 0.017 | 0.013 |
| JRC2018F | 0.20 | 0.21 | 0.17 | 0.035 | 0.025 | 0.018 |
| FCWB | 0.47 | 0.32 | 0.23 | 0.051 | 0.038 | 0.0047 |
| Tefor | 0.26 | 0.23 | 0.14 | 0.040 | 0.019 | 0.0033 |

**Table 5:** Summaries of algorithm performance, tabulating mean skeleton distance values across templates of the summary statistics given in Table 7, i.e., mean-of-means, mean-of-medians, and mean-of-ranks. Mean rank is computed by ranking the algorithms by mean skeleton distance when the template is fixed, then averaging over templates. Algorithms are sorted by their mean skeleton distance for the JRC 2018 template.

| Algorithm | Mean distance |  | Mean of medians | Mean rank |
| --- | --- | --- | --- | --- |
|  | JRC 2018 | All |  |  |
| ANTs A | 3.97 | 4.56 | 3.48 | 1.80 |
| Elastix A | 4.05 | 4.50 | 3.53 | 2.20 |
| Elastix B | 4.13 | 4.53 | 3.54 | 2.80 |
| ANTs B | 4.34 | 4.68 | 3.71 | 3.20 |
| CMTK A | 4.03 | 5.08 | 3.78 | 4.20 |
| CMTK B | 4.04 | 4.99 | 3.70 | 4.20 |
| ANTs C | 4.53 | 4.87 | 3.83 | 4.60 |
| CMTK C | 4.13 | 5.07 | 3.90 | 5.00 |

The statistics we report in this section are computed over the whole brain. Appendix B contains tables with additional statistics computed over subsets of the brain defined by 76 compartment labels.

#### 3.2.2 Deformation measure

In addition to measuring accuracy, we also report the standard deviation of the Jacobian determinant (JSD), a scalar measure that describes the deformation or distortion of a brain’s shape after undergoing a transformation. This is important if downstream analyses in template space rely on anatomical morphology, see Section 4.3 for a discussion. The Jacobian determinant of a transformation at a particular location describes the amount of local shrinking/stretching, where a value of 1.0 indicates that volume is locally preserved, less that 1.0 indicates volume decrease, and greater than 1.0 indicates volume increase. The standard deviation of the distribution of the Jacobian determinant map reflects deformation because the spread of this distribution (not the mean) captures the extent to which a transformation is *simultaneously* stretching and shrinking space. A similarity transformation will have a fixed value of the Jacobian determinant equal to its scale parameter at all points. As a result, its mean Jacobian determinant will equal that value. Therefore, the mean value of the Jacobian determinant is not indicative of a deformation, but rather shows average scaling. In the supplement, Table 8 shows that the *mean* value of of the Jacobian determinant is very nearly 1.0 for most templates, as we would expect. We computed the JSD from displacement fields and do not include the affine component to avoid confusing global scale changes with deformation. We consider an alternative measure of deformation in Section 3.2.3.

In Figure 5, we plot the JSD against the mean skeleton distance described above. We observed that algorithms with stronger regularization (CMTK C, ANTs B) have a lower JSD, as we would expect. The mean distance between skeletons is lowest for the JRC 2018F template. Using the best overall algorithm (ANTs A), the deformation when using JRC 2018F is lower than when using Tefor, the next best template. Among the fastest algorithms (Elastix A and Elastix B), the JSD for JRC 2018F is lower than that for JFRC 2013, but higher than Tefor and JFRC 2010.

#### 3.2.3 Another deformation measure

In Section 3.2.2, we explain that the standard deviation of the Jacobian determinant over space measures deformation. One reason to use that measure is that the registration libraries we tested here (ANTs, CMTK, elastix) provide functions to compute the Jacobian determinant. Despite its prevalent use, it may not always measure the kind of deformation researchers care to avoid.

For example, a transformation with regions of volume decrease and regions of volume increase would have a large JSD, but we might still want to call that transform “smooth” if those regions are spatially far from each other and the intermediate space varies smoothly from shrinking to stretching. On the other hand, it could be useful to describe transforms with less extreme values of the Jacobian determinant as “unsmooth” if changes from stretch to shrink occur over smaller spatial distances. In other words, it could be useful to describe *how quickly* (over space) transform changes occur, rather than describing *that* transform changes occur over all of space as JSD does. Next, we describe that the norm of the Hessian matrix could serve as a useful and complementary measure to the JSD.

The Hessian matrix is the matrix containing all partial second derivatives of a transformation, and describes how spatially quickly changes in the transformation occur. The Hessian of any linear transformation will be identically zero everywhere. Therefore the “size” of the matrix as measured by a matrix norm describes the extent to which a transformation is locally non-linear. The mean value of the Hessian matrix norm is an appropriate statistic for summarizing over space. We compare the Hessian matrix Frobenius norm mean (HFM) to the JSD in Section 3.2.3, and report both values in Table 4.

Table 4 shows the JSD and HFM for the five templates when using ANTs A as the registration algorithm (the best algorithm on average, measured by mean skeleton distance). This suggests that the JRC 2018F template may be “closer” on average to the shape of individual brains (in our testing cohort) in the sense that less deformation is required to transform those brains to the template. The conclusions for HFM are different, though. The JRC 2018 template is third smoothest on average according to HFM. This could be due to different tissue preparation methods used for evaluation brains and template building brains. The tissue preparation used for the JFRC 2010 and JFRC 2013 templates is more similar to the evaluation images than the preparation used for the images used to build our template. Table 8 in Appendix A.5 shows a complete listing of JSD and HFM statistics across templates and algorithms.

### 3.3 Computation time and supplementary results

Figure 6 plots mean skeleton distance against mean computation time (in CPU-hours). The choice of algorithm and parameters influences computation expense more than the choice of template. Our particular parameter choices for CMTK are generally faster than ANTs, and our parameter choices for elastix are about ten times faster still. We see that the best performing algorithm (ANTs A) is also the most computationally demanding. Finally, for the fastest algorithms (Elastix A and Elastix B) mean skeleton distance is smallest using the JRC 2018F template. JFRC 2013 is the next best template.

We also performed experiments varying the resolution of the two best templates JRC 2018F and Tefor (see Appendix A.4). The results show that a marked speed up in computation time can be achieved by registering downsampled images, for only a modest decrease in accuracy.

### 3.4 Sexual Dimorphism

Next, as a demonstration of efficacy, we briefly examine sexual dimorphism in the *Drosophila* brain, reproducing the results of Cachero et al. (2010), using the brain templates we created. Consistent with those previous findings, we observe enlargement of anatomy associated with *fruitless* gene expression in the male JRC 2018 template relative to the female JRC 2018 template.

We compared the Jacobian determinant maps of two transformations: the transformation between the male and unisex template and that between the female and unisex template. These transformations should reflect the differences between the mean female and male shapes, respectively. Figure 7 shows the differences of the Jacobian determinant map for the transformations between the male-to-unisex and female-to-unisex templates. This qualitatively agrees with the result in Cachero et al. (2010) that male enlarged regions correlate with *fruitless* gene expression.

**Figure 7:**
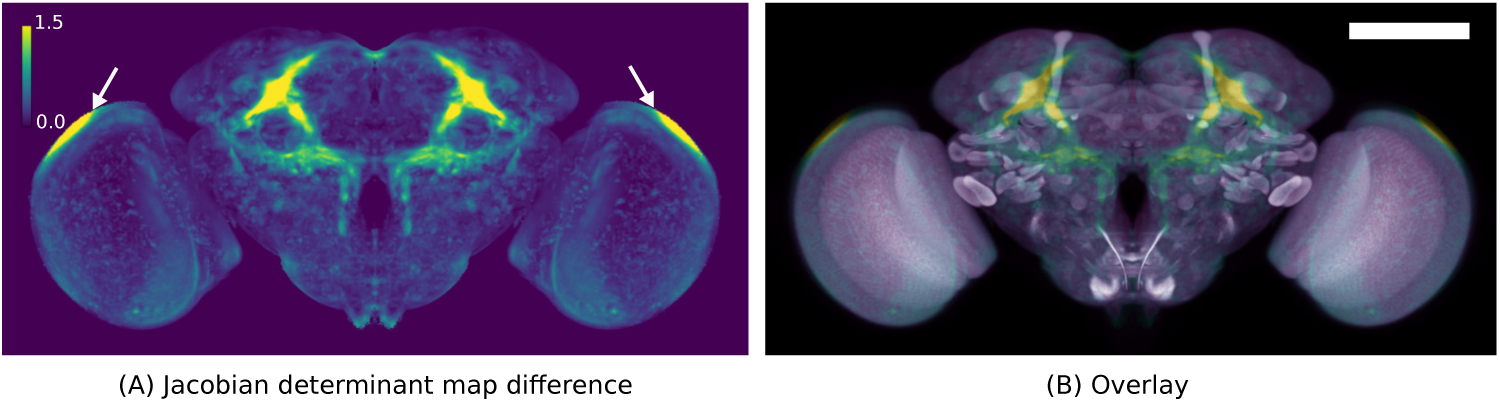
A visualization of female vs male morphological differences. (A) the maximum intensity projection (MIP) of the difference between male and female Jacobian determinant maps where values greater than 1 (yellow) indicate the male template is locally larger. (B) the MIP of the Jacobian determinant map overlayed with a MIP of the unisex template. Arrows indicate an artifact of this analysis in regions where the templates have no contrast, but the registration algorithm applies a strong deformation. Scale bar 100 µm.

## 4 Discussion

We created six *Drosophila* templates for female, male, and unisex central brains and ventral nerve cords. In Figure 1, we see that the unisex templates are generally more blurry than the single sex templates, likely because the unisex templates include inter-sex variation in addition to inter-individual variation. This is more pronounced for the VNC than for the central brain. Furthermore, male templates appear to be more blurry than the female templates both for the brain and VNC. This could be due to higher variability across male individuals, or poorer performance by registration algorithms for males, though we believe the former to be more likely.

It is difficult to determine the extent to which each aspect of template construction affects registration performance. Nevertheless, here we put our results in the context of other work attempting to estimate performance of various templates. In Figure 3, we see that the anatomical features in the *Drosophila* brain in our template are qualitatively more pronounced, having higher contrast than other existing templates. As a result, pixel-based similarity measures used by automatic registration algorithms may be more effective at optimization when the signal for the target image is less obscured by noise of different kinds (e.g., anatomical variability, imaging artifacts). This is potentially one of the reasons that the JRC 2018F template outperforms others.

### 4.1 Estimating registration quality

In this work we chose pairwise neuronal skeleton distance as the primary measure of registration performance. Most templates are not explicitly evaluated for registration accuracy at time of publication (Jenett et al., 2012; Aso et al., 2014), though Arganda-Carreras et al. (2018) use the overlap of anatomical labels to compare performance. Unlike reference image similarity, both skeleton distance and label overlap are relevant in the sense that these measures are not directly optimized for by the registration algorithm which can lead to meaningless results (Rohlfing, 2012). Both of these measures have advantages and drawbacks.

Label overlap is sensitive to errors in the spatial location of anatomical compartments that are (usually manually) labeled by human annotators. Therefore, it specifically focuses on regions that are of potential interest to researchers. However, human annotation includes arbitrary choices, and therefore this measure may miss errors in biologically relevant areas that annotators overlook. Furthermore, annotating many individual images can be costly and introduces the variability of manual labeling. On the other hand, having multiple individuals annotated is advantageous in that the anatomical labels themselves can also have uncertainty associated with them. Multiple labeled individuals enable multi-atlas segmentation (Iglesias and Sabuncu, 2015), an extension of single-atlas segmentation, at the cost of increased computational cost. Another potential drawback of using label overlap is that only changes near the boundaries of labels effects the overlap measure, and so it cannot differentiate methods that perform differently at the interior of labels. Registration errors parallel to or along the boundary are also not distinguishable.

The skeleton distance measure we use in this work evaluates registration using automatically extracted anatomy *directly* rather than indirectly as labeled by human annotators and so avoids manual effort in obtaining a registration accuracy measure. It instead requires that an additional independent image channel be acquired and must deal with biological variability and algorithmic errors as confounding factors. Specifically, näıve skeletonization and pairwise matching rely on the assumption that the nearest points on two skeletons are in correspondence, and will result in underestimates of distance. This also means that skeleton distance cannot differentiate errors parallel to the skeleton. Furthermore, any errors in the automatic segmentation of the skeletons will affect the distance computation. Biological variability could also be a significant limitation if this variability is much larger than the differences between templates/registration algorithms. In fact, we do observe a large variance in the distribution of skeleton distance for all templates and all algorithms. The errors we observe are consistent with estimates of registration accuracy / biological variability in previous studies (see Section 4.2). In this work, differences in the distributions of this measure between templates and algorithms were detectable, even in the presence of these sources of variability.

Currently, the JRC 2018 templates do not have anatomical labels of their own superimposed, except for those that are inherited implicitly through bridging transformations to other templates. As a result, the accuracy of these labels depends on the accuracy of bridging transformation. In future work, we plan to develop a set of anatomical labels in the space of the JRC 2018 templates.

### 4.2 Studies of registration accuracy and anatomical variability

In a previous study, Jefferis et al. (2007) measured biological variability and registration accuracy by measuring the branch point of a projection neuron (PN) after entering the lateral horn, and estimated spatial variability of 2.64, 1.80, and 2.81 µm, for the three axes. This yields a mean euclidean distance of about 4.3 µm. Separately, they found that the mean axon positions of PNs within the inner antennocerebral tract (iACT) were 3.4 µm apart. In another study Peng et al. (2011), developed a pointwise image registration method, “Brainaligner,” and assessed its accuracy, as well as natural biological variability. They estimated the spatial variability of the axons in the iACT to 3.26 µm, quite similar to the 3.4 µm estimate in Jefferis et al. (2007). These estimates, taken together are consistent with our estimate of a mean skeleton distance of about 4.0 µm for the best registration algorithm and the best template. This figure is slightly less than the 4.3 µm distance of the lateral horn PN branch point from and slightly greater than the 3.4 µm estimate of the iACT from Jefferis et al. (2007). The 3.26 µm estimate of Peng et al. (2011) is lower still. We expect part of this difference has to do with the fact that their measure focuses on a single, perhaps easily localizable region, whereas we examine distances across the entire brain.

### 4.3 Deformation

As observed in Figure 5, one of the benefits of the JRC 2018 template is that an accurate registration can be obtained with less local deformation than for other brain templates. This is beneficial because the morphology of anatomy is better preserved, and therefore the influence of the transformation is less likely to be a confounding factor in downstream analyses. For example, neuronal morphology is an important signal when searching across driver lines for a particular neuron or neurons of interest. This task benefits from normalizing spatial location (registration) and from comparing morphology. Therefore, minimizing deformation is beneficial for this task. For example, NBLAST (Costa et al., 2016) makes use of morphology of neurons in computing pairwise scores, and therefore any distortion in shape could prevent effective matching.

While the Jacobian- and Hessian-based measures of deformation are correlated, they do measure somewhat different properties of the transformation. The fact that some transformations with similar values of JSD can have markedly different values of HFM is evidence of this. Transformations produced by ANTs are less smooth than those produced by CMTK and elastix when measured by HFM.

The extent of deformation is just one of several factors to be considered when choosing a template and registration algorithm. A reasonable choice would be to accept larger deformations only if they produce more accurate registration results. A danger described in Rohlfing (2012) is that transformations with an unreasonably high degree of deformations can “trick” bad proxy measures of registration accuracy. Better measures of registration quality, such as the skeleton distance used here, can help to avoid this. For example, the ANTs C parameters were not regularized, and produced very unsmooth transforms, corroborated by high values of JSD in Figure 5. That algorithm also scored relatively poorly according to skeleton distance, which indicates that the measure is robust. This poor performance is visually apparent as well, as can be seen in the examples in Appendix C.

In the following sections, we consider a few possible reasons for the improvements in performance we see when using JRC 2018 as a target for registration.

### 4.4 Influence of groupwise averaging

A potential concern in using an average of many individuals as a registration target is that some anatomical features are blurred or lost due to imperfect registration followed by averaging. In this case, it could be that the loss or blurring of these anatomical features removes a useful signal for the registration algorithm and could result in worse alignment. If this is true, then individual templates should perform better than average templates, since features are not blurred away. Our results suggest the opposite, that groupwise-average templates outperform individual templates This agrees with other work on the human hippocampus from MRI (Avants et al., 2010), in ants (Arganda-Carreras et al., 2017), and *Drosophila* (Arganda-Carreras et al., 2018).

If an anatomical feature is blurred or lost in the process of template construction, then that feature must have been poorly aligned on average across the population, due perhaps to large anatomical variability. It could be that the presence of unreliable/ highly variable anatomy is useless or detrimental on average for registration across a large population. For example, highly variable anatomy could result in over-warping, especially when using algorithms with very flexible transformation models, since they can “force” improvements to the similarity measure even for incompatible anatomy, analogous to overfitting in machine learning. This is another possible reason that average templates outperform individual templates.

### 4.5 Symmetry

The central brain of *Drosophila* is largely left-right symmetric, but it does have a noteworthy asymmetry (Pascual et al., 2004). Any template created without enforcing symmetry would have small but widespread asymmetries caused by the particular specimens and images that contributed to the average, not *biologically meaningful* asymmetries. We therefore created a set of left-right symmetric templates. Of course, this has the disadvantage of removing *real* asymmetries from the template brain. For researchers interested in analyzing asymmetries, they can be recovered by measuring left-right differences in the deformations to a symmetric template using deformation based morphometry (Ashburner et al., 1998). An advantage of symmetry is that the estimation of a transform mapping the left hemisphere to the right and vice-versa (“a mirroring transformation”) is straightforward. Manton et al. (2014) and Schlegel et al. (2017), for example, describe how such mirroring transforms are useful in performing neuronal comparisons across hemispheres, since many neurons in one hemisphere correspond to a partner in the other.

### 4.6 Influence of number of individuals to build atlas

Our templates were built using many more individuals than other average templates. The FCWB template averaged 26 individuals, and the Tefor brain averaged 10 individuals, where we average 36 female individuals, and 26 male individuals for the sex-specific but consider twice as many images by left-right flipping each individual. As a result, we are more confident that our templates are near the “mean shape” of the central brain for each sex. Arganda-Carreras et al. (2018) explored how the number of individuals that comprised a template effects the registration quality to those templates. They found a plateau of performance for templates built using between seven and ten individuals. Yet, JRC 2018 generally outperforms Tefor. Next, we outline some of the potential reasons for this.

If the conclusion of Arganda-Carreras et al. (2018) is true, then whatever performance improvement JRC 2018 achieves over the Tefor brain is not due to the number of individuals comprising the templates. We note first that the mean distance measures for JRC 2018 and Tefor are very similar when using the best performing but most costly registration algorithm (ANTs A). Using this algorithm, the average deformation energy (measured by Jacobian determinant) is smaller for JRC 2018 (0.17) than for Tefor (0.23) (see Figure 5), suggesting that it is “closer” to the mean of our test subjects than the Tefor brain. This could also explain why JRC 2018 achieves better performance than Tefor when using registration methods that are more highly regularized (e.g. ANTs B and CMTK C). It could be the case that the template shape does not converge after only seven to ten individuals have been averaged, but that this shape change is not reflected in registration performance measures. It seems unlikely that image acquisition would affect the average deformation when registering to a template. Given that, our results support the conclusion that using more individuals for a template yields a more representative shape which reduces deformations for a given level of performance, or enables better performance for a smaller, fixed amount of deformation.

Finally, we note that one of the most widely used anatomical templates, the Allen Mouse Common Coordinate Framework, used many more individuals (1675) than we used in this work, and, as we did, also included left-right flips as well as the original images for a total of 3350 (Allen Institute for Brain Science, 2015).

### 4.7 Influence of tissue staining and preparation

Arganda-Carreras et al. (2018) compare their Tefor template to the FCWB template (Ostrovsky and Jefferis, 2014) and show that Tefor has improved performance, a conclusion that our results corroborate. They suggest that the difference between tissue staining protocols could be a contributing factor in the observed performance improvement. Nc82 images were used both to build the Tefor brain and for testing in Arganda-Carreras et al. (2018). In our work, we also use nc82 images for testing, but instead use brp-SNAP tag for images that contributed to our template. Still, evidence suggests that nc82 images are more reliably registered to our template than to Tefor, despite the fact that images contributing to our template underwent different sample preparation and imaging, and that similar methodology (groupwise registration) was used to construct our template and Tefor. The SNAP labeling results in more even contrast throughout the brain. Our results suggest that a template built with images having differing but improved contrast is a better registration target than a template of the same modality but worse contrast.

### 4.8 Influence of template resolution

The resolutions of a brain template could affect the ability of registration algorithms to produce an accurate alignment. It could be that templates rendered at isotropic resolution could outperform those with anisotropic voxels even when the moving image is anisotropic, as is the case with our testing data that were acquired with a confocal microscope. One of the anisotropic templates we evaluated here performs well in the best case (Tefor), while the other (FCWB) performs worse than isotropic templates generally.

We also explored how varying the resolution of the two best templates (JRC 2018 and Tefor) affects registration performance (see Appendix A.4). We conclude that it is possible for lower resolution templates to achieve similar performance to high resolution templates with the benefit of computational savings. However, it is worth exploring different choices of algorithms in this case, since any given algorithm may not have uniform performance across different resolutions.

### 4.9 Accuracy/time tradeoff

The required level of accuracy of registration will vary from task to task. We recommend that researchers first experiment with the less computationally demanding algorithms/parameter sets, determine whether the results are adequate for their particular task, and if not, to try the potentially more accurate but more computationally demanding options. Specifically, among the parameter sets we tested, elastix is about ten times faster than the fastest parameter settings for CMTK and ANTs, and is therefore worth experimenting with first.

As a result, Figure 6 shows a cluster of templates with good performance that are fast to compute when using elastix. JRC 2018 is the best performing template in this cluster. Furthermore, the performance is also qualitatively much better using JRC 2018 than others when examining the registration results. Many examples of registration results are provided in Appendix C.

## Acknowledgements

The authors would like to thank Philipp Hanslovsky, Igor Pisarev, Alice Robie, Yoshi Aso, Greg Jefferis, and Michael Reiser for helpful discussions, Zhihao Zheng for help with EM-LM registration analysis, Arnim Jenett and Ignacio Arganda-Carreras for the Tefor brain template. This work was supported by the Howard Hughes Medical Institute.

## A Supplementary Notes

### A.1 Reference image overlay

Figure 8 shows two examples of test images (randomly selected from the 20 test images) aligned to four different templates using the “best” algorithm (ANTs A above). Some differences are visible upon close inspection. For example, the right mushroom body is poorly aligned for Fly B using the FCWB template. Small misalignments near the mushroom body are evident for both flies after registration to JFRC 2013. Otherwise, alignment appears generally good.

### A.2 Algorithm comparison

Table 5 compares the eight different registration algorithms we tested by tabulating statistics of the skeleton distance described above in Section 3.1. ANTs A performed best overall, though the two parameter settings of elastix were similarly good according to these measures. The two worst performing algorithms were ANTs C, the least smooth parameter settings among ANTs algorithms, and CMTK C the smoothest of the CMTK algorithms.

**Figure 8:**
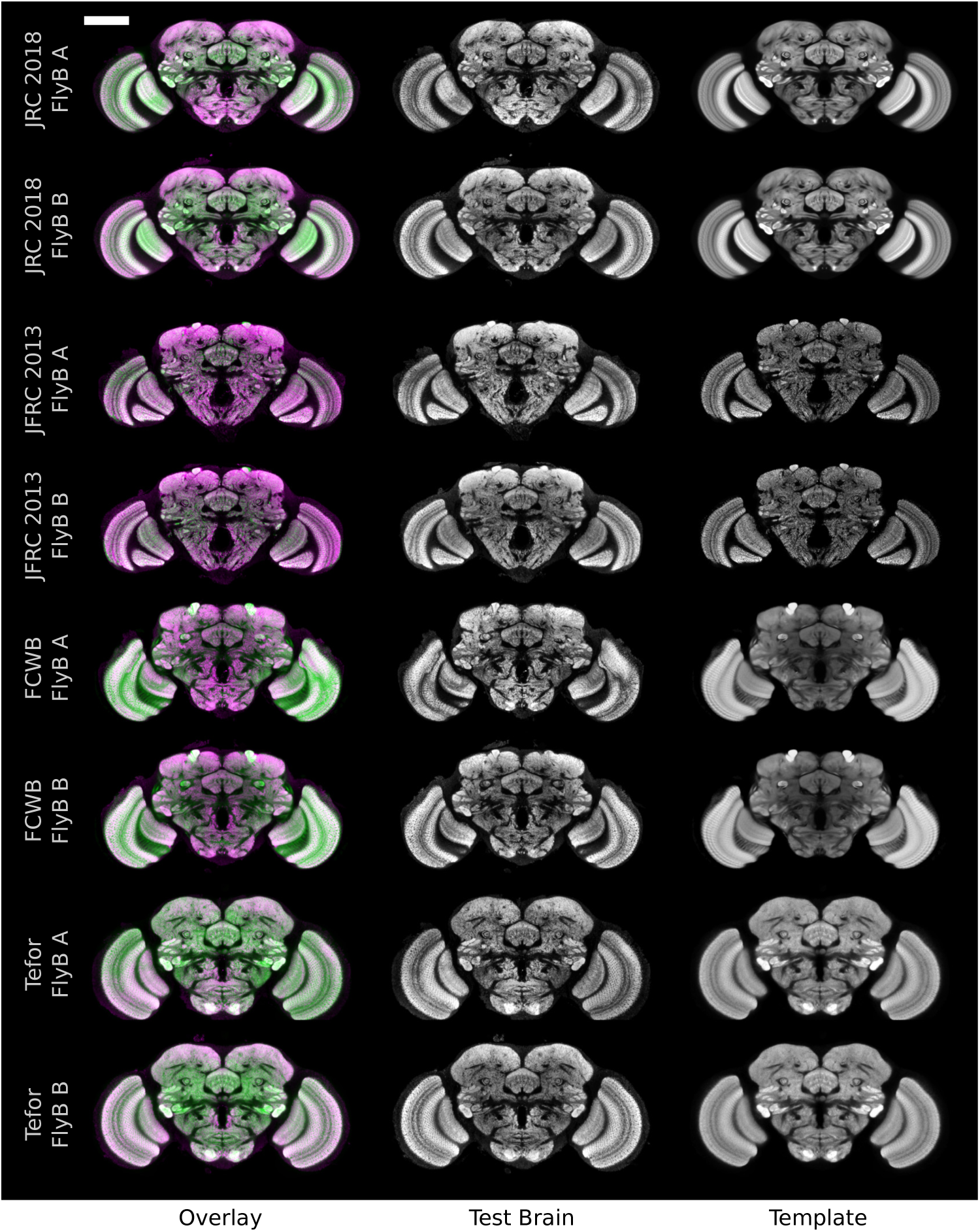
Overlays of the reference (nc82) channel. Scale bar 100 µm.

**Figure 9:**
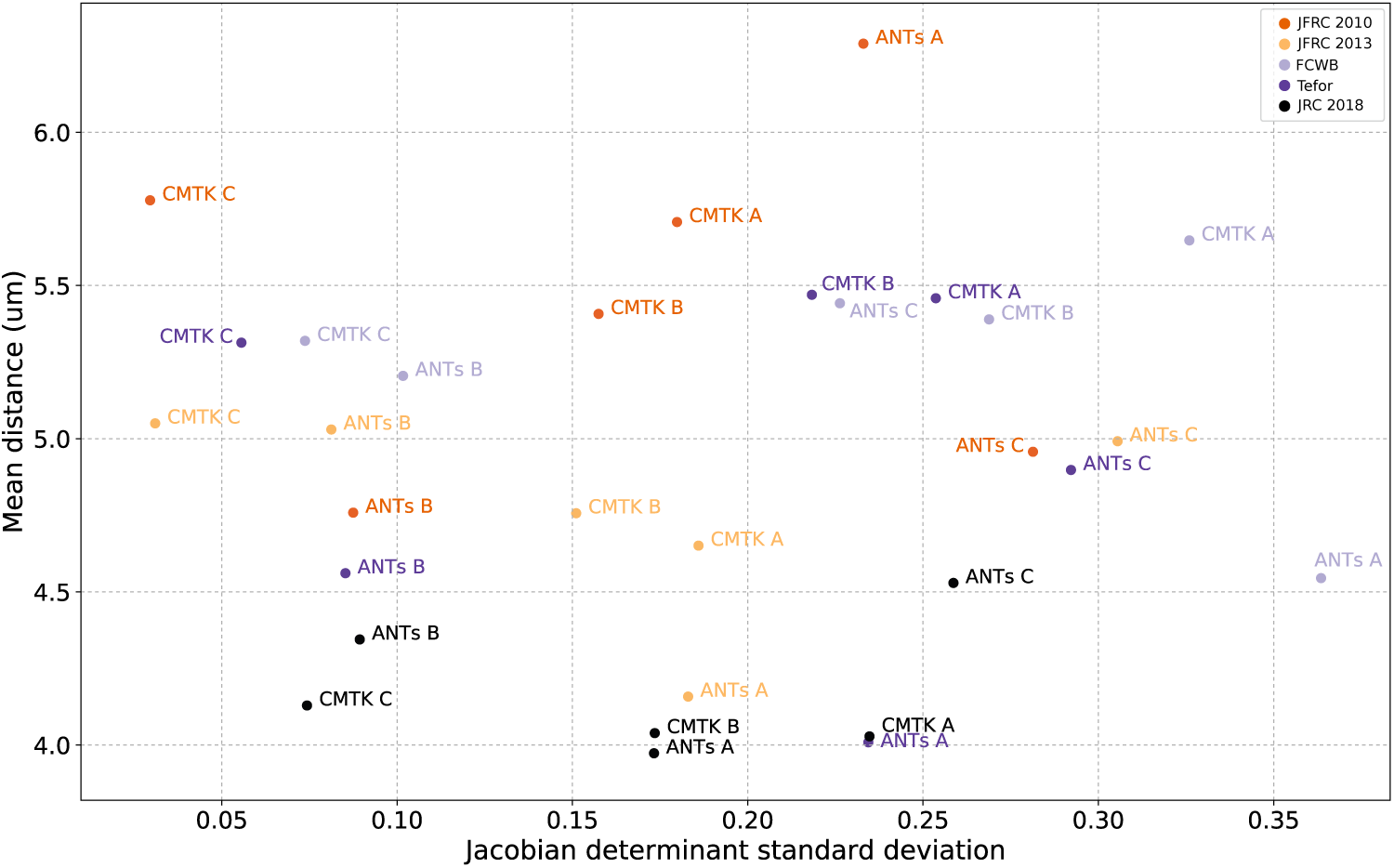
Scatterplot showing mean skeleton distance and standard deviation of Jacobian transform map for template-algorithm pairs using affine-only normalization.

### A.3 Size and shape normalization for evaluation

Figure 9 shows the results of mean skeleton distance and Jacobian determinant standard deviation (JSD) after normalization by using only the affine part of the bridging transformation. This is in contrast to Figure 5, in which both the affine and deformable parts of the transformation contributed to normalization. We observed that results are generally very similar. The largest difference we observe is that the distance measures for the FCWB template are notably larger (worse performance) when using affine-only normalization.

### A.4 Effect of resolution

We performed experiments to explore the effect that the resolution of a template has on the tradeoff between registration quality and computational cost. We repeated the evaluation described in section Section 3.2 using downsampled versions of the two best performing templates: JRC 2018 and Tefor. We resampled each of these two templates from its native resolution to three lower, isotropic resolutions: 1.2, 2.4, and 3.6 µm. These resolutions correspond to downsampling in *xy* by factors of approximately 2, 3, and 4. Figure 10 shows a scatterplot of mean skeleton distance (our measure of registration accuracy) against mean CPU time for template-algorithm pairs at various resolutions. The same results are listed in Table 6.

**Figure 10:**
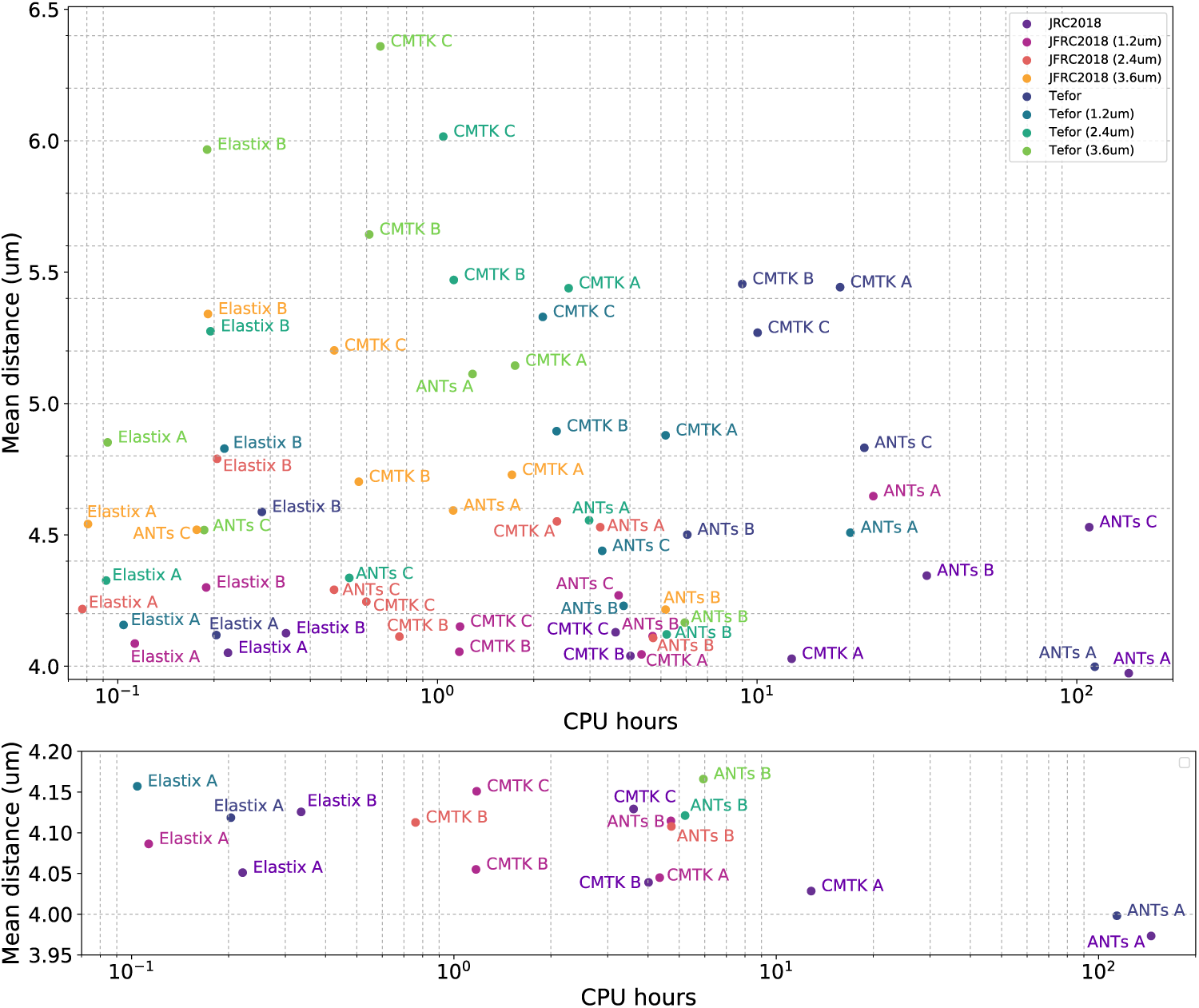
Scatterplot showing mean skeleton distance and the mean computation time in CPU-hours for the JRC 2018 and Tefor templates at various resolutions (above), and the best performing pairs (below).

We observed that some downsampled templates perform as well or nearly as well as the full-resolution templates. For instance, JRC 2018, sampled at 1.2 µm performs competitively with the full-resolution template using CMTK A, CMTK B, and Elastix A. Downsampled versions of Tefor seem to suffer somewhat more in terms of performance, though are competitive when using Elastix A or ANTs B. Detailed exploration of algorithm parameters will influence the time-performance tradeoff, and may be worthwhile for some tasks. Here we give values for reasonable ranges of parameters from faster and more highly regularized to slower with potentially greater accuracy.

More computationally expensive algorithms tend to enjoy larger speedups when operating on downsampled images. For example, using JRC 2018, ANTs A (the slowest) speeds up by six times on downsampled by two images relative to full resolution, whereas Elastix A (the fastest), speeds up by two times.

Furthermore, more highly regularized ANTs parameter settings may perform better on very highly downsampled images. For example, ANTs B is the best performing algorithm on the most downsampled versions of both templates. This does not seem to be the case for CMTK, since CMTK C performs relatively poorly for the most downsampled template versions.

**Table 6:** Mean skeleton distances and mean CPU time by template and algorithm at different resolutions.

| Template | Algorithm | Mean Distance (um) | Mean CPU time (hrs) |
| --- | --- | --- | --- |
| JRC2018 | ANTs A | 3.97 | 145.48 |
|  | ANTs B | 4.34 | 33.96 |
|  | ANTs C | 4.53 | 109.43 |
|  | CMTK A | 4.03 | 12.83 |
|  | CMTK B | 4.04 | 4.02 |
|  | CMTK C | 4.13 | 3.61 |
|  | Elastix A | 4.05 | 0.22 |
|  | Elastix B | 4.13 | 0.34 |
| JFRC2018 (1.2um) | ANTs A | 4.65 | 23.10 |
|  | ANTs B | 4.11 | 4.71 |
|  | ANTs C | 4.27 | 3.69 |
|  | CMTK A | 4.04 | 4.35 |
|  | CMTK B | 4.06 | 1.17 |
|  | CMTK C | 4.15 | 1.18 |
|  | Elastix A | 4.09 | 0.11 |
|  | Elastix B | 4.30 | 0.19 |
| JFRC2018 (2.4um) | ANTs A | 4.53 | 3.23 |
|  | ANTs B | 4.11 | 4.73 |
|  | ANTs C | 4.29 | 0.47 |
|  | CMTK A | 4.55 | 2.36 |
|  | CMTK B | 4.11 | 0.76 |
|  | CMTK C | 4.25 | 0.60 |
|  | Elastix A | 4.22 | 0.08 |
|  | Elastix B | 4.79 | 0.20 |
| JFRC2018 (3.6um) | ANTs A | 4.59 | 1.12 |
|  | ANTs B | 4.22 | 5.17 |
|  | ANTs C | 4.52 | 0.18 |
|  | CMTK A | 4.73 | 1.71 |
|  | CMTK B | 4.70 | 0.57 |
|  | CMTK C | 5.20 | 0.48 |
|  | Elastix A | 4.54 | 0.08 |
|  | Elastix B | 5.34 | 0.19 |
| Tefor | ANTs A | 4.00 | 113.92 |
|  | ANTs B | 4.50 | 6.05 |
|  | ANTs C | 4.83 | 21.65 |
|  | CMTK A | 5.44 | 18.19 |
|  | CMTK B | 5.45 | 8.99 |
|  | CMTK C | 5.27 | 10.04 |
|  | Elastix A | 4.12 | 0.20 |
|  | Elastix B | 4.59 | 0.28 |
| Tefor (1.2um) | ANTs A | 4.51 | 19.58 |
|  | ANTs B | 4.23 | 3.82 |
|  | ANTs C | 4.44 | 3.28 |
|  | CMTK A | 4.88 | 5.17 |
|  | CMTK B | 4.89 | 2.36 |
|  | CMTK C | 5.33 | 2.14 |
|  | Elastix A | 4.16 | 0.10 |
|  | Elastix B | 4.83 | 0.22 |
| Tefor (2.4um) | ANTs A | 4.56 | 2.98 |
|  | ANTs B | 4.12 | 5.21 |
|  | ANTs C | 4.34 | 0.53 |
|  | CMTK A | 5.44 | 2.57 |
|  | CMTK B | 5.47 | 1.12 |
|  | CMTK C | 6.02 | 1.04 |
|  | Elastix A | 4.33 | 0.09 |
|  | Elastix B | 5.27 | 0.19 |
| Tefor (3.6um) | ANTs A | 5.11 | 1.29 |
|  | ANTs B | 4.17 | 5.94 |
|  | ANTs C | 4.52 | 0.19 |
|  | CMTK A | 5.14 | 1.75 |
|  | CMTK B | 5.64 | 0.61 |
|  | CMTK C | 6.36 | 0.66 |
|  | Elastix A | 4.85 | 0.09 |
|  | Elastix B | 5.97 | 0.19 |

**Table 7:** Skeleton distance statistics (in µm) for all algorithm/template pairs, sorted by mean distance.

| Template | Algorithm | Mean | Std dev | Median |
| --- | --- | --- | --- | --- |
| JRC2018 | ANTs A | 3.97 | 3.65 | 3.08 |
| Tefor | ANTs A | 4.00 | 3.68 | 3.16 |
| JRC2018 | CMTK A | 4.03 | 3.70 | 3.16 |
| JRC2018 | CMTK B | 4.04 | 3.70 | 3.16 |
| JRC2018 | Elastix A | 4.05 | 3.73 | 3.24 |
| JFRC2013 | Elastix A | 4.06 | 3.68 | 3.24 |
| JFRC2013 | Elastix B | 4.10 | 3.73 | 3.24 |
| JFRC2010 | Elastix A | 4.11 | 3.68 | 3.24 |
| Tefor | Elastix A | 4.12 | 3.69 | 3.24 |
| JRC2018 | Elastix B | 4.13 | 3.78 | 3.24 |
| JRC2018 | CMTK C | 4.13 | 3.74 | 3.24 |
| JFRC2013 | ANTs A | 4.15 | 3.84 | 3.24 |
| JRC2018 | ANTs B | 4.34 | 3.94 | 3.46 |
| FCWB | ANTs A | 4.35 | 3.97 | 3.46 |
| JFRC2010 | Elastix B | 4.42 | 3.95 | 3.54 |
| Tefor | ANTs B | 4.50 | 4.02 | 3.61 |
| JRC2018 | ANTs C | 4.53 | 4.07 | 3.61 |
| Tefor | Elastix B | 4.59 | 4.13 | 3.61 |
| JFRC2010 | ANTs B | 4.59 | 4.06 | 3.61 |
| JFRC2013 | CMTK A | 4.73 | 4.71 | 3.61 |
| JFRC2010 | ANTs C | 4.81 | 4.15 | 3.81 |
| Tefor | ANTs C | 4.83 | 4.32 | 3.81 |
| JFRC2013 | CMTK B | 4.85 | 4.89 | 3.61 |
| JFRC2013 | ANTs B | 4.97 | 4.40 | 3.81 |
| JFRC2013 | ANTs C | 4.98 | 4.41 | 3.81 |
| FCWB | ANTs B | 5.00 | 4.26 | 4.06 |
| JFRC2013 | CMTK C | 5.10 | 4.23 | 4.12 |
| FCWB | CMTK C | 5.17 | 4.54 | 4.06 |
| FCWB | ANTs C | 5.21 | 4.41 | 4.12 |
| FCWB | CMTK B | 5.26 | 4.75 | 4.06 |
| Tefor | CMTK C | 5.27 | 5.44 | 3.81 |
| JFRC2010 | CMTK B | 5.36 | 5.13 | 3.87 |
| FCWB | Elastix B | 5.40 | 5.59 | 4.06 |
| Tefor | CMTK A | 5.44 | 5.75 | 3.81 |
| Tefor | CMTK B | 5.45 | 5.74 | 3.81 |
| FCWB | CMTK A | 5.52 | 4.97 | 4.18 |
| JFRC2010 | CMTK A | 5.69 | 5.50 | 4.12 |
| JFRC2010 | CMTK C | 5.69 | 5.28 | 4.24 |
| FCWB | Elastix A | 6.18 | 5.93 | 4.69 |
| JFRC2010 | ANTs A | 6.31 | 6.20 | 4.47 |

### A.5 Deformation measure

Table 8 shows the mean and standard deviations of the distributions of the Jacobian determinants for all templates and algorithms. As expected, the means for all methods are generally close to 1.0, and the standard deviations are smaller for algorithms with stronger regularization (ANTs B and CMTK C). This is in line with our reasoning above, suggesting that the standard deviation of the Jacobian determinant maps is a more informative statistic than the mean.

The mean values of the Jacobian determinant for JFRC 2013 appear to be significantly different both from 1.0 and the other templates. This could be due to a difference in size of that individual template and the cohort of individuals that we tested with, though the affine part of the transformation should account for gross size differences, and this statistic is computed from the deformation only, making it difficult to reach a definitive conclusion.

**Table 8:** Mean (Standard deviation) of Jacobian determinant standard deviation and Hessian frobenius norm mean by template. Observe that all mean values are close to 1.0, whereas standard deviations vary much more across template/algorithm pairs. For instance, ANTs B and CMTK C have consistently smaller standard deviations because they are more regularized.

| Template | Algorithm | Jacobian determinant<br>Mean (std dev) | Hessian frobenius norm<br>Mean (std dev) |
| --- | --- | --- | --- |
| JFRC2013 | ANTs A | 0.98 (0.23) | 0.039 (0.033) |
| JFRC2013 | ANTs B | 0.98 (0.11) | 0.013 (0.013) |
| JFRC2013 | ANTs C | 0.99 (0.32) | 0.062 (0.058) |
| JFRC2013 | CMTK A | 1.02 (0.26) | 0.017 (0.016) |
| JFRC2013 | CMTK B | 1.01 (0.23) | 0.013 (0.013) |
| JFRC2013 | CMTK C | 1.00 (0.09) | 0.0018 (0.0016) |
| JFRC2013 | Elastix A | 1.01 (0.23) | 0.013 (0.013) |
| JFRC2013 | Elastix B | 1.01 (0.23) | 0.013 (0.013) |
| JFRC2010 | ANTs A | 0.99 (0.26) | 0.032 (0.028) |
| JFRC2010 | ANTs B | 0.97 (0.12) | 0.0124 (0.011) |
| JFRC2010 | ANTs C | 0.95 (0.30) | 0.049 (0.045) |
| JFRC2010 | CMTK A | 1.03 (0.19) | 0.014 (0.015) |
| JFRC2010 | CMTK B | 1.04 (0.17) | 0.013 (0.014) |
| JFRC2010 | CMTK C | 1.01 (0.05) | 0.0019 (0.0021) |
| JFRC2010 | Elastix A | 1.02 (0.15) | 0.0029 (0.0023) |
| JFRC2010 | Elastix B | 1.03 (0.29) | 0.011 (0.0083) |
| Tefor | ANTs A | 1.01 (0.26) | 0.041 (0.034) |
| Tefor | ANTs B | 0.97 (0.11) | 0.023 (0.020) |
| Tefor | ANTs C | 0.97 (0.31) | 0.053 (0.050) |
| Tefor | CMTK A | 1.04 (0.23) | 0.019 (0.019) |
| Tefor | CMTK B | 1.03 (0.21) | 0.016 (0.017) |
| Tefor | CMTK C | 1.03 (0.08) | 0.0039 (0.0050) |
| Tefor | Elastix A | 1.02 (0.14) | 0.0033 (0.0024) |
| Tefor | Elastix B | 1.02 (0.30) | 0.013 (0.010) |
| JRC2018 | ANTs A | 1.01 (0.21) | 0.035 (0.029) |
| JRC2018 | ANTs B | 0.98 (0.11) | 0.017 (0.015) |
| JRC2018 | ANTs C | 0.98 (0.27) | 0.047 (0.042) |
| JRC2018 | CMTK A | 1.03 (0.21) | 0.025 (0.023) |
| JRC2018 | CMTK B | 1.03 (0.17) | 0.018 (0.017) |
| JRC2018 | CMTK C | 1.02 (0.10) | 0.0068 (0.0069) |
| JRC2018 | Elastix A | 1.03 (0.17) | 0.018 (0.017) |
| JRC2018 | Elastix B | 1.03 (0.17) | 0.018 (0.017) |
| FCWB | ANTs A | 1.00 (0.47) | 0.051 (0.054) |
| FCWB | ANTs B | 0.98 (0.13) | 0.0267 (0.023) |
| FCWB | ANTs C | 0.97 (0.24) | 0.056 (0.055) |
| FCWB | CMTK A | 1.06 (0.32) | 0.038 (0.035) |
| FCWB | CMTK B | 1.05 (0.28) | 0.033 (0.031) |
| FCWB | CMTK C | 1.01 (0.09) | 0.0097 (0.012) |
| FCWB | Elastix A | 1.08 (0.23) | 0.0047 (0.0035) |
| FCWB | Elastix B | 1.11 (0.39) | 0.014 (0.011) |

### A.6 Compare Jacobian and Hessian measures

In Section 3.2.2, we used the standard deviation of the Jacobian determinant over space to measure deformation. One reason for this is that the registration libraries we tested here (ANTs, CMTK, elastix) provide functions to compute that quantity. In the sections above, we explained why the standard deviation of the distribution of of the Jacobian determinant (JSD) is a sensible measure of deformation. To reiterate, JSD quantifies the extent to which a transformation *simultaneously* stretches and shrinks space.

Despite its prevalent use, it may not always measure the kind of deformation researchers care to avoid, because there are transformations with large values of JSD that may not reflect deformation in the intuitive sense. In Section 3.2.3, we describe how another measure, the Hessian matrix Frobenius norm mean (HFM), could be useful. Here we compare the HFM to the JSD.

Figure 11 plots JSD against HFM. As we would expect, these measure are positively correlated (correlation coefficient 0.63), but we observe some systematic patterns. Most interestingly, transformations estimated with ANTs tend to be less smooth when measured by HFM than transformations from either elastix or CMTK, even when the measures of JSD are the same or similar. This could be because the ANTs parameter sets we tested generally produce diffeomorphisms (ANTs A, ANTs C), whereas elastix and CMTK parametrize the transformations using splines.

Figure 12 is similar to Figure 5, but plots the mean skeleton distance (our proxy for registration accuracy) against the Hessian based smoothness measure (HFM) rather than the Jacobian based measure (JSD).

### A.7 Inter-modality registration

Zheng et al. (2017) present a complete electron microscopy (EM) volume of the *Drosophila* brain (FAFB), including a manual registration between the EM volume and a LM brain template. The transformation described therein used a Fiji (Schindelin et al., 2012) plugin called ELM (https://github.com/saalfeldlab/elm) based on BigWarp (Bogovic et al., 2016) to manually place landmark point correspondences between FAFB and JFRC 2013. While this approach can yield good results, it is limited in the following ways. Humans are limited to exploring the 3D EM data in one (arbitrary, i.e. not necessarily axis-aligned) 2D cross section at a time, and need to infer 3D structure from these views. Landmark placement can be difficult where the primary structures are planes, lines or curves rather than points. Furthermore, the EM volume is very large, and it is time consuming for a human to even look at the entire volume at a fine scale, and therefore accuracy of the transformation is likely to vary significantly over space. An automatically generated registration between these two modalities would improve upon some of these shortcomings.

**Figure 11:**
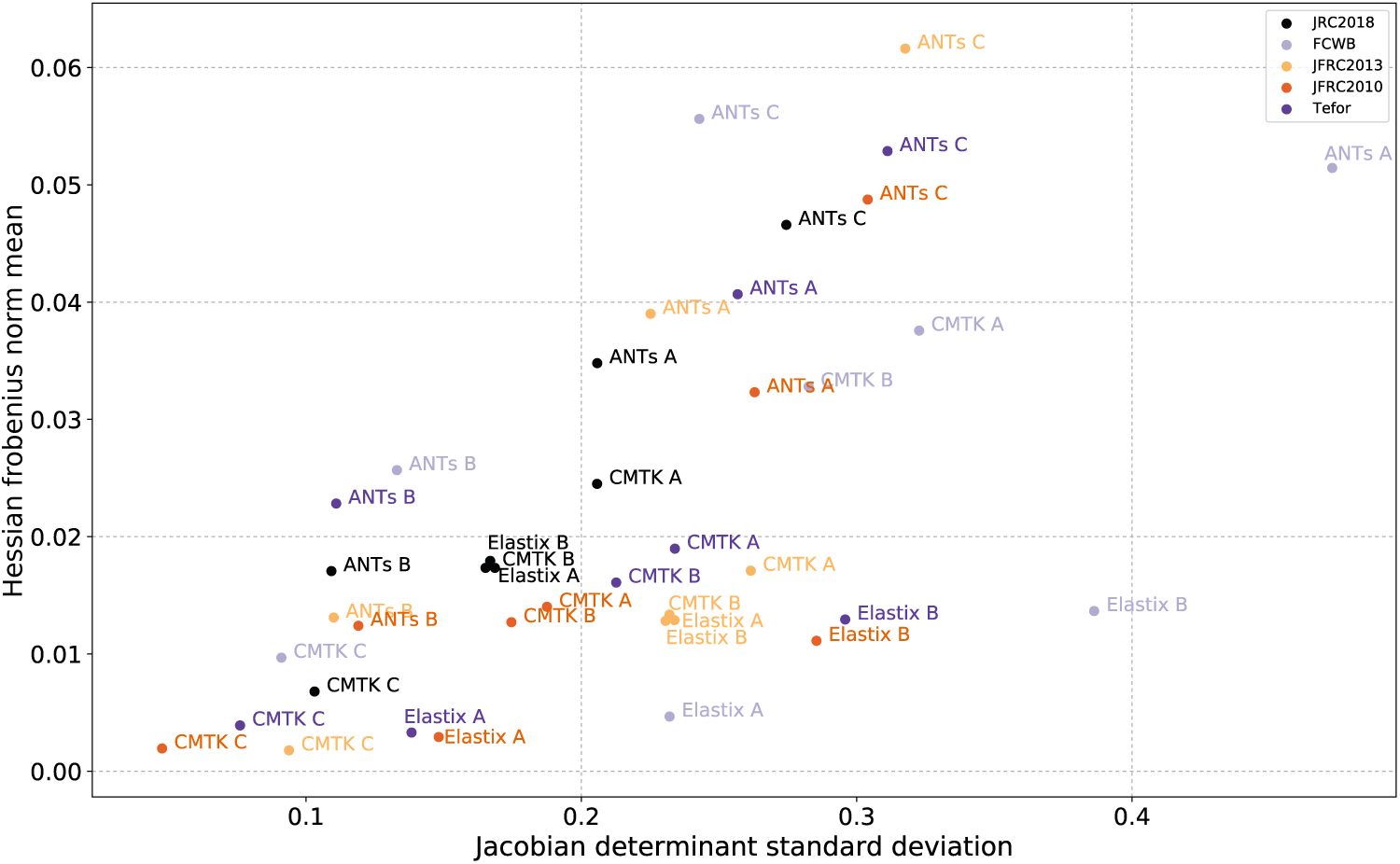
Scatterplot showing the standard deviation of the Jacobian determinant (JSD) and the mean of the Hessian’s Frobenius norm for template-algorithm pairs.

**Figure 12:**
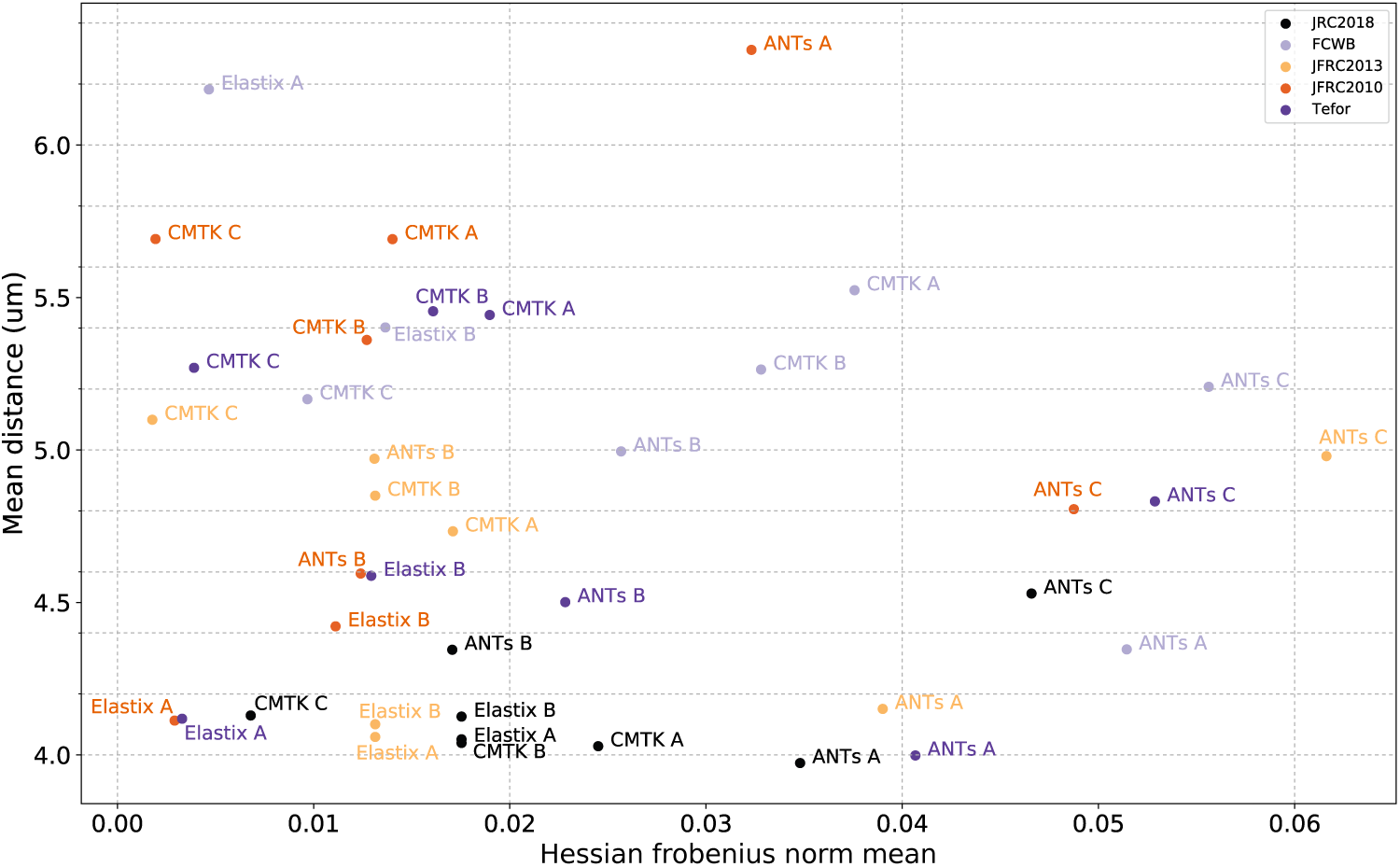
Scatterplot showing mean skeleton distance and the mean of the Hessian’s Frobenius norm for template-algorithm pairs.

In this work, we created a new, automatic alignment between that FAFB and the JRC 2018F brain template. The primary challenge is that the image contrast from EM and LM are at vastly different scales and measure different tissue properties. Typical approaches for inter-modality registration usually rely on choosing a similarity measure such as mutual information that works well across modalities, but this is highly unlikely to produce adequate results given the differences of scale and contrast. Our approach is to use an image derived from the EM instead of the EM image directly. Since brp-SNAP tag produces contrast at synapses, we created an image of synapses resembling our chemical tag stains (or nc82 images) using the automated synapse predictions by Heinrich et al. (2018), applied to FAFB. The bottom row of Figure 3 shows that the image of synapses estimated from EM does indeed resemble an LM template brain.

We reproduce the analyses by Zheng et al. (2017) that lead to figures 5B, 5C, S7B, and S7C in that work in order to demonstrate that our proposed brain template and this EM to LM transformation will be useful in this domain. The conclusions drawn by Zheng et al. (2017) are that projection neurons (PNs) arising from the same glomerulus cluster more tightly in the calyx of the mushroom body than previous observed using light microscopy.

One set of neurons was manually traced in FAFB by Zheng et al. (2017), while the other set came from the Fly Circuit (Chiang et al., 2011) database of neurons from light microscopy, pooling over several individual fly brains that have been co-registered to the FCWB template, as described by Costa et al. (2016). The neurons traced from LM were transformed to the space of JFRC 2010 using bridging registrations from Manton et al. (2014). The neurons traced from EM went through two transformations: the first from FAFB to JFRC 2013 generated manually as described above. The second was the automatically estimated transformation between JFRC 2013 and FCWB (Manton et al., 2014).

**Figure 13:**
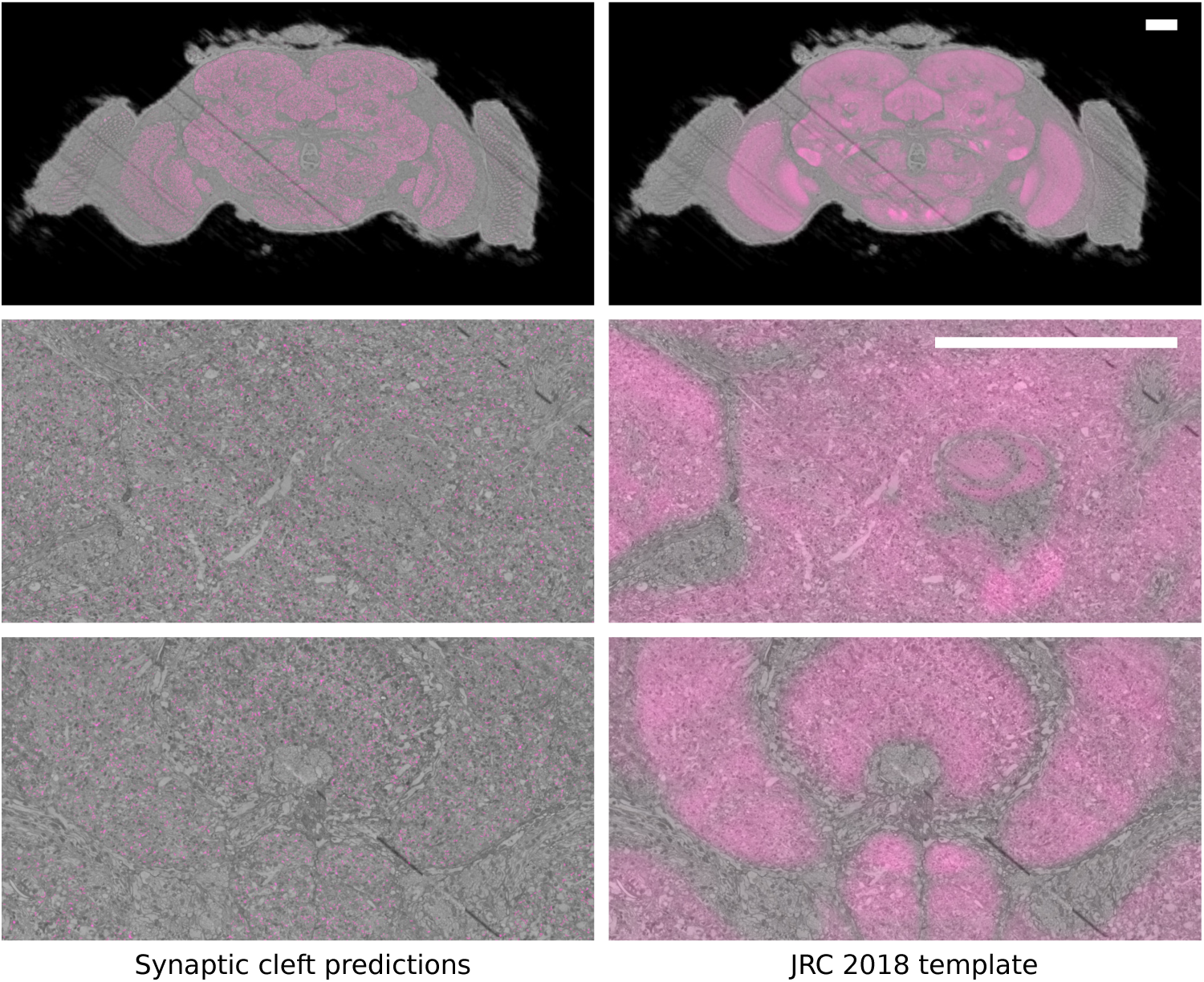
A visualization of the registration between JRC 2018F (magenta) and the FAFB electron microscopy volume (gray). The left column shows synaptic cleft predictions (magenta). The right column shows the resulting registration. Scale bar 50 µm.

The details of the analysis can be found in Zheng et al. (2017), but briefly, it consists of identifying projection neurons (PNs) in the calyx and grouping these by the glomerulus to which they project, then computing similarity/distance measures between pairs of PNs in the same group. Two related similarity scores between pairs of neurons were computed: the NBLAST score (Costa et al., 2016) and the mean Euclidean distance (NBLAST uses distance as part of its computation of score).

We repeated this analysis using our automatically generated transformation from FAFB to JRC 2018F followed by our bridging transformation from JRC 2018F to FCWB. This shows that our new automatic EM to LM transform can be used to draw the same conclusions. Figure 14 shows the results. Red and blue points and bars show the measurements of FAFB traced neurons and LM database neurons, respectively, exactly as in Zheng et al. (2017) with ELM. Black points and bars show the same results using our EM to LM transformation. Notice that the distributions of red and black are very similar as they should be, given that they are measures of the same neurons transformed in different ways. Note that the LM neurons in this analysis were co-registered to the FCWB template by Costa et al. (2016); Jefferis et al. (2007), which we have shown to be a less reliable/accurate registration target than JRC 2018F. Better results (more similar to the EM) could be possible by co-registering to an improved brain template.

### A.8 Block format for transformations

In this section, we motivate and describe a format for storing displacement fields in a block-based (HDF5) file format. Details of the file specification and examples of usage can be found online at https://github.com/saalfeldlab/template-building/wiki/Hdf5-Deformation-fields.

Displacement fields are a common and portable representation for deformable transformations. The ANTs library outputs its transformation as displacement fields. CMTK and elastix use a b-spline representation, and therefore produce output in a more compact (memory-efficient) format, but this comes at the cost of portability. CMTK and elastix each use a custom format that cannot be interchanged between software packages, and therefore both also provide tools for converting transforms to displacement fields.

Displacement fields are typically stored as (32-bit) float valued vector fields, and are therefore costly to store and load from disk. The overhead of loading is especially costly when transforming a sparse set of points rather than an entire image because the entire field is typically loaded from disk before any point coordinates can be transformed. A block-based storage scheme would incur less overhead because it enables efficient access of subset of the displacement field data. HDF5 (“h5”) is a popular and ubiquitous choice for such storage needs (The HDF Group, 1997-2019).

**Figure 14:**
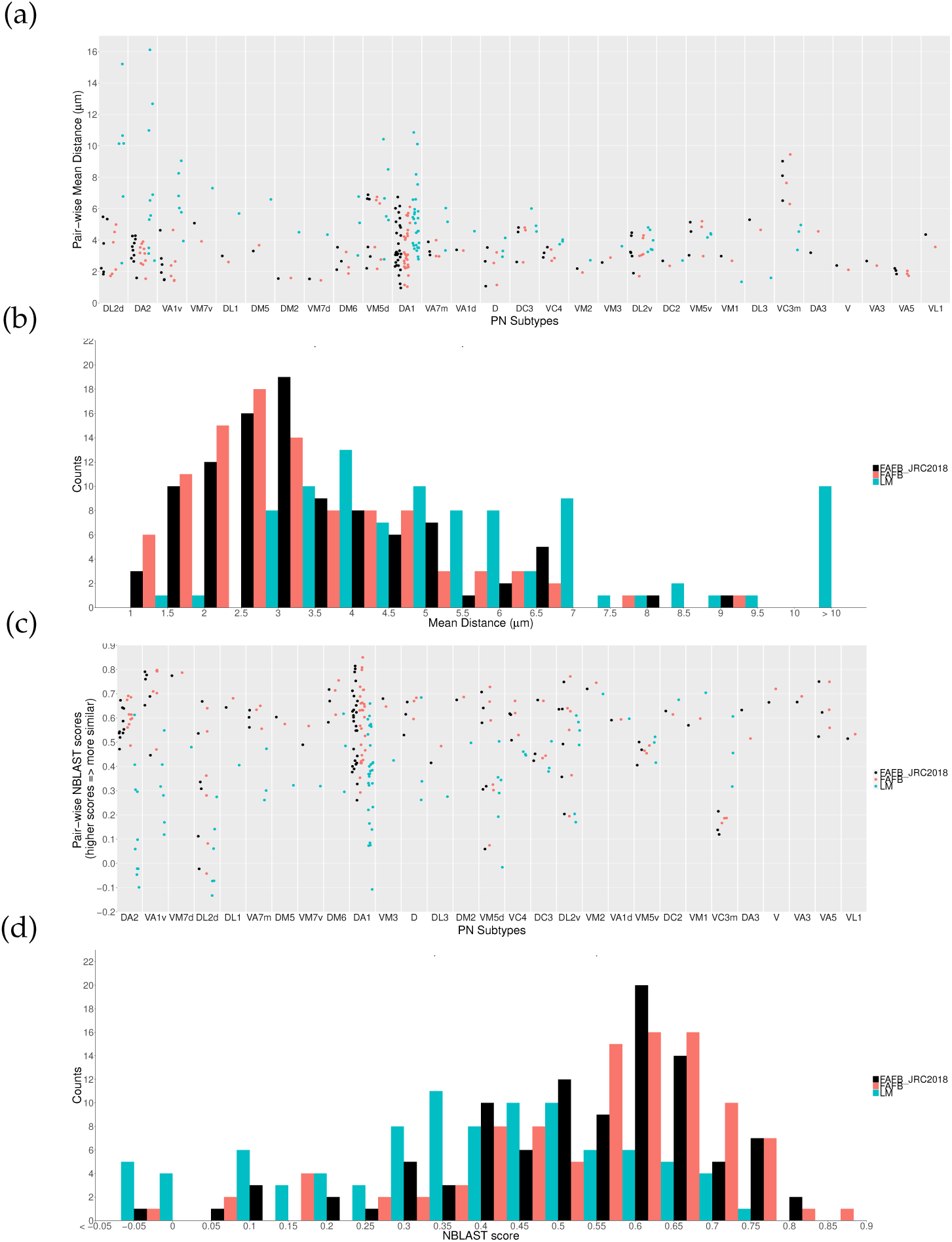
Scatterplots and histograms of (a)-(b) mean skeleton distance and (c)-(d) NBLAST scores for various projection neuron subtypes traced from electron microscopy.

Storage and loading overhead can be improved even further by quantizing and/or downsampling displacement fields. Here we show results quantizing displacement fields to 16-bit precision reducing the raw memory burden by a factor of two. In practice, we observe much greater reductions (over factor of 10 because the quantized displacement fields can be compressed much more efficiently than the original fields, see Table 9).

**Table 9:**
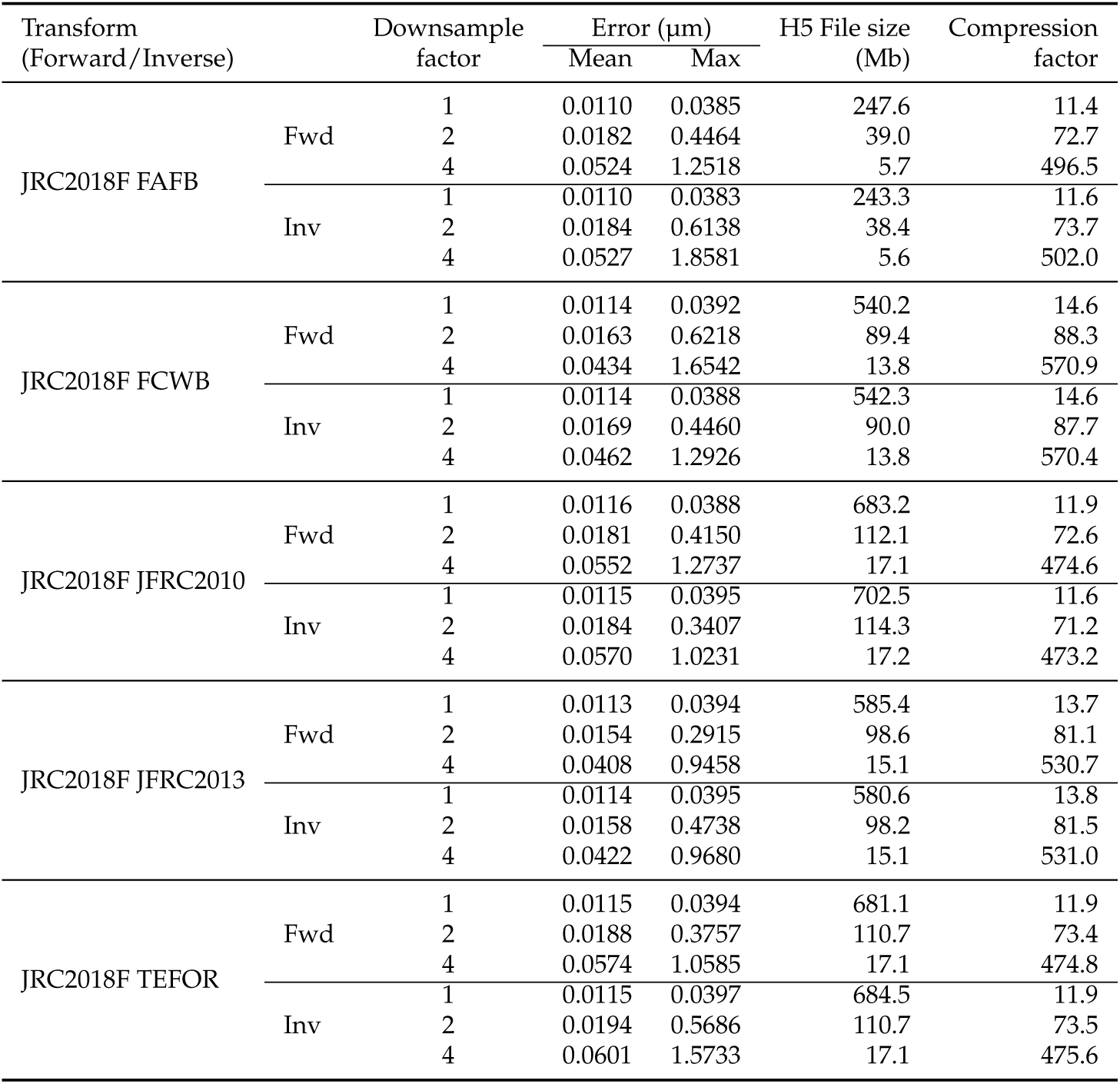
Errors due to quantization and downsampling, and reduction in file size of h5 displacement fields. For a given transformation, specified by (TARGET and SOURCE), and direction (Forward or Inverse), we report the mean and max errors due to quantization and downsampling. Rows with a downsample factor of one, show errors due to quantization only. The two rightmost columns show the size of the quantized and possibly downsampled displacement fields in Megabytes, and the compression factor relative to the compressed original 32-bit float field at full resolution.

| Transform<br>(Forward/Inverse) | | Downsample<br>factor | Error ($\mu\text{m}$) | | H5 File size<br>(Mb) | Compression<br>factor |
| --- | --- | --- | --- | --- | --- | --- |
|  |  |  | Mean | Max |  |  |
| JRC2018F FAFB | Fwd | 1 | 0.0110 | 0.0385 | 247.6 | 11.4 |
|  |  | 2 | 0.0182 | 0.4464 | 39.0 | 72.7 |
|  |  | 4 | 0.0524 | 1.2518 | 5.7 | 496.5 |
|  | Inv | 1 | 0.0110 | 0.0383 | 243.3 | 11.6 |
|  |  | 2 | 0.0184 | 0.6138 | 38.4 | 73.7 |
|  |  | 4 | 0.0527 | 1.8581 | 5.6 | 502.0 |
| JRC2018F FCWB | Fwd | 1 | 0.0114 | 0.0392 | 540.2 | 14.6 |
|  |  | 2 | 0.0163 | 0.6218 | 89.4 | 88.3 |
|  |  | 4 | 0.0434 | 1.6542 | 13.8 | 570.9 |
|  | Inv | 1 | 0.0114 | 0.0388 | 542.3 | 14.6 |
|  |  | 2 | 0.0169 | 0.4460 | 90.0 | 87.7 |
|  |  | 4 | 0.0462 | 1.2926 | 13.8 | 570.4 |
| JRC2018F JFRC2010 | Fwd | 1 | 0.0116 | 0.0388 | 683.2 | 11.9 |
|  |  | 2 | 0.0181 | 0.4150 | 112.1 | 72.6 |
|  |  | 4 | 0.0552 | 1.2737 | 17.1 | 474.6 |
|  | Inv | 1 | 0.0115 | 0.0395 | 702.5 | 11.6 |
|  |  | 2 | 0.0184 | 0.3407 | 114.3 | 71.2 |
|  |  | 4 | 0.0570 | 1.0231 | 17.2 | 473.2 |
| JRC2018F JFRC2013 | Fwd | 1 | 0.0113 | 0.0394 | 585.4 | 13.7 |
|  |  | 2 | 0.0154 | 0.2915 | 98.6 | 81.1 |
|  |  | 4 | 0.0408 | 0.9458 | 15.1 | 530.7 |
|  | Inv | 1 | 0.0114 | 0.0395 | 580.6 | 13.8 |
|  |  | 2 | 0.0158 | 0.4738 | 98.2 | 81.5 |
|  |  | 4 | 0.0422 | 0.9680 | 15.1 | 531.0 |
| JRC2018F TEFOR | Fwd | 1 | 0.0115 | 0.0394 | 681.1 | 11.9 |
|  |  | 2 | 0.0188 | 0.3757 | 110.7 | 73.4 |
|  |  | 4 | 0.0574 | 1.0585 | 17.1 | 474.8 |
|  | Inv | 1 | 0.0115 | 0.0397 | 684.5 | 11.9 |
|  |  | 2 | 0.0194 | 0.5686 | 110.7 | 73.5 |
|  |  | 4 | 0.0601 | 1.5733 | 17.1 | 475.6 |

Quantization inevitably will result in errors relative to the original data, but the magnitude of these errors can be controlled. In our case, we specify the maximum tolerable error as 0.04 µm (less than one-tenth of the pixel spacing). Table 9 shows mean and maximum quantization error (rows with a downsample factor of one), and we observe that errors are less than the specified 0.04 µm threshold as expected. Table 9 also shows errors incurred when both quantizing and downsampling displacement fields by factors of two or four. Downsampling errors are not controllable, as they depend on how quickly the transform changes over space. We observe a maximum error of 1.86 µm (about three voxel widths) for a downsample factor of four.

Our software can apply transformations to neuronal skeletons (stored as SWC files (Stockley et al., 1993)) using transformations stored in several formats. Here we compare the wall clock time required to transform the a skeleton with identical transformations stored either as nifti files (the direct output of ANTs) or stored in the block based format introduced here, using a set of 166 neurons traced in FAFB using CATMAID (Schneider-Mizell et al., 2016). This experiment measures how much block-based loading speeds up the transformation of sparse points. Transformation using nifti displacement fields required an average (standard deviation) of 28.8(1.8) seconds, whereas transforming a neuron with an h5 displacement field took 2.5(0.4) seconds, an 11x speedup.

We also compared the speed at which images can be transformed using displacement fields stored either as uncompressed nifti files, or compressed, quantized h5 file. In this case, we expect any differences in speed to result from two factors. First, the h5 displacement fields comprise half as many bytes (before compression), so fewer bytes will have to be loaded when using h5 files. However, this may be offset somewhat by the decompression necessary when loading from the h5, that is not necessary when loading from the nifti. We transformed 10 images from JRC2018F space to FAFB space. Using displacement fields stored either as nifti files took an average (standard deviation) of 343.3(54.7) seconds, whereas the same transformation using h5 took 292.2(11.5) seconds (wall clock time).

## B. Supplementary Tables

**Table S1:**
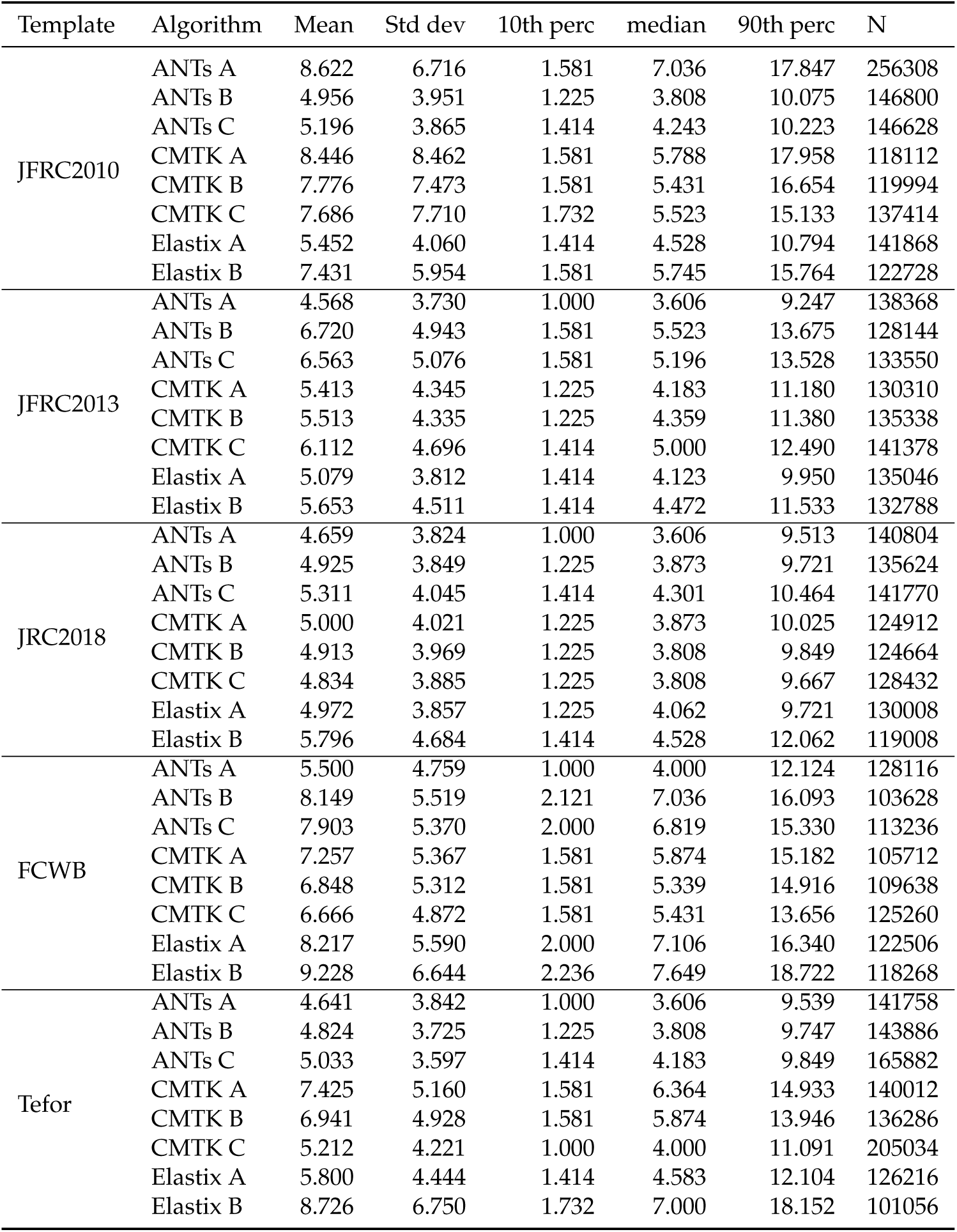
CRE_R : crepine

**Table S2:** CAN_R : cantle

| Template | Algorithm | Mean | Std dev | 10th perc | median | 90th perc | N |
| --- | --- | --- | --- | --- | --- | --- | --- |
| JFRC2010 | ANTs A | 7.403 | 6.977 | 1.581 | 4.950 | 18.221 | 16290 |
|  | ANTs B | 4.236 | 2.557 | 1.414 | 3.808 | 7.583 | 30124 |
|  | ANTs C | 4.126 | 2.587 | 1.225 | 3.606 | 7.416 | 37836 |
|  | CMTK A | 6.262 | 6.624 | 1.225 | 4.062 | 15.000 | 35436 |
|  | CMTK B | 5.936 | 6.301 | 1.225 | 3.808 | 14.916 | 30950 |
|  | CMTK C | 6.016 | 5.701 | 1.581 | 4.183 | 14.933 | 33842 |
|  | Elastix A | 4.645 | 3.253 | 1.225 | 3.808 | 9.028 | 14486 |
|  | Elastix B | 4.522 | 3.060 | 1.225 | 3.808 | 8.868 | 14868 |
| JFRC2013 | ANTs A | 3.932 | 2.799 | 1.000 | 3.240 | 7.649 | 21630 |
|  | ANTs B | 3.346 | 2.276 | 1.000 | 2.915 | 6.325 | 47016 |
|  | ANTs C | 3.153 | 2.230 | 0.707 | 2.646 | 6.000 | 44028 |
|  | CMTK A | 3.529 | 2.666 | 1.000 | 2.915 | 6.856 | 24354 |
|  | CMTK B | 3.556 | 2.687 | 1.000 | 2.915 | 6.964 | 24016 |
|  | CMTK C | 4.098 | 2.588 | 1.225 | 3.606 | 7.583 | 34654 |
|  | Elastix A | 3.719 | 2.763 | 1.000 | 3.000 | 7.382 | 23464 |
|  | Elastix B | 4.062 | 2.963 | 1.000 | 3.240 | 8.093 | 21682 |
| JRC2018 | ANTs A | 3.768 | 2.749 | 1.000 | 3.000 | 7.382 | 21558 |
|  | ANTs B | 4.132 | 2.791 | 1.000 | 3.606 | 7.649 | 23310 |
|  | ANTs C | 4.038 | 2.633 | 1.225 | 3.536 | 7.416 | 26778 |
|  | CMTK A | 3.868 | 2.804 | 1.000 | 3.162 | 7.616 | 21388 |
|  | CMTK B | 3.908 | 2.907 | 1.000 | 3.082 | 7.842 | 21008 |
|  | CMTK C | 3.809 | 2.839 | 1.000 | 3.082 | 7.550 | 21214 |
|  | Elastix A | 3.924 | 2.821 | 1.000 | 3.240 | 7.810 | 21484 |
|  | Elastix B | 4.162 | 2.984 | 1.000 | 3.536 | 8.185 | 19802 |
| FCWB | ANTs A | 3.755 | 2.665 | 1.000 | 3.082 | 7.246 | 22464 |
|  | ANTs B | 3.844 | 2.335 | 1.225 | 3.536 | 6.745 | 26150 |
|  | ANTs C | 3.759 | 2.273 | 1.225 | 3.464 | 6.708 | 22568 |
|  | CMTK A | 3.814 | 2.649 | 1.000 | 3.240 | 7.280 | 46938 |
|  | CMTK B | 3.866 | 2.702 | 1.000 | 3.240 | 7.382 | 31124 |
|  | CMTK C | 4.068 | 2.855 | 1.000 | 3.536 | 7.906 | 33454 |
|  | Elastix A | 3.782 | 3.067 | 1.000 | 3.240 | 6.856 | 39318 |
|  | Elastix B | 4.687 | 3.322 | 1.000 | 3.808 | 9.618 | 56956 |
| Tefor | ANTs A | 4.258 | 2.985 | 1.225 | 3.606 | 8.185 | 19428 |
|  | ANTs B | 4.267 | 2.671 | 1.414 | 3.808 | 7.649 | 24896 |
|  | ANTs C | 4.942 | 2.938 | 1.581 | 4.528 | 8.631 | 30864 |
|  | CMTK A | 4.763 | 3.168 | 1.414 | 4.123 | 9.274 | 23110 |
|  | CMTK B | 4.860 | 3.102 | 1.414 | 4.243 | 9.343 | 23078 |
|  | CMTK C | 4.490 | 3.093 | 1.414 | 3.808 | 8.544 | 21146 |
|  | Elastix A | 4.494 | 2.952 | 1.225 | 3.808 | 8.399 | 15300 |
|  | Elastix B | 4.062 | 2.450 | 1.225 | 3.606 | 7.382 | 18314 |

**Table S3:** AMMC_R : antennal mechanosensory and motor center

| Template | Algorithm | Mean | Std dev | 10th perc | median | 90th perc | N |
| --- | --- | --- | --- | --- | --- | --- | --- |
| JFRC2010 | ANTs A | 7.150 | 7.956 | 1.581 | 4.528 | 19.770 | 225304 |
|  | ANTs B | 5.234 | 4.993 | 1.581 | 4.183 | 8.746 | 187944 |
|  | ANTs C | 5.416 | 5.060 | 1.581 | 4.359 | 9.165 | 193640 |
|  | CMTK A | 5.177 | 4.705 | 1.414 | 4.123 | 9.192 | 189018 |
|  | CMTK B | 5.022 | 4.600 | 1.414 | 4.062 | 8.602 | 194876 |
|  | CMTK C | 5.253 | 5.016 | 1.581 | 4.183 | 8.746 | 187834 |
|  | Elastix A | 4.623 | 4.374 | 1.414 | 3.674 | 7.810 | 187814 |
|  | Elastix B | 4.646 | 4.339 | 1.414 | 3.808 | 7.842 | 198088 |
| JFRC2013 | ANTs A | 4.671 | 4.553 | 1.225 | 3.674 | 7.906 | 194144 |
|  | ANTs B | 6.253 | 6.122 | 1.581 | 4.743 | 11.180 | 163990 |
|  | ANTs C | 6.093 | 6.197 | 1.581 | 4.528 | 11.269 | 159530 |
|  | CMTK A | 5.040 | 4.726 | 1.581 | 4.062 | 8.631 | 178554 |
|  | CMTK B | 5.070 | 4.724 | 1.414 | 4.062 | 8.660 | 172350 |
|  | CMTK C | 5.755 | 4.961 | 1.581 | 4.528 | 10.440 | 139908 |
|  | Elastix A | 4.571 | 4.670 | 1.225 | 3.606 | 7.649 | 192020 |
|  | Elastix B | 4.659 | 4.590 | 1.414 | 3.674 | 7.810 | 187546 |
| JRC2018 | ANTs A | 4.582 | 4.503 | 1.225 | 3.606 | 7.649 | 196510 |
|  | ANTs B | 4.853 | 4.557 | 1.414 | 3.873 | 8.155 | 200496 |
|  | ANTs C | 5.003 | 4.938 | 1.414 | 3.873 | 8.515 | 220340 |
|  | CMTK A | 4.663 | 4.622 | 1.225 | 3.674 | 7.842 | 192616 |
|  | CMTK B | 4.673 | 4.505 | 1.414 | 3.674 | 7.906 | 190402 |
|  | CMTK C | 4.754 | 4.397 | 1.414 | 3.808 | 8.031 | 193484 |
|  | Elastix A | 4.570 | 4.344 | 1.414 | 3.674 | 7.616 | 196662 |
|  | Elastix B | 4.712 | 4.633 | 1.414 | 3.808 | 7.906 | 190378 |
| FCWB | ANTs A | 4.766 | 4.218 | 1.414 | 3.873 | 8.276 | 198206 |
|  | ANTs B | 5.606 | 5.450 | 1.581 | 4.472 | 9.539 | 118958 |
|  | ANTs C | 5.231 | 4.522 | 1.581 | 4.183 | 9.247 | 129264 |
|  | CMTK A | 5.296 | 4.588 | 1.581 | 4.359 | 9.301 | 189414 |
|  | CMTK B | 5.277 | 4.672 | 1.581 | 4.301 | 9.220 | 187246 |
|  | CMTK C | 5.168 | 4.630 | 1.581 | 4.243 | 8.746 | 153568 |
|  | Elastix A | 5.405 | 6.600 | 1.414 | 4.123 | 8.631 | 197108 |
|  | Elastix B | 4.817 | 5.207 | 1.414 | 3.808 | 8.000 | 201070 |
| Tefor | ANTs A | 4.629 | 4.464 | 1.225 | 3.674 | 7.810 | 195792 |
|  | ANTs B | 4.936 | 4.866 | 1.414 | 3.808 | 8.544 | 206934 |
|  | ANTs C | 5.295 | 5.410 | 1.581 | 4.183 | 8.718 | 221890 |
|  | CMTK A | 5.096 | 4.741 | 1.414 | 3.873 | 9.028 | 198934 |
|  | CMTK B | 4.896 | 4.528 | 1.414 | 3.873 | 8.515 | 203194 |
|  | CMTK C | 5.165 | 5.229 | 1.581 | 4.123 | 8.515 | 201560 |
|  | Elastix A | 4.603 | 4.346 | 1.414 | 3.674 | 7.746 | 187738 |
|  | Elastix B | 4.745 | 4.064 | 1.414 | 3.873 | 8.093 | 197718 |

**Table S4:** ICL_R : inferior clamp

| Template | Algorithm | Mean | Std dev | 10th perc | median | 90th perc | N |
| --- | --- | --- | --- | --- | --- | --- | --- |
| JFRC2010 | ANTs A | 8.835 | 7.213 | 1.414 | 6.745 | 19.506 | 501078 |
|  | ANTs B | 3.167 | 2.986 | 0.707 | 2.236 | 6.671 | 316016 |
|  | ANTs C | 3.315 | 3.202 | 0.707 | 2.345 | 7.106 | 307394 |
|  | CMTK A | 4.211 | 5.414 | 0.707 | 2.236 | 11.203 | 293368 |
|  | CMTK B | 4.158 | 5.566 | 0.707 | 2.236 | 9.539 | 289230 |
|  | CMTK C | 4.177 | 4.478 | 0.707 | 2.828 | 9.220 | 331886 |
|  | Elastix A | 2.948 | 2.765 | 0.707 | 2.121 | 6.205 | 300488 |
|  | Elastix B | 3.122 | 3.178 | 0.707 | 2.121 | 7.000 | 311002 |
| JFRC2013 | ANTs A | 2.697 | 2.771 | 0.707 | 2.000 | 5.568 | 321788 |
|  | ANTs B | 3.434 | 3.348 | 0.707 | 2.345 | 7.681 | 299558 |
|  | ANTs C | 3.219 | 3.168 | 0.707 | 2.236 | 7.036 | 303386 |
|  | CMTK A | 3.011 | 2.874 | 0.707 | 2.121 | 6.403 | 293092 |
|  | CMTK B | 3.169 | 2.972 | 0.707 | 2.236 | 6.819 | 290932 |
|  | CMTK C | 4.035 | 3.561 | 0.707 | 3.000 | 8.485 | 263762 |
|  | Elastix A | 2.726 | 2.688 | 0.707 | 2.000 | 5.701 | 320686 |
|  | Elastix B | 2.674 | 2.733 | 0.707 | 2.000 | 5.568 | 328936 |
| JRC2018 | ANTs A | 2.673 | 2.700 | 0.707 | 2.000 | 5.568 | 319308 |
|  | ANTs B | 2.861 | 2.664 | 0.707 | 2.121 | 5.745 | 326204 |
|  | ANTs C | 2.902 | 2.819 | 0.707 | 2.121 | 6.000 | 329458 |
|  | CMTK A | 2.679 | 2.803 | 0.707 | 2.000 | 5.568 | 298028 |
|  | CMTK B | 2.660 | 2.771 | 0.707 | 1.732 | 5.568 | 296780 |
|  | CMTK C | 2.717 | 2.683 | 0.707 | 2.000 | 5.568 | 303460 |
|  | Elastix A | 2.742 | 2.706 | 0.707 | 2.121 | 5.657 | 312578 |
|  | Elastix B | 2.711 | 2.714 | 0.707 | 2.000 | 5.568 | 310436 |
| FCWB | ANTs A | 2.758 | 2.812 | 0.707 | 2.000 | 5.874 | 334680 |
|  | ANTs B | 3.551 | 3.415 | 0.707 | 2.449 | 7.937 | 310126 |
|  | ANTs C | 3.377 | 3.280 | 0.707 | 2.345 | 7.416 | 305450 |
|  | CMTK A | 5.384 | 5.552 | 0.707 | 3.536 | 12.981 | 283240 |
|  | CMTK B | 4.969 | 5.255 | 0.707 | 3.240 | 11.269 | 264802 |
|  | CMTK C | 3.833 | 4.041 | 0.707 | 2.550 | 8.631 | 304944 |
|  | Elastix A | 6.306 | 6.023 | 0.707 | 4.183 | 15.313 | 290364 |
|  | Elastix B | 3.448 | 4.041 | 0.707 | 2.121 | 7.906 | 304660 |
| Tefor | ANTs A | 2.671 | 2.668 | 0.707 | 2.000 | 5.568 | 321736 |
|  | ANTs B | 3.090 | 2.853 | 0.707 | 2.236 | 6.557 | 322866 |
|  | ANTs C | 3.239 | 3.132 | 0.707 | 2.236 | 7.036 | 319676 |
|  | CMTK A | 3.468 | 4.315 | 0.707 | 2.121 | 8.031 | 296402 |
|  | CMTK B | 3.586 | 4.498 | 0.707 | 2.121 | 8.337 | 299568 |
|  | CMTK C | 3.490 | 3.250 | 0.707 | 2.550 | 7.583 | 319366 |
|  | Elastix A | 2.837 | 2.731 | 0.707 | 2.121 | 5.916 | 305932 |
|  | Elastix B | 3.276 | 3.410 | 0.707 | 2.121 | 7.314 | 315504 |

**Table S5:** VES_R : vest

| Template | Algorithm | Mean | Std dev | 10th perc | median | 90th perc | N |
| --- | --- | --- | --- | --- | --- | --- | --- |
| JFRC2010 | ANTs A | 8.273 | 6.716 | 1.732 | 6.205 | 18.125 | 124266 |
|  | ANTs B | 4.619 | 3.635 | 1.225 | 3.808 | 8.972 | 171134 |
|  | ANTs C | 4.846 | 4.001 | 1.000 | 3.808 | 9.925 | 176908 |
|  | CMTK A | 7.379 | 6.750 | 1.414 | 5.000 | 17.277 | 162750 |
|  | CMTK B | 7.217 | 6.801 | 1.414 | 4.743 | 17.132 | 139584 |
|  | CMTK C | 6.430 | 6.078 | 1.414 | 4.301 | 15.199 | 184334 |
|  | Elastix A | 4.157 | 3.351 | 1.000 | 3.240 | 8.276 | 136550 |
|  | Elastix B | 4.352 | 3.374 | 1.000 | 3.536 | 8.660 | 122002 |
| JFRC2013 | ANTs A | 4.537 | 3.631 | 1.000 | 3.606 | 9.055 | 113054 |
|  | ANTs B | 4.515 | 3.459 | 1.225 | 3.606 | 8.860 | 279266 |
|  | ANTs C | 4.575 | 3.668 | 1.000 | 3.606 | 9.247 | 244618 |
|  | CMTK A | 4.719 | 3.648 | 1.225 | 3.808 | 9.539 | 98662 |
|  | CMTK B | 4.759 | 3.657 | 1.225 | 3.808 | 9.513 | 99628 |
|  | CMTK C | 4.877 | 3.859 | 1.225 | 3.808 | 10.025 | 182464 |
|  | Elastix A | 4.506 | 3.545 | 1.225 | 3.606 | 8.972 | 116176 |
|  | Elastix B | 4.487 | 3.474 | 1.000 | 3.606 | 8.746 | 111036 |
| JRC2018 | ANTs A | 4.464 | 3.631 | 1.000 | 3.606 | 8.972 | 111894 |
|  | ANTs B | 4.932 | 3.717 | 1.225 | 4.062 | 9.823 | 136380 |
|  | ANTs C | 4.973 | 3.597 | 1.414 | 4.062 | 9.925 | 135970 |
|  | CMTK A | 4.531 | 3.617 | 1.000 | 3.606 | 9.000 | 95754 |
|  | CMTK B | 4.536 | 3.632 | 1.000 | 3.606 | 9.055 | 92400 |
|  | CMTK C | 4.594 | 3.672 | 1.000 | 3.674 | 9.220 | 100094 |
|  | Elastix A | 4.487 | 3.531 | 1.000 | 3.606 | 8.718 | 118292 |
|  | Elastix B | 4.466 | 3.374 | 1.000 | 3.674 | 8.660 | 112558 |
| FCWB | ANTs A | 4.787 | 3.985 | 1.000 | 3.674 | 9.592 | 112756 |
|  | ANTs B | 4.878 | 3.636 | 1.414 | 4.062 | 9.513 | 221906 |
|  | ANTs C | 4.971 | 3.861 | 1.414 | 4.062 | 9.849 | 184380 |
|  | CMTK A | 7.352 | 5.921 | 1.581 | 5.431 | 16.340 | 189818 |
|  | CMTK B | 6.945 | 5.472 | 1.581 | 5.292 | 14.868 | 139980 |
|  | CMTK C | 6.271 | 4.814 | 1.581 | 4.950 | 13.285 | 153026 |
|  | Elastix A | 7.003 | 5.628 | 1.581 | 5.431 | 14.474 | 264300 |
|  | Elastix B | 7.055 | 5.435 | 1.581 | 5.431 | 15.330 | 205946 |
| Tefor | ANTs A | 4.442 | 3.502 | 1.000 | 3.606 | 8.746 | 120918 |
|  | ANTs B | 4.467 | 3.393 | 1.225 | 3.606 | 8.660 | 201810 |
|  | ANTs C | 5.259 | 3.794 | 1.414 | 4.359 | 10.271 | 194238 |
|  | CMTK A | 7.314 | 5.607 | 1.581 | 5.788 | 16.016 | 156436 |
|  | CMTK B | 7.404 | 5.644 | 1.581 | 5.874 | 16.016 | 160204 |
|  | CMTK C | 6.877 | 5.227 | 1.581 | 5.292 | 14.849 | 165862 |
|  | Elastix A | 4.217 | 3.378 | 1.000 | 3.464 | 8.185 | 147790 |
|  | Elastix B | 4.809 | 3.647 | 1.225 | 3.808 | 9.925 | 168850 |

**Table S6:** IB_R : inferior bridge

| Template | Algorithm | Mean | Std dev | 10th perc | median | 90th perc | N |
| --- | --- | --- | --- | --- | --- | --- | --- |
| JFRC2010 | ANTs A | 5.841 | 5.356 | 1.000 | 3.873 | 13.416 | 212072 |
|  | ANTs B | 4.067 | 3.357 | 1.000 | 3.082 | 8.660 | 255490 |
|  | ANTs C | 4.098 | 3.194 | 1.000 | 3.240 | 8.426 | 364138 |
|  | CMTK A | 4.722 | 4.537 | 1.000 | 3.240 | 10.630 | 247624 |
|  | CMTK B | 4.457 | 4.372 | 1.000 | 3.082 | 9.644 | 257816 |
|  | CMTK C | 4.417 | 4.448 | 1.000 | 3.082 | 9.220 | 370982 |
|  | Elastix A | 3.060 | 2.493 | 0.707 | 2.449 | 5.916 | 259080 |
|  | Elastix B | 3.670 | 3.090 | 0.707 | 2.828 | 7.906 | 211838 |
| JFRC2013 | ANTs A | 3.163 | 2.767 | 0.707 | 2.345 | 6.557 | 210128 |
|  | ANTs B | 4.260 | 3.368 | 1.000 | 3.317 | 8.775 | 324356 |
|  | ANTs C | 4.210 | 3.293 | 1.000 | 3.317 | 8.775 | 325204 |
|  | CMTK A | 3.366 | 2.867 | 0.707 | 2.550 | 6.964 | 181380 |
|  | CMTK B | 3.509 | 2.882 | 0.707 | 2.646 | 7.280 | 178796 |
|  | CMTK C | 4.819 | 3.925 | 1.000 | 3.606 | 11.068 | 207494 |
|  | Elastix A | 3.105 | 2.621 | 0.707 | 2.345 | 6.325 | 213226 |
|  | Elastix B | 3.115 | 2.625 | 0.707 | 2.345 | 6.403 | 201400 |
| JRC2018 | ANTs A | 3.101 | 2.674 | 0.707 | 2.345 | 6.403 | 216404 |
|  | ANTs B | 3.405 | 2.886 | 0.707 | 2.646 | 7.071 | 226268 |
|  | ANTs C | 3.214 | 2.667 | 0.707 | 2.550 | 6.557 | 227828 |
|  | CMTK A | 3.321 | 3.016 | 0.707 | 2.449 | 7.106 | 183046 |
|  | CMTK B | 3.190 | 2.844 | 0.707 | 2.345 | 6.671 | 181056 |
|  | CMTK C | 3.128 | 2.739 | 0.707 | 2.345 | 6.442 | 195562 |
|  | Elastix A | 3.155 | 2.681 | 0.707 | 2.449 | 6.403 | 203250 |
|  | Elastix B | 3.184 | 2.710 | 0.707 | 2.449 | 6.633 | 198934 |
| FCWB | ANTs A | 3.343 | 2.863 | 0.707 | 2.550 | 6.856 | 208368 |
|  | ANTs B | 4.804 | 3.684 | 1.000 | 3.808 | 10.149 | 295074 |
|  | ANTs C | 3.921 | 2.902 | 1.000 | 3.162 | 7.906 | 345268 |
|  | CMTK A | 6.347 | 5.291 | 1.225 | 4.743 | 13.964 | 204434 |
|  | CMTK B | 5.750 | 4.965 | 1.000 | 4.123 | 12.923 | 198092 |
|  | CMTK C | 4.281 | 3.728 | 1.000 | 3.082 | 9.513 | 246048 |
|  | Elastix A | 7.917 | 7.926 | 1.414 | 5.292 | 18.152 | 242588 |
|  | Elastix B | 4.365 | 4.009 | 1.000 | 3.162 | 9.434 | 222796 |
| Tefor | ANTs A | 3.107 | 2.665 | 0.707 | 2.345 | 6.364 | 212832 |
|  | ANTs B | 3.848 | 3.211 | 1.000 | 2.915 | 8.031 | 239752 |
|  | ANTs C | 3.998 | 3.250 | 1.000 | 3.082 | 8.515 | 275152 |
|  | CMTK A | 4.508 | 4.170 | 0.707 | 3.000 | 10.863 | 206740 |
|  | CMTK B | 4.501 | 4.218 | 0.707 | 3.000 | 10.977 | 208652 |
|  | CMTK C | 4.508 | 3.874 | 1.000 | 3.240 | 10.223 | 206512 |
|  | Elastix A | 3.207 | 2.703 | 0.707 | 2.550 | 6.481 | 219002 |
|  | Elastix B | 3.689 | 3.088 | 0.707 | 2.828 | 8.000 | 211786 |

**Table S7:** ATL_R : antler

| Template | Algorithm | Mean | Std dev | 10th perc | median | 90th perc | N |
| --- | --- | --- | --- | --- | --- | --- | --- |
| JFRC2010 | ANTs A | 4.343 | 3.451 | 1.000 | 3.536 | 8.485 | 125792 |
|  | ANTs B | 4.230 | 2.710 | 1.225 | 3.674 | 8.031 | 99154 |
|  | ANTs C | 4.473 | 3.157 | 1.000 | 3.674 | 9.000 | 116250 |
|  | CMTK A | 4.913 | 3.653 | 1.000 | 4.000 | 10.000 | 100040 |
|  | CMTK B | 4.735 | 3.419 | 1.000 | 3.873 | 9.539 | 100534 |
|  | CMTK C | 4.179 | 3.048 | 1.000 | 3.536 | 8.276 | 124706 |
|  | Elastix A | 3.471 | 2.269 | 1.000 | 3.000 | 6.557 | 98164 |
|  | Elastix B | 3.774 | 2.437 | 1.000 | 3.240 | 7.141 | 108068 |
| JFRC2013 | ANTs A | 3.491 | 2.498 | 1.000 | 2.915 | 6.557 | 102024 |
|  | ANTs B | 4.523 | 3.082 | 1.225 | 3.808 | 8.888 | 113776 |
|  | ANTs C | 4.536 | 3.198 | 1.000 | 3.808 | 9.110 | 110298 |
|  | CMTK A | 3.449 | 2.294 | 1.000 | 2.915 | 6.481 | 93474 |
|  | CMTK B | 3.571 | 2.345 | 1.000 | 3.082 | 6.671 | 90208 |
|  | CMTK C | 4.575 | 2.978 | 1.414 | 3.873 | 8.718 | 90024 |
|  | Elastix A | 3.454 | 2.241 | 1.000 | 3.000 | 6.403 | 100400 |
|  | Elastix B | 3.377 | 2.305 | 1.000 | 2.915 | 6.519 | 104878 |
| JRC2018 | ANTs A | 3.239 | 2.275 | 1.000 | 2.646 | 6.083 | 105940 |
|  | ANTs B | 3.613 | 2.450 | 1.000 | 3.082 | 6.708 | 101916 |
|  | ANTs C | 3.546 | 2.438 | 1.000 | 3.000 | 6.671 | 101848 |
|  | CMTK A | 3.383 | 2.437 | 1.000 | 2.915 | 6.481 | 100240 |
|  | CMTK B | 3.374 | 2.375 | 1.000 | 2.915 | 6.442 | 99604 |
|  | CMTK C | 3.408 | 2.293 | 1.000 | 2.915 | 6.364 | 97688 |
|  | Elastix A | 3.435 | 2.284 | 1.000 | 3.000 | 6.519 | 96144 |
|  | Elastix B | 3.345 | 2.313 | 1.000 | 2.915 | 6.442 | 105946 |
| FCWB | ANTs A | 3.665 | 2.486 | 1.000 | 3.162 | 6.856 | 95158 |
|  | ANTs B | 4.521 | 2.948 | 1.225 | 3.808 | 8.746 | 107846 |
|  | ANTs C | 4.260 | 2.853 | 1.225 | 3.606 | 8.367 | 118618 |
|  | CMTK A | 5.780 | 5.223 | 1.414 | 4.359 | 11.358 | 72358 |
|  | CMTK B | 5.523 | 5.176 | 1.414 | 4.062 | 11.180 | 76574 |
|  | CMTK C | 4.258 | 2.995 | 1.225 | 3.606 | 8.093 | 88152 |
|  | Elastix A | 6.748 | 6.980 | 1.581 | 4.950 | 12.207 | 91484 |
|  | Elastix B | 4.385 | 3.467 | 1.000 | 3.536 | 8.660 | 93730 |
| Tefor | ANTs A | 3.239 | 2.249 | 1.000 | 2.828 | 6.083 | 104392 |
|  | ANTs B | 4.069 | 2.637 | 1.225 | 3.606 | 7.616 | 99864 |
|  | ANTs C | 4.376 | 3.001 | 1.225 | 3.674 | 8.544 | 108260 |
|  | CMTK A | 3.885 | 2.724 | 1.000 | 3.240 | 7.583 | 112418 |
|  | CMTK B | 3.824 | 2.622 | 1.000 | 3.240 | 7.348 | 114492 |
|  | CMTK C | 4.280 | 2.708 | 1.225 | 3.808 | 8.000 | 98510 |
|  | Elastix A | 3.430 | 2.297 | 1.000 | 2.915 | 6.633 | 104642 |
|  | Elastix B | 3.480 | 2.372 | 1.000 | 3.000 | 6.671 | 117174 |

**Table S8:**
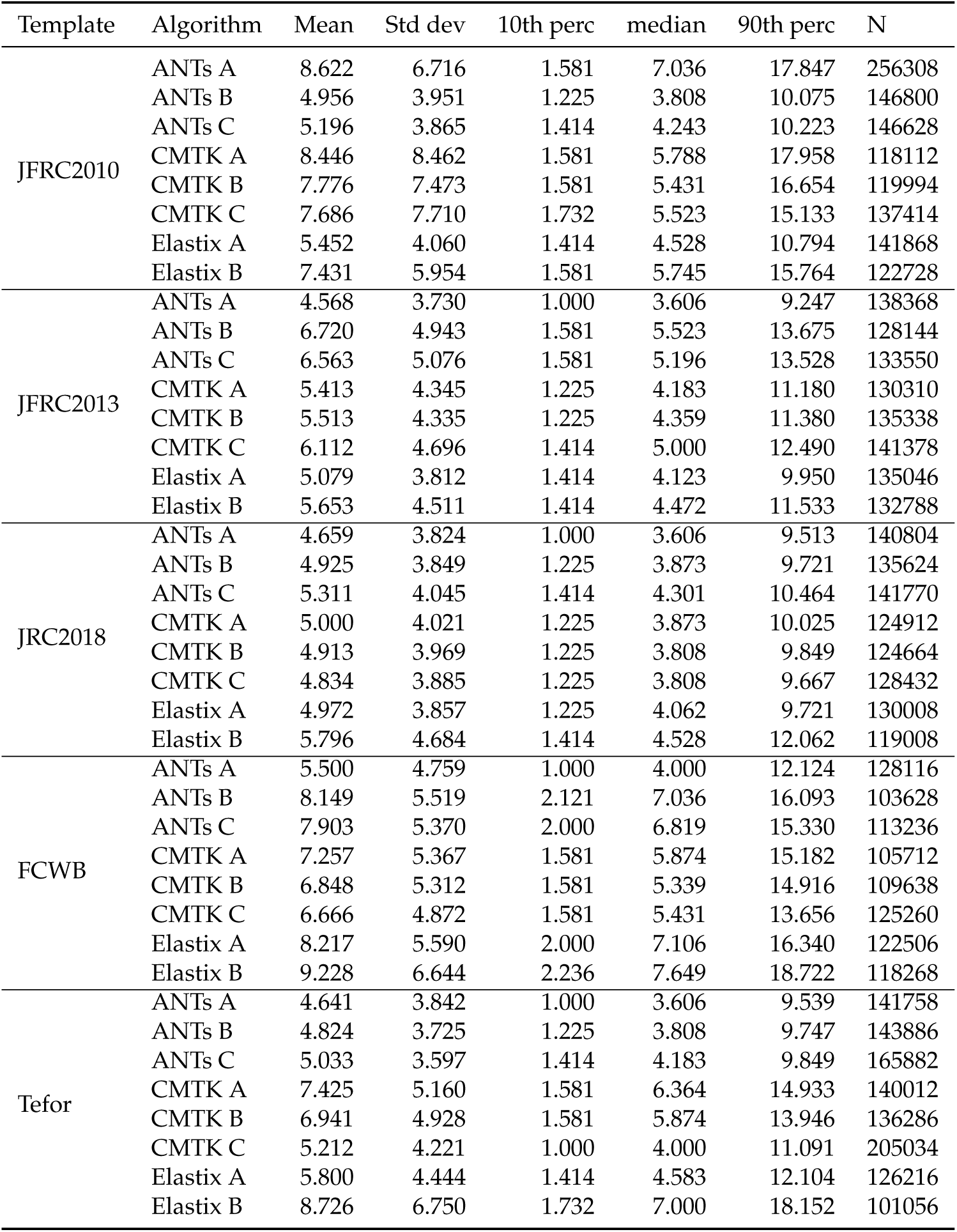
CRE_R : crepine

**Table S9:**
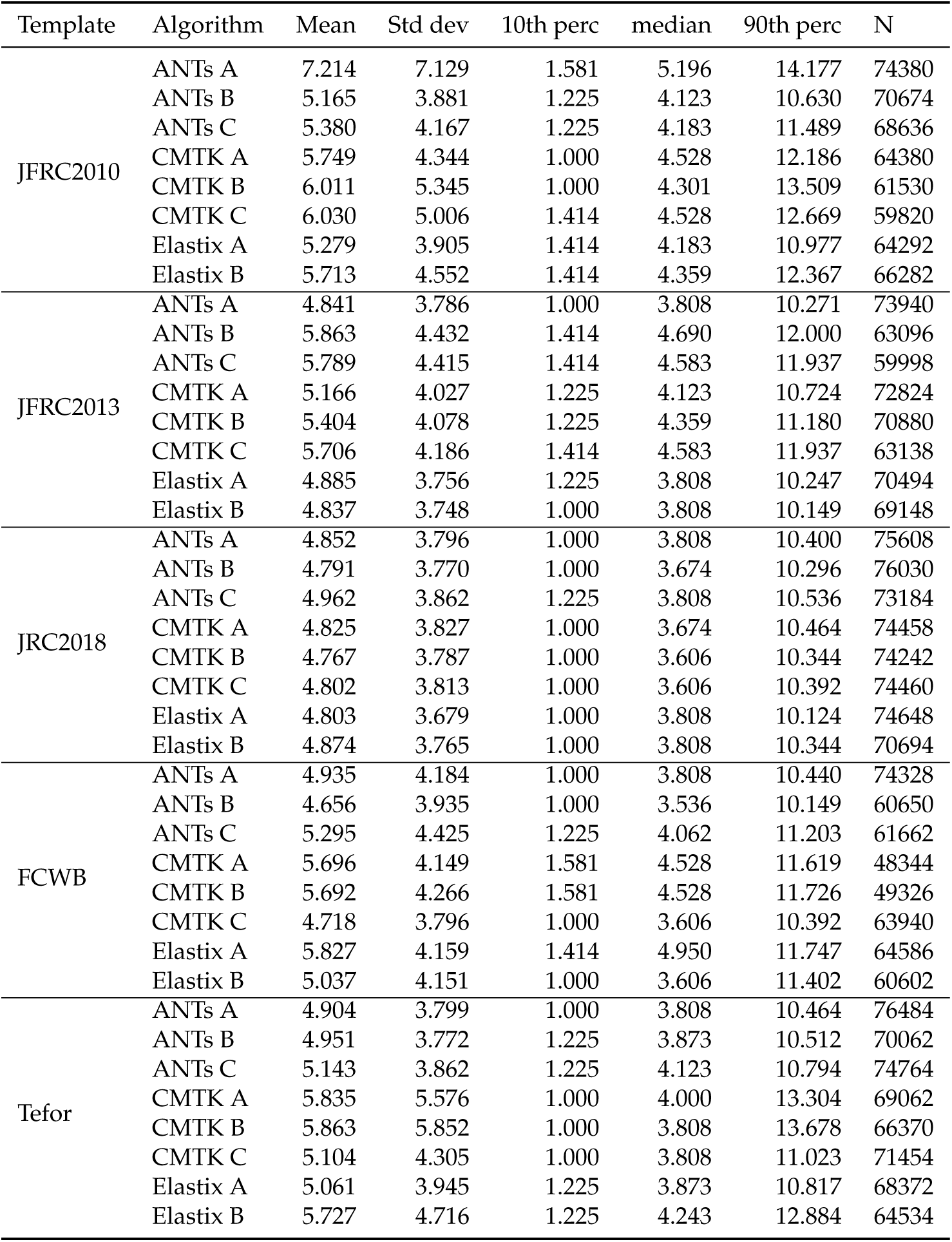
MB_PED_R : pedunculus of adult mushroom body

**Table S10:**
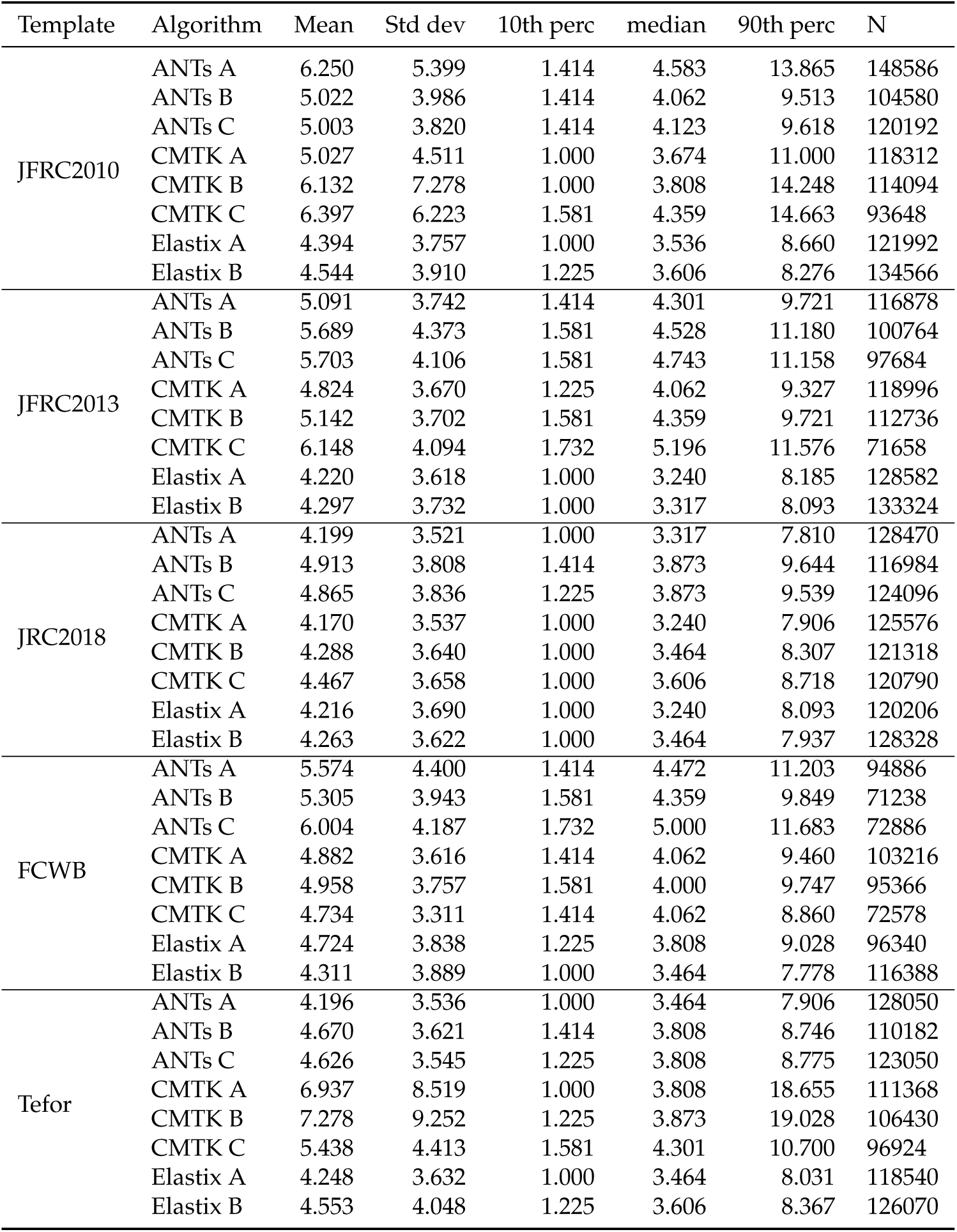
MB_VL_R : vertical lobe of adult mushroom body

| Template | Algorithm | Mean | Std dev | 10th perc | median | 90th perc | N |
| --- | --- | --- | --- | --- | --- | --- | --- |
| JFRC2010 | ANTs A | 6.250 | 5.399 | 1.414 | 4.583 | 13.865 | 148586 |
|  | ANTs B | 5.022 | 3.986 | 1.414 | 4.062 | 9.513 | 104580 |
|  | ANTs C | 5.003 | 3.820 | 1.414 | 4.123 | 9.618 | 120192 |
|  | CMTK A | 5.027 | 4.511 | 1.000 | 3.674 | 11.000 | 118312 |
|  | CMTK B | 6.132 | 7.278 | 1.000 | 3.808 | 14.248 | 114094 |
|  | CMTK C | 6.397 | 6.223 | 1.581 | 4.359 | 14.663 | 93648 |
|  | Elastix A | 4.394 | 3.757 | 1.000 | 3.536 | 8.660 | 121992 |
|  | Elastix B | 4.544 | 3.910 | 1.225 | 3.606 | 8.276 | 134566 |
| JFRC2013 | ANTs A | 5.091 | 3.742 | 1.414 | 4.301 | 9.721 | 116878 |
|  | ANTs B | 5.689 | 4.373 | 1.581 | 4.528 | 11.180 | 100764 |
|  | ANTs C | 5.703 | 4.106 | 1.581 | 4.743 | 11.158 | 97684 |
|  | CMTK A | 4.824 | 3.670 | 1.225 | 4.062 | 9.327 | 118996 |
|  | CMTK B | 5.142 | 3.702 | 1.581 | 4.359 | 9.721 | 112736 |
|  | CMTK C | 6.148 | 4.094 | 1.732 | 5.196 | 11.576 | 71658 |
|  | Elastix A | 4.220 | 3.618 | 1.000 | 3.240 | 8.185 | 128582 |
|  | Elastix B | 4.297 | 3.732 | 1.000 | 3.317 | 8.093 | 133324 |
| JRC2018 | ANTs A | 4.199 | 3.521 | 1.000 | 3.317 | 7.810 | 128470 |
|  | ANTs B | 4.913 | 3.808 | 1.414 | 3.873 | 9.644 | 116984 |
|  | ANTs C | 4.865 | 3.836 | 1.225 | 3.873 | 9.539 | 124096 |
|  | CMTK A | 4.170 | 3.537 | 1.000 | 3.240 | 7.906 | 125576 |
|  | CMTK B | 4.288 | 3.640 | 1.000 | 3.464 | 8.307 | 121318 |
|  | CMTK C | 4.467 | 3.658 | 1.000 | 3.606 | 8.718 | 120790 |
|  | Elastix A | 4.216 | 3.690 | 1.000 | 3.240 | 8.093 | 120206 |
|  | Elastix B | 4.263 | 3.622 | 1.000 | 3.464 | 7.937 | 128328 |
| FCWB | ANTs A | 5.574 | 4.400 | 1.414 | 4.472 | 11.203 | 94886 |
|  | ANTs B | 5.305 | 3.943 | 1.581 | 4.359 | 9.849 | 71238 |
|  | ANTs C | 6.004 | 4.187 | 1.732 | 5.000 | 11.683 | 72886 |
|  | CMTK A | 4.882 | 3.616 | 1.414 | 4.062 | 9.460 | 103216 |
|  | CMTK B | 4.958 | 3.757 | 1.581 | 4.000 | 9.747 | 95366 |
|  | CMTK C | 4.734 | 3.311 | 1.414 | 4.062 | 8.860 | 72578 |
|  | Elastix A | 4.724 | 3.838 | 1.225 | 3.808 | 9.028 | 96340 |
|  | Elastix B | 4.311 | 3.889 | 1.000 | 3.464 | 7.778 | 116388 |
| Tefor | ANTs A | 4.196 | 3.536 | 1.000 | 3.464 | 7.906 | 128050 |
|  | ANTs B | 4.670 | 3.621 | 1.414 | 3.808 | 8.746 | 110182 |
|  | ANTs C | 4.626 | 3.545 | 1.225 | 3.808 | 8.775 | 123050 |
|  | CMTK A | 6.937 | 8.519 | 1.000 | 3.808 | 18.655 | 111368 |
|  | CMTK B | 7.278 | 9.252 | 1.225 | 3.873 | 19.028 | 106430 |
|  | CMTK C | 5.438 | 4.413 | 1.581 | 4.301 | 10.700 | 96924 |
|  | Elastix A | 4.248 | 3.632 | 1.000 | 3.464 | 8.031 | 118540 |
|  | Elastix B | 4.553 | 4.048 | 1.225 | 3.606 | 8.367 | 126070 |

**Table S11:**
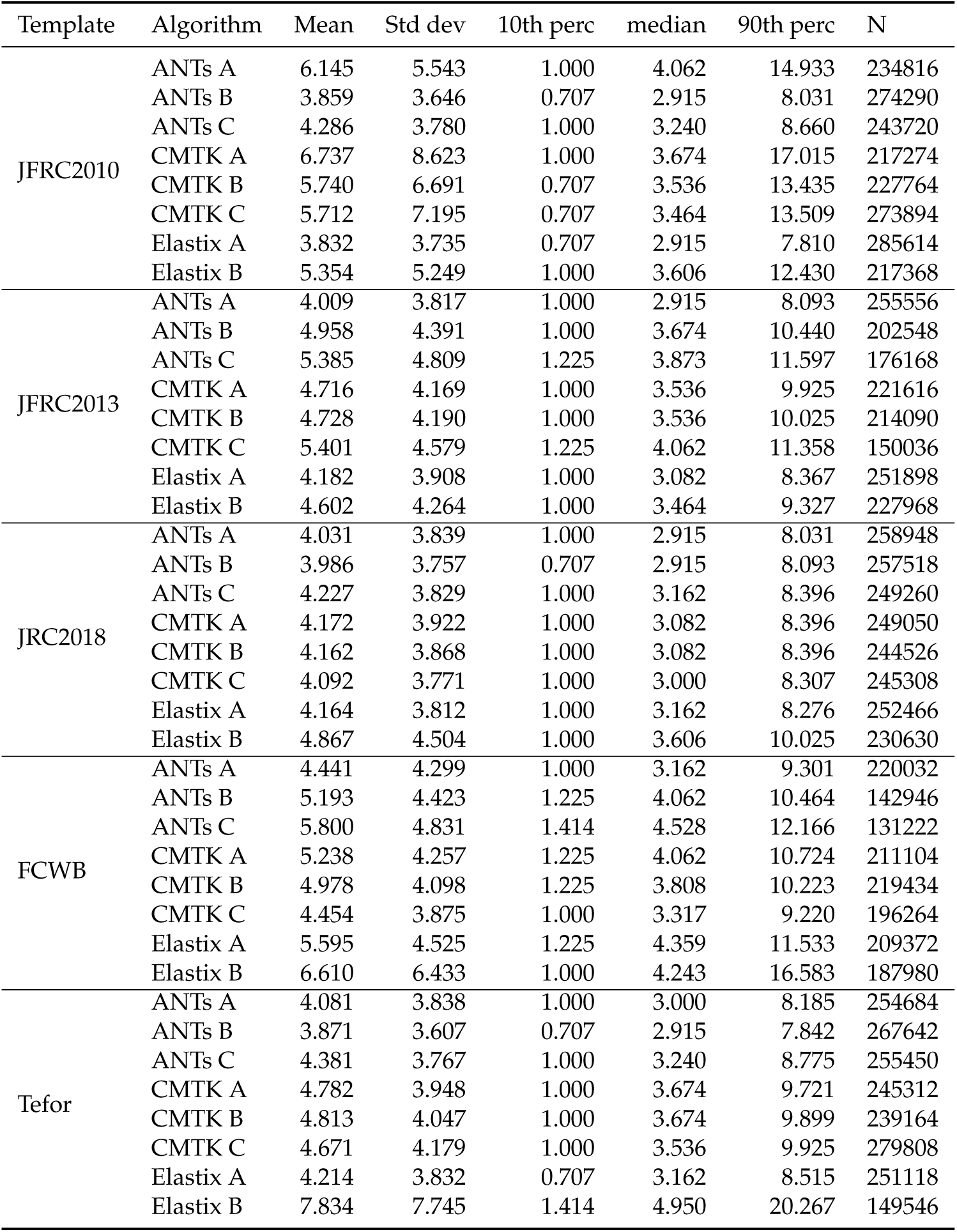
MB_ML_R : medial lobe of adult mushroom body

| Template | Algorithm | Mean | Std dev | 10th perc | median | 90th perc | N |
| --- | --- | --- | --- | --- | --- | --- | --- |
| JFRC2010 | ANTs A | 6.145 | 5.543 | 1.000 | 4.062 | 14.933 | 234816 |
|  | ANTs B | 3.859 | 3.646 | 0.707 | 2.915 | 8.031 | 274290 |
|  | ANTs C | 4.286 | 3.780 | 1.000 | 3.240 | 8.660 | 243720 |
|  | CMTK A | 6.737 | 8.623 | 1.000 | 3.674 | 17.015 | 217274 |
|  | CMTK B | 5.740 | 6.691 | 0.707 | 3.536 | 13.435 | 227764 |
|  | CMTK C | 5.712 | 7.195 | 0.707 | 3.464 | 13.509 | 273894 |
|  | Elastix A | 3.832 | 3.735 | 0.707 | 2.915 | 7.810 | 285614 |
|  | Elastix B | 5.354 | 5.249 | 1.000 | 3.606 | 12.430 | 217368 |
| JFRC2013 | ANTs A | 4.009 | 3.817 | 1.000 | 2.915 | 8.093 | 255556 |
|  | ANTs B | 4.958 | 4.391 | 1.000 | 3.674 | 10.440 | 202548 |
|  | ANTs C | 5.385 | 4.809 | 1.225 | 3.873 | 11.597 | 176168 |
|  | CMTK A | 4.716 | 4.169 | 1.000 | 3.536 | 9.925 | 221616 |
|  | CMTK B | 4.728 | 4.190 | 1.000 | 3.536 | 10.025 | 214090 |
|  | CMTK C | 5.401 | 4.579 | 1.225 | 4.062 | 11.358 | 150036 |
|  | Elastix A | 4.182 | 3.908 | 1.000 | 3.082 | 8.367 | 251898 |
|  | Elastix B | 4.602 | 4.264 | 1.000 | 3.464 | 9.327 | 227968 |
| JRC2018 | ANTs A | 4.031 | 3.839 | 1.000 | 2.915 | 8.031 | 258948 |
|  | ANTs B | 3.986 | 3.757 | 0.707 | 2.915 | 8.093 | 257518 |
|  | ANTs C | 4.227 | 3.829 | 1.000 | 3.162 | 8.396 | 249260 |
|  | CMTK A | 4.172 | 3.922 | 1.000 | 3.082 | 8.396 | 249050 |
|  | CMTK B | 4.162 | 3.868 | 1.000 | 3.082 | 8.396 | 244526 |
|  | CMTK C | 4.092 | 3.771 | 1.000 | 3.000 | 8.307 | 245308 |
|  | Elastix A | 4.164 | 3.812 | 1.000 | 3.162 | 8.276 | 252466 |
|  | Elastix B | 4.867 | 4.504 | 1.000 | 3.606 | 10.025 | 230630 |
| FCWB | ANTs A | 4.441 | 4.299 | 1.000 | 3.162 | 9.301 | 220032 |
|  | ANTs B | 5.193 | 4.423 | 1.225 | 4.062 | 10.464 | 142946 |
|  | ANTs C | 5.800 | 4.831 | 1.414 | 4.528 | 12.166 | 131222 |
|  | CMTK A | 5.238 | 4.257 | 1.225 | 4.062 | 10.724 | 211104 |
|  | CMTK B | 4.978 | 4.098 | 1.225 | 3.808 | 10.223 | 219434 |
|  | CMTK C | 4.454 | 3.875 | 1.000 | 3.317 | 9.220 | 196264 |
|  | Elastix A | 5.595 | 4.525 | 1.225 | 4.359 | 11.533 | 209372 |
|  | Elastix B | 6.610 | 6.433 | 1.000 | 4.243 | 16.583 | 187980 |
| Tefor | ANTs A | 4.081 | 3.838 | 1.000 | 3.000 | 8.185 | 254684 |
|  | ANTs B | 3.871 | 3.607 | 0.707 | 2.915 | 7.842 | 267642 |
|  | ANTs C | 4.381 | 3.767 | 1.000 | 3.240 | 8.775 | 255450 |
|  | CMTK A | 4.782 | 3.948 | 1.000 | 3.674 | 9.721 | 245312 |
|  | CMTK B | 4.813 | 4.047 | 1.000 | 3.674 | 9.899 | 239164 |
|  | CMTK C | 4.671 | 4.179 | 1.000 | 3.536 | 9.925 | 279808 |
|  | Elastix A | 4.214 | 3.832 | 0.707 | 3.162 | 8.515 | 251118 |
|  | Elastix B | 7.834 | 7.745 | 1.414 | 4.950 | 20.267 | 149546 |

**Table S12:** FLA_R : flange

| Template | Algorithm | Mean | Std dev | 10th perc | median | 90th perc | N |
| --- | --- | --- | --- | --- | --- | --- | --- |
| JFRC2010 | ANTs A | 5.474 | 8.066 | 0.707 | 2.915 | 11.597 | 354210 |
|  | ANTs B | 3.321 | 2.687 | 0.707 | 2.828 | 6.364 | 377092 |
|  | ANTs C | 3.267 | 2.787 | 0.707 | 2.646 | 6.442 | 419414 |
|  | CMTK A | 3.680 | 4.159 | 0.707 | 2.550 | 7.382 | 352410 |
|  | CMTK B | 3.637 | 4.142 | 0.707 | 2.550 | 7.280 | 361674 |
|  | CMTK C | 3.641 | 3.605 | 0.707 | 2.646 | 7.416 | 348282 |
|  | Elastix A | 2.724 | 2.391 | 0.707 | 2.236 | 5.339 | 368494 |
|  | Elastix B | 2.707 | 2.404 | 0.707 | 2.236 | 5.292 | 365130 |
| JFRC2013 | ANTs A | 2.676 | 2.259 | 0.707 | 2.121 | 5.339 | 399670 |
|  | ANTs B | 4.493 | 3.953 | 1.000 | 3.606 | 8.631 | 262524 |
|  | ANTs C | 4.066 | 3.673 | 1.000 | 3.240 | 7.681 | 272910 |
|  | CMTK A | 2.641 | 2.233 | 0.707 | 2.121 | 5.196 | 424002 |
|  | CMTK B | 2.628 | 2.202 | 0.707 | 2.121 | 5.196 | 410600 |
|  | CMTK C | 3.378 | 2.734 | 0.707 | 2.915 | 6.403 | 303820 |
|  | Elastix A | 2.547 | 2.201 | 0.707 | 2.121 | 5.000 | 364092 |
|  | Elastix B | 2.534 | 2.236 | 0.707 | 2.121 | 5.000 | 396174 |
| JRC2018 | ANTs A | 2.420 | 2.106 | 0.707 | 2.000 | 4.743 | 408766 |
|  | ANTs B | 3.128 | 2.713 | 0.707 | 2.550 | 6.205 | 384048 |
|  | ANTs C | 3.450 | 3.071 | 0.707 | 2.646 | 6.819 | 379300 |
|  | CMTK A | 2.459 | 2.190 | 0.707 | 2.121 | 4.899 | 449412 |
|  | CMTK B | 2.465 | 2.233 | 0.707 | 2.121 | 4.796 | 421730 |
|  | CMTK C | 2.471 | 2.173 | 0.707 | 2.121 | 4.950 | 402876 |
|  | Elastix A | 2.551 | 2.158 | 0.707 | 2.121 | 5.000 | 398878 |
|  | Elastix B | 2.616 | 2.482 | 0.707 | 2.121 | 5.148 | 389906 |
| FCWB | ANTs A | 2.524 | 2.465 | 0.707 | 2.121 | 5.000 | 422600 |
|  | ANTs B | 3.848 | 3.358 | 1.000 | 3.082 | 7.246 | 303022 |
|  | ANTs C | 3.835 | 3.391 | 1.000 | 3.000 | 7.348 | 316062 |
|  | CMTK A | 3.217 | 2.742 | 0.707 | 2.646 | 6.325 | 393608 |
|  | CMTK B | 3.221 | 2.928 | 0.707 | 2.646 | 6.124 | 391004 |
|  | CMTK C | 3.130 | 2.665 | 0.707 | 2.550 | 6.124 | 336788 |
|  | Elastix A | 4.111 | 4.140 | 0.707 | 3.240 | 7.937 | 305028 |
|  | Elastix B | 3.161 | 2.643 | 0.707 | 2.550 | 6.364 | 371708 |
| Tefor | ANTs A | 2.495 | 2.341 | 0.707 | 2.121 | 5.000 | 426156 |
|  | ANTs B | 3.402 | 2.944 | 0.707 | 2.828 | 6.557 | 366804 |
|  | ANTs C | 3.480 | 3.206 | 0.707 | 2.646 | 6.745 | 414964 |
|  | CMTK A | 3.079 | 2.677 | 0.707 | 2.449 | 6.364 | 379488 |
|  | CMTK B | 3.319 | 3.008 | 0.707 | 2.550 | 6.964 | 363494 |
|  | CMTK C | 3.703 | 3.878 | 0.707 | 2.646 | 6.964 | 363462 |
|  | Elastix A | 2.881 | 2.397 | 0.707 | 2.345 | 5.657 | 369246 |
|  | Elastix B | 3.062 | 2.759 | 0.707 | 2.550 | 6.000 | 344700 |

**Table S13:** LOP_R : lobula plate

| Template | Algorithm | Mean | Std dev | 10th perc | median | 90th perc | N |
| --- | --- | --- | --- | --- | --- | --- | --- |
| JFRC2010 | ANTs A | 8.661 | 6.730 | 2.345 | 6.928 | 17.132 | 475654 |
|  | ANTs B | 6.802 | 4.843 | 2.121 | 5.745 | 12.767 | 388000 |
|  | ANTs C | 7.298 | 4.926 | 2.121 | 6.325 | 13.583 | 458102 |
|  | CMTK A | 8.485 | 5.918 | 2.236 | 7.106 | 16.628 | 509662 |
|  | CMTK B | 7.640 | 5.088 | 2.121 | 6.519 | 14.832 | 460728 |
|  | CMTK C | 7.212 | 4.914 | 2.121 | 6.083 | 13.946 | 497312 |
|  | Elastix A | 5.645 | 4.406 | 1.581 | 4.528 | 10.863 | 368684 |
|  | Elastix B | 5.415 | 4.242 | 1.581 | 4.472 | 10.296 | 403768 |
| JFRC2013 | ANTs A | 6.053 | 5.289 | 1.581 | 4.743 | 11.937 | 442126 |
|  | ANTs B | 6.657 | 5.179 | 1.732 | 5.385 | 13.000 | 445352 |
|  | ANTs C | 7.051 | 5.202 | 2.000 | 5.874 | 13.528 | 444334 |
|  | CMTK A | 7.234 | 6.544 | 1.581 | 5.196 | 15.182 | 426364 |
|  | CMTK B | 7.264 | 6.489 | 1.732 | 5.339 | 14.866 | 424412 |
|  | CMTK C | 6.756 | 4.706 | 2.121 | 5.701 | 12.767 | 449680 |
|  | Elastix A | 5.801 | 4.865 | 1.581 | 4.528 | 11.424 | 450286 |
|  | Elastix B | 5.711 | 4.972 | 1.414 | 4.472 | 11.336 | 482592 |
| JRC2018 | ANTs A | 5.652 | 4.929 | 1.581 | 4.359 | 11.068 | 439706 |
|  | ANTs B | 5.683 | 4.802 | 1.581 | 4.528 | 11.000 | 435238 |
|  | ANTs C | 6.006 | 4.981 | 1.581 | 4.743 | 11.832 | 457370 |
|  | CMTK A | 5.610 | 4.934 | 1.414 | 4.359 | 11.045 | 468788 |
|  | CMTK B | 5.604 | 4.911 | 1.581 | 4.359 | 10.909 | 458608 |
|  | CMTK C | 5.760 | 5.062 | 1.581 | 4.528 | 11.068 | 448426 |
|  | Elastix A | 5.604 | 4.814 | 1.581 | 4.359 | 10.886 | 437394 |
|  | Elastix B | 5.545 | 5.104 | 1.581 | 4.359 | 10.724 | 469178 |
| FCWB | ANTs A | 6.767 | 5.505 | 1.732 | 5.431 | 13.153 | 411806 |
|  | ANTs B | 6.731 | 4.922 | 2.000 | 5.568 | 12.748 | 359238 |
|  | ANTs C | 6.791 | 4.888 | 2.121 | 5.701 | 12.767 | 392490 |
|  | CMTK A | 7.542 | 5.486 | 2.121 | 6.245 | 14.387 | 416848 |
|  | CMTK B | 7.397 | 5.463 | 2.121 | 6.083 | 14.160 | 391324 |
|  | CMTK C | 7.686 | 5.412 | 2.236 | 6.364 | 14.866 | 404082 |
|  | Elastix A | 8.417 | 7.709 | 2.121 | 6.671 | 15.827 | 426276 |
|  | Elastix B | 7.598 | 9.293 | 1.732 | 5.568 | 13.528 | 391602 |
| Tefor | ANTs A | 5.807 | 5.075 | 1.581 | 4.528 | 11.358 | 436920 |
|  | ANTs B | 6.259 | 4.860 | 1.732 | 5.148 | 11.790 | 378356 |
|  | ANTs C | 6.948 | 5.085 | 2.000 | 5.831 | 13.304 | 416008 |
|  | CMTK A | 7.139 | 5.660 | 1.732 | 5.701 | 14.230 | 471104 |
|  | CMTK B | 7.117 | 5.625 | 1.732 | 5.701 | 13.946 | 455220 |
|  | CMTK C | 6.553 | 5.365 | 1.732 | 5.196 | 12.430 | 409754 |
|  | Elastix A | 5.593 | 4.687 | 1.581 | 4.416 | 10.886 | 376232 |
|  | Elastix B | 5.431 | 4.587 | 1.581 | 4.301 | 10.630 | 384092 |

**Table S14:** EB : ellipsoid body

| Template | Algorithm | Mean | Std dev | 10th perc | median | 90th perc | N |
| --- | --- | --- | --- | --- | --- | --- | --- |
| JFRC2010 | ANTs A | 5.630 | 5.639 | 1.414 | 3.808 | 13.675 | 239966 |
|  | ANTs B | 3.587 | 2.022 | 1.225 | 3.240 | 6.245 | 267694 |
|  | ANTs C | 3.702 | 2.098 | 1.225 | 3.464 | 6.481 | 259974 |
|  | CMTK A | 6.327 | 7.018 | 1.581 | 4.359 | 11.336 | 246854 |
|  | CMTK B | 5.277 | 4.698 | 1.581 | 4.062 | 9.849 | 244064 |
|  | CMTK C | 5.338 | 4.636 | 1.581 | 4.183 | 9.721 | 246128 |
|  | Elastix A | 3.402 | 1.932 | 1.000 | 3.082 | 6.000 | 293972 |
|  | Elastix B | 4.024 | 2.695 | 1.225 | 3.536 | 7.517 | 293234 |
| JFRC2013 | ANTs A | 3.489 | 1.992 | 1.225 | 3.240 | 6.083 | 269494 |
|  | ANTs B | 3.909 | 2.298 | 1.414 | 3.606 | 6.964 | 287216 |
|  | ANTs C | 3.825 | 2.245 | 1.225 | 3.536 | 6.819 | 279940 |
|  | CMTK A | 3.627 | 2.071 | 1.225 | 3.317 | 6.364 | 263754 |
|  | CMTK B | 3.718 | 2.138 | 1.225 | 3.464 | 6.557 | 271152 |
|  | CMTK C | 3.374 | 2.116 | 1.000 | 2.915 | 6.205 | 355066 |
|  | Elastix A | 3.543 | 1.984 | 1.225 | 3.240 | 6.124 | 281728 |
|  | Elastix B | 3.470 | 1.948 | 1.225 | 3.240 | 6.083 | 293524 |
| JRC2018 | ANTs A | 3.477 | 1.982 | 1.225 | 3.162 | 6.083 | 269700 |
|  | ANTs B | 3.547 | 1.995 | 1.225 | 3.240 | 6.164 | 266400 |
|  | ANTs C | 3.597 | 2.017 | 1.225 | 3.317 | 6.245 | 266182 |
|  | CMTK A | 3.601 | 2.110 | 1.225 | 3.240 | 6.403 | 242150 |
|  | CMTK B | 3.603 | 2.092 | 1.225 | 3.240 | 6.364 | 243970 |
|  | CMTK C | 3.583 | 2.018 | 1.225 | 3.240 | 6.205 | 256428 |
|  | Elastix A | 3.558 | 1.989 | 1.225 | 3.240 | 6.124 | 265772 |
|  | Elastix B | 3.570 | 2.055 | 1.225 | 3.240 | 6.325 | 276532 |
| FCWB | ANTs A | 3.468 | 1.976 | 1.225 | 3.162 | 6.042 | 265968 |
|  | ANTs B | 3.525 | 2.414 | 1.000 | 3.000 | 6.856 | 373646 |
|  | ANTs C | 3.636 | 2.399 | 1.000 | 3.082 | 6.964 | 345156 |
|  | CMTK A | 5.688 | 4.228 | 1.581 | 4.528 | 11.683 | 302648 |
|  | CMTK B | 5.225 | 3.956 | 1.414 | 4.123 | 10.817 | 302658 |
|  | CMTK C | 3.877 | 2.675 | 1.000 | 3.240 | 7.583 | 330860 |
|  | Elastix A | 4.572 | 3.166 | 1.414 | 3.873 | 8.746 | 384774 |
|  | Elastix B | 4.932 | 3.963 | 1.414 | 3.808 | 9.721 | 247436 |
| Tefor | ANTs A | 3.474 | 1.984 | 1.000 | 3.162 | 6.083 | 267228 |
|  | ANTs B | 3.589 | 2.025 | 1.225 | 3.317 | 6.205 | 258562 |
|  | ANTs C | 3.596 | 2.055 | 1.225 | 3.317 | 6.364 | 272178 |
|  | CMTK A | 4.521 | 2.941 | 1.414 | 3.873 | 8.396 | 240954 |
|  | CMTK B | 4.452 | 2.870 | 1.414 | 3.808 | 8.307 | 242188 |
|  | CMTK C | 4.120 | 2.555 | 1.414 | 3.606 | 7.550 | 302688 |
|  | Elastix A | 3.495 | 2.007 | 1.225 | 3.162 | 6.124 | 281684 |
|  | Elastix B | 4.676 | 3.586 | 1.414 | 3.808 | 9.220 | 260680 |

**Table S15:** AL_R : adult antennal lobe

| Template | Algorithm | Mean | Std dev | 10th perc | median | 90th perc | N |
| --- | --- | --- | --- | --- | --- | --- | --- |
| JFRC2010 | ANTs A | 5.255 | 6.244 | 1.225 | 3.464 | 10.817 | 1405498 |
|  | ANTs B | 3.445 | 2.357 | 1.000 | 3.000 | 6.000 | 1443546 |
|  | ANTs C | 3.525 | 2.492 | 1.225 | 3.082 | 6.083 | 1542696 |
|  | CMTK A | 4.328 | 4.241 | 1.225 | 3.240 | 7.649 | 1383128 |
|  | CMTK B | 4.419 | 4.625 | 1.225 | 3.240 | 7.746 | 1366692 |
|  | CMTK C | 4.983 | 5.726 | 1.225 | 3.464 | 9.110 | 1307010 |
|  | Elastix A | 3.232 | 2.074 | 1.000 | 2.915 | 5.568 | 1456306 |
|  | Elastix B | 3.285 | 2.093 | 1.000 | 3.000 | 5.701 | 1500946 |
| JFRC2013 | ANTs A | 3.215 | 2.012 | 1.000 | 2.915 | 5.568 | 1517212 |
|  | ANTs B | 4.297 | 3.658 | 1.225 | 3.317 | 8.246 | 1094568 |
|  | ANTs C | 4.446 | 4.027 | 1.225 | 3.317 | 8.746 | 1157590 |
|  | CMTK A | 3.505 | 2.526 | 1.000 | 3.000 | 6.083 | 1436332 |
|  | CMTK B | 3.548 | 2.519 | 1.000 | 3.082 | 6.205 | 1403924 |
|  | CMTK C | 3.778 | 2.778 | 1.225 | 3.162 | 6.964 | 1085438 |
|  | Elastix A | 3.216 | 2.060 | 1.000 | 2.915 | 5.523 | 1497794 |
|  | Elastix B | 3.194 | 2.043 | 1.000 | 2.915 | 5.523 | 1549406 |
| JRC2018 | ANTs A | 3.144 | 1.980 | 1.000 | 2.915 | 5.431 | 1549380 |
|  | ANTs B | 3.343 | 2.076 | 1.000 | 3.000 | 5.745 | 1520126 |
|  | ANTs C | 3.473 | 2.145 | 1.225 | 3.162 | 6.083 | 1530696 |
|  | CMTK A | 3.152 | 1.987 | 1.000 | 2.915 | 5.431 | 1545220 |
|  | CMTK B | 3.165 | 1.989 | 1.000 | 2.915 | 5.431 | 1555946 |
|  | CMTK C | 3.213 | 2.012 | 1.000 | 2.915 | 5.523 | 1561296 |
|  | Elastix A | 3.220 | 2.019 | 1.000 | 2.915 | 5.523 | 1557730 |
|  | Elastix B | 3.201 | 2.031 | 1.000 | 2.915 | 5.523 | 1547092 |
| FCWB | ANTs A | 3.468 | 2.336 | 1.000 | 3.082 | 6.042 | 1524310 |
|  | ANTs B | 3.959 | 3.016 | 1.225 | 3.240 | 7.280 | 1045354 |
|  | ANTs C | 4.367 | 3.836 | 1.225 | 3.464 | 8.246 | 1108210 |
|  | CMTK A | 3.575 | 2.365 | 1.225 | 3.162 | 6.205 | 1536540 |
|  | CMTK B | 3.578 | 2.458 | 1.225 | 3.162 | 6.205 | 1510018 |
|  | CMTK C | 3.720 | 2.562 | 1.225 | 3.240 | 6.557 | 1328746 |
|  | Elastix A | 3.644 | 2.546 | 1.225 | 3.240 | 6.364 | 1463492 |
|  | Elastix B | 3.451 | 2.230 | 1.225 | 3.082 | 6.083 | 1487654 |
| Tefor | ANTs A | 3.175 | 2.013 | 1.000 | 2.915 | 5.523 | 1543712 |
|  | ANTs B | 3.390 | 2.263 | 1.000 | 3.000 | 5.874 | 1526548 |
|  | ANTs C | 3.696 | 3.147 | 1.225 | 3.162 | 6.205 | 1584950 |
|  | CMTK A | 3.725 | 2.591 | 1.225 | 3.240 | 6.708 | 1540136 |
|  | CMTK B | 3.703 | 2.566 | 1.225 | 3.240 | 6.671 | 1554432 |
|  | CMTK C | 3.884 | 3.081 | 1.225 | 3.240 | 6.964 | 1531560 |
|  | Elastix A | 3.291 | 2.106 | 1.000 | 3.000 | 5.701 | 1509564 |
|  | Elastix B | 3.646 | 2.421 | 1.225 | 3.240 | 6.519 | 1498242 |

**Table S16:** ME_R : medulla

| Template | Algorithm | Mean | Std dev | 10th perc | median | 90th perc | N |
| --- | --- | --- | --- | --- | --- | --- | --- |
| JFRC2010 | ANTs A | 6.367 | 6.837 | 1.581 | 4.743 | 11.402 | 3492090 |
|  | ANTs B | 5.040 | 4.515 | 1.581 | 4.243 | 8.860 | 2962300 |
|  | ANTs C | 5.358 | 4.467 | 1.581 | 4.528 | 9.513 | 3300890 |
|  | CMTK A | 5.502 | 4.495 | 1.581 | 4.637 | 10.050 | 3257940 |
|  | CMTK B | 5.198 | 4.364 | 1.581 | 4.528 | 9.220 | 3184846 |
|  | CMTK C | 5.766 | 4.471 | 1.732 | 5.000 | 10.271 | 3006892 |
|  | Elastix A | 4.629 | 4.184 | 1.581 | 4.062 | 8.031 | 3067542 |
|  | Elastix B | 4.741 | 4.283 | 1.581 | 4.123 | 8.246 | 3114026 |
| JFRC2013 | ANTs A | 4.812 | 4.251 | 1.581 | 4.123 | 8.544 | 3210516 |
|  | ANTs B | 5.407 | 4.875 | 1.581 | 4.528 | 9.721 | 3020702 |
|  | ANTs C | 5.796 | 4.699 | 1.732 | 5.000 | 10.344 | 3127004 |
|  | CMTK A | 5.781 | 5.388 | 1.581 | 4.690 | 10.025 | 2897718 |
|  | CMTK B | 6.030 | 5.700 | 1.732 | 4.796 | 10.416 | 2829810 |
|  | CMTK C | 6.238 | 5.048 | 2.121 | 5.385 | 10.630 | 2760140 |
|  | Elastix A | 4.653 | 4.131 | 1.581 | 4.062 | 8.185 | 3280756 |
|  | Elastix B | 4.628 | 4.071 | 1.414 | 4.062 | 8.124 | 3247838 |
| JRC2018 | ANTs A | 4.601 | 4.073 | 1.414 | 4.000 | 8.093 | 3275738 |
|  | ANTs B | 5.017 | 4.679 | 1.581 | 4.183 | 8.775 | 3215812 |
|  | ANTs C | 5.493 | 5.040 | 1.581 | 4.528 | 9.899 | 3275980 |
|  | CMTK A | 4.632 | 4.189 | 1.414 | 4.062 | 8.093 | 3215146 |
|  | CMTK B | 4.670 | 4.144 | 1.581 | 4.062 | 8.155 | 3236520 |
|  | CMTK C | 4.832 | 4.261 | 1.581 | 4.183 | 8.515 | 3271916 |
|  | Elastix A | 4.615 | 4.321 | 1.581 | 4.062 | 7.937 | 3205366 |
|  | Elastix B | 4.653 | 4.120 | 1.581 | 4.062 | 8.093 | 3201122 |
| FCWB | ANTs A | 4.795 | 4.176 | 1.581 | 4.123 | 8.426 | 3302824 |
|  | ANTs B | 5.485 | 4.647 | 1.581 | 4.690 | 9.644 | 3156232 |
|  | ANTs C | 6.412 | 5.015 | 2.000 | 5.523 | 11.446 | 3176428 |
|  | CMTK A | 6.061 | 5.185 | 1.732 | 5.000 | 11.068 | 3333848 |
|  | CMTK B | 5.863 | 5.094 | 1.732 | 4.796 | 10.583 | 3310852 |
|  | CMTK C | 6.024 | 4.836 | 1.732 | 5.196 | 10.770 | 3069468 |
|  | Elastix A | 6.627 | 6.804 | 2.000 | 5.385 | 12.490 | 3344776 |
|  | Elastix B | 5.859 | 6.269 | 1.581 | 4.690 | 10.886 | 3331994 |
| Tefor | ANTs A | 4.633 | 4.197 | 1.414 | 4.000 | 8.185 | 3254842 |
|  | ANTs B | 5.102 | 4.661 | 1.581 | 4.183 | 9.055 | 2995846 |
|  | ANTs C | 5.812 | 4.841 | 1.581 | 4.950 | 10.392 | 3193460 |
|  | CMTK A | 6.927 | 7.527 | 1.581 | 4.743 | 14.370 | 2953692 |
|  | CMTK B | 6.898 | 7.427 | 1.581 | 4.743 | 14.177 | 2969518 |
|  | CMTK C | 6.883 | 8.348 | 1.581 | 4.743 | 12.124 | 2878476 |
|  | Elastix A | 4.589 | 4.112 | 1.581 | 4.000 | 8.031 | 3103026 |
|  | Elastix B | 4.674 | 4.183 | 1.581 | 4.062 | 8.185 | 3084188 |

**Table S17:**
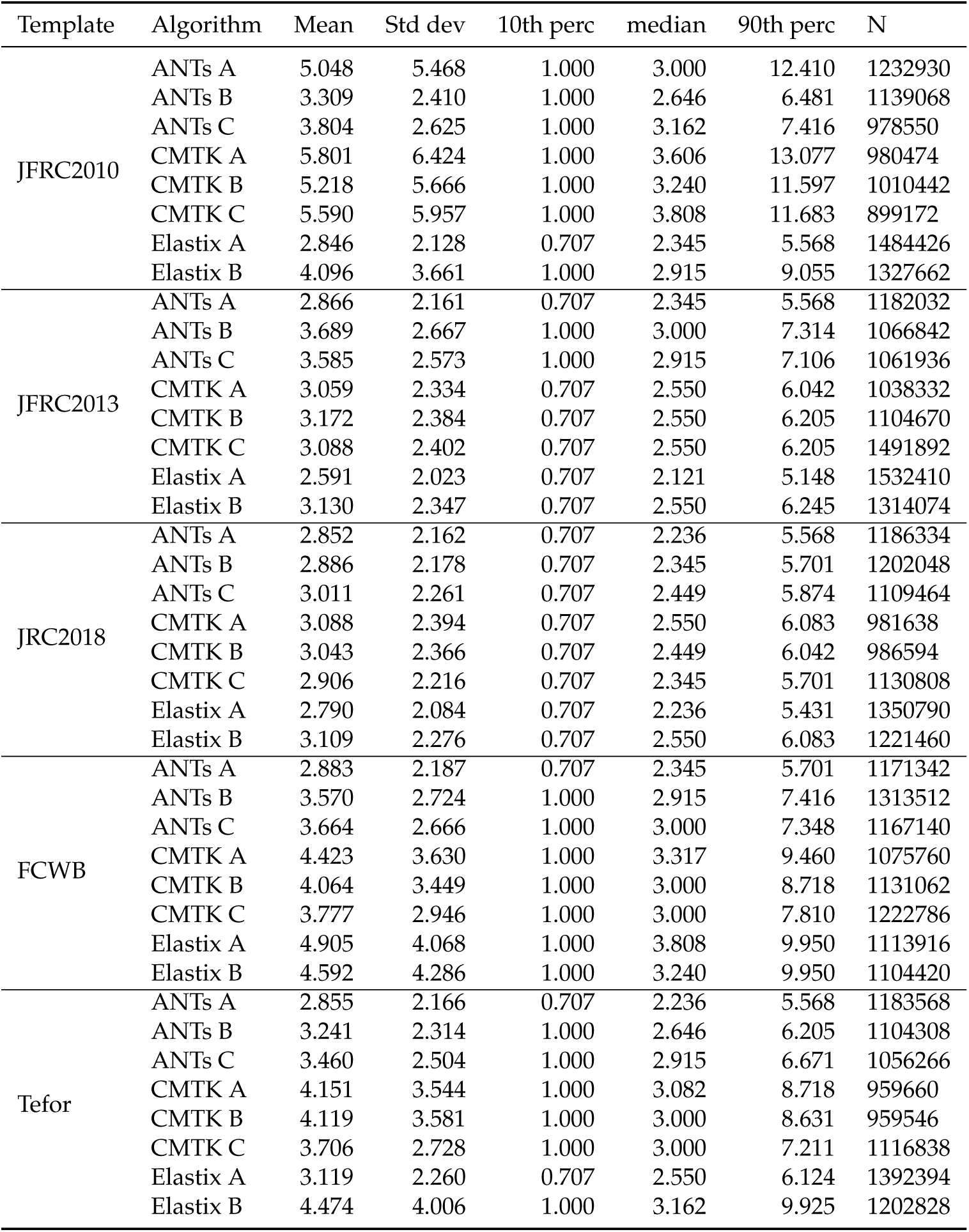
FB : fan-shaped body

| Template | Algorithm | Mean | Std dev | 10th perc | median | 90th perc | N |
| --- | --- | --- | --- | --- | --- | --- | --- |
| JFRC2010 | ANTs A | 5.048 | 5.468 | 1.000 | 3.000 | 12.410 | 1232930 |
|  | ANTs B | 3.309 | 2.410 | 1.000 | 2.646 | 6.481 | 1139068 |
|  | ANTs C | 3.804 | 2.625 | 1.000 | 3.162 | 7.416 | 978550 |
|  | CMTK A | 5.801 | 6.424 | 1.000 | 3.606 | 13.077 | 980474 |
|  | CMTK B | 5.218 | 5.666 | 1.000 | 3.240 | 11.597 | 1010442 |
|  | CMTK C | 5.590 | 5.957 | 1.000 | 3.808 | 11.683 | 899172 |
|  | Elastix A | 2.846 | 2.128 | 0.707 | 2.345 | 5.568 | 1484426 |
|  | Elastix B | 4.096 | 3.661 | 1.000 | 2.915 | 9.055 | 1327662 |
| JFRC2013 | ANTs A | 2.866 | 2.161 | 0.707 | 2.345 | 5.568 | 1182032 |
|  | ANTs B | 3.689 | 2.667 | 1.000 | 3.000 | 7.314 | 1066842 |
|  | ANTs C | 3.585 | 2.573 | 1.000 | 2.915 | 7.106 | 1061936 |
|  | CMTK A | 3.059 | 2.334 | 0.707 | 2.550 | 6.042 | 1038332 |
|  | CMTK B | 3.172 | 2.384 | 0.707 | 2.550 | 6.205 | 1104670 |
|  | CMTK C | 3.088 | 2.402 | 0.707 | 2.550 | 6.205 | 1491892 |
|  | Elastix A | 2.591 | 2.023 | 0.707 | 2.121 | 5.148 | 1532410 |
|  | Elastix B | 3.130 | 2.347 | 0.707 | 2.550 | 6.245 | 1314074 |
| JRC2018 | ANTs A | 2.852 | 2.162 | 0.707 | 2.236 | 5.568 | 1186334 |
|  | ANTs B | 2.886 | 2.178 | 0.707 | 2.345 | 5.701 | 1202048 |
|  | ANTs C | 3.011 | 2.261 | 0.707 | 2.449 | 5.874 | 1109464 |
|  | CMTK A | 3.088 | 2.394 | 0.707 | 2.550 | 6.083 | 981638 |
|  | CMTK B | 3.043 | 2.366 | 0.707 | 2.449 | 6.042 | 986594 |
|  | CMTK C | 2.906 | 2.216 | 0.707 | 2.345 | 5.701 | 1130808 |
|  | Elastix A | 2.790 | 2.084 | 0.707 | 2.236 | 5.431 | 1350790 |
|  | Elastix B | 3.109 | 2.276 | 0.707 | 2.550 | 6.083 | 1221460 |
| FCWB | ANTs A | 2.883 | 2.187 | 0.707 | 2.345 | 5.701 | 1171342 |
|  | ANTs B | 3.570 | 2.724 | 1.000 | 2.915 | 7.416 | 1313512 |
|  | ANTs C | 3.664 | 2.666 | 1.000 | 3.000 | 7.348 | 1167140 |
|  | CMTK A | 4.423 | 3.630 | 1.000 | 3.317 | 9.460 | 1075760 |
|  | CMTK B | 4.064 | 3.449 | 1.000 | 3.000 | 8.718 | 1131062 |
|  | CMTK C | 3.777 | 2.946 | 1.000 | 3.000 | 7.810 | 1222786 |
|  | Elastix A | 4.905 | 4.068 | 1.000 | 3.808 | 9.950 | 1113916 |
|  | Elastix B | 4.592 | 4.286 | 1.000 | 3.240 | 9.950 | 1104420 |
| Tefor | ANTs A | 2.855 | 2.166 | 0.707 | 2.236 | 5.568 | 1183568 |
|  | ANTs B | 3.241 | 2.314 | 1.000 | 2.646 | 6.205 | 1104308 |
|  | ANTs C | 3.460 | 2.504 | 1.000 | 2.915 | 6.671 | 1056266 |
|  | CMTK A | 4.151 | 3.544 | 1.000 | 3.082 | 8.718 | 959660 |
|  | CMTK B | 4.119 | 3.581 | 1.000 | 3.000 | 8.631 | 959546 |
|  | CMTK C | 3.706 | 2.728 | 1.000 | 3.000 | 7.211 | 1116838 |
|  | Elastix A | 3.119 | 2.260 | 0.707 | 2.550 | 6.124 | 1392394 |
|  | Elastix B | 4.474 | 4.006 | 1.000 | 3.162 | 9.925 | 1202828 |

**Table S18:**
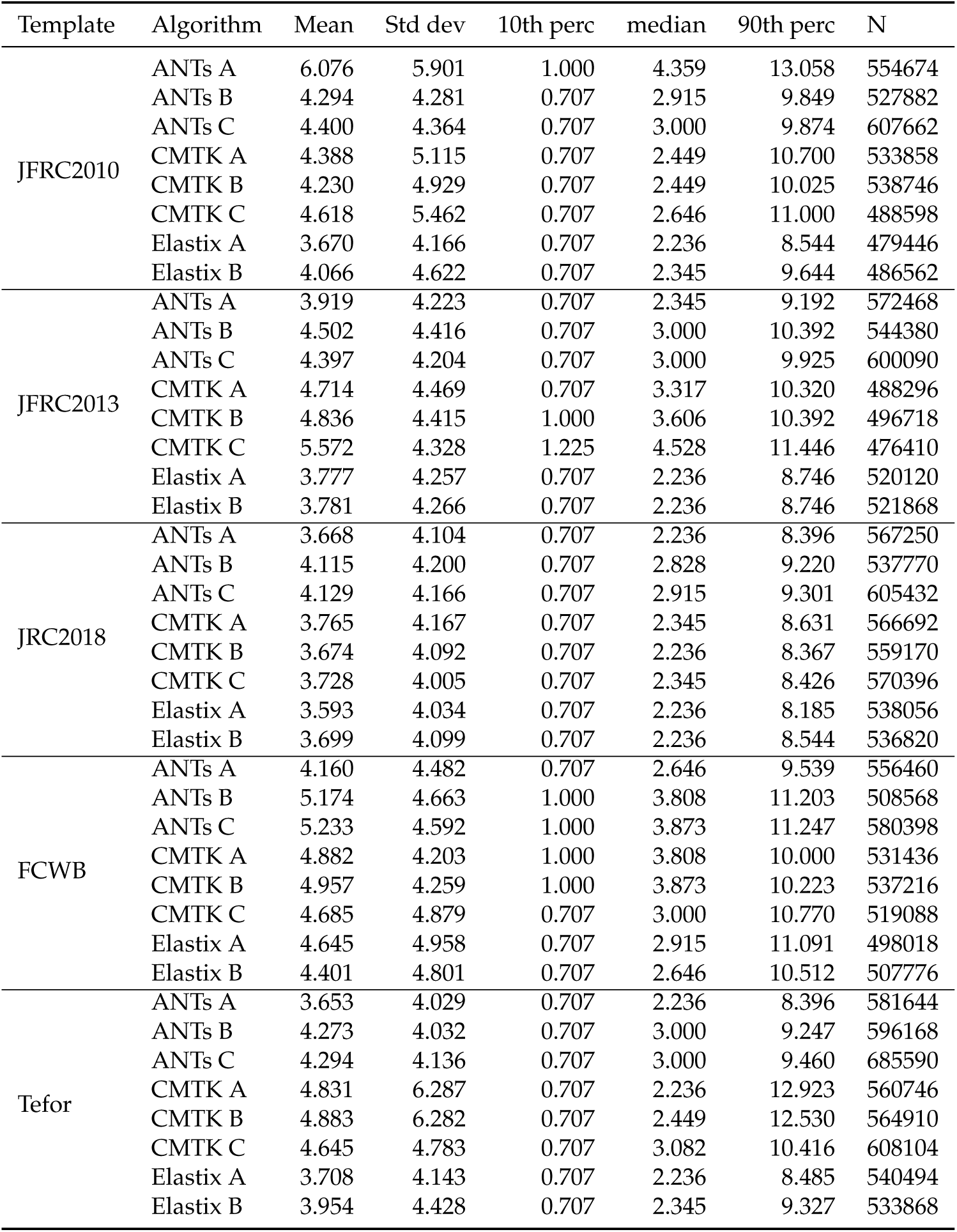
SLP_R : superior lateral protocerebrum

**Table S19:**
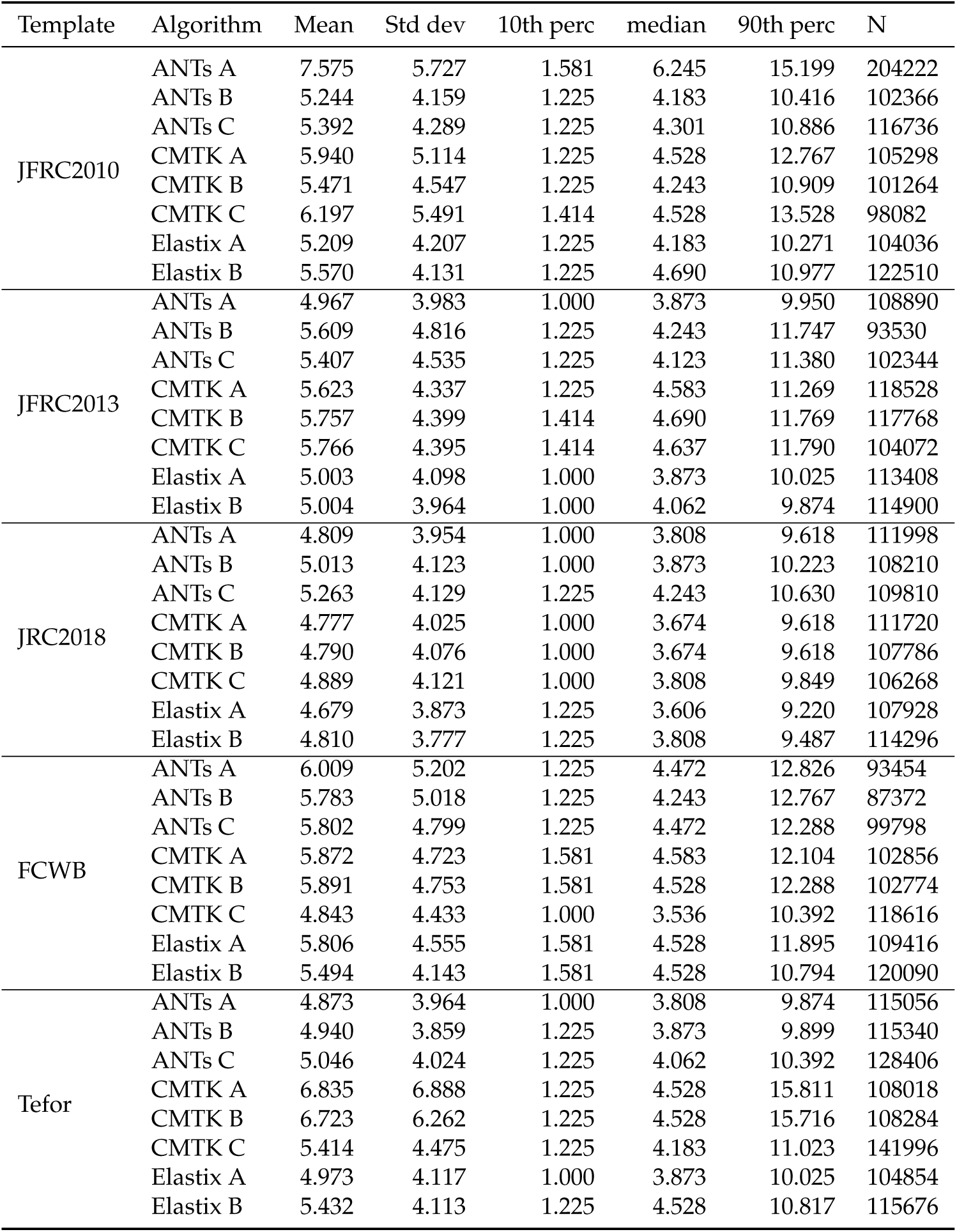
SIP_R : superior intermediate protocerebrum

| Template | Algorithm | Mean | Std dev | 10th perc | median | 90th perc | N |
| --- | --- | --- | --- | --- | --- | --- | --- |
| JFRC2010 | ANTs A | 7.575 | 5.727 | 1.581 | 6.245 | 15.199 | 204222 |
|  | ANTs B | 5.244 | 4.159 | 1.225 | 4.183 | 10.416 | 102366 |
|  | ANTs C | 5.392 | 4.289 | 1.225 | 4.301 | 10.886 | 116736 |
|  | CMTK A | 5.940 | 5.114 | 1.225 | 4.528 | 12.767 | 105298 |
|  | CMTK B | 5.471 | 4.547 | 1.225 | 4.243 | 10.909 | 101264 |
|  | CMTK C | 6.197 | 5.491 | 1.414 | 4.528 | 13.528 | 98082 |
|  | Elastix A | 5.209 | 4.207 | 1.225 | 4.183 | 10.271 | 104036 |
|  | Elastix B | 5.570 | 4.131 | 1.225 | 4.690 | 10.977 | 122510 |
| JFRC2013 | ANTs A | 4.967 | 3.983 | 1.000 | 3.873 | 9.950 | 108890 |
|  | ANTs B | 5.609 | 4.816 | 1.225 | 4.243 | 11.747 | 93530 |
|  | ANTs C | 5.407 | 4.535 | 1.225 | 4.123 | 11.380 | 102344 |
|  | CMTK A | 5.623 | 4.337 | 1.225 | 4.583 | 11.269 | 118528 |
|  | CMTK B | 5.757 | 4.399 | 1.414 | 4.690 | 11.769 | 117768 |
|  | CMTK C | 5.766 | 4.395 | 1.414 | 4.637 | 11.790 | 104072 |
|  | Elastix A | 5.003 | 4.098 | 1.000 | 3.873 | 10.025 | 113408 |
|  | Elastix B | 5.004 | 3.964 | 1.000 | 4.062 | 9.874 | 114900 |
| JRC2018 | ANTs A | 4.809 | 3.954 | 1.000 | 3.808 | 9.618 | 111998 |
|  | ANTs B | 5.013 | 4.123 | 1.000 | 3.873 | 10.223 | 108210 |
|  | ANTs C | 5.263 | 4.129 | 1.225 | 4.243 | 10.630 | 109810 |
|  | CMTK A | 4.777 | 4.025 | 1.000 | 3.674 | 9.618 | 111720 |
|  | CMTK B | 4.790 | 4.076 | 1.000 | 3.674 | 9.618 | 107786 |
|  | CMTK C | 4.889 | 4.121 | 1.000 | 3.808 | 9.849 | 106268 |
|  | Elastix A | 4.679 | 3.873 | 1.225 | 3.606 | 9.220 | 107928 |
|  | Elastix B | 4.810 | 3.777 | 1.225 | 3.808 | 9.487 | 114296 |
| FCWB | ANTs A | 6.009 | 5.202 | 1.225 | 4.472 | 12.826 | 93454 |
|  | ANTs B | 5.783 | 5.018 | 1.225 | 4.243 | 12.767 | 87372 |
|  | ANTs C | 5.802 | 4.799 | 1.225 | 4.472 | 12.288 | 99798 |
|  | CMTK A | 5.872 | 4.723 | 1.581 | 4.583 | 12.104 | 102856 |
|  | CMTK B | 5.891 | 4.753 | 1.581 | 4.528 | 12.288 | 102774 |
|  | CMTK C | 4.843 | 4.433 | 1.000 | 3.536 | 10.392 | 118616 |
|  | Elastix A | 5.806 | 4.555 | 1.581 | 4.528 | 11.895 | 109416 |
|  | Elastix B | 5.494 | 4.143 | 1.581 | 4.528 | 10.794 | 120090 |
| Tefor | ANTs A | 4.873 | 3.964 | 1.000 | 3.808 | 9.874 | 115056 |
|  | ANTs B | 4.940 | 3.859 | 1.225 | 3.873 | 9.899 | 115340 |
|  | ANTs C | 5.046 | 4.024 | 1.225 | 4.062 | 10.392 | 128406 |
|  | CMTK A | 6.835 | 6.888 | 1.225 | 4.528 | 15.811 | 108018 |
|  | CMTK B | 6.723 | 6.262 | 1.225 | 4.528 | 15.716 | 108284 |
|  | CMTK C | 5.414 | 4.475 | 1.225 | 4.183 | 11.023 | 141996 |
|  | Elastix A | 4.973 | 4.117 | 1.000 | 3.873 | 10.025 | 104854 |
|  | Elastix B | 5.432 | 4.113 | 1.225 | 4.528 | 10.817 | 115676 |

**Table S20:** SMP_R : superior medial protocerebrum

| Template | Algorithm | Mean | Std dev | 10th perc | median | 90th perc | N |
| --- | --- | --- | --- | --- | --- | --- | --- |
| JFRC2010 | ANTs A | 4.344 | 4.449 | 0.707 | 2.915 | 10.149 | 2145362 |
|  | ANTs B | 2.678 | 2.441 | 0.707 | 2.121 | 5.701 | 2003074 |
|  | ANTs C | 2.726 | 2.483 | 0.707 | 2.121 | 5.788 | 2169630 |
|  | CMTK A | 4.002 | 5.297 | 0.707 | 2.345 | 8.544 | 1631608 |
|  | CMTK B | 3.727 | 4.456 | 0.707 | 2.236 | 8.337 | 1653970 |
|  | CMTK C | 3.889 | 5.259 | 0.707 | 2.236 | 8.396 | 1886180 |
|  | Elastix A | 2.773 | 2.542 | 0.707 | 2.121 | 5.831 | 1719396 |
|  | Elastix B | 3.650 | 3.492 | 0.707 | 2.550 | 8.093 | 1589316 |
| JFRC2013 | ANTs A | 2.489 | 2.351 | 0.707 | 1.732 | 5.292 | 1971190 |
|  | ANTs B | 2.799 | 2.583 | 0.707 | 2.121 | 5.916 | 1889764 |
|  | ANTs C | 2.804 | 2.629 | 0.707 | 2.121 | 6.042 | 1873850 |
|  | CMTK A | 2.739 | 2.587 | 0.707 | 2.000 | 5.874 | 1856670 |
|  | CMTK B | 2.860 | 2.638 | 0.707 | 2.121 | 6.164 | 1832456 |
|  | CMTK C | 3.399 | 2.853 | 0.707 | 2.646 | 7.141 | 1670644 |
|  | Elastix A | 2.595 | 2.461 | 0.707 | 2.000 | 5.431 | 1868890 |
|  | Elastix B | 2.817 | 2.723 | 0.707 | 2.121 | 6.000 | 1851748 |
| JRC2018 | ANTs A | 2.531 | 2.454 | 0.707 | 1.732 | 5.431 | 1976502 |
|  | ANTs B | 2.569 | 2.464 | 0.707 | 2.000 | 5.431 | 1908536 |
|  | ANTs C | 2.583 | 2.501 | 0.707 | 1.732 | 5.523 | 1983574 |
|  | CMTK A | 2.515 | 2.362 | 0.707 | 2.000 | 5.196 | 1963182 |
|  | CMTK B | 2.547 | 2.379 | 0.707 | 2.000 | 5.196 | 1884060 |
|  | CMTK C | 2.598 | 2.432 | 0.707 | 2.000 | 5.385 | 1856570 |
|  | Elastix A | 2.639 | 2.456 | 0.707 | 2.000 | 5.523 | 1831450 |
|  | Elastix B | 2.779 | 2.623 | 0.707 | 2.121 | 5.874 | 1873824 |
| FCWB | ANTs A | 2.694 | 2.593 | 0.707 | 2.000 | 5.745 | 1984280 |
|  | ANTs B | 3.214 | 2.842 | 0.707 | 2.449 | 6.708 | 1554920 |
|  | ANTs C | 3.343 | 3.080 | 0.707 | 2.550 | 7.071 | 1532000 |
|  | CMTK A | 3.123 | 2.787 | 0.707 | 2.345 | 6.557 | 1858790 |
|  | CMTK B | 3.052 | 2.743 | 0.707 | 2.236 | 6.442 | 1759028 |
|  | CMTK C | 3.273 | 2.850 | 0.707 | 2.449 | 7.036 | 1688338 |
|  | Elastix A | 3.892 | 3.776 | 0.707 | 2.646 | 8.631 | 1591698 |
|  | Elastix B | 3.542 | 3.348 | 0.707 | 2.550 | 7.778 | 1661438 |
| Tefor | ANTs A | 2.519 | 2.438 | 0.707 | 1.732 | 5.385 | 1968522 |
|  | ANTs B | 2.654 | 2.423 | 0.707 | 2.000 | 5.701 | 2000184 |
|  | ANTs C | 2.682 | 2.415 | 0.707 | 2.121 | 5.701 | 2215018 |
|  | CMTK A | 3.435 | 4.027 | 0.707 | 2.236 | 7.382 | 1750938 |
|  | CMTK B | 3.379 | 3.966 | 0.707 | 2.236 | 7.106 | 1758512 |
|  | CMTK C | 3.123 | 3.003 | 0.707 | 2.236 | 6.708 | 1928770 |
|  | Elastix A | 2.797 | 2.604 | 0.707 | 2.121 | 5.874 | 1803676 |
|  | Elastix B | 3.623 | 3.452 | 0.707 | 2.550 | 8.185 | 1746914 |

**Table S21:**
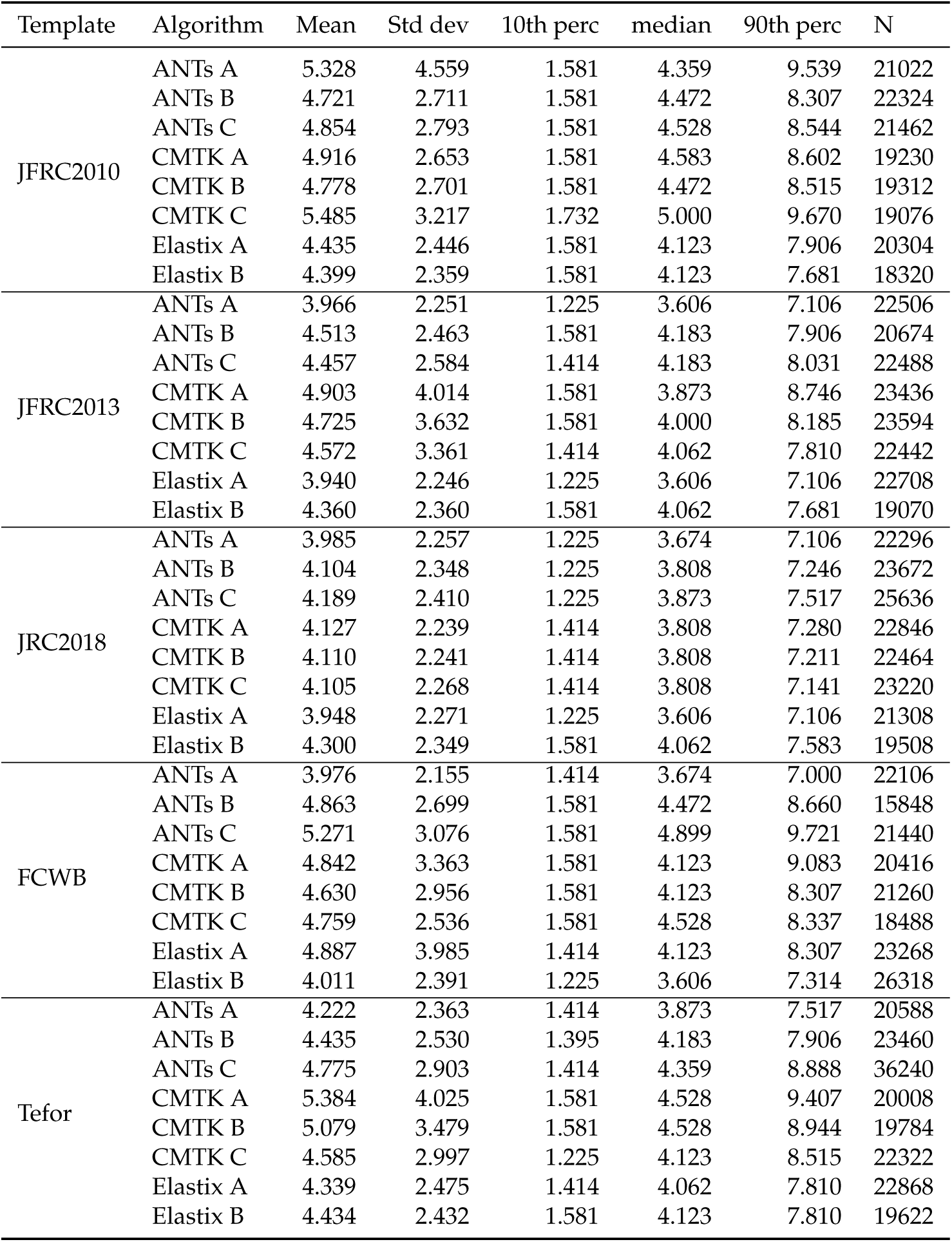
AME_R : accessory medulla

| Template | Algorithm | Mean | Std dev | 10th perc | median | 90th perc | N |
| --- | --- | --- | --- | --- | --- | --- | --- |
| JFRC2010 | ANTs A | 5.328 | 4.559 | 1.581 | 4.359 | 9.539 | 21022 |
|  | ANTs B | 4.721 | 2.711 | 1.581 | 4.472 | 8.307 | 22324 |
|  | ANTs C | 4.854 | 2.793 | 1.581 | 4.528 | 8.544 | 21462 |
|  | CMTK A | 4.916 | 2.653 | 1.581 | 4.583 | 8.602 | 19230 |
|  | CMTK B | 4.778 | 2.701 | 1.581 | 4.472 | 8.515 | 19312 |
|  | CMTK C | 5.485 | 3.217 | 1.732 | 5.000 | 9.670 | 19076 |
|  | Elastix A | 4.435 | 2.446 | 1.581 | 4.123 | 7.906 | 20304 |
|  | Elastix B | 4.399 | 2.359 | 1.581 | 4.123 | 7.681 | 18320 |
| JFRC2013 | ANTs A | 3.966 | 2.251 | 1.225 | 3.606 | 7.106 | 22506 |
|  | ANTs B | 4.513 | 2.463 | 1.581 | 4.183 | 7.906 | 20674 |
|  | ANTs C | 4.457 | 2.584 | 1.414 | 4.183 | 8.031 | 22488 |
|  | CMTK A | 4.903 | 4.014 | 1.581 | 3.873 | 8.746 | 23436 |
|  | CMTK B | 4.725 | 3.632 | 1.581 | 4.000 | 8.185 | 23594 |
|  | CMTK C | 4.572 | 3.361 | 1.414 | 4.062 | 7.810 | 22442 |
|  | Elastix A | 3.940 | 2.246 | 1.225 | 3.606 | 7.106 | 22708 |
|  | Elastix B | 4.360 | 2.360 | 1.581 | 4.062 | 7.681 | 19070 |
| JRC2018 | ANTs A | 3.985 | 2.257 | 1.225 | 3.674 | 7.106 | 22296 |
|  | ANTs B | 4.104 | 2.348 | 1.225 | 3.808 | 7.246 | 23672 |
|  | ANTs C | 4.189 | 2.410 | 1.225 | 3.873 | 7.517 | 25636 |
|  | CMTK A | 4.127 | 2.239 | 1.414 | 3.808 | 7.280 | 22846 |
|  | CMTK B | 4.110 | 2.241 | 1.414 | 3.808 | 7.211 | 22464 |
|  | CMTK C | 4.105 | 2.268 | 1.414 | 3.808 | 7.141 | 23220 |
|  | Elastix A | 3.948 | 2.271 | 1.225 | 3.606 | 7.106 | 21308 |
|  | Elastix B | 4.300 | 2.349 | 1.581 | 4.062 | 7.583 | 19508 |
| FCWB | ANTs A | 3.976 | 2.155 | 1.414 | 3.674 | 7.000 | 22106 |
|  | ANTs B | 4.863 | 2.699 | 1.581 | 4.472 | 8.660 | 15848 |
|  | ANTs C | 5.271 | 3.076 | 1.581 | 4.899 | 9.721 | 21440 |
|  | CMTK A | 4.842 | 3.363 | 1.581 | 4.123 | 9.083 | 20416 |
|  | CMTK B | 4.630 | 2.956 | 1.581 | 4.123 | 8.307 | 21260 |
|  | CMTK C | 4.759 | 2.536 | 1.581 | 4.528 | 8.337 | 18488 |
|  | Elastix A | 4.887 | 3.985 | 1.414 | 4.123 | 8.307 | 23268 |
|  | Elastix B | 4.011 | 2.391 | 1.225 | 3.606 | 7.314 | 26318 |
| Tefor | ANTs A | 4.222 | 2.363 | 1.414 | 3.873 | 7.517 | 20588 |
|  | ANTs B | 4.435 | 2.530 | 1.395 | 4.183 | 7.906 | 23460 |
|  | ANTs C | 4.775 | 2.903 | 1.414 | 4.359 | 8.888 | 36240 |
|  | CMTK A | 5.384 | 4.025 | 1.581 | 4.528 | 9.407 | 20008 |
|  | CMTK B | 5.079 | 3.479 | 1.581 | 4.528 | 8.944 | 19784 |
|  | CMTK C | 4.585 | 2.997 | 1.225 | 4.123 | 8.515 | 22322 |
|  | Elastix A | 4.339 | 2.475 | 1.414 | 4.062 | 7.810 | 22868 |
|  | Elastix B | 4.434 | 2.432 | 1.581 | 4.123 | 7.810 | 19622 |

**Table S22:** AVLP_R : anterior ventrolateral protocerebrum

| Template | Algorithm | Mean | Std dev | 10th perc | median | 90th perc | N |
| --- | --- | --- | --- | --- | --- | --- | --- |
| JFRC2010 | ANTs A | 7.648 | 6.824 | 1.732 | 5.568 | 16.263 | 636130 |
|  | ANTs B | 5.618 | 3.997 | 1.581 | 4.743 | 10.886 | 609838 |
|  | ANTs C | 5.737 | 4.157 | 1.581 | 4.743 | 11.247 | 675888 |
|  | CMTK A | 6.115 | 4.523 | 1.581 | 5.050 | 12.104 | 690236 |
|  | CMTK B | 6.010 | 4.515 | 1.581 | 4.950 | 12.000 | 669340 |
|  | CMTK C | 6.487 | 5.247 | 1.581 | 5.196 | 12.767 | 577554 |
|  | Elastix A | 5.060 | 3.534 | 1.414 | 4.359 | 9.539 | 665630 |
|  | Elastix B | 5.627 | 4.177 | 1.414 | 4.637 | 11.045 | 714568 |
| JFRC2013 | ANTs A | 4.934 | 3.495 | 1.414 | 4.183 | 9.301 | 682792 |
|  | ANTs B | 6.238 | 4.374 | 1.581 | 5.196 | 12.124 | 610754 |
|  | ANTs C | 6.265 | 4.588 | 1.581 | 5.196 | 12.510 | 631998 |
|  | CMTK A | 5.675 | 4.003 | 1.581 | 4.743 | 10.840 | 639244 |
|  | CMTK B | 5.795 | 4.097 | 1.581 | 4.950 | 11.136 | 616978 |
|  | CMTK C | 6.599 | 4.397 | 2.000 | 5.701 | 12.430 | 491772 |
|  | Elastix A | 4.974 | 3.459 | 1.414 | 4.301 | 9.301 | 691716 |
|  | Elastix B | 4.997 | 3.527 | 1.414 | 4.243 | 9.460 | 709604 |
| JRC2018 | ANTs A | 4.907 | 3.465 | 1.414 | 4.183 | 9.301 | 690704 |
|  | ANTs B | 5.110 | 3.605 | 1.414 | 4.359 | 9.644 | 665526 |
|  | ANTs C | 5.286 | 3.727 | 1.414 | 4.528 | 10.100 | 701290 |
|  | CMTK A | 4.881 | 3.416 | 1.414 | 4.183 | 9.165 | 680930 |
|  | CMTK B | 4.901 | 3.459 | 1.414 | 4.183 | 9.220 | 670036 |
|  | CMTK C | 4.990 | 3.538 | 1.414 | 4.243 | 9.434 | 655926 |
|  | Elastix A | 4.956 | 3.476 | 1.414 | 4.243 | 9.301 | 669066 |
|  | Elastix B | 4.975 | 3.490 | 1.414 | 4.243 | 9.434 | 684326 |
| FCWB | ANTs A | 5.468 | 4.045 | 1.414 | 4.528 | 10.677 | 661724 |
|  | ANTs B | 6.114 | 4.267 | 1.581 | 5.196 | 11.790 | 519096 |
|  | ANTs C | 6.485 | 4.682 | 1.581 | 5.431 | 12.748 | 542078 |
|  | CMTK A | 6.167 | 4.299 | 1.581 | 5.196 | 11.958 | 642618 |
|  | CMTK B | 5.998 | 4.256 | 1.581 | 5.050 | 11.662 | 618072 |
|  | CMTK C | 6.080 | 4.249 | 1.581 | 5.148 | 11.853 | 534082 |
|  | Elastix A | 6.115 | 4.193 | 1.581 | 5.196 | 11.683 | 653536 |
|  | Elastix B | 5.540 | 4.195 | 1.414 | 4.528 | 10.724 | 657850 |
| Tefor | ANTs A | 4.941 | 3.490 | 1.414 | 4.183 | 9.301 | 707608 |
|  | ANTs B | 5.279 | 3.596 | 1.581 | 4.528 | 9.849 | 654438 |
|  | ANTs C | 5.499 | 3.863 | 1.581 | 4.690 | 10.464 | 698276 |
|  | CMTK A | 6.070 | 5.111 | 1.414 | 4.743 | 12.207 | 751188 |
|  | CMTK B | 6.105 | 5.215 | 1.414 | 4.743 | 12.430 | 731016 |
|  | CMTK C | 5.735 | 4.566 | 1.581 | 4.637 | 11.023 | 686572 |
|  | Elastix A | 5.028 | 3.540 | 1.414 | 4.359 | 9.460 | 667658 |
|  | Elastix B | 5.790 | 4.438 | 1.414 | 4.743 | 11.424 | 750524 |

**Table S23:**
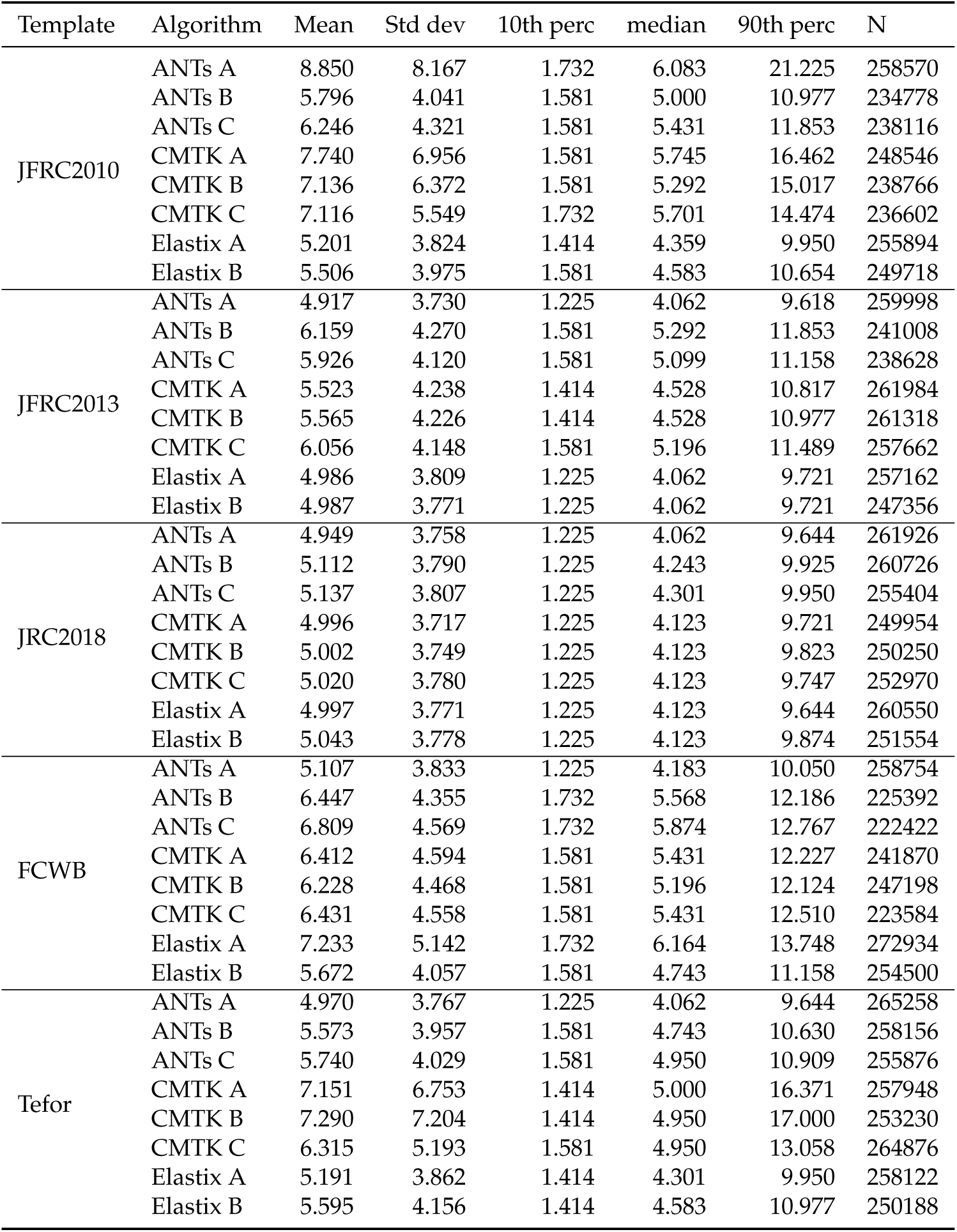
PVLP_R : posterior ventrolateral protocerebrum

**Table S24:**
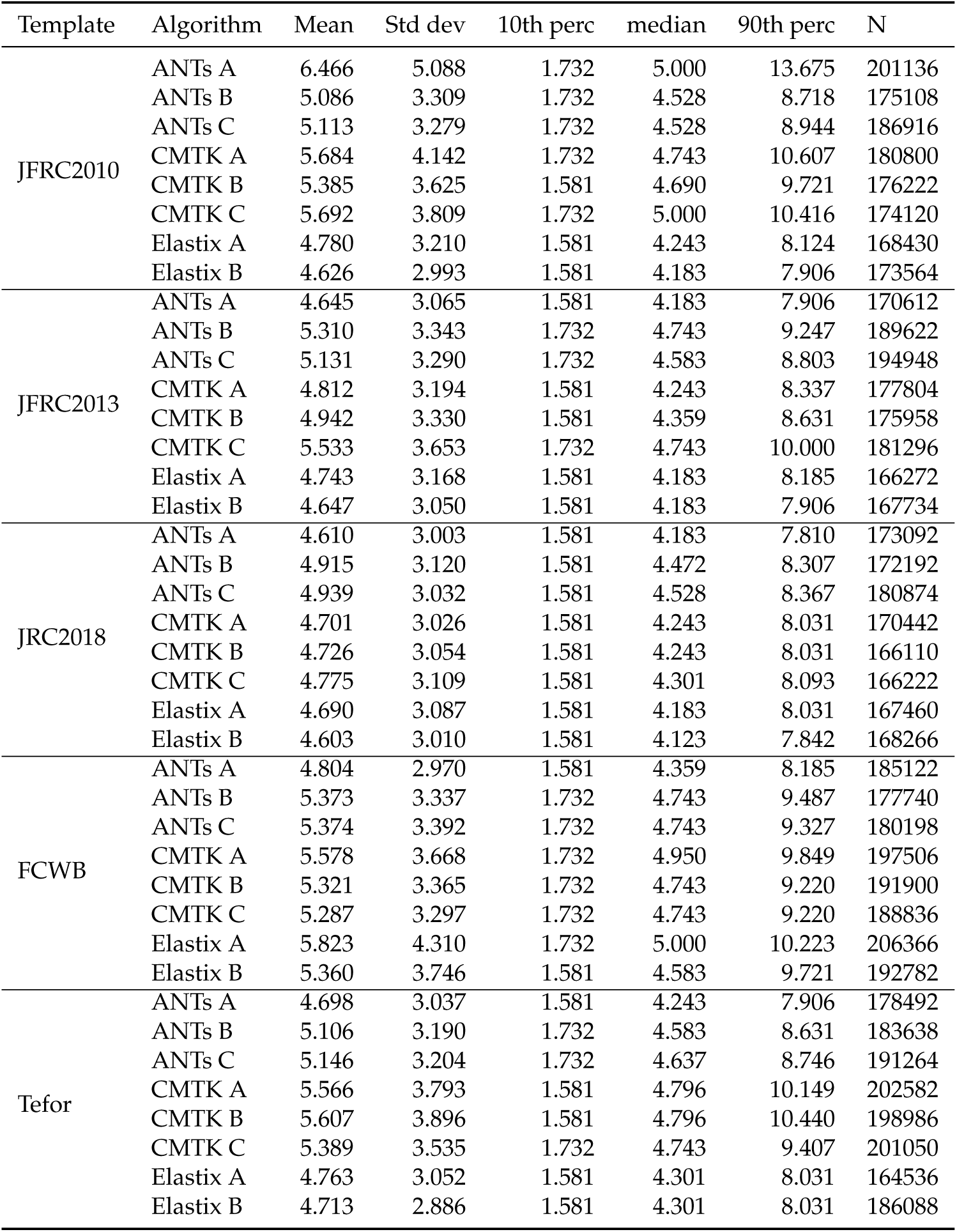
IVLP_R : wedge

**Table S25:** PLP_R : posterior lateral protocerebrum

| Template | Algorithm | Mean | Std dev | 10th perc | median | 90th perc | N |
| --- | --- | --- | --- | --- | --- | --- | --- |
| JFRC2010 | ANTs A | 8.138 | 7.834 | 1.000 | 5.292 | 20.012 | 436308 |
|  | ANTs B | 3.868 | 4.330 | 0.707 | 2.345 | 9.301 | 431512 |
|  | ANTs C | 4.288 | 4.362 | 0.707 | 2.828 | 9.950 | 405618 |
|  | CMTK A | 4.167 | 5.672 | 0.707 | 2.121 | 11.000 | 408410 |
|  | CMTK B | 4.087 | 5.280 | 0.707 | 2.121 | 10.464 | 408162 |
|  | CMTK C | 4.759 | 5.428 | 0.707 | 2.915 | 11.619 | 392684 |
|  | Elastix A | 3.515 | 4.163 | 0.707 | 2.121 | 8.337 | 420146 |
|  | Elastix B | 3.369 | 4.185 | 0.707 | 2.000 | 8.307 | 416076 |
| JFRC2013 | ANTs A | 3.491 | 4.434 | 0.707 | 2.000 | 8.631 | 426466 |
|  | ANTs B | 4.013 | 4.123 | 0.707 | 2.646 | 9.220 | 436614 |
|  | ANTs C | 3.941 | 4.192 | 0.707 | 2.550 | 9.220 | 446714 |
|  | CMTK A | 4.096 | 4.460 | 0.707 | 2.550 | 9.925 | 394782 |
|  | CMTK B | 4.233 | 4.546 | 0.707 | 2.646 | 9.925 | 396906 |
|  | CMTK C | 4.599 | 4.552 | 0.707 | 3.000 | 10.724 | 417880 |
|  | Elastix A | 3.444 | 4.263 | 0.707 | 2.000 | 8.485 | 407894 |
|  | Elastix B | 3.320 | 4.261 | 0.707 | 1.732 | 8.276 | 416212 |
| JRC2018 | ANTs A | 3.424 | 4.378 | 0.707 | 1.732 | 8.573 | 433598 |
|  | ANTs B | 3.581 | 4.304 | 0.707 | 2.121 | 8.544 | 421340 |
|  | ANTs C | 3.550 | 4.278 | 0.707 | 2.121 | 8.544 | 437168 |
|  | CMTK A | 3.426 | 4.491 | 0.707 | 1.732 | 8.485 | 412332 |
|  | CMTK B | 3.393 | 4.419 | 0.707 | 1.732 | 8.276 | 408696 |
|  | CMTK C | 3.416 | 4.308 | 0.707 | 2.000 | 8.246 | 406596 |
|  | Elastix A | 3.353 | 4.264 | 0.707 | 2.000 | 8.093 | 416490 |
|  | Elastix B | 3.350 | 4.296 | 0.707 | 1.732 | 8.185 | 411904 |
| FCWB | ANTs A | 3.640 | 4.550 | 0.707 | 2.121 | 9.000 | 471818 |
|  | ANTs B | 4.230 | 4.373 | 0.707 | 2.646 | 9.849 | 425034 |
|  | ANTs C | 4.165 | 4.300 | 0.707 | 2.646 | 9.460 | 428146 |
|  | CMTK A | 6.287 | 6.209 | 1.000 | 4.062 | 15.067 | 357542 |
|  | CMTK B | 5.840 | 5.666 | 1.000 | 3.873 | 13.342 | 326860 |
|  | CMTK C | 4.421 | 5.056 | 0.707 | 2.646 | 10.700 | 408458 |
|  | Elastix A | 6.252 | 5.924 | 0.707 | 4.359 | 14.491 | 382002 |
|  | Elastix B | 4.167 | 4.870 | 0.707 | 2.236 | 10.886 | 423918 |
| Tefor | ANTs A | 3.436 | 4.397 | 0.707 | 1.732 | 8.631 | 443520 |
|  | ANTs B | 3.818 | 4.130 | 0.707 | 2.345 | 9.028 | 435004 |
|  | ANTs C | 3.974 | 4.276 | 0.707 | 2.449 | 9.460 | 434722 |
|  | CMTK A | 4.141 | 5.693 | 0.707 | 2.121 | 10.607 | 405412 |
|  | CMTK B | 4.286 | 5.878 | 0.707 | 2.121 | 11.068 | 406718 |
|  | CMTK C | 4.370 | 5.011 | 0.707 | 2.550 | 10.607 | 418774 |
|  | Elastix A | 3.531 | 4.427 | 0.707 | 2.000 | 8.660 | 421524 |
|  | Elastix B | 3.351 | 4.235 | 0.707 | 1.732 | 8.485 | 416838 |

**Table S26:** AOTU_R : anterior optic tubercle

| Template | Algorithm | Mean | Std dev | 10th perc | median | 90th perc | N |
| --- | --- | --- | --- | --- | --- | --- | --- |
| JFRC2010 | ANTs A | 13.329 | 10.282 | 2.121 | 10.863 | 28.328 | 17992 |
|  | ANTs B | 7.807 | 8.316 | 1.000 | 4.062 | 21.331 | 14808 |
|  | ANTs C | 8.087 | 7.943 | 1.225 | 5.000 | 20.917 | 14984 |
|  | CMTK A | 8.705 | 8.213 | 1.225 | 5.339 | 21.953 | 16612 |
|  | CMTK B | 9.651 | 9.371 | 1.225 | 5.385 | 25.159 | 16454 |
|  | CMTK C | 9.807 | 9.479 | 1.581 | 5.701 | 25.060 | 13412 |
|  | Elastix A | 8.053 | 7.717 | 1.225 | 5.148 | 19.873 | 17078 |
|  | Elastix B | 7.727 | 7.692 | 1.000 | 4.528 | 20.075 | 17320 |
| JFRC2013 | ANTs A | 7.026 | 7.092 | 1.000 | 4.301 | 17.479 | 17460 |
|  | ANTs B | 6.472 | 7.142 | 1.000 | 3.606 | 16.703 | 14228 |
|  | ANTs C | 6.378 | 7.364 | 1.000 | 3.240 | 17.944 | 13420 |
|  | CMTK A | 10.651 | 9.174 | 1.414 | 7.583 | 24.208 | 19620 |
|  | CMTK B | 10.417 | 8.960 | 1.414 | 7.681 | 23.633 | 19254 |
|  | CMTK C | 10.034 | 8.353 | 2.000 | 7.280 | 22.372 | 18320 |
|  | Elastix A | 7.791 | 7.466 | 1.000 | 5.000 | 19.085 | 16856 |
|  | Elastix B | 7.241 | 7.267 | 1.000 | 4.472 | 18.083 | 18682 |
| JRC2018 | ANTs A | 7.155 | 7.195 | 1.000 | 4.301 | 17.776 | 17150 |
|  | ANTs B | 7.451 | 7.594 | 1.000 | 4.301 | 19.209 | 16210 |
|  | ANTs C | 7.851 | 7.668 | 1.225 | 4.690 | 20.261 | 15504 |
|  | CMTK A | 7.115 | 6.907 | 1.000 | 4.472 | 17.331 | 17054 |
|  | CMTK B | 7.108 | 7.046 | 1.000 | 4.359 | 17.819 | 17370 |
|  | CMTK C | 7.315 | 7.216 | 1.000 | 4.528 | 18.371 | 15946 |
|  | Elastix A | 7.141 | 7.254 | 1.000 | 4.183 | 18.044 | 16760 |
|  | Elastix B | 6.963 | 6.918 | 1.000 | 4.359 | 17.517 | 18784 |
| FCWB | ANTs A | 7.659 | 7.969 | 1.000 | 4.472 | 19.456 | 16620 |
|  | ANTs B | 7.112 | 8.124 | 1.000 | 4.359 | 17.972 | 10226 |
|  | ANTs C | 8.176 | 8.485 | 1.225 | 5.000 | 22.683 | 13788 |
|  | CMTK A | 9.582 | 8.345 | 1.581 | 6.708 | 22.159 | 15260 |
|  | CMTK B | 9.158 | 8.109 | 1.581 | 6.245 | 21.691 | 15174 |
|  | CMTK C | 7.059 | 8.317 | 1.000 | 3.464 | 20.846 | 17600 |
|  | Elastix A | 8.114 | 7.969 | 1.000 | 4.796 | 21.483 | 16684 |
|  | Elastix B | 6.995 | 6.898 | 1.000 | 4.359 | 17.751 | 18760 |
| Tefor | ANTs A | 7.077 | 6.970 | 1.000 | 4.359 | 17.578 | 17286 |
|  | ANTs B | 7.911 | 7.031 | 1.581 | 5.701 | 18.152 | 13698 |
|  | ANTs C | 6.938 | 6.834 | 1.000 | 4.472 | 17.357 | 17156 |
|  | CMTK A | 10.065 | 8.432 | 1.414 | 7.416 | 22.858 | 19664 |
|  | CMTK B | 9.895 | 8.227 | 1.414 | 7.280 | 22.749 | 19914 |
|  | CMTK C | 7.788 | 7.484 | 1.225 | 4.583 | 19.515 | 23164 |
|  | Elastix A | 7.023 | 7.338 | 1.000 | 3.808 | 18.155 | 18980 |
|  | Elastix B | 7.508 | 7.435 | 1.225 | 4.528 | 18.345 | 19840 |

**Table S27:** GOR_R : gorget

| Template | Algorithm | Mean | Std dev | 10th perc | median | 90th perc | N |
| --- | --- | --- | --- | --- | --- | --- | --- |
| JFRC2010 | ANTs A | 7.788 | 8.293 | 1.225 | 5.000 | 19.962 | 113000 |
|  | ANTs B | 3.837 | 3.175 | 0.707 | 2.915 | 8.185 | 105928 |
|  | ANTs C | 4.132 | 3.291 | 0.707 | 3.240 | 8.775 | 103418 |
|  | CMTK A | 5.092 | 6.903 | 0.707 | 2.915 | 11.269 | 84104 |
|  | CMTK B | 5.052 | 6.506 | 0.707 | 2.915 | 11.068 | 85226 |
|  | CMTK C | 5.238 | 6.329 | 1.000 | 3.162 | 11.769 | 98656 |
|  | Elastix A | 3.411 | 2.662 | 0.707 | 2.915 | 6.708 | 94852 |
|  | Elastix B | 3.594 | 2.928 | 0.707 | 2.915 | 7.246 | 88476 |
| JFRC2013 | ANTs A | 3.239 | 2.488 | 0.707 | 2.646 | 6.671 | 101422 |
|  | ANTs B | 4.387 | 3.451 | 1.000 | 3.536 | 9.220 | 99430 |
|  | ANTs C | 4.344 | 3.432 | 1.000 | 3.464 | 9.220 | 97998 |
|  | CMTK A | 3.365 | 2.706 | 0.707 | 2.646 | 7.071 | 84030 |
|  | CMTK B | 3.425 | 2.794 | 0.707 | 2.646 | 7.106 | 86578 |
|  | CMTK C | 4.234 | 3.308 | 1.000 | 3.536 | 8.000 | 99184 |
|  | Elastix A | 3.355 | 2.576 | 0.707 | 2.646 | 6.856 | 92320 |
|  | Elastix B | 3.236 | 2.484 | 0.707 | 2.646 | 6.671 | 90826 |
| JRC2018 | ANTs A | 3.212 | 2.472 | 0.707 | 2.646 | 6.557 | 99712 |
|  | ANTs B | 3.614 | 2.878 | 0.707 | 2.915 | 7.616 | 103552 |
|  | ANTs C | 3.364 | 2.580 | 0.707 | 2.646 | 6.856 | 97756 |
|  | CMTK A | 3.384 | 2.692 | 0.707 | 2.646 | 7.141 | 81058 |
|  | CMTK B | 3.361 | 2.638 | 0.707 | 2.646 | 7.071 | 82722 |
|  | CMTK C | 3.223 | 2.431 | 0.707 | 2.646 | 6.481 | 96996 |
|  | Elastix A | 3.298 | 2.399 | 0.707 | 2.828 | 6.481 | 107920 |
|  | Elastix B | 3.197 | 2.385 | 0.707 | 2.646 | 6.442 | 99008 |
| FCWB | ANTs A | 3.224 | 2.556 | 0.707 | 2.550 | 6.671 | 97924 |
|  | ANTs B | 3.967 | 2.743 | 1.000 | 3.317 | 7.649 | 94372 |
|  | ANTs C | 3.714 | 2.654 | 1.000 | 3.082 | 7.246 | 94252 |
|  | CMTK A | 6.950 | 7.080 | 1.000 | 4.301 | 18.348 | 67048 |
|  | CMTK B | 6.223 | 6.145 | 1.000 | 4.062 | 15.232 | 80174 |
|  | CMTK C | 3.882 | 3.410 | 0.707 | 2.915 | 8.031 | 102780 |
|  | Elastix A | 9.584 | 9.620 | 1.000 | 5.523 | 26.182 | 86106 |
|  | Elastix B | 4.043 | 3.799 | 0.707 | 2.915 | 8.631 | 99540 |
| Tefor | ANTs A | 3.170 | 2.431 | 0.707 | 2.550 | 6.481 | 98684 |
|  | ANTs B | 3.797 | 3.090 | 0.707 | 2.915 | 8.031 | 107810 |
|  | ANTs C | 4.030 | 3.142 | 0.707 | 3.240 | 8.307 | 101590 |
|  | CMTK A | 4.006 | 4.605 | 0.707 | 2.646 | 8.185 | 82046 |
|  | CMTK B | 3.913 | 4.397 | 0.707 | 2.646 | 8.031 | 82136 |
|  | CMTK C | 3.854 | 3.416 | 1.000 | 3.000 | 7.517 | 106304 |
|  | Elastix A | 3.301 | 2.545 | 0.707 | 2.646 | 6.671 | 94924 |
|  | Elastix B | 3.731 | 2.989 | 1.000 | 2.915 | 7.616 | 90522 |

**Table S28:** MB_CA_R : calyx of adult mushroom body

| Template | Algorithm | Mean | Std dev | 10th perc | median | 90th perc | N |
| --- | --- | --- | --- | --- | --- | --- | --- |
| JFRC2010 | ANTs A | 4.842 | 3.579 | 1.225 | 4.000 | 9.644 | 449372 |
|  | ANTs B | 5.478 | 4.268 | 1.414 | 4.359 | 11.091 | 284214 |
|  | ANTs C | 4.769 | 3.397 | 1.225 | 4.062 | 9.301 | 374730 |
|  | CMTK A | 5.117 | 4.587 | 1.000 | 3.808 | 10.724 | 364534 |
|  | CMTK B | 4.714 | 3.852 | 1.000 | 3.674 | 9.301 | 360984 |
|  | CMTK C | 5.814 | 5.695 | 1.225 | 4.183 | 11.597 | 292448 |
|  | Elastix A | 4.104 | 2.753 | 1.225 | 3.536 | 7.906 | 353136 |
|  | Elastix B | 3.748 | 2.496 | 1.000 | 3.240 | 7.141 | 385158 |
| JFRC2013 | ANTs A | 5.162 | 3.945 | 1.225 | 4.123 | 10.607 | 309960 |
|  | ANTs B | 5.486 | 4.208 | 1.414 | 4.359 | 11.068 | 312332 |
|  | ANTs C | 5.548 | 4.330 | 1.414 | 4.359 | 11.424 | 305446 |
|  | CMTK A | 3.896 | 2.553 | 1.000 | 3.317 | 7.416 | 238400 |
|  | CMTK B | 4.203 | 2.846 | 1.225 | 3.606 | 8.093 | 228912 |
|  | CMTK C | 5.285 | 3.729 | 1.414 | 4.472 | 10.271 | 184862 |
|  | Elastix A | 3.729 | 2.415 | 1.000 | 3.240 | 7.071 | 319332 |
|  | Elastix B | 3.576 | 2.326 | 1.000 | 3.082 | 6.819 | 328444 |
| JRC2018 | ANTs A | 3.546 | 2.446 | 1.000 | 2.915 | 7.000 | 421666 |
|  | ANTs B | 4.481 | 3.102 | 1.225 | 3.808 | 8.660 | 400478 |
|  | ANTs C | 4.315 | 2.930 | 1.225 | 3.674 | 8.276 | 417118 |
|  | CMTK A | 3.648 | 2.524 | 1.000 | 3.000 | 7.106 | 423462 |
|  | CMTK B | 3.774 | 2.534 | 1.000 | 3.162 | 7.280 | 407944 |
|  | CMTK C | 3.967 | 2.564 | 1.000 | 3.536 | 7.517 | 390700 |
|  | Elastix A | 4.153 | 2.860 | 1.000 | 3.536 | 8.093 | 400538 |
|  | Elastix B | 3.720 | 2.532 | 1.000 | 3.162 | 7.211 | 406424 |
| FCWB | ANTs A | 4.132 | 3.032 | 1.000 | 3.464 | 8.093 | 315408 |
|  | ANTs B | 5.556 | 4.440 | 1.414 | 4.359 | 11.424 | 262056 |
|  | ANTs C | 5.035 | 3.930 | 1.225 | 4.062 | 10.223 | 304860 |
|  | CMTK A | 5.056 | 4.063 | 1.000 | 3.873 | 10.512 | 375234 |
|  | CMTK B | 4.835 | 3.779 | 1.000 | 3.808 | 9.670 | 370578 |
|  | CMTK C | 5.338 | 4.104 | 1.225 | 4.301 | 10.817 | 240104 |
|  | Elastix A | 6.264 | 5.314 | 1.225 | 4.583 | 14.195 | 267858 |
|  | Elastix B | 4.907 | 4.293 | 1.000 | 3.606 | 10.223 | 308952 |
| Tefor | ANTs A | 3.444 | 2.382 | 1.000 | 2.915 | 6.708 | 414600 |
|  | ANTs B | 5.301 | 4.154 | 1.225 | 4.123 | 11.000 | 262940 |
|  | ANTs C | 5.297 | 4.199 | 1.225 | 4.123 | 11.269 | 316732 |
|  | CMTK A | 4.965 | 4.998 | 1.000 | 3.464 | 10.700 | 338350 |
|  | CMTK B | 5.116 | 5.223 | 1.000 | 3.536 | 10.817 | 320050 |
|  | CMTK C | 5.524 | 4.721 | 1.225 | 4.183 | 11.336 | 242994 |
|  | Elastix A | 3.923 | 2.675 | 1.000 | 3.317 | 7.517 | 344336 |
|  | Elastix B | 3.640 | 2.457 | 1.000 | 3.082 | 7.000 | 381100 |

**Table S29:** SPS_R : superior posterior slope

| Template | Algorithm | Mean | Std dev | 10th perc | median | 90th perc | N |
| --- | --- | --- | --- | --- | --- | --- | --- |
| JFRC2010 | ANTs A | 7.065 | 5.820 | 1.581 | 5.431 | 15.017 | 416104 |
|  | ANTs B | 5.342 | 4.028 | 1.414 | 4.301 | 10.817 | 403054 |
|  | ANTs C | 5.702 | 4.104 | 1.414 | 4.743 | 11.402 | 430918 |
|  | CMTK A | 7.125 | 6.977 | 1.414 | 4.743 | 16.279 | 309582 |
|  | CMTK B | 6.271 | 5.801 | 1.414 | 4.528 | 13.454 | 318590 |
|  | CMTK C | 6.349 | 5.999 | 1.414 | 4.472 | 13.910 | 423128 |
|  | Elastix A | 4.247 | 3.271 | 1.000 | 3.536 | 8.396 | 367332 |
|  | Elastix B | 4.470 | 3.646 | 1.000 | 3.536 | 9.000 | 331736 |
| JFRC2013 | ANTs A | 4.777 | 3.821 | 1.000 | 3.808 | 9.644 | 365592 |
|  | ANTs B | 5.869 | 4.310 | 1.581 | 4.796 | 11.853 | 354298 |
|  | ANTs C | 5.653 | 4.082 | 1.581 | 4.690 | 11.136 | 379870 |
|  | CMTK A | 4.890 | 3.882 | 1.225 | 3.808 | 10.000 | 309642 |
|  | CMTK B | 4.890 | 3.848 | 1.225 | 3.808 | 9.925 | 319706 |
|  | CMTK C | 5.413 | 4.224 | 1.414 | 4.301 | 11.045 | 394108 |
|  | Elastix A | 4.472 | 3.561 | 1.000 | 3.606 | 8.972 | 359646 |
|  | Elastix B | 4.512 | 3.673 | 1.000 | 3.536 | 9.220 | 343302 |
| JRC2018 | ANTs A | 4.609 | 3.635 | 1.000 | 3.606 | 9.460 | 365344 |
|  | ANTs B | 5.140 | 3.904 | 1.225 | 4.123 | 10.392 | 368634 |
|  | ANTs C | 5.137 | 3.830 | 1.414 | 4.183 | 10.124 | 366652 |
|  | CMTK A | 4.998 | 3.878 | 1.225 | 4.062 | 10.124 | 313232 |
|  | CMTK B | 4.955 | 3.877 | 1.225 | 4.000 | 10.000 | 311026 |
|  | CMTK C | 4.998 | 3.844 | 1.225 | 4.062 | 9.925 | 329436 |
|  | Elastix A | 4.650 | 3.672 | 1.000 | 3.674 | 9.381 | 353656 |
|  | Elastix B | 4.652 | 3.781 | 1.000 | 3.606 | 9.460 | 333390 |
| FCWB | ANTs A | 5.113 | 3.889 | 1.225 | 4.123 | 10.271 | 392366 |
|  | ANTs B | 5.396 | 4.056 | 1.414 | 4.359 | 10.977 | 426566 |
|  | ANTs C | 5.244 | 3.855 | 1.414 | 4.301 | 10.512 | 415894 |
|  | CMTK A | 6.272 | 5.706 | 1.414 | 4.690 | 13.000 | 314618 |
|  | CMTK B | 5.998 | 5.601 | 1.414 | 4.359 | 12.530 | 331114 |
|  | CMTK C | 5.694 | 4.890 | 1.414 | 4.301 | 12.104 | 368580 |
|  | Elastix A | 6.563 | 5.646 | 1.414 | 4.743 | 14.933 | 406550 |
|  | Elastix B | 5.316 | 4.694 | 1.000 | 3.873 | 11.247 | 386478 |
| Tefor | ANTs A | 4.693 | 3.644 | 1.000 | 3.808 | 9.539 | 388030 |
|  | ANTs B | 5.100 | 3.892 | 1.225 | 4.123 | 10.247 | 446012 |
|  | ANTs C | 5.550 | 4.065 | 1.414 | 4.528 | 10.977 | 415622 |
|  | CMTK A | 5.513 | 4.931 | 1.225 | 4.123 | 11.597 | 323322 |
|  | CMTK B | 5.493 | 4.770 | 1.225 | 4.123 | 11.424 | 327944 |
|  | CMTK C | 5.336 | 4.139 | 1.414 | 4.243 | 10.724 | 438296 |
|  | Elastix A | 4.558 | 3.545 | 1.000 | 3.606 | 9.000 | 352978 |
|  | Elastix B | 4.667 | 3.701 | 1.000 | 3.674 | 9.301 | 317442 |

**Table S30:** IPS_R : inferior posterior slope

| Template | Algorithm | Mean | Std dev | 10th perc | median | 90th perc | N |
| --- | --- | --- | --- | --- | --- | --- | --- |
| JFRC2010 | ANTs A | 8.126 | 5.735 | 2.121 | 6.671 | 16.401 | 342866 |
|  | ANTs B | 6.740 | 4.685 | 1.732 | 5.701 | 13.342 | 245712 |
|  | ANTs C | 7.951 | 5.083 | 2.449 | 6.856 | 15.182 | 201122 |
|  | CMTK A | 7.647 | 5.359 | 2.000 | 6.205 | 15.668 | 229274 |
|  | CMTK B | 7.248 | 5.118 | 1.732 | 5.831 | 14.983 | 231834 |
|  | CMTK C | 7.370 | 4.969 | 2.000 | 6.205 | 14.714 | 222776 |
|  | Elastix A | 4.699 | 3.819 | 1.000 | 3.606 | 9.925 | 306590 |
|  | Elastix B | 4.888 | 4.050 | 1.225 | 3.674 | 10.416 | 308932 |
| JFRC2013 | ANTs A | 5.746 | 4.481 | 1.414 | 4.528 | 12.166 | 301130 |
|  | ANTs B | 7.402 | 4.879 | 2.236 | 6.205 | 14.577 | 219492 |
|  | ANTs C | 7.204 | 4.724 | 2.121 | 6.124 | 13.946 | 230954 |
|  | CMTK A | 5.630 | 4.317 | 1.414 | 4.528 | 11.576 | 282136 |
|  | CMTK B | 5.786 | 4.335 | 1.581 | 4.690 | 11.705 | 277218 |
|  | CMTK C | 6.513 | 4.554 | 1.732 | 5.339 | 13.134 | 293938 |
|  | Elastix A | 4.714 | 3.943 | 1.000 | 3.606 | 10.000 | 325016 |
|  | Elastix B | 4.913 | 4.014 | 1.000 | 3.808 | 10.512 | 302166 |
| JRC2018 | ANTs A | 4.866 | 3.894 | 1.225 | 3.808 | 10.025 | 313820 |
|  | ANTs B | 6.137 | 4.565 | 1.581 | 5.000 | 12.550 | 273266 |
|  | ANTs C | 6.024 | 4.265 | 1.581 | 5.000 | 11.832 | 268456 |
|  | CMTK A | 5.262 | 4.008 | 1.414 | 4.243 | 10.512 | 290294 |
|  | CMTK B | 5.347 | 4.095 | 1.414 | 4.301 | 10.724 | 271122 |
|  | CMTK C | 5.732 | 4.272 | 1.581 | 4.583 | 11.489 | 249432 |
|  | Elastix A | 4.668 | 3.871 | 1.000 | 3.606 | 9.721 | 322516 |
|  | Elastix B | 4.906 | 3.956 | 1.225 | 3.808 | 10.223 | 304868 |
| FCWB | ANTs A | 5.664 | 4.176 | 1.414 | 4.583 | 11.424 | 323078 |
|  | ANTs B | 6.484 | 4.423 | 1.732 | 5.431 | 12.728 | 261148 |
|  | ANTs C | 6.926 | 4.620 | 2.121 | 5.916 | 13.229 | 225102 |
|  | CMTK A | 7.538 | 5.218 | 2.000 | 6.245 | 15.248 | 304856 |
|  | CMTK B | 6.912 | 4.943 | 1.732 | 5.701 | 14.053 | 299012 |
|  | CMTK C | 6.947 | 4.662 | 2.000 | 5.874 | 13.675 | 254194 |
|  | Elastix A | 7.752 | 5.436 | 2.121 | 6.519 | 15.297 | 333362 |
|  | Elastix B | 6.437 | 5.103 | 1.581 | 5.000 | 13.583 | 373960 |
| Tefor | ANTs A | 5.070 | 3.997 | 1.225 | 4.062 | 10.536 | 320492 |
|  | ANTs B | 6.728 | 4.517 | 2.000 | 5.701 | 13.172 | 265242 |
|  | ANTs C | 7.193 | 4.684 | 2.121 | 6.124 | 13.730 | 240284 |
|  | CMTK A | 7.271 | 6.902 | 1.414 | 4.796 | 17.000 | 316638 |
|  | CMTK B | 7.316 | 6.931 | 1.581 | 4.950 | 17.015 | 312412 |
|  | CMTK C | 6.579 | 5.477 | 1.581 | 5.000 | 14.036 | 346162 |
|  | Elastix A | 4.592 | 3.740 | 1.000 | 3.606 | 9.513 | 324824 |
|  | Elastix B | 4.867 | 3.865 | 1.225 | 3.808 | 10.025 | 317084 |

**Table S31:**
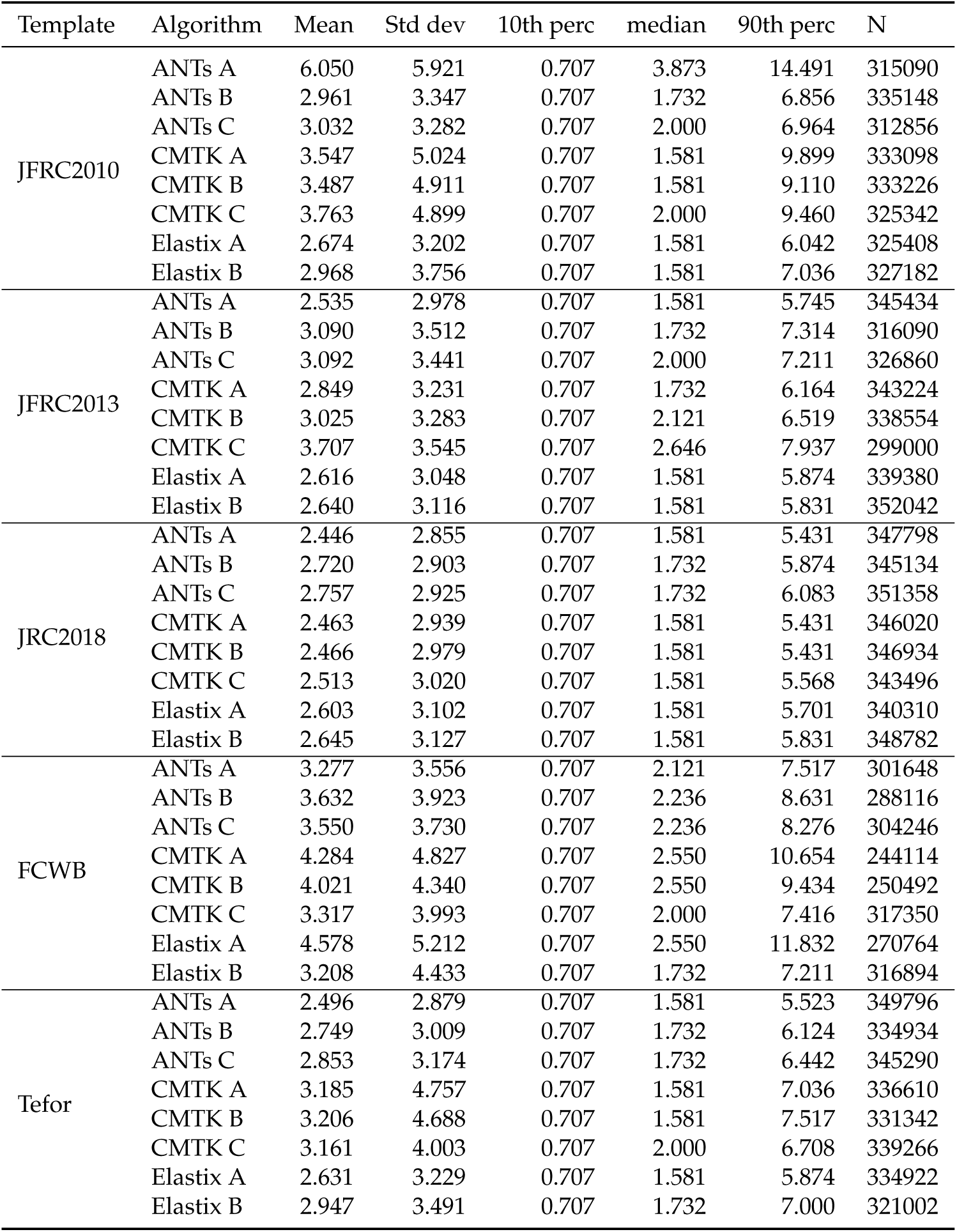
SCL_R : superior clamp

**Table S32:** LO_R : lobula

| Template | Algorithm | Mean | Std dev | 10th perc | median | 90th perc | N |
| --- | --- | --- | --- | --- | --- | --- | --- |
| JFRC2010 | ANTs A | 8.079 | 8.073 | 1.581 | 5.196 | 19.300 | 765240 |
|  | ANTs B | 5.244 | 4.284 | 1.581 | 4.183 | 9.950 | 635108 |
|  | ANTs C | 5.892 | 4.745 | 1.581 | 4.583 | 11.937 | 612046 |
|  | CMTK A | 8.762 | 7.817 | 1.732 | 5.874 | 20.857 | 674518 |
|  | CMTK B | 7.278 | 6.555 | 1.581 | 5.148 | 16.202 | 633590 |
|  | CMTK C | 7.782 | 6.612 | 1.732 | 5.745 | 16.523 | 709042 |
|  | Elastix A | 4.646 | 3.713 | 1.414 | 3.808 | 8.515 | 600342 |
|  | Elastix B | 4.479 | 3.533 | 1.414 | 3.674 | 8.185 | 611184 |
| JFRC2013 | ANTs A | 4.590 | 4.037 | 1.225 | 3.606 | 8.515 | 630492 |
|  | ANTs B | 5.339 | 4.529 | 1.581 | 4.123 | 10.512 | 625254 |
|  | ANTs C | 5.221 | 4.461 | 1.414 | 4.123 | 10.025 | 601610 |
|  | CMTK A | 6.059 | 5.748 | 1.581 | 4.243 | 13.304 | 689216 |
|  | CMTK B | 6.337 | 6.124 | 1.581 | 4.359 | 14.457 | 681688 |
|  | CMTK C | 5.830 | 4.451 | 1.581 | 4.583 | 11.790 | 660042 |
|  | Elastix A | 4.439 | 3.599 | 1.225 | 3.606 | 7.937 | 639570 |
|  | Elastix B | 4.400 | 3.598 | 1.225 | 3.606 | 7.937 | 654960 |
| JRC2018 | ANTs A | 4.368 | 3.696 | 1.225 | 3.606 | 7.906 | 633808 |
|  | ANTs B | 4.542 | 3.821 | 1.414 | 3.606 | 8.307 | 649512 |
|  | ANTs C | 4.659 | 3.911 | 1.414 | 3.808 | 8.544 | 633012 |
|  | CMTK A | 4.361 | 3.625 | 1.225 | 3.606 | 7.906 | 650942 |
|  | CMTK B | 4.359 | 3.594 | 1.225 | 3.606 | 7.906 | 628218 |
|  | CMTK C | 4.443 | 3.657 | 1.225 | 3.606 | 8.062 | 623624 |
|  | Elastix A | 4.386 | 3.570 | 1.225 | 3.606 | 7.842 | 649538 |
|  | Elastix B | 4.541 | 3.737 | 1.414 | 3.674 | 8.185 | 644400 |
| FCWB | ANTs A | 4.659 | 4.054 | 1.225 | 3.606 | 8.631 | 629916 |
|  | ANTs B | 5.370 | 4.123 | 1.581 | 4.359 | 10.025 | 738100 |
|  | ANTs C | 6.037 | 4.515 | 1.732 | 4.950 | 11.597 | 704222 |
|  | CMTK A | 8.843 | 7.562 | 1.732 | 6.164 | 20.408 | 621926 |
|  | CMTK B | 7.667 | 6.647 | 1.732 | 5.339 | 17.692 | 579792 |
|  | CMTK C | 6.573 | 5.154 | 1.732 | 5.099 | 13.620 | 739520 |
|  | Elastix A | 8.836 | 7.008 | 2.121 | 6.708 | 18.520 | 766336 |
|  | Elastix B | 7.715 | 7.530 | 1.581 | 5.148 | 17.776 | 692590 |
| Tefor | ANTs A | 4.395 | 3.671 | 1.225 | 3.606 | 8.031 | 639674 |
|  | ANTs B | 4.926 | 4.055 | 1.414 | 3.873 | 9.220 | 612498 |
|  | ANTs C | 5.096 | 4.334 | 1.414 | 4.062 | 9.644 | 582010 |
|  | CMTK A | 7.184 | 6.627 | 1.581 | 4.796 | 16.688 | 722676 |
|  | CMTK B | 7.202 | 6.647 | 1.581 | 4.950 | 16.583 | 714756 |
|  | CMTK C | 6.569 | 5.881 | 1.581 | 4.690 | 14.577 | 613750 |
|  | Elastix A | 4.570 | 3.677 | 1.414 | 3.808 | 8.307 | 597264 |
|  | Elastix B | 4.457 | 3.577 | 1.225 | 3.606 | 8.093 | 613856 |

**Table S33:** EPA_R : epaulette

| Template | Algorithm | Mean | Std dev | 10th perc | median | 90th perc | N |
| --- | --- | --- | --- | --- | --- | --- | --- |
| JFRC2010 | ANTs A | 10.110 | 9.192 | 1.581 | 7.000 | 24.870 | 20032 |
|  | ANTs B | 5.442 | 5.156 | 1.000 | 3.606 | 13.229 | 15862 |
|  | ANTs C | 7.146 | 6.935 | 1.000 | 4.123 | 18.507 | 13508 |
|  | CMTK A | 6.936 | 9.153 | 1.000 | 3.240 | 18.180 | 12426 |
|  | CMTK B | 6.429 | 8.057 | 1.000 | 3.240 | 15.716 | 12974 |
|  | CMTK C | 6.017 | 6.591 | 1.000 | 3.536 | 15.379 | 10854 |
|  | Elastix A | 3.909 | 3.004 | 1.000 | 3.162 | 7.810 | 16124 |
|  | Elastix B | 4.172 | 2.910 | 1.000 | 3.536 | 8.307 | 22564 |
| JFRC2013 | ANTs A | 3.902 | 3.014 | 1.000 | 3.000 | 8.276 | 20466 |
|  | ANTs B | 7.986 | 6.650 | 1.732 | 5.568 | 19.157 | 10842 |
|  | ANTs C | 8.399 | 6.698 | 1.581 | 5.701 | 18.722 | 12826 |
|  | CMTK A | 3.623 | 2.706 | 1.000 | 2.915 | 7.649 | 16792 |
|  | CMTK B | 3.685 | 2.840 | 1.000 | 2.915 | 7.681 | 17748 |
|  | CMTK C | 5.969 | 5.275 | 1.225 | 4.359 | 12.590 | 16420 |
|  | Elastix A | 3.793 | 2.980 | 1.000 | 2.915 | 8.031 | 21572 |
|  | Elastix B | 3.781 | 3.007 | 1.000 | 2.915 | 8.093 | 21214 |
| JRC2018 | ANTs A | 3.883 | 2.876 | 1.000 | 3.082 | 8.031 | 20550 |
|  | ANTs B | 4.840 | 4.506 | 1.000 | 3.240 | 11.023 | 19126 |
|  | ANTs C | 4.767 | 4.333 | 1.000 | 3.536 | 10.000 | 16668 |
|  | CMTK A | 3.478 | 2.658 | 0.707 | 2.915 | 7.036 | 17250 |
|  | CMTK B | 3.395 | 2.573 | 0.707 | 2.646 | 7.036 | 17552 |
|  | CMTK C | 3.707 | 2.808 | 1.000 | 2.915 | 7.906 | 18484 |
|  | Elastix A | 3.956 | 3.037 | 1.000 | 3.000 | 8.396 | 18924 |
|  | Elastix B | 3.709 | 2.816 | 1.000 | 2.915 | 7.649 | 19554 |
| FCWB | ANTs A | 4.427 | 3.643 | 1.000 | 3.240 | 9.407 | 18974 |
|  | ANTs B | 5.636 | 4.773 | 1.414 | 4.183 | 12.266 | 20592 |
|  | ANTs C | 5.450 | 4.966 | 1.000 | 3.873 | 12.207 | 20774 |
|  | CMTK A | 5.640 | 5.752 | 1.000 | 3.808 | 12.826 | 11616 |
|  | CMTK B | 6.464 | 5.187 | 1.414 | 5.050 | 12.981 | 14828 |
|  | CMTK C | 5.664 | 5.591 | 1.000 | 3.536 | 13.601 | 20246 |
|  | Elastix A | 11.371 | 8.555 | 1.225 | 10.392 | 23.611 | 14924 |
|  | Elastix B | 7.076 | 5.746 | 1.000 | 5.196 | 15.588 | 27258 |
| Tefor | ANTs A | 3.765 | 2.803 | 1.000 | 3.000 | 7.906 | 20450 |
|  | ANTs B | 6.679 | 6.131 | 1.000 | 4.301 | 15.906 | 15576 |
|  | ANTs C | 6.193 | 5.221 | 1.414 | 4.610 | 13.366 | 14578 |
|  | CMTK A | 5.613 | 8.651 | 0.707 | 2.646 | 10.794 | 15534 |
|  | CMTK B | 5.580 | 8.523 | 0.707 | 2.646 | 10.607 | 16134 |
|  | CMTK C | 5.280 | 4.433 | 1.000 | 4.000 | 11.180 | 20264 |
|  | Elastix A | 4.129 | 3.043 | 1.000 | 3.240 | 8.746 | 19768 |
|  | Elastix B | 4.306 | 3.201 | 1.000 | 3.536 | 8.718 | 20036 |

**Table S34:** GNG : adult gnathal ganglion

| Template | Algorithm | Mean | Std dev | 10th perc | median | 90th perc | N |
| --- | --- | --- | --- | --- | --- | --- | --- |
| JFRC2010 | ANTs A | 7.024 | 5.606 | 1.732 | 5.568 | 13.802 | 1889608 |
|  | ANTs B | 6.817 | 4.626 | 2.121 | 5.788 | 12.748 | 1696632 |
|  | ANTs C | 6.951 | 4.770 | 2.121 | 5.874 | 13.077 | 1812676 |
|  | CMTK A | 6.567 | 4.732 | 1.732 | 5.431 | 12.767 | 1859654 |
|  | CMTK B | 6.409 | 4.684 | 1.732 | 5.292 | 12.369 | 1736788 |
|  | CMTK C | 6.917 | 4.713 | 2.121 | 5.874 | 13.000 | 1711540 |
|  | Elastix A | 5.671 | 4.258 | 1.581 | 4.583 | 11.045 | 1588272 |
|  | Elastix B | 5.702 | 4.328 | 1.581 | 4.583 | 11.180 | 1604856 |
| JFRC2013 | ANTs A | 5.917 | 4.639 | 1.581 | 4.743 | 11.853 | 1676118 |
|  | ANTs B | 7.171 | 5.031 | 2.121 | 6.000 | 13.601 | 1611632 |
|  | ANTs C | 7.246 | 5.104 | 2.121 | 6.083 | 13.802 | 1658494 |
|  | CMTK A | 5.859 | 4.324 | 1.581 | 4.796 | 11.336 | 1614130 |
|  | CMTK B | 5.933 | 4.318 | 1.581 | 4.950 | 11.402 | 1613038 |
|  | CMTK C | 6.907 | 4.692 | 2.121 | 5.831 | 13.058 | 1507918 |
|  | Elastix A | 5.507 | 4.373 | 1.581 | 4.359 | 10.863 | 1641038 |
|  | Elastix B | 5.450 | 4.396 | 1.414 | 4.301 | 10.817 | 1621388 |
| JRC2018 | ANTs A | 5.419 | 4.431 | 1.414 | 4.243 | 10.886 | 1645444 |
|  | ANTs B | 6.766 | 4.646 | 2.000 | 5.701 | 12.826 | 1637668 |
|  | ANTs C | 6.908 | 4.798 | 2.121 | 5.788 | 13.115 | 1688056 |
|  | CMTK A | 5.416 | 4.405 | 1.414 | 4.243 | 10.817 | 1654410 |
|  | CMTK B | 5.454 | 4.421 | 1.581 | 4.301 | 10.817 | 1601386 |
|  | CMTK C | 5.715 | 4.440 | 1.581 | 4.528 | 11.136 | 1595976 |
|  | Elastix A | 5.504 | 4.367 | 1.581 | 4.359 | 10.817 | 1570950 |
|  | Elastix B | 5.533 | 4.386 | 1.581 | 4.359 | 10.886 | 1596056 |
| FCWB | ANTs A | 5.715 | 4.390 | 1.581 | 4.583 | 11.180 | 1668296 |
|  | ANTs B | 6.499 | 4.468 | 2.121 | 5.523 | 12.166 | 1722224 |
|  | ANTs C | 6.654 | 4.623 | 2.121 | 5.568 | 12.510 | 1752242 |
|  | CMTK A | 6.772 | 4.818 | 2.000 | 5.701 | 13.000 | 1771876 |
|  | CMTK B | 6.638 | 4.791 | 2.000 | 5.523 | 12.689 | 1691152 |
|  | CMTK C | 6.546 | 4.528 | 2.000 | 5.523 | 12.590 | 1636296 |
|  | Elastix A | 7.302 | 5.378 | 2.121 | 6.083 | 13.802 | 1681564 |
|  | Elastix B | 6.459 | 4.579 | 1.732 | 5.385 | 12.410 | 1917728 |
| Tefor | ANTs A | 5.610 | 4.474 | 1.581 | 4.472 | 11.158 | 1660500 |
|  | ANTs B | 7.064 | 4.853 | 2.121 | 6.000 | 13.304 | 1680796 |
|  | ANTs C | 7.435 | 5.237 | 2.121 | 6.205 | 14.213 | 1731516 |
|  | CMTK A | 6.500 | 5.088 | 1.581 | 5.148 | 13.210 | 1815200 |
|  | CMTK B | 6.564 | 5.118 | 1.581 | 5.148 | 13.285 | 1809270 |
|  | CMTK C | 7.222 | 5.560 | 2.121 | 5.831 | 13.892 | 1738810 |
|  | Elastix A | 5.597 | 4.333 | 1.581 | 4.528 | 10.977 | 1646730 |
|  | Elastix B | 6.153 | 4.573 | 1.581 | 5.000 | 12.062 | 1619762 |

**Table S35:**
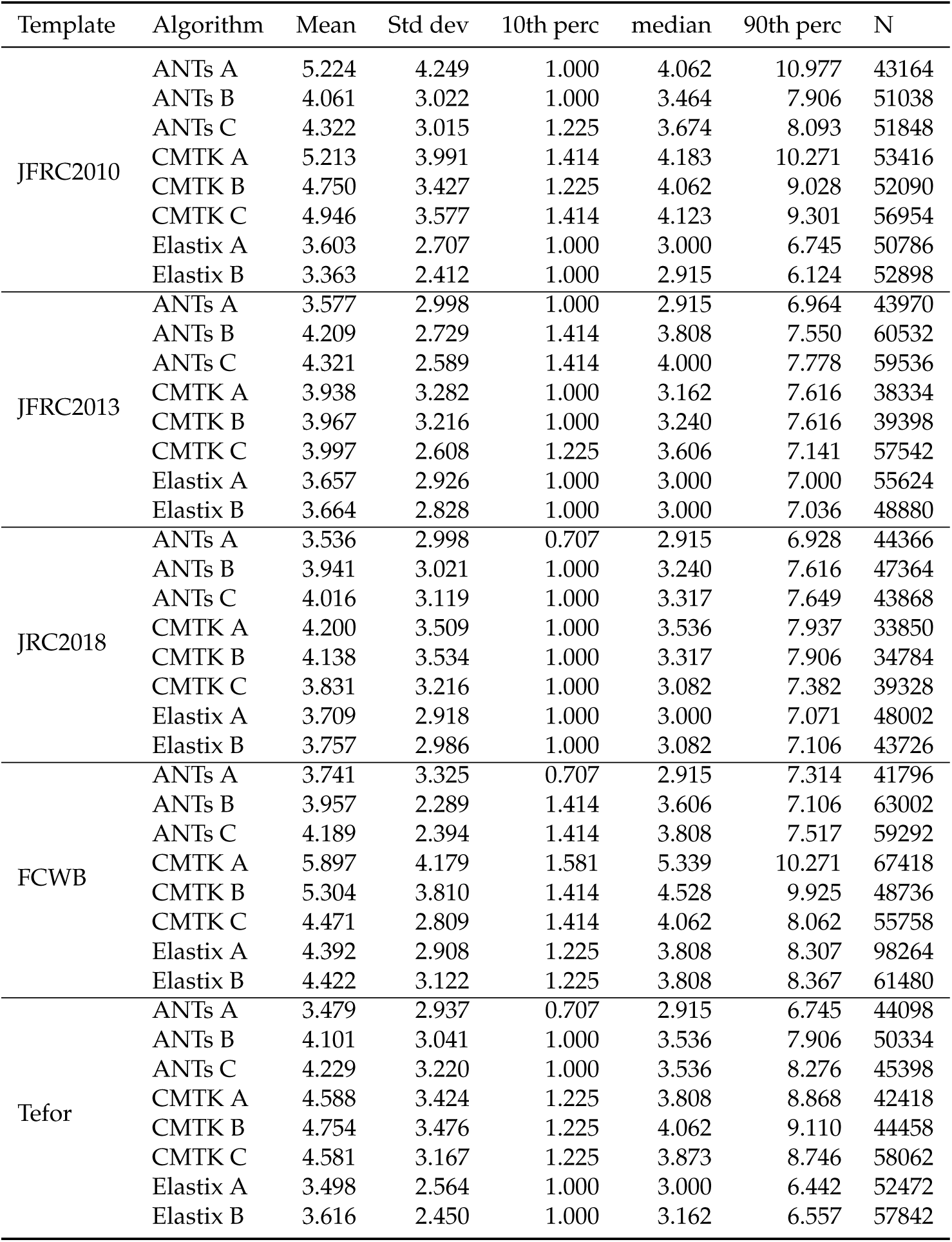
NO : nodulus

**Table S36:** PRW : prow

| Template | Algorithm | Mean | Std dev | 10th perc | median | 90th perc | N |
| --- | --- | --- | --- | --- | --- | --- | --- |
| JFRC2010 | ANTs A | 5.451 | 6.308 | 1.414 | 3.873 | 8.660 | 410994 |
|  | ANTs B | 4.951 | 3.782 | 1.581 | 4.301 | 8.544 | 390412 |
|  | ANTs C | 4.836 | 3.714 | 1.581 | 4.243 | 8.337 | 452552 |
|  | CMTK A | 4.467 | 4.126 | 1.225 | 3.674 | 7.517 | 417922 |
|  | CMTK B | 4.306 | 3.549 | 1.414 | 3.808 | 7.280 | 410878 |
|  | CMTK C | 4.613 | 3.522 | 1.414 | 4.062 | 7.906 | 383418 |
|  | Elastix A | 3.839 | 3.271 | 1.000 | 3.464 | 6.557 | 452338 |
|  | Elastix B | 3.936 | 3.128 | 1.000 | 3.536 | 6.671 | 448592 |
| JFRC2013 | ANTs A | 5.287 | 4.569 | 1.414 | 4.123 | 10.583 | 368712 |
|  | ANTs B | 7.394 | 5.886 | 2.000 | 5.523 | 16.016 | 197600 |
|  | ANTs C | 6.672 | 5.305 | 1.732 | 5.099 | 14.036 | 215962 |
|  | CMTK A | 4.308 | 3.505 | 1.000 | 3.674 | 7.550 | 425532 |
|  | CMTK B | 4.414 | 3.515 | 1.225 | 3.808 | 7.746 | 402130 |
|  | CMTK C | 5.602 | 4.139 | 1.581 | 4.583 | 10.817 | 245386 |
|  | Elastix A | 3.935 | 3.234 | 1.000 | 3.536 | 6.671 | 416116 |
|  | Elastix B | 3.938 | 3.324 | 1.000 | 3.536 | 6.708 | 419422 |
| JRC2018 | ANTs A | 3.771 | 3.098 | 1.000 | 3.317 | 6.403 | 452490 |
|  | ANTs B | 5.119 | 4.005 | 1.581 | 4.301 | 9.381 | 385182 |
|  | ANTs C | 5.495 | 4.301 | 1.581 | 4.528 | 10.247 | 403048 |
|  | CMTK A | 3.694 | 3.086 | 1.000 | 3.240 | 6.245 | 467804 |
|  | CMTK B | 3.749 | 3.136 | 1.000 | 3.317 | 6.364 | 458750 |
|  | CMTK C | 3.860 | 3.118 | 1.000 | 3.536 | 6.519 | 443426 |
|  | Elastix A | 3.770 | 3.107 | 1.000 | 3.464 | 6.364 | 441406 |
|  | Elastix B | 3.861 | 3.516 | 1.000 | 3.464 | 6.481 | 435064 |
| FCWB | ANTs A | 4.263 | 3.615 | 1.000 | 3.808 | 7.314 | 460912 |
|  | ANTs B | 6.585 | 5.218 | 1.732 | 5.099 | 13.675 | 280454 |
|  | ANTs C | 7.174 | 6.479 | 1.581 | 5.000 | 16.763 | 341950 |
|  | CMTK A | 4.270 | 3.651 | 1.000 | 3.674 | 7.649 | 463556 |
|  | CMTK B | 4.281 | 3.740 | 1.000 | 3.808 | 7.550 | 449328 |
|  | CMTK C | 4.521 | 3.681 | 1.000 | 3.808 | 8.246 | 373002 |
|  | Elastix A | 4.855 | 4.014 | 1.225 | 4.183 | 8.660 | 408028 |
|  | Elastix B | 4.065 | 3.510 | 1.000 | 3.606 | 7.106 | 532386 |
| Tefor | ANTs A | 3.862 | 3.146 | 1.000 | 3.536 | 6.557 | 446818 |
|  | ANTs B | 5.076 | 4.005 | 1.581 | 4.301 | 8.944 | 387362 |
|  | ANTs C | 5.220 | 4.459 | 1.581 | 4.243 | 9.220 | 421106 |
|  | CMTK A | 4.090 | 3.315 | 1.000 | 3.606 | 7.106 | 457468 |
|  | CMTK B | 4.399 | 3.671 | 1.225 | 3.674 | 7.681 | 445144 |
|  | CMTK C | 5.448 | 5.667 | 1.581 | 4.183 | 8.631 | 389430 |
|  | Elastix A | 3.905 | 3.120 | 1.225 | 3.536 | 6.557 | 424520 |
|  | Elastix B | 4.507 | 3.591 | 1.414 | 3.808 | 7.778 | 394342 |

**Table S37:** GA_R : gall

| Template | Algorithm | Mean | Std dev | 10th perc | median | 90th perc | N |
| --- | --- | --- | --- | --- | --- | --- | --- |
| JFRC2010 | ANTs A | 8.650 | 6.583 | 3.240 | 7.280 | 13.058 | 12776 |
|  | ANTs B | 6.726 | 3.628 | 2.121 | 6.325 | 11.853 | 5332 |
|  | ANTs C | 6.919 | 3.736 | 2.236 | 6.442 | 12.124 | 5164 |
|  | CMTK A | 8.269 | 4.077 | 3.240 | 7.921 | 13.675 | 4902 |
|  | CMTK B | 8.348 | 4.733 | 2.828 | 7.649 | 15.100 | 5636 |
|  | CMTK C | 8.120 | 4.036 | 3.082 | 7.649 | 13.802 | 4300 |
|  | Elastix A | 7.205 | 4.097 | 2.121 | 6.671 | 12.683 | 5184 |
|  | Elastix B | 7.425 | 4.349 | 2.000 | 7.106 | 13.019 | 5252 |
| JFRC2013 | ANTs A | 6.483 | 3.928 | 1.732 | 5.745 | 11.937 | 6140 |
|  | ANTs B | 7.392 | 3.944 | 2.636 | 6.856 | 13.058 | 5440 |
|  | ANTs C | 7.707 | 3.666 | 3.082 | 7.280 | 12.865 | 5362 |
|  | CMTK A | 7.079 | 3.788 | 2.550 | 6.557 | 12.186 | 6426 |
|  | CMTK B | 7.120 | 4.025 | 2.449 | 6.519 | 12.530 | 6260 |
|  | CMTK C | 8.871 | 3.816 | 3.606 | 8.888 | 13.675 | 4532 |
|  | Elastix A | 6.524 | 3.889 | 2.121 | 5.831 | 11.726 | 5546 |
|  | Elastix B | 6.757 | 4.029 | 2.121 | 6.083 | 12.104 | 5690 |
| JRC2018 | ANTs A | 6.542 | 4.016 | 1.732 | 5.745 | 12.104 | 6208 |
|  | ANTs B | 6.646 | 4.030 | 1.732 | 5.895 | 12.166 | 6202 |
|  | ANTs C | 6.856 | 4.018 | 2.121 | 6.083 | 12.367 | 6582 |
|  | CMTK A | 6.834 | 4.189 | 2.121 | 6.083 | 12.245 | 5732 |
|  | CMTK B | 6.755 | 4.151 | 2.000 | 5.916 | 12.430 | 6162 |
|  | CMTK C | 6.748 | 4.089 | 2.121 | 6.083 | 12.186 | 5888 |
|  | Elastix A | 6.671 | 4.026 | 2.000 | 6.000 | 12.104 | 5226 |
|  | Elastix B | 7.546 | 4.414 | 2.345 | 6.708 | 13.600 | 5462 |
| FCWB | ANTs A | 6.297 | 3.421 | 2.000 | 5.916 | 11.136 | 5590 |
|  | ANTs B | 8.126 | 4.053 | 2.550 | 8.031 | 13.611 | 3196 |
|  | ANTs C | 8.335 | 4.082 | 2.915 | 8.124 | 13.946 | 4078 |
|  | CMTK A | 7.034 | 4.278 | 2.345 | 6.124 | 12.767 | 6136 |
|  | CMTK B | 6.829 | 4.076 | 2.236 | 6.083 | 12.124 | 5632 |
|  | CMTK C | 7.609 | 3.586 | 3.000 | 7.550 | 12.349 | 4046 |
|  | Elastix A | 8.167 | 4.621 | 2.646 | 7.550 | 14.526 | 6008 |
|  | Elastix B | 7.350 | 4.343 | 2.236 | 6.442 | 13.601 | 5688 |
| Tefor | ANTs A | 6.669 | 4.165 | 1.732 | 5.831 | 12.104 | 6206 |
|  | ANTs B | 6.988 | 4.027 | 2.000 | 6.442 | 12.081 | 6332 |
|  | ANTs C | 7.073 | 3.991 | 2.236 | 6.557 | 12.590 | 7242 |
|  | CMTK A | 7.247 | 3.744 | 2.449 | 7.000 | 12.166 | 7508 |
|  | CMTK B | 7.146 | 3.944 | 2.550 | 6.557 | 12.410 | 6372 |
|  | CMTK C | 7.636 | 3.922 | 2.828 | 7.106 | 12.748 | 5630 |
|  | Elastix A | 6.943 | 4.088 | 2.000 | 6.519 | 12.349 | 5154 |
|  | Elastix B | 7.311 | 4.060 | 2.345 | 6.856 | 12.826 | 5682 |

**Table S38:** AME_L : accessory medulla

| Template | Algorithm | Mean | Std dev | 10th perc | median | 90th perc | N |
| --- | --- | --- | --- | --- | --- | --- | --- |
| JFRC2010 | ANTs A | 6.332 | 6.591 | 1.414 | 4.183 | 15.556 | 30288 |
|  | ANTs B | 4.387 | 2.400 | 1.581 | 4.123 | 7.649 | 26644 |
|  | ANTs C | 4.460 | 2.496 | 1.581 | 4.183 | 7.778 | 27714 |
|  | CMTK A | 5.377 | 4.685 | 1.414 | 4.123 | 10.100 | 29566 |
|  | CMTK B | 5.202 | 4.786 | 1.414 | 4.000 | 9.381 | 32256 |
|  | CMTK C | 6.248 | 5.083 | 2.000 | 4.950 | 11.958 | 23192 |
|  | Elastix A | 4.103 | 2.433 | 1.225 | 3.808 | 7.382 | 37556 |
|  | Elastix B | 4.014 | 2.260 | 1.225 | 3.808 | 7.106 | 32506 |
| JFRC2013 | ANTs A | 3.770 | 2.246 | 1.225 | 3.536 | 6.819 | 31780 |
|  | ANTs B | 4.478 | 2.626 | 1.581 | 4.123 | 7.810 | 20492 |
|  | ANTs C | 4.489 | 2.555 | 1.581 | 4.123 | 8.016 | 24386 |
|  | CMTK A | 4.145 | 2.878 | 1.225 | 3.606 | 7.382 | 33056 |
|  | CMTK B | 4.430 | 2.991 | 1.414 | 3.808 | 8.031 | 31064 |
|  | CMTK C | 4.468 | 2.792 | 1.581 | 4.062 | 7.550 | 24506 |
|  | Elastix A | 3.665 | 2.178 | 1.000 | 3.317 | 6.557 | 35180 |
|  | Elastix B | 3.831 | 2.174 | 1.225 | 3.606 | 6.708 | 34084 |
| JRC2018 | ANTs A | 3.791 | 2.250 | 1.000 | 3.536 | 6.964 | 34898 |
|  | ANTs B | 3.872 | 2.352 | 1.000 | 3.536 | 7.106 | 32926 |
|  | ANTs C | 4.057 | 2.519 | 1.000 | 3.606 | 7.416 | 32226 |
|  | CMTK A | 3.733 | 2.176 | 1.225 | 3.464 | 6.671 | 35186 |
|  | CMTK B | 3.774 | 2.195 | 1.225 | 3.464 | 6.745 | 35516 |
|  | CMTK C | 3.781 | 2.232 | 1.000 | 3.536 | 6.856 | 35504 |
|  | Elastix A | 3.832 | 2.249 | 1.225 | 3.536 | 6.856 | 36706 |
|  | Elastix B | 3.967 | 2.208 | 1.414 | 3.674 | 6.964 | 34706 |
| FCWB | ANTs A | 3.977 | 2.203 | 1.225 | 3.808 | 6.964 | 30514 |
|  | ANTs B | 4.686 | 2.763 | 1.581 | 4.243 | 8.307 | 18914 |
|  | ANTs C | 4.556 | 2.921 | 1.225 | 4.062 | 8.544 | 20840 |
|  | CMTK A | 4.182 | 2.362 | 1.414 | 3.808 | 7.382 | 31992 |
|  | CMTK B | 4.035 | 2.340 | 1.414 | 3.606 | 7.246 | 31278 |
|  | CMTK C | 5.682 | 4.936 | 1.581 | 4.359 | 10.536 | 18984 |
|  | Elastix A | 5.871 | 6.539 | 1.581 | 4.472 | 10.794 | 27614 |
|  | Elastix B | 4.667 | 6.330 | 1.225 | 3.606 | 7.382 | 36610 |
| Tefor | ANTs A | 3.938 | 2.297 | 1.225 | 3.606 | 7.106 | 31502 |
|  | ANTs B | 4.125 | 2.447 | 1.225 | 3.808 | 7.517 | 30136 |
|  | ANTs C | 4.171 | 2.517 | 1.225 | 3.808 | 7.616 | 31770 |
|  | CMTK A | 4.897 | 3.875 | 1.414 | 3.873 | 9.487 | 33304 |
|  | CMTK B | 4.968 | 3.794 | 1.414 | 4.062 | 9.747 | 33040 |
|  | CMTK C | 4.499 | 3.312 | 1.225 | 3.606 | 8.986 | 27266 |
|  | Elastix A | 4.147 | 2.548 | 1.000 | 3.808 | 7.649 | 34694 |
|  | Elastix B | 4.103 | 2.512 | 1.225 | 3.674 | 7.550 | 33598 |

**Table S39:**
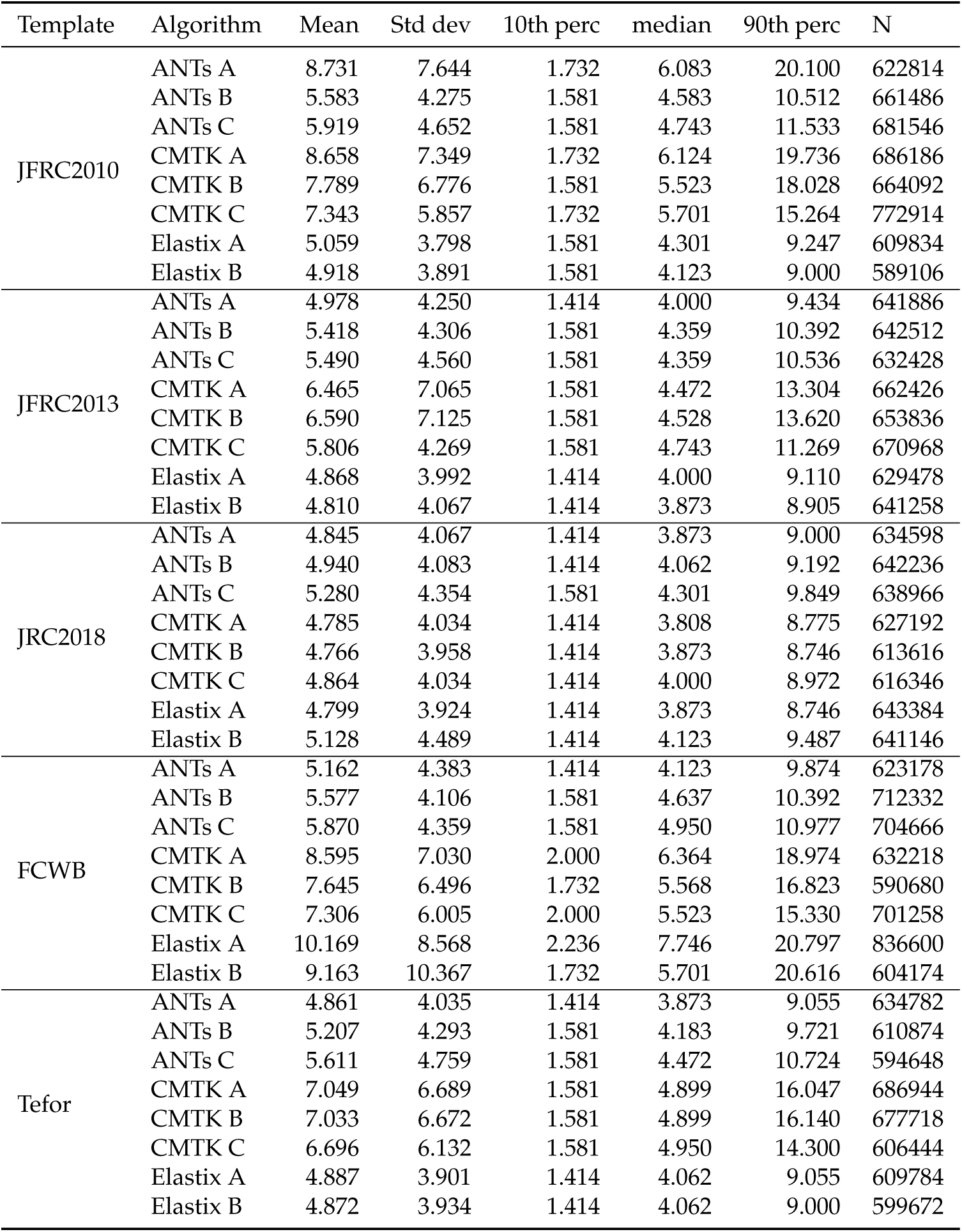
LO_L : lobula

**Table S40:**
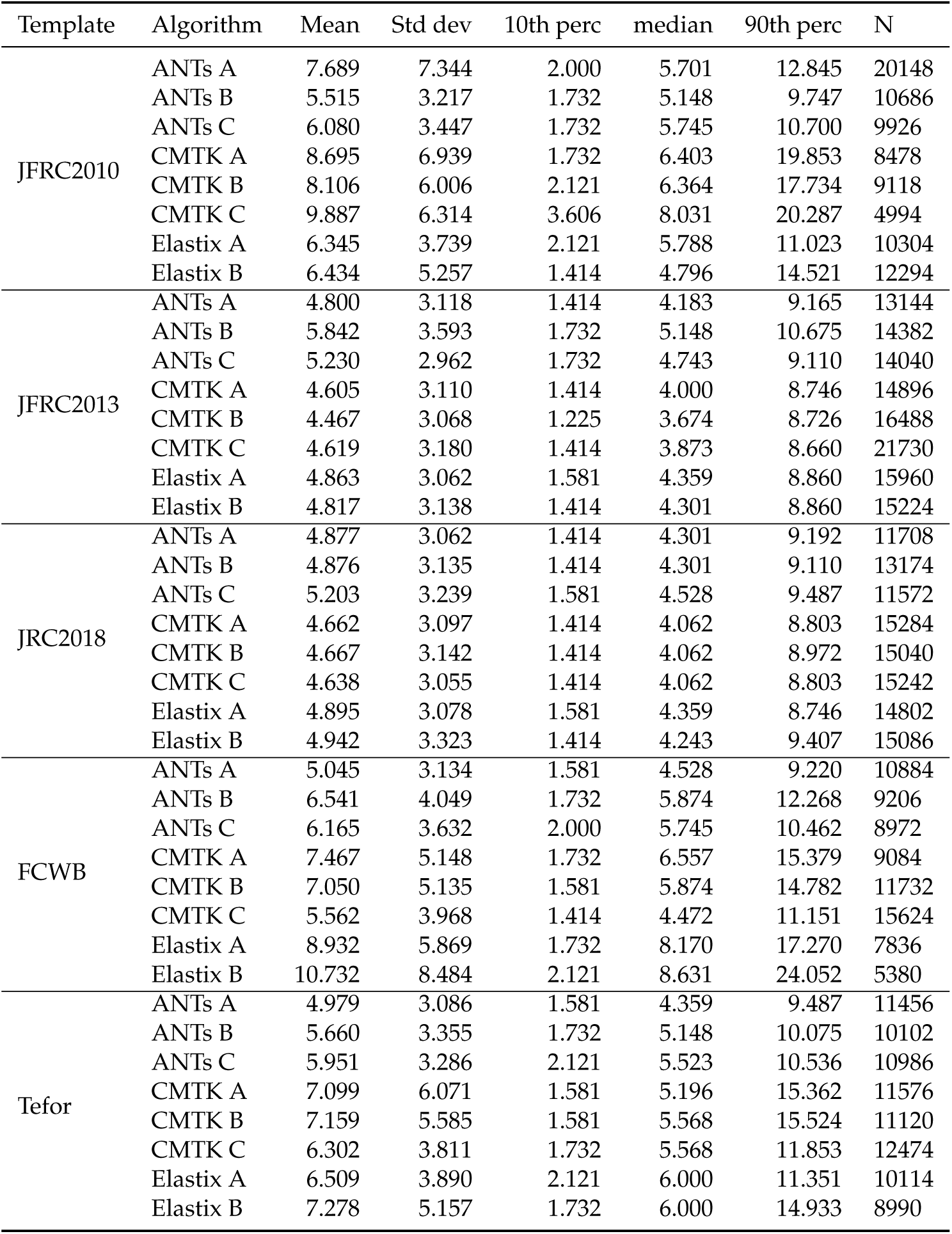
BU_L : bulb

| Template | Algorithm | Mean | Std dev | 10th perc | median | 90th perc | N |
| --- | --- | --- | --- | --- | --- | --- | --- |
| JFRC2010 | ANTs A | 7.689 | 7.344 | 2.000 | 5.701 | 12.845 | 20148 |
|  | ANTs B | 5.515 | 3.217 | 1.732 | 5.148 | 9.747 | 10686 |
|  | ANTs C | 6.080 | 3.447 | 1.732 | 5.745 | 10.700 | 9926 |
|  | CMTK A | 8.695 | 6.939 | 1.732 | 6.403 | 19.853 | 8478 |
|  | CMTK B | 8.106 | 6.006 | 2.121 | 6.364 | 17.734 | 9118 |
|  | CMTK C | 9.887 | 6.314 | 3.606 | 8.031 | 20.287 | 4994 |
|  | Elastix A | 6.345 | 3.739 | 2.121 | 5.788 | 11.023 | 10304 |
|  | Elastix B | 6.434 | 5.257 | 1.414 | 4.796 | 14.521 | 12294 |
| JFRC2013 | ANTs A | 4.800 | 3.118 | 1.414 | 4.183 | 9.165 | 13144 |
|  | ANTs B | 5.842 | 3.593 | 1.732 | 5.148 | 10.675 | 14382 |
|  | ANTs C | 5.230 | 2.962 | 1.732 | 4.743 | 9.110 | 14040 |
|  | CMTK A | 4.605 | 3.110 | 1.414 | 4.000 | 8.746 | 14896 |
|  | CMTK B | 4.467 | 3.068 | 1.225 | 3.674 | 8.726 | 16488 |
|  | CMTK C | 4.619 | 3.180 | 1.414 | 3.873 | 8.660 | 21730 |
|  | Elastix A | 4.863 | 3.062 | 1.581 | 4.359 | 8.860 | 15960 |
|  | Elastix B | 4.817 | 3.138 | 1.414 | 4.301 | 8.860 | 15224 |
| JRC2018 | ANTs A | 4.877 | 3.062 | 1.414 | 4.301 | 9.192 | 11708 |
|  | ANTs B | 4.876 | 3.135 | 1.414 | 4.301 | 9.110 | 13174 |
|  | ANTs C | 5.203 | 3.239 | 1.581 | 4.528 | 9.487 | 11572 |
|  | CMTK A | 4.662 | 3.097 | 1.414 | 4.062 | 8.803 | 15284 |
|  | CMTK B | 4.667 | 3.142 | 1.414 | 4.062 | 8.972 | 15040 |
|  | CMTK C | 4.638 | 3.055 | 1.414 | 4.062 | 8.803 | 15242 |
|  | Elastix A | 4.895 | 3.078 | 1.581 | 4.359 | 8.746 | 14802 |
|  | Elastix B | 4.942 | 3.323 | 1.414 | 4.243 | 9.407 | 15086 |
| FCWB | ANTs A | 5.045 | 3.134 | 1.581 | 4.528 | 9.220 | 10884 |
|  | ANTs B | 6.541 | 4.049 | 1.732 | 5.874 | 12.268 | 9206 |
|  | ANTs C | 6.165 | 3.632 | 2.000 | 5.745 | 10.462 | 8972 |
|  | CMTK A | 7.467 | 5.148 | 1.732 | 6.557 | 15.379 | 9084 |
|  | CMTK B | 7.050 | 5.135 | 1.581 | 5.874 | 14.782 | 11732 |
|  | CMTK C | 5.562 | 3.968 | 1.414 | 4.472 | 11.151 | 15624 |
|  | Elastix A | 8.932 | 5.869 | 1.732 | 8.170 | 17.270 | 7836 |
|  | Elastix B | 10.732 | 8.484 | 2.121 | 8.631 | 24.052 | 5380 |
| Tefor | ANTs A | 4.979 | 3.086 | 1.581 | 4.359 | 9.487 | 11456 |
|  | ANTs B | 5.660 | 3.355 | 1.732 | 5.148 | 10.075 | 10102 |
|  | ANTs C | 5.951 | 3.286 | 2.121 | 5.523 | 10.536 | 10986 |
|  | CMTK A | 7.099 | 6.071 | 1.581 | 5.196 | 15.362 | 11576 |
|  | CMTK B | 7.159 | 5.585 | 1.581 | 5.568 | 15.524 | 11120 |
|  | CMTK C | 6.302 | 3.811 | 1.732 | 5.568 | 11.853 | 12474 |
|  | Elastix A | 6.509 | 3.890 | 2.121 | 6.000 | 11.351 | 10114 |
|  | Elastix B | 7.278 | 5.157 | 1.732 | 6.000 | 14.933 | 8990 |

**Table S41:** LH_L : lateral horn

| Template | Algorithm | Mean | Std dev | 10th perc | median | 90th perc | N |
| --- | --- | --- | --- | --- | --- | --- | --- |
| JFRC2010 | ANTs A | 6.131 | 4.686 | 1.581 | 5.000 | 12.104 | 273626 |
|  | ANTs B | 5.764 | 4.971 | 1.414 | 4.528 | 11.402 | 271974 |
|  | ANTs C | 6.337 | 5.179 | 1.581 | 5.000 | 12.430 | 282188 |
|  | CMTK A | 6.438 | 4.694 | 1.581 | 5.339 | 12.826 | 438598 |
|  | CMTK B | 6.349 | 4.623 | 1.581 | 5.196 | 12.530 | 401034 |
|  | CMTK C | 7.004 | 4.965 | 1.732 | 5.788 | 13.946 | 397448 |
|  | Elastix A | 4.973 | 3.639 | 1.414 | 4.183 | 9.327 | 269426 |
|  | Elastix B | 5.390 | 3.897 | 1.581 | 4.528 | 10.271 | 277598 |
| JFRC2013 | ANTs A | 4.914 | 3.565 | 1.225 | 4.183 | 9.327 | 305420 |
|  | ANTs B | 6.759 | 5.441 | 1.732 | 5.431 | 13.077 | 268188 |
|  | ANTs C | 6.399 | 5.541 | 1.581 | 5.000 | 12.530 | 295292 |
|  | CMTK A | 5.221 | 3.812 | 1.414 | 4.359 | 10.124 | 329340 |
|  | CMTK B | 5.047 | 3.592 | 1.414 | 4.243 | 9.670 | 321452 |
|  | CMTK C | 5.661 | 4.459 | 1.225 | 4.528 | 11.511 | 344122 |
|  | Elastix A | 4.969 | 3.514 | 1.414 | 4.183 | 9.460 | 272544 |
|  | Elastix B | 4.888 | 3.525 | 1.414 | 4.123 | 9.220 | 295686 |
| JRC2018 | ANTs A | 4.787 | 3.415 | 1.225 | 4.123 | 9.028 | 320566 |
|  | ANTs B | 5.584 | 4.404 | 1.414 | 4.528 | 10.794 | 297424 |
|  | ANTs C | 5.671 | 4.244 | 1.581 | 4.690 | 11.068 | 326468 |
|  | CMTK A | 4.845 | 3.348 | 1.414 | 4.183 | 9.000 | 322432 |
|  | CMTK B | 4.854 | 3.356 | 1.414 | 4.183 | 9.028 | 309956 |
|  | CMTK C | 5.107 | 3.580 | 1.414 | 4.359 | 9.618 | 297512 |
|  | Elastix A | 4.932 | 3.403 | 1.414 | 4.243 | 9.220 | 286506 |
|  | Elastix B | 5.025 | 3.441 | 1.414 | 4.359 | 9.301 | 300684 |
| FCWB | ANTs A | 5.304 | 3.605 | 1.414 | 4.583 | 10.050 | 333478 |
|  | ANTs B | 6.691 | 5.234 | 1.732 | 5.431 | 12.590 | 251700 |
|  | ANTs C | 6.766 | 5.062 | 1.732 | 5.568 | 13.229 | 280818 |
|  | CMTK A | 6.283 | 5.506 | 1.581 | 5.000 | 11.489 | 283790 |
|  | CMTK B | 5.714 | 4.789 | 1.581 | 4.743 | 10.223 | 292828 |
|  | CMTK C | 6.162 | 4.331 | 1.732 | 5.196 | 11.619 | 272040 |
|  | Elastix A | 7.355 | 5.445 | 1.732 | 6.000 | 14.933 | 240540 |
|  | Elastix B | 6.728 | 6.016 | 1.581 | 5.148 | 13.210 | 300976 |
| Tefor | ANTs A | 4.792 | 3.468 | 1.225 | 4.123 | 9.028 | 318946 |
|  | ANTs B | 5.730 | 4.661 | 1.414 | 4.583 | 11.091 | 292186 |
|  | ANTs C | 6.158 | 5.294 | 1.581 | 4.796 | 11.853 | 323302 |
|  | CMTK A | 7.814 | 9.068 | 1.414 | 4.690 | 19.812 | 270274 |
|  | CMTK B | 8.033 | 9.262 | 1.414 | 4.743 | 21.083 | 266200 |
|  | CMTK C | 6.962 | 7.775 | 1.414 | 4.743 | 13.675 | 283214 |
|  | Elastix A | 4.958 | 3.537 | 1.414 | 4.183 | 9.327 | 291076 |
|  | Elastix B | 5.504 | 3.924 | 1.581 | 4.690 | 10.512 | 291692 |

**Table S42:**
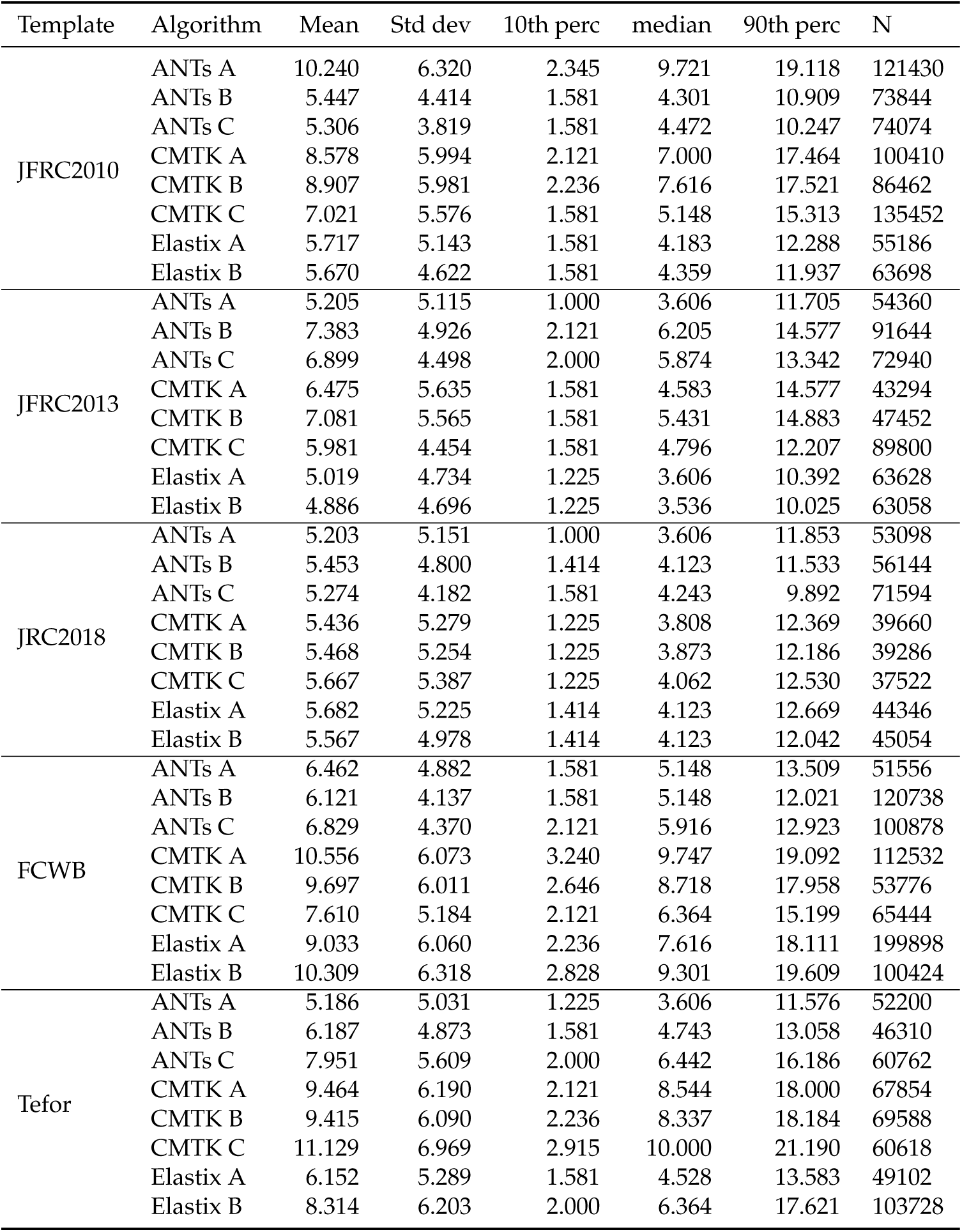
LAL_L : lateral accessory lobe

| Template | Algorithm | Mean | Std dev | 10th perc | median | 90th perc | N |
| --- | --- | --- | --- | --- | --- | --- | --- |
| JFRC2010 | ANTs A | 10.240 | 6.320 | 2.345 | 9.721 | 19.118 | 121430 |
|  | ANTs B | 5.447 | 4.414 | 1.581 | 4.301 | 10.909 | 73844 |
|  | ANTs C | 5.306 | 3.819 | 1.581 | 4.472 | 10.247 | 74074 |
|  | CMTK A | 8.578 | 5.994 | 2.121 | 7.000 | 17.464 | 100410 |
|  | CMTK B | 8.907 | 5.981 | 2.236 | 7.616 | 17.521 | 86462 |
|  | CMTK C | 7.021 | 5.576 | 1.581 | 5.148 | 15.313 | 135452 |
|  | Elastix A | 5.717 | 5.143 | 1.581 | 4.183 | 12.288 | 55186 |
|  | Elastix B | 5.670 | 4.622 | 1.581 | 4.359 | 11.937 | 63698 |
| JFRC2013 | ANTs A | 5.205 | 5.115 | 1.000 | 3.606 | 11.705 | 54360 |
|  | ANTs B | 7.383 | 4.926 | 2.121 | 6.205 | 14.577 | 91644 |
|  | ANTs C | 6.899 | 4.498 | 2.000 | 5.874 | 13.342 | 72940 |
|  | CMTK A | 6.475 | 5.635 | 1.581 | 4.583 | 14.577 | 43294 |
|  | CMTK B | 7.081 | 5.565 | 1.581 | 5.431 | 14.883 | 47452 |
|  | CMTK C | 5.981 | 4.454 | 1.581 | 4.796 | 12.207 | 89800 |
|  | Elastix A | 5.019 | 4.734 | 1.225 | 3.606 | 10.392 | 63628 |
|  | Elastix B | 4.886 | 4.696 | 1.225 | 3.536 | 10.025 | 63058 |
| JRC2018 | ANTs A | 5.203 | 5.151 | 1.000 | 3.606 | 11.853 | 53098 |
|  | ANTs B | 5.453 | 4.800 | 1.414 | 4.123 | 11.533 | 56144 |
|  | ANTs C | 5.274 | 4.182 | 1.581 | 4.243 | 9.892 | 71594 |
|  | CMTK A | 5.436 | 5.279 | 1.225 | 3.808 | 12.369 | 39660 |
|  | CMTK B | 5.468 | 5.254 | 1.225 | 3.873 | 12.186 | 39286 |
|  | CMTK C | 5.667 | 5.387 | 1.225 | 4.062 | 12.530 | 37522 |
|  | Elastix A | 5.682 | 5.225 | 1.414 | 4.123 | 12.669 | 44346 |
|  | Elastix B | 5.567 | 4.978 | 1.414 | 4.123 | 12.042 | 45054 |
| FCWB | ANTs A | 6.462 | 4.882 | 1.581 | 5.148 | 13.509 | 51556 |
|  | ANTs B | 6.121 | 4.137 | 1.581 | 5.148 | 12.021 | 120738 |
|  | ANTs C | 6.829 | 4.370 | 2.121 | 5.916 | 12.923 | 100878 |
|  | CMTK A | 10.556 | 6.073 | 3.240 | 9.747 | 19.092 | 112532 |
|  | CMTK B | 9.697 | 6.011 | 2.646 | 8.718 | 17.958 | 53776 |
|  | CMTK C | 7.610 | 5.184 | 2.121 | 6.364 | 15.199 | 65444 |
|  | Elastix A | 9.033 | 6.060 | 2.236 | 7.616 | 18.111 | 199898 |
|  | Elastix B | 10.309 | 6.318 | 2.828 | 9.301 | 19.609 | 100424 |
| Tefor | ANTs A | 5.186 | 5.031 | 1.225 | 3.606 | 11.576 | 52200 |
|  | ANTs B | 6.187 | 4.873 | 1.581 | 4.743 | 13.058 | 46310 |
|  | ANTs C | 7.951 | 5.609 | 2.000 | 6.442 | 16.186 | 60762 |
|  | CMTK A | 9.464 | 6.190 | 2.121 | 8.544 | 18.000 | 67854 |
|  | CMTK B | 9.415 | 6.090 | 2.236 | 8.337 | 18.184 | 69588 |
|  | CMTK C | 11.129 | 6.969 | 2.915 | 10.000 | 21.190 | 60618 |
|  | Elastix A | 6.152 | 5.289 | 1.581 | 4.528 | 13.583 | 49102 |
|  | Elastix B | 8.314 | 6.203 | 2.000 | 6.364 | 17.621 | 103728 |

**Table S43:** CAN_L : cantle

| Template | Algorithm | Mean | Std dev | 10th perc | median | 90th perc | N |
| --- | --- | --- | --- | --- | --- | --- | --- |
| JFRC2010 | ANTs A | 9.178 | 8.034 | 1.732 | 6.519 | 22.471 | 19372 |
|  | ANTs B | 4.415 | 2.860 | 1.225 | 3.873 | 8.276 | 29064 |
|  | ANTs C | 4.149 | 3.045 | 1.000 | 3.464 | 8.307 | 47732 |
|  | CMTK A | 6.167 | 4.918 | 1.414 | 4.950 | 12.767 | 55044 |
|  | CMTK B | 5.427 | 3.647 | 1.414 | 4.690 | 10.440 | 48774 |
|  | CMTK C | 5.454 | 3.460 | 1.581 | 4.975 | 9.823 | 40106 |
|  | Elastix A | 4.994 | 2.999 | 1.581 | 4.528 | 8.972 | 16106 |
|  | Elastix B | 4.680 | 2.960 | 1.414 | 4.243 | 8.544 | 22274 |
| JFRC2013 | ANTs A | 4.549 | 3.076 | 1.225 | 3.873 | 8.631 | 23450 |
|  | ANTs B | 3.661 | 2.605 | 1.000 | 3.082 | 7.071 | 43696 |
|  | ANTs C | 3.624 | 2.660 | 1.000 | 3.000 | 7.280 | 42390 |
|  | CMTK A | 4.543 | 3.047 | 1.414 | 3.873 | 8.738 | 24904 |
|  | CMTK B | 4.474 | 3.080 | 1.225 | 3.808 | 8.573 | 25474 |
|  | CMTK C | 4.436 | 2.772 | 1.414 | 3.873 | 8.062 | 40470 |
|  | Elastix A | 4.353 | 3.057 | 1.000 | 3.674 | 8.515 | 26294 |
|  | Elastix B | 4.620 | 3.131 | 1.225 | 4.000 | 8.775 | 24252 |
| JRC2018 | ANTs A | 4.378 | 3.041 | 1.225 | 3.674 | 8.485 | 24512 |
|  | ANTs B | 4.506 | 3.037 | 1.225 | 3.808 | 8.515 | 24614 |
|  | ANTs C | 4.316 | 2.850 | 1.000 | 3.808 | 8.093 | 28654 |
|  | CMTK A | 4.710 | 3.125 | 1.414 | 4.062 | 8.888 | 26174 |
|  | CMTK B | 4.584 | 3.055 | 1.225 | 3.873 | 8.746 | 25826 |
|  | CMTK C | 4.543 | 3.108 | 1.225 | 3.808 | 8.746 | 25846 |
|  | Elastix A | 4.491 | 3.149 | 1.225 | 3.808 | 8.631 | 24382 |
|  | Elastix B | 4.691 | 3.151 | 1.414 | 4.062 | 8.746 | 24012 |
| FCWB | ANTs A | 4.290 | 2.902 | 1.225 | 3.606 | 8.185 | 24878 |
|  | ANTs B | 4.211 | 3.012 | 1.000 | 3.536 | 8.185 | 27174 |
|  | ANTs C | 3.897 | 2.760 | 1.000 | 3.317 | 7.382 | 28356 |
|  | CMTK A | 4.296 | 2.864 | 1.000 | 3.808 | 8.062 | 73796 |
|  | CMTK B | 4.522 | 3.127 | 1.225 | 3.873 | 8.367 | 39270 |
|  | CMTK C | 4.203 | 2.886 | 1.000 | 3.606 | 8.093 | 45984 |
|  | Elastix A | 4.496 | 3.587 | 1.225 | 3.808 | 8.062 | 47902 |
|  | Elastix B | 4.921 | 3.166 | 1.414 | 4.301 | 9.407 | 50172 |
| Tefor | ANTs A | 4.597 | 3.061 | 1.414 | 3.873 | 8.660 | 23362 |
|  | ANTs B | 4.360 | 2.945 | 1.414 | 3.674 | 8.276 | 22966 |
|  | ANTs C | 4.423 | 2.995 | 1.000 | 3.808 | 8.426 | 34566 |
|  | CMTK A | 4.499 | 2.806 | 1.414 | 4.062 | 8.155 | 30862 |
|  | CMTK B | 4.396 | 2.804 | 1.414 | 3.873 | 8.093 | 30122 |
|  | CMTK C | 5.168 | 3.147 | 1.581 | 4.743 | 9.513 | 20416 |
|  | Elastix A | 4.475 | 2.692 | 1.414 | 4.123 | 8.031 | 16314 |
|  | Elastix B | 4.425 | 2.658 | 1.414 | 4.062 | 7.906 | 27170 |

**Table S44:** AMMC_L : antennal mechanosensory and motor center

| Template | Algorithm | Mean | Std dev | 10th perc | median | 90th perc | N |
| --- | --- | --- | --- | --- | --- | --- | --- |
| JFRC2010 | ANTs A | 5.702 | 5.274 | 1.581 | 4.472 | 10.817 | 199226 |
|  | ANTs B | 5.715 | 6.225 | 1.581 | 4.301 | 9.618 | 165386 |
|  | ANTs C | 5.753 | 5.989 | 1.581 | 4.528 | 9.618 | 187140 |
|  | CMTK A | 5.132 | 5.145 | 1.414 | 4.062 | 8.803 | 218338 |
|  | CMTK B | 4.996 | 4.845 | 1.414 | 4.062 | 8.631 | 218734 |
|  | CMTK C | 5.457 | 5.774 | 1.581 | 4.183 | 9.407 | 200846 |
|  | Elastix A | 4.937 | 5.199 | 1.225 | 3.808 | 8.426 | 189012 |
|  | Elastix B | 4.852 | 4.941 | 1.225 | 3.808 | 8.246 | 200538 |
| JFRC2013 | ANTs A | 4.993 | 5.454 | 1.225 | 3.808 | 8.602 | 178108 |
|  | ANTs B | 6.124 | 6.457 | 1.581 | 4.528 | 10.607 | 144664 |
|  | ANTs C | 5.968 | 6.282 | 1.581 | 4.359 | 10.440 | 150050 |
|  | CMTK A | 5.528 | 5.617 | 1.581 | 4.243 | 9.644 | 173606 |
|  | CMTK B | 5.683 | 5.882 | 1.581 | 4.359 | 9.950 | 164712 |
|  | CMTK C | 6.089 | 5.811 | 1.581 | 4.637 | 10.770 | 145314 |
|  | Elastix A | 4.909 | 5.689 | 1.225 | 3.606 | 8.276 | 192508 |
|  | Elastix B | 4.845 | 5.514 | 1.225 | 3.606 | 8.185 | 198670 |
| JRC2018 | ANTs A | 4.730 | 5.167 | 1.000 | 3.674 | 7.937 | 197440 |
|  | ANTs B | 5.373 | 5.741 | 1.414 | 4.123 | 9.301 | 181890 |
|  | ANTs C | 5.681 | 5.736 | 1.581 | 4.359 | 10.050 | 192824 |
|  | CMTK A | 4.842 | 5.356 | 1.000 | 3.674 | 8.185 | 197642 |
|  | CMTK B | 4.868 | 5.361 | 1.000 | 3.674 | 8.307 | 194334 |
|  | CMTK C | 4.977 | 5.309 | 1.225 | 3.808 | 8.515 | 193026 |
|  | Elastix A | 4.875 | 5.211 | 1.225 | 3.808 | 8.185 | 195568 |
|  | Elastix B | 5.161 | 5.565 | 1.225 | 3.873 | 8.860 | 197934 |
| FCWB | ANTs A | 5.020 | 5.213 | 1.225 | 3.808 | 8.746 | 191558 |
|  | ANTs B | 5.986 | 6.269 | 1.581 | 4.528 | 10.607 | 145952 |
|  | ANTs C | 6.040 | 6.381 | 1.581 | 4.528 | 10.607 | 155918 |
|  | CMTK A | 5.365 | 4.788 | 1.581 | 4.359 | 9.247 | 192852 |
|  | CMTK B | 5.442 | 4.986 | 1.581 | 4.359 | 9.434 | 193322 |
|  | CMTK C | 5.592 | 5.494 | 1.581 | 4.243 | 9.950 | 173954 |
|  | Elastix A | 6.366 | 7.889 | 1.581 | 4.583 | 10.607 | 194944 |
|  | Elastix B | 5.446 | 6.128 | 1.414 | 4.123 | 9.301 | 198834 |
| Tefor | ANTs A | 4.738 | 5.086 | 1.225 | 3.674 | 7.937 | 199464 |
|  | ANTs B | 5.561 | 6.031 | 1.581 | 4.183 | 9.434 | 174346 |
|  | ANTs C | 5.980 | 6.428 | 1.581 | 4.472 | 10.512 | 183560 |
|  | CMTK A | 5.149 | 5.134 | 1.225 | 4.000 | 9.434 | 218888 |
|  | CMTK B | 5.087 | 5.109 | 1.225 | 4.062 | 9.000 | 219208 |
|  | CMTK C | 5.239 | 5.104 | 1.581 | 4.183 | 8.972 | 207136 |
|  | Elastix A | 4.970 | 5.408 | 1.225 | 3.808 | 8.515 | 197682 |
|  | Elastix B | 4.964 | 4.944 | 1.414 | 4.062 | 8.337 | 210416 |

**Table S45:** ICL_L : inferior clamp

| Template | Algorithm | Mean | Std dev | 10th perc | median | 90th perc | N |
| --- | --- | --- | --- | --- | --- | --- | --- |
| JFRC2010 | ANTs A | 4.630 | 5.270 | 0.707 | 3.000 | 9.874 | 320720 |
|  | ANTs B | 3.816 | 3.557 | 0.707 | 2.646 | 8.573 | 258504 |
|  | ANTs C | 4.282 | 3.797 | 0.707 | 3.162 | 9.460 | 250196 |
|  | CMTK A | 5.171 | 6.012 | 0.707 | 2.646 | 14.577 | 265902 |
|  | CMTK B | 4.996 | 5.497 | 0.707 | 2.828 | 13.134 | 270070 |
|  | CMTK C | 5.192 | 5.249 | 0.707 | 3.240 | 12.610 | 266652 |
|  | Elastix A | 3.301 | 2.947 | 0.707 | 2.345 | 7.314 | 263970 |
|  | Elastix B | 3.519 | 3.570 | 0.707 | 2.236 | 8.185 | 271836 |
| JFRC2013 | ANTs A | 2.964 | 2.740 | 0.707 | 2.121 | 6.519 | 274140 |
|  | ANTs B | 3.956 | 3.560 | 0.707 | 2.915 | 8.775 | 273296 |
|  | ANTs C | 3.621 | 3.400 | 0.707 | 2.550 | 8.093 | 272964 |
|  | CMTK A | 3.401 | 3.014 | 0.707 | 2.550 | 7.211 | 251372 |
|  | CMTK B | 3.548 | 3.139 | 0.707 | 2.646 | 7.517 | 250490 |
|  | CMTK C | 4.381 | 3.392 | 1.000 | 3.536 | 9.000 | 252590 |
|  | Elastix A | 3.141 | 2.831 | 0.707 | 2.236 | 6.856 | 269952 |
|  | Elastix B | 3.077 | 2.832 | 0.707 | 2.236 | 6.708 | 274714 |
| JRC2018 | ANTs A | 2.936 | 2.696 | 0.707 | 2.121 | 6.442 | 277906 |
|  | ANTs B | 3.112 | 2.860 | 0.707 | 2.236 | 6.856 | 275250 |
|  | ANTs C | 3.129 | 2.864 | 0.707 | 2.236 | 6.856 | 270738 |
|  | CMTK A | 2.915 | 2.794 | 0.707 | 2.121 | 6.442 | 247334 |
|  | CMTK B | 2.904 | 2.782 | 0.707 | 2.121 | 6.364 | 250582 |
|  | CMTK C | 2.940 | 2.756 | 0.707 | 2.121 | 6.442 | 256194 |
|  | Elastix A | 3.018 | 2.764 | 0.707 | 2.121 | 6.557 | 270890 |
|  | Elastix B | 3.035 | 2.836 | 0.707 | 2.121 | 6.671 | 271324 |
| FCWB | ANTs A | 3.200 | 2.942 | 0.707 | 2.236 | 7.106 | 285442 |
|  | ANTs B | 3.882 | 3.600 | 0.707 | 2.646 | 8.660 | 266108 |
|  | ANTs C | 3.829 | 3.367 | 0.707 | 2.828 | 8.573 | 261238 |
|  | CMTK A | 5.664 | 5.290 | 1.000 | 3.808 | 12.923 | 244344 |
|  | CMTK B | 5.217 | 5.143 | 1.000 | 3.606 | 11.424 | 228770 |
|  | CMTK C | 4.132 | 4.121 | 0.707 | 2.646 | 9.670 | 249120 |
|  | Elastix A | 7.432 | 6.613 | 1.000 | 5.292 | 17.132 | 243274 |
|  | Elastix B | 4.629 | 4.765 | 0.707 | 2.915 | 10.998 | 273922 |
| Tefor | ANTs A | 2.932 | 2.682 | 0.707 | 2.121 | 6.364 | 274682 |
|  | ANTs B | 3.530 | 3.179 | 0.707 | 2.550 | 7.810 | 263076 |
|  | ANTs C | 3.692 | 3.349 | 0.707 | 2.646 | 8.276 | 259718 |
|  | CMTK A | 4.028 | 4.995 | 0.707 | 2.236 | 9.721 | 250548 |
|  | CMTK B | 4.129 | 5.001 | 0.707 | 2.236 | 10.392 | 254716 |
|  | CMTK C | 3.852 | 3.775 | 0.707 | 2.646 | 8.631 | 251828 |
|  | Elastix A | 3.176 | 2.807 | 0.707 | 2.236 | 6.856 | 270686 |
|  | Elastix B | 3.731 | 3.463 | 0.707 | 2.646 | 8.276 | 266274 |

**Table S46:** BU_R : bulb

| Template | Algorithm | Mean | Std dev | 10th perc | median | 90th perc | N |
| --- | --- | --- | --- | --- | --- | --- | --- |
| JFRC2010 | ANTs A | 6.841 | 6.256 | 1.581 | 5.099 | 13.379 | 22424 |
|  | ANTs B | 4.086 | 2.934 | 1.000 | 3.240 | 8.307 | 11390 |
|  | ANTs C | 4.507 | 3.029 | 1.225 | 3.808 | 8.944 | 10434 |
|  | CMTK A | 6.861 | 6.411 | 0.707 | 4.637 | 17.117 | 8222 |
|  | CMTK B | 7.242 | 7.921 | 1.000 | 4.743 | 17.641 | 7780 |
|  | CMTK C | 8.124 | 5.609 | 2.449 | 7.000 | 16.509 | 5620 |
|  | Elastix A | 4.611 | 3.085 | 1.000 | 4.062 | 9.110 | 8760 |
|  | Elastix B | 5.793 | 5.397 | 0.707 | 3.808 | 13.675 | 10622 |
| JFRC2013 | ANTs A | 3.843 | 2.835 | 1.000 | 3.000 | 7.937 | 11872 |
|  | ANTs B | 5.089 | 3.287 | 1.225 | 4.528 | 9.670 | 8524 |
|  | ANTs C | 4.723 | 3.322 | 1.000 | 3.873 | 9.618 | 9558 |
|  | CMTK A | 3.947 | 2.867 | 1.000 | 3.240 | 7.906 | 13272 |
|  | CMTK B | 4.215 | 2.919 | 1.000 | 3.606 | 8.276 | 13264 |
|  | CMTK C | 4.151 | 2.845 | 1.000 | 3.536 | 8.093 | 14454 |
|  | Elastix A | 4.034 | 2.971 | 1.000 | 3.162 | 8.544 | 11234 |
|  | Elastix B | 4.181 | 2.877 | 1.000 | 3.606 | 8.276 | 11774 |
| JRC2018 | ANTs A | 3.780 | 2.795 | 1.000 | 3.000 | 7.778 | 11756 |
|  | ANTs B | 3.733 | 2.846 | 0.707 | 2.915 | 7.906 | 11878 |
|  | ANTs C | 3.867 | 2.873 | 1.000 | 3.000 | 8.093 | 10400 |
|  | CMTK A | 3.581 | 2.687 | 0.707 | 2.915 | 7.246 | 13240 |
|  | CMTK B | 3.521 | 2.683 | 0.707 | 2.828 | 7.382 | 13728 |
|  | CMTK C | 3.545 | 2.768 | 0.707 | 2.646 | 7.649 | 13292 |
|  | Elastix A | 3.646 | 2.756 | 1.000 | 2.915 | 7.681 | 12872 |
|  | Elastix B | 3.947 | 2.887 | 0.707 | 3.240 | 8.124 | 12502 |
| FCWB | ANTs A | 3.985 | 2.916 | 1.000 | 3.162 | 8.093 | 11384 |
|  | ANTs B | 5.016 | 3.673 | 1.000 | 4.243 | 10.149 | 6992 |
|  | ANTs C | 4.931 | 3.364 | 1.000 | 4.243 | 9.849 | 7812 |
|  | CMTK A | 4.931 | 4.047 | 1.000 | 3.606 | 10.886 | 10626 |
|  | CMTK B | 5.382 | 4.323 | 1.000 | 4.183 | 11.726 | 12736 |
|  | CMTK C | 4.243 | 3.157 | 1.000 | 3.464 | 8.944 | 12936 |
|  | Elastix A | 6.866 | 5.075 | 1.225 | 5.745 | 14.089 | 9312 |
|  | Elastix B | 7.656 | 6.481 | 1.732 | 6.042 | 15.925 | 7320 |
| Tefor | ANTs A | 3.797 | 2.739 | 1.000 | 3.000 | 7.810 | 12254 |
|  | ANTs B | 4.168 | 2.814 | 1.000 | 3.536 | 8.246 | 10092 |
|  | ANTs C | 4.115 | 2.670 | 1.000 | 3.536 | 8.093 | 12988 |
|  | CMTK A | 5.519 | 5.626 | 0.707 | 4.031 | 10.886 | 12094 |
|  | CMTK B | 5.515 | 5.538 | 0.707 | 3.808 | 11.023 | 11236 |
|  | CMTK C | 4.592 | 2.858 | 1.414 | 4.062 | 8.860 | 14446 |
|  | Elastix A | 4.895 | 3.334 | 1.000 | 4.243 | 9.513 | 9732 |
|  | Elastix B | 6.338 | 5.427 | 1.000 | 4.528 | 14.560 | 8606 |

**Table S47:** VES_L : vest

| Template | Algorithm | Mean | Std dev | 10th perc | median | 90th perc | N |
| --- | --- | --- | --- | --- | --- | --- | --- |
| JFRC2010 | ANTs A | 8.553 | 6.912 | 1.581 | 6.364 | 18.722 | 127354 |
|  | ANTs B | 4.516 | 3.277 | 1.225 | 3.674 | 8.888 | 247654 |
|  | ANTs C | 4.862 | 3.569 | 1.225 | 4.062 | 9.747 | 276772 |
|  | CMTK A | 6.364 | 5.110 | 1.414 | 4.950 | 13.656 | 181760 |
|  | CMTK B | 6.464 | 5.207 | 1.414 | 4.950 | 14.160 | 161820 |
|  | CMTK C | 5.854 | 4.722 | 1.414 | 4.528 | 12.227 | 258396 |
|  | Elastix A | 4.170 | 3.497 | 1.000 | 3.240 | 8.246 | 127674 |
|  | Elastix B | 4.304 | 3.391 | 1.000 | 3.464 | 8.544 | 117868 |
| JFRC2013 | ANTs A | 4.252 | 3.412 | 1.000 | 3.464 | 8.426 | 115754 |
|  | ANTs B | 4.414 | 3.302 | 1.225 | 3.606 | 8.631 | 279706 |
|  | ANTs C | 4.619 | 3.383 | 1.225 | 3.808 | 9.192 | 254522 |
|  | CMTK A | 4.824 | 3.874 | 1.000 | 3.808 | 9.849 | 104886 |
|  | CMTK B | 4.860 | 3.763 | 1.225 | 3.808 | 9.849 | 109968 |
|  | CMTK C | 4.822 | 3.441 | 1.414 | 4.062 | 9.434 | 194618 |
|  | Elastix A | 4.441 | 3.515 | 1.000 | 3.606 | 8.631 | 107718 |
|  | Elastix B | 4.440 | 3.540 | 1.000 | 3.606 | 8.775 | 106336 |
| JRC2018 | ANTs A | 4.268 | 3.460 | 1.000 | 3.464 | 8.544 | 114502 |
|  | ANTs B | 4.623 | 3.490 | 1.225 | 3.808 | 9.083 | 141690 |
|  | ANTs C | 4.590 | 3.383 | 1.225 | 3.808 | 9.192 | 145422 |
|  | CMTK A | 4.504 | 3.553 | 1.000 | 3.606 | 9.000 | 106820 |
|  | CMTK B | 4.472 | 3.555 | 1.000 | 3.606 | 8.944 | 101196 |
|  | CMTK C | 4.553 | 3.671 | 1.000 | 3.606 | 8.972 | 103616 |
|  | Elastix A | 4.296 | 3.409 | 1.000 | 3.536 | 8.337 | 113174 |
|  | Elastix B | 4.257 | 3.389 | 1.000 | 3.464 | 8.396 | 112718 |
| FCWB | ANTs A | 4.399 | 3.629 | 1.000 | 3.464 | 8.860 | 116374 |
|  | ANTs B | 4.546 | 3.278 | 1.414 | 3.808 | 8.631 | 230142 |
|  | ANTs C | 4.721 | 3.336 | 1.414 | 4.062 | 8.972 | 212550 |
|  | CMTK A | 7.129 | 5.518 | 1.581 | 5.701 | 15.067 | 201758 |
|  | CMTK B | 7.042 | 5.441 | 1.581 | 5.568 | 14.866 | 151708 |
|  | CMTK C | 6.447 | 5.039 | 1.581 | 5.000 | 13.675 | 164182 |
|  | Elastix A | 6.541 | 5.142 | 1.581 | 5.148 | 13.472 | 263744 |
|  | Elastix B | 7.034 | 5.573 | 1.581 | 5.523 | 14.731 | 178076 |
| Tefor | ANTs A | 4.269 | 3.440 | 1.000 | 3.464 | 8.485 | 120644 |
|  | ANTs B | 4.489 | 3.298 | 1.225 | 3.674 | 8.803 | 210210 |
|  | ANTs C | 5.060 | 3.693 | 1.414 | 4.123 | 10.075 | 177482 |
|  | CMTK A | 7.087 | 5.766 | 1.414 | 5.339 | 15.906 | 165070 |
|  | CMTK B | 7.404 | 6.048 | 1.581 | 5.523 | 16.643 | 161920 |
|  | CMTK C | 6.849 | 5.351 | 1.581 | 5.292 | 14.731 | 164558 |
|  | Elastix A | 4.036 | 3.284 | 1.000 | 3.240 | 7.906 | 161986 |
|  | Elastix B | 4.821 | 3.675 | 1.225 | 3.873 | 9.618 | 185564 |

**Table S48:** IB_L : inferior bridge

| Template | Algorithm | Mean | Std dev | 10th perc | median | 90th perc | N |
| --- | --- | --- | --- | --- | --- | --- | --- |
| JFRC2010 | ANTs A | 4.882 | 4.210 | 1.000 | 3.606 | 10.583 | 232736 |
|  | ANTs B | 4.080 | 3.540 | 0.707 | 3.000 | 8.718 | 256620 |
|  | ANTs C | 4.103 | 3.172 | 1.000 | 3.240 | 8.544 | 390228 |
|  | CMTK A | 4.728 | 4.691 | 0.707 | 3.162 | 10.583 | 242754 |
|  | CMTK B | 4.884 | 5.420 | 0.707 | 3.082 | 11.269 | 254096 |
|  | CMTK C | 4.620 | 4.982 | 1.000 | 3.000 | 10.512 | 338184 |
|  | Elastix A | 2.999 | 2.482 | 0.707 | 2.345 | 5.874 | 255814 |
|  | Elastix B | 3.499 | 2.962 | 0.707 | 2.646 | 7.280 | 213680 |
| JFRC2013 | ANTs A | 3.054 | 2.715 | 0.707 | 2.236 | 6.442 | 223036 |
|  | ANTs B | 4.110 | 3.312 | 1.000 | 3.240 | 8.396 | 358766 |
|  | ANTs C | 4.159 | 3.331 | 1.000 | 3.240 | 8.944 | 370482 |
|  | CMTK A | 3.197 | 2.739 | 0.707 | 2.550 | 6.442 | 190126 |
|  | CMTK B | 3.244 | 2.740 | 0.707 | 2.550 | 6.557 | 188414 |
|  | CMTK C | 4.745 | 3.770 | 1.000 | 3.606 | 10.440 | 230108 |
|  | Elastix A | 2.953 | 2.545 | 0.707 | 2.236 | 5.831 | 229984 |
|  | Elastix B | 3.015 | 2.666 | 0.707 | 2.236 | 6.083 | 204902 |
| JRC2018 | ANTs A | 2.930 | 2.662 | 0.707 | 2.236 | 6.042 | 223680 |
|  | ANTs B | 3.316 | 3.020 | 0.707 | 2.345 | 7.211 | 224980 |
|  | ANTs C | 3.240 | 2.907 | 0.707 | 2.345 | 7.036 | 228840 |
|  | CMTK A | 3.193 | 2.959 | 0.707 | 2.345 | 6.708 | 195902 |
|  | CMTK B | 3.107 | 2.839 | 0.707 | 2.236 | 6.557 | 190138 |
|  | CMTK C | 3.035 | 2.702 | 0.707 | 2.236 | 6.364 | 197050 |
|  | Elastix A | 3.059 | 2.679 | 0.707 | 2.345 | 6.205 | 206600 |
|  | Elastix B | 2.988 | 2.646 | 0.707 | 2.236 | 6.083 | 201614 |
| FCWB | ANTs A | 3.394 | 3.016 | 0.707 | 2.550 | 7.280 | 226180 |
|  | ANTs B | 4.588 | 3.574 | 1.000 | 3.606 | 9.618 | 298536 |
|  | ANTs C | 3.784 | 2.918 | 1.000 | 3.000 | 7.746 | 366938 |
|  | CMTK A | 6.488 | 5.469 | 1.414 | 4.796 | 14.107 | 190292 |
|  | CMTK B | 5.715 | 5.134 | 1.000 | 4.062 | 12.767 | 195356 |
|  | CMTK C | 4.265 | 3.892 | 0.707 | 3.000 | 9.539 | 250136 |
|  | Elastix A | 7.710 | 7.839 | 1.225 | 5.050 | 18.083 | 267584 |
|  | Elastix B | 4.410 | 4.285 | 0.707 | 3.000 | 9.618 | 245972 |
| Tefor | ANTs A | 3.053 | 2.744 | 0.707 | 2.236 | 6.364 | 224680 |
|  | ANTs B | 3.920 | 3.501 | 0.707 | 2.915 | 8.544 | 228986 |
|  | ANTs C | 3.954 | 3.505 | 0.707 | 2.915 | 8.860 | 252914 |
|  | CMTK A | 4.384 | 3.963 | 0.707 | 3.000 | 10.296 | 206084 |
|  | CMTK B | 4.337 | 3.946 | 0.707 | 3.000 | 10.124 | 208340 |
|  | CMTK C | 4.332 | 3.818 | 0.707 | 3.162 | 9.539 | 203452 |
|  | Elastix A | 3.112 | 2.640 | 0.707 | 2.449 | 6.205 | 228276 |
|  | Elastix B | 3.587 | 3.086 | 0.707 | 2.646 | 7.616 | 206388 |

**Table S49:** ATL_L : antler

| Template | Algorithm | Mean | Std dev | 10th perc | median | 90th perc | N |
| --- | --- | --- | --- | --- | --- | --- | --- |
| JFRC2010 | ANTs A | 3.926 | 2.876 | 1.000 | 3.240 | 7.550 | 81610 |
|  | ANTs B | 3.925 | 2.699 | 1.000 | 3.317 | 7.583 | 80346 |
|  | ANTs C | 4.335 | 2.831 | 1.225 | 3.808 | 8.276 | 87234 |
|  | CMTK A | 5.173 | 5.092 | 1.000 | 3.808 | 9.849 | 76872 |
|  | CMTK B | 4.921 | 4.715 | 1.000 | 3.606 | 9.434 | 77040 |
|  | CMTK C | 4.630 | 3.994 | 1.000 | 3.674 | 8.367 | 81250 |
|  | Elastix A | 3.316 | 2.154 | 1.000 | 2.915 | 6.124 | 87064 |
|  | Elastix B | 3.583 | 2.412 | 1.000 | 3.000 | 7.000 | 91986 |
| JFRC2013 | ANTs A | 3.112 | 2.133 | 1.000 | 2.646 | 5.831 | 79242 |
|  | ANTs B | 4.120 | 2.866 | 1.000 | 3.536 | 8.062 | 96460 |
|  | ANTs C | 4.268 | 3.111 | 1.000 | 3.536 | 8.972 | 84404 |
|  | CMTK A | 3.225 | 2.191 | 1.000 | 2.828 | 6.083 | 74162 |
|  | CMTK B | 3.450 | 2.314 | 1.000 | 3.000 | 6.519 | 70046 |
|  | CMTK C | 4.372 | 2.934 | 1.225 | 3.808 | 8.337 | 67292 |
|  | Elastix A | 3.155 | 2.036 | 1.000 | 2.828 | 5.701 | 77368 |
|  | Elastix B | 3.127 | 2.053 | 1.000 | 2.646 | 5.788 | 81338 |
| JRC2018 | ANTs A | 3.006 | 2.071 | 0.707 | 2.550 | 5.701 | 80720 |
|  | ANTs B | 3.131 | 2.137 | 1.000 | 2.646 | 5.788 | 74448 |
|  | ANTs C | 3.297 | 2.190 | 1.000 | 2.915 | 6.083 | 74372 |
|  | CMTK A | 3.208 | 2.179 | 1.000 | 2.828 | 6.083 | 79102 |
|  | CMTK B | 3.173 | 2.136 | 1.000 | 2.646 | 6.042 | 78520 |
|  | CMTK C | 3.158 | 2.128 | 1.000 | 2.646 | 5.874 | 77660 |
|  | Elastix A | 3.248 | 2.125 | 1.000 | 2.915 | 6.042 | 76480 |
|  | Elastix B | 3.083 | 2.079 | 1.000 | 2.646 | 5.831 | 83538 |
| FCWB | ANTs A | 3.393 | 2.406 | 1.000 | 2.915 | 6.245 | 77720 |
|  | ANTs B | 4.198 | 2.897 | 1.225 | 3.536 | 8.515 | 85288 |
|  | ANTs C | 4.050 | 2.615 | 1.225 | 3.536 | 7.649 | 95052 |
|  | CMTK A | 5.627 | 5.160 | 1.414 | 4.123 | 11.597 | 63522 |
|  | CMTK B | 5.410 | 5.394 | 1.225 | 3.674 | 11.790 | 63758 |
|  | CMTK C | 3.874 | 2.625 | 1.000 | 3.317 | 7.211 | 66826 |
|  | Elastix A | 6.525 | 7.232 | 1.414 | 4.528 | 11.790 | 83412 |
|  | Elastix B | 4.357 | 3.178 | 1.000 | 3.606 | 8.544 | 89336 |
| Tefor | ANTs A | 3.012 | 2.067 | 0.707 | 2.550 | 5.701 | 80992 |
|  | ANTs B | 3.660 | 2.400 | 1.000 | 3.162 | 6.708 | 77408 |
|  | ANTs C | 3.975 | 2.649 | 1.000 | 3.536 | 7.550 | 72348 |
|  | CMTK A | 3.683 | 2.532 | 1.000 | 3.082 | 7.211 | 90296 |
|  | CMTK B | 3.724 | 2.548 | 1.000 | 3.162 | 7.314 | 92468 |
|  | CMTK C | 3.980 | 2.613 | 1.000 | 3.536 | 7.382 | 81148 |
|  | Elastix A | 3.209 | 2.125 | 1.000 | 2.828 | 6.042 | 91852 |
|  | Elastix B | 3.361 | 2.214 | 1.000 | 2.915 | 6.442 | 99390 |

**Table S50:** CRE_L : crepine

| Template | Algorithm | Mean | Std dev | 10th perc | median | 90th perc | N |
| --- | --- | --- | --- | --- | --- | --- | --- |
| JFRC2010 | ANTs A | 7.843 | 6.117 | 1.414 | 6.364 | 16.310 | 239816 |
|  | ANTs B | 4.882 | 4.247 | 1.000 | 3.606 | 10.536 | 169636 |
|  | ANTs C | 5.226 | 4.288 | 1.225 | 4.062 | 10.817 | 145186 |
|  | CMTK A | 7.478 | 6.081 | 1.581 | 5.745 | 16.093 | 112646 |
|  | CMTK B | 6.826 | 5.505 | 1.414 | 5.339 | 14.440 | 115154 |
|  | CMTK C | 6.944 | 5.331 | 1.732 | 5.568 | 13.675 | 145274 |
|  | Elastix A | 5.228 | 4.305 | 1.225 | 4.062 | 10.863 | 138834 |
|  | Elastix B | 7.382 | 6.197 | 1.414 | 5.339 | 16.523 | 114000 |
| JFRC2013 | ANTs A | 4.344 | 3.985 | 1.000 | 3.082 | 9.301 | 156584 |
|  | ANTs B | 6.904 | 5.199 | 1.581 | 5.523 | 14.248 | 130784 |
|  | ANTs C | 5.645 | 4.975 | 1.225 | 4.123 | 12.268 | 149384 |
|  | CMTK A | 5.374 | 4.354 | 1.225 | 4.123 | 11.247 | 139624 |
|  | CMTK B | 5.432 | 4.392 | 1.225 | 4.183 | 11.358 | 142824 |
|  | CMTK C | 5.417 | 4.644 | 1.225 | 4.123 | 11.180 | 178982 |
|  | Elastix A | 5.023 | 4.069 | 1.225 | 3.873 | 10.124 | 141276 |
|  | Elastix B | 5.548 | 4.683 | 1.225 | 4.183 | 11.597 | 129230 |
| JRC2018 | ANTs A | 4.425 | 3.963 | 1.000 | 3.162 | 9.487 | 150742 |
|  | ANTs B | 4.932 | 4.289 | 1.000 | 3.606 | 10.700 | 145352 |
|  | ANTs C | 5.012 | 4.227 | 1.000 | 3.808 | 10.583 | 144126 |
|  | CMTK A | 4.675 | 4.011 | 1.000 | 3.536 | 9.849 | 128454 |
|  | CMTK B | 4.576 | 3.993 | 1.000 | 3.464 | 9.644 | 127400 |
|  | CMTK C | 4.661 | 3.978 | 1.000 | 3.536 | 9.706 | 135654 |
|  | Elastix A | 4.838 | 3.958 | 1.000 | 3.808 | 9.899 | 130144 |
|  | Elastix B | 5.844 | 5.171 | 1.000 | 4.183 | 13.077 | 121474 |
| FCWB | ANTs A | 5.445 | 5.399 | 1.000 | 3.536 | 12.806 | 154238 |
|  | ANTs B | 8.573 | 5.625 | 2.236 | 7.416 | 16.538 | 96860 |
|  | ANTs C | 7.817 | 5.851 | 1.732 | 6.519 | 15.588 | 124264 |
|  | CMTK A | 6.758 | 5.417 | 1.581 | 5.099 | 14.832 | 115928 |
|  | CMTK B | 6.119 | 5.018 | 1.414 | 4.528 | 13.533 | 124330 |
|  | CMTK C | 6.238 | 5.030 | 1.414 | 4.637 | 13.675 | 142558 |
|  | Elastix A | 8.272 | 5.892 | 1.732 | 7.036 | 16.447 | 117624 |
|  | Elastix B | 9.648 | 7.001 | 2.121 | 7.937 | 20.100 | 94772 |
| Tefor | ANTs A | 4.256 | 3.872 | 1.000 | 3.000 | 9.192 | 153390 |
|  | ANTs B | 4.722 | 4.137 | 1.000 | 3.464 | 10.223 | 161208 |
|  | ANTs C | 4.981 | 4.023 | 1.000 | 3.808 | 10.607 | 169534 |
|  | CMTK A | 7.305 | 5.380 | 1.581 | 6.000 | 15.379 | 129854 |
|  | CMTK B | 6.964 | 5.274 | 1.581 | 5.568 | 14.782 | 122228 |
|  | CMTK C | 5.181 | 4.484 | 1.000 | 3.808 | 11.336 | 209880 |
|  | Elastix A | 5.339 | 4.350 | 1.225 | 4.062 | 11.380 | 121224 |
|  | Elastix B | 9.298 | 7.235 | 1.732 | 7.314 | 19.925 | 97360 |

**Table S51:** MB_PED_L : pedunculus of adult mushroom body

| Template | Algorithm | Mean | Std dev | 10th perc | median | 90th perc | N |
| --- | --- | --- | --- | --- | --- | --- | --- |
| JFRC2010 | ANTs A | 7.145 | 8.007 | 1.581 | 4.743 | 13.964 | 91380 |
|  | ANTs B | 6.229 | 4.282 | 1.581 | 5.292 | 12.410 | 60494 |
|  | ANTs C | 6.416 | 4.304 | 1.581 | 5.523 | 12.590 | 58210 |
|  | CMTK A | 6.655 | 5.238 | 1.414 | 5.050 | 14.387 | 60120 |
|  | CMTK B | 6.406 | 4.971 | 1.581 | 5.000 | 13.583 | 81492 |
|  | CMTK C | 6.463 | 4.580 | 1.581 | 5.339 | 13.285 | 74938 |
|  | Elastix A | 5.762 | 4.296 | 1.581 | 4.528 | 12.137 | 62808 |
|  | Elastix B | 6.287 | 4.869 | 1.581 | 4.950 | 13.229 | 61956 |
| JFRC2013 | ANTs A | 5.118 | 3.964 | 1.225 | 3.873 | 10.817 | 67314 |
|  | ANTs B | 6.594 | 4.532 | 1.732 | 5.568 | 12.806 | 54366 |
|  | ANTs C | 5.766 | 4.191 | 1.581 | 4.743 | 11.489 | 54350 |
|  | CMTK A | 5.550 | 4.136 | 1.414 | 4.359 | 11.511 | 68540 |
|  | CMTK B | 5.647 | 4.233 | 1.414 | 4.528 | 11.597 | 69752 |
|  | CMTK C | 5.630 | 4.180 | 1.414 | 4.583 | 11.424 | 58542 |
|  | Elastix A | 5.424 | 4.127 | 1.414 | 4.243 | 11.511 | 64554 |
|  | Elastix B | 5.225 | 4.026 | 1.414 | 4.062 | 11.091 | 65702 |
| JRC2018 | ANTs A | 5.159 | 3.957 | 1.225 | 4.062 | 10.724 | 69302 |
|  | ANTs B | 5.290 | 4.014 | 1.225 | 4.183 | 10.909 | 68308 |
|  | ANTs C | 5.245 | 3.992 | 1.414 | 4.123 | 10.817 | 66266 |
|  | CMTK A | 5.158 | 4.013 | 1.225 | 3.873 | 10.886 | 67844 |
|  | CMTK B | 5.155 | 3.989 | 1.225 | 3.873 | 10.863 | 67512 |
|  | CMTK C | 5.207 | 4.011 | 1.414 | 4.062 | 10.977 | 67636 |
|  | Elastix A | 5.180 | 3.918 | 1.414 | 4.062 | 10.817 | 69618 |
|  | Elastix B | 5.303 | 4.103 | 1.414 | 4.123 | 11.358 | 64394 |
| FCWB | ANTs A | 5.690 | 4.372 | 1.414 | 4.528 | 11.769 | 56720 |
|  | ANTs B | 5.220 | 4.126 | 1.000 | 4.062 | 11.068 | 55698 |
|  | ANTs C | 5.146 | 3.926 | 1.000 | 4.123 | 10.464 | 59852 |
|  | CMTK A | 6.471 | 4.991 | 1.414 | 5.099 | 13.472 | 51692 |
|  | CMTK B | 6.402 | 4.772 | 1.581 | 5.196 | 13.151 | 52692 |
|  | CMTK C | 5.146 | 4.140 | 1.000 | 3.873 | 11.068 | 63016 |
|  | Elastix A | 6.927 | 4.825 | 1.581 | 5.874 | 13.802 | 62632 |
|  | Elastix B | 6.015 | 5.928 | 1.000 | 3.873 | 13.638 | 57326 |
| Tefor | ANTs A | 5.088 | 3.903 | 1.225 | 4.000 | 10.607 | 71160 |
|  | ANTs B | 5.438 | 3.996 | 1.414 | 4.359 | 11.068 | 62306 |
|  | ANTs C | 5.736 | 4.044 | 1.581 | 4.743 | 11.380 | 67984 |
|  | CMTK A | 6.099 | 5.023 | 1.414 | 4.472 | 13.416 | 67056 |
|  | CMTK B | 6.055 | 4.971 | 1.414 | 4.301 | 13.472 | 64964 |
|  | CMTK C | 5.667 | 4.403 | 1.414 | 4.359 | 12.042 | 67684 |
|  | Elastix A | 5.611 | 4.253 | 1.414 | 4.359 | 11.726 | 63094 |
|  | Elastix B | 6.480 | 5.079 | 1.581 | 4.950 | 13.892 | 57146 |

**Table S52:** MB_VL_L : vertical lobe of adult mushroom body

| Template | Algorithm | Mean | Std dev | 10th perc | median | 90th perc | N |
| --- | --- | --- | --- | --- | --- | --- | --- |
| JFRC2010 | ANTs A | 5.895 | 5.046 | 1.581 | 4.528 | 11.597 | 135494 |
|  | ANTs B | 5.647 | 4.232 | 1.581 | 4.583 | 10.794 | 93942 |
|  | ANTs C | 6.059 | 4.522 | 1.581 | 5.000 | 11.619 | 98208 |
|  | CMTK A | 5.182 | 3.897 | 1.581 | 4.301 | 9.618 | 124950 |
|  | CMTK B | 5.238 | 4.127 | 1.581 | 4.243 | 9.849 | 121466 |
|  | CMTK C | 5.962 | 4.319 | 1.732 | 5.050 | 10.977 | 88886 |
|  | Elastix A | 4.942 | 4.069 | 1.414 | 3.873 | 9.513 | 122750 |
|  | Elastix B | 5.055 | 4.310 | 1.414 | 3.873 | 10.025 | 131996 |
| JFRC2013 | ANTs A | 5.296 | 4.152 | 1.581 | 4.359 | 9.721 | 111872 |
|  | ANTs B | 6.160 | 4.326 | 1.732 | 5.196 | 11.554 | 93068 |
|  | ANTs C | 5.942 | 4.297 | 1.581 | 5.000 | 10.977 | 101550 |
|  | CMTK A | 5.691 | 4.508 | 1.581 | 4.528 | 11.068 | 115624 |
|  | CMTK B | 5.785 | 4.455 | 1.581 | 4.743 | 10.863 | 108634 |
|  | CMTK C | 6.098 | 4.247 | 1.732 | 5.196 | 11.203 | 95824 |
|  | Elastix A | 4.934 | 3.981 | 1.414 | 3.873 | 9.487 | 130894 |
|  | Elastix B | 4.890 | 4.075 | 1.225 | 3.808 | 9.644 | 133500 |
| JRC2018 | ANTs A | 4.678 | 4.021 | 1.225 | 3.606 | 9.000 | 131582 |
|  | ANTs B | 5.279 | 4.156 | 1.581 | 4.243 | 10.000 | 115318 |
|  | ANTs C | 5.384 | 4.236 | 1.581 | 4.359 | 10.464 | 122964 |
|  | CMTK A | 4.724 | 4.148 | 1.225 | 3.606 | 9.247 | 128888 |
|  | CMTK B | 4.944 | 4.139 | 1.414 | 3.808 | 9.434 | 126064 |
|  | CMTK C | 5.323 | 4.164 | 1.581 | 4.301 | 9.950 | 119020 |
|  | Elastix A | 4.905 | 4.078 | 1.414 | 3.808 | 9.460 | 123552 |
|  | Elastix B | 4.965 | 4.286 | 1.414 | 3.808 | 9.849 | 123732 |
| FCWB | ANTs A | 5.525 | 4.317 | 1.581 | 4.472 | 10.724 | 111644 |
|  | ANTs B | 5.282 | 3.922 | 1.581 | 4.359 | 10.124 | 79092 |
|  | ANTs C | 5.348 | 3.989 | 1.581 | 4.472 | 9.950 | 89654 |
|  | CMTK A | 5.570 | 4.269 | 1.581 | 4.528 | 10.724 | 102496 |
|  | CMTK B | 5.517 | 4.203 | 1.581 | 4.472 | 10.654 | 97158 |
|  | CMTK C | 5.266 | 3.814 | 1.581 | 4.359 | 9.899 | 80306 |
|  | Elastix A | 5.428 | 4.186 | 1.581 | 4.472 | 10.149 | 100274 |
|  | Elastix B | 5.563 | 4.439 | 1.581 | 4.359 | 10.909 | 112672 |
| Tefor | ANTs A | 4.681 | 3.993 | 1.225 | 3.606 | 9.055 | 133834 |
|  | ANTs B | 5.442 | 4.207 | 1.581 | 4.359 | 10.223 | 98620 |
|  | ANTs C | 5.421 | 4.099 | 1.581 | 4.472 | 10.392 | 106004 |
|  | CMTK A | 6.814 | 6.928 | 1.414 | 4.359 | 17.044 | 138644 |
|  | CMTK B | 6.370 | 5.807 | 1.581 | 4.528 | 14.765 | 129842 |
|  | CMTK C | 6.186 | 4.424 | 1.732 | 5.148 | 11.853 | 98746 |
|  | Elastix A | 4.993 | 4.136 | 1.414 | 4.000 | 9.539 | 125584 |
|  | Elastix B | 5.336 | 4.609 | 1.414 | 4.062 | 10.724 | 122576 |

**Table S53:** MB_ML_L : medial lobe of adult mushroom body

| Template | Algorithm | Mean | Std dev | 10th perc | median | 90th perc | N |
| --- | --- | --- | --- | --- | --- | --- | --- |
| JFRC2010 | ANTs A | 6.261 | 6.063 | 1.225 | 4.183 | 14.036 | 219070 |
|  | ANTs B | 4.228 | 3.847 | 1.000 | 3.162 | 8.631 | 246608 |
|  | ANTs C | 4.538 | 3.955 | 1.000 | 3.464 | 9.327 | 226850 |
|  | CMTK A | 6.603 | 6.355 | 1.414 | 4.472 | 15.556 | 201488 |
|  | CMTK B | 5.414 | 4.843 | 1.000 | 3.873 | 11.769 | 209146 |
|  | CMTK C | 5.735 | 5.508 | 1.225 | 4.000 | 12.410 | 235838 |
|  | Elastix A | 4.264 | 3.803 | 1.000 | 3.240 | 8.544 | 252902 |
|  | Elastix B | 5.639 | 5.303 | 1.000 | 3.873 | 12.826 | 221790 |
| JFRC2013 | ANTs A | 4.092 | 3.702 | 1.000 | 3.000 | 8.276 | 229810 |
|  | ANTs B | 5.867 | 4.824 | 1.581 | 4.528 | 12.021 | 161752 |
|  | ANTs C | 5.250 | 4.780 | 1.000 | 3.808 | 10.794 | 168914 |
|  | CMTK A | 5.086 | 4.257 | 1.225 | 3.808 | 10.630 | 207446 |
|  | CMTK B | 5.164 | 4.277 | 1.225 | 3.873 | 10.677 | 206382 |
|  | CMTK C | 4.820 | 4.394 | 1.000 | 3.536 | 10.100 | 182974 |
|  | Elastix A | 4.470 | 3.881 | 1.000 | 3.536 | 8.860 | 219734 |
|  | Elastix B | 4.781 | 4.117 | 1.000 | 3.606 | 9.721 | 209946 |
| JRC2018 | ANTs A | 4.045 | 3.695 | 1.000 | 3.000 | 8.093 | 239818 |
|  | ANTs B | 4.313 | 3.797 | 1.000 | 3.240 | 8.746 | 224826 |
|  | ANTs C | 4.363 | 3.819 | 1.000 | 3.317 | 8.573 | 224532 |
|  | CMTK A | 4.353 | 3.779 | 1.000 | 3.317 | 8.573 | 223708 |
|  | CMTK B | 4.317 | 3.745 | 1.000 | 3.317 | 8.515 | 219828 |
|  | CMTK C | 4.439 | 3.793 | 1.000 | 3.464 | 8.775 | 217188 |
|  | Elastix A | 4.363 | 3.737 | 1.000 | 3.464 | 8.544 | 230346 |
|  | Elastix B | 5.155 | 4.515 | 1.225 | 3.808 | 10.724 | 211664 |
| FCWB | ANTs A | 4.683 | 4.373 | 1.000 | 3.317 | 10.025 | 194636 |
|  | ANTs B | 5.355 | 4.389 | 1.414 | 4.123 | 11.158 | 155400 |
|  | ANTs C | 5.271 | 4.507 | 1.225 | 3.873 | 10.977 | 184730 |
|  | CMTK A | 5.598 | 4.750 | 1.414 | 4.183 | 11.790 | 193610 |
|  | CMTK B | 5.410 | 4.572 | 1.414 | 4.062 | 11.489 | 192134 |
|  | CMTK C | 5.022 | 4.183 | 1.225 | 3.808 | 10.607 | 178880 |
|  | Elastix A | 6.116 | 4.738 | 1.581 | 4.950 | 12.288 | 183072 |
|  | Elastix B | 7.718 | 7.008 | 1.581 | 5.196 | 18.628 | 162264 |
| Tefor | ANTs A | 4.103 | 3.678 | 1.000 | 3.082 | 8.185 | 238708 |
|  | ANTs B | 4.272 | 3.742 | 1.000 | 3.240 | 8.746 | 229654 |
|  | ANTs C | 4.688 | 4.236 | 1.000 | 3.536 | 9.618 | 222946 |
|  | CMTK A | 5.672 | 4.650 | 1.414 | 4.359 | 11.937 | 218430 |
|  | CMTK B | 5.610 | 4.568 | 1.414 | 4.359 | 11.597 | 218734 |
|  | CMTK C | 5.211 | 4.507 | 1.225 | 3.808 | 11.180 | 226658 |
|  | Elastix A | 4.530 | 3.883 | 1.000 | 3.536 | 9.220 | 245986 |
|  | Elastix B | 8.394 | 7.716 | 1.581 | 5.431 | 20.445 | 150204 |

**Table S54:** FLA_L : flange

| Template | Algorithm | Mean | Std dev | 10th perc | median | 90th perc | N |
| --- | --- | --- | --- | --- | --- | --- | --- |
| JFRC2010 | ANTs A | 5.633 | 6.448 | 0.707 | 3.240 | 15.636 | 336696 |
|  | ANTs B | 3.763 | 3.134 | 0.707 | 3.000 | 7.314 | 351018 |
|  | ANTs C | 3.552 | 3.075 | 0.707 | 2.915 | 7.071 | 401822 |
|  | CMTK A | 3.703 | 3.276 | 0.707 | 2.915 | 7.280 | 369328 |
|  | CMTK B | 3.687 | 3.450 | 0.707 | 2.915 | 7.106 | 373918 |
|  | CMTK C | 4.291 | 3.846 | 1.000 | 3.240 | 8.544 | 336458 |
|  | Elastix A | 2.995 | 2.373 | 0.707 | 2.550 | 5.745 | 388882 |
|  | Elastix B | 2.908 | 2.325 | 0.707 | 2.345 | 5.701 | 398758 |
| JFRC2013 | ANTs A | 2.870 | 2.450 | 0.707 | 2.236 | 5.701 | 392302 |
|  | ANTs B | 4.250 | 3.381 | 1.000 | 3.536 | 8.000 | 277230 |
|  | ANTs C | 3.783 | 3.161 | 1.000 | 3.000 | 7.211 | 295372 |
|  | CMTK A | 2.938 | 2.383 | 0.707 | 2.345 | 5.788 | 420952 |
|  | CMTK B | 2.952 | 2.406 | 0.707 | 2.345 | 5.788 | 412536 |
|  | CMTK C | 3.427 | 2.583 | 1.000 | 2.915 | 6.557 | 335392 |
|  | Elastix A | 2.970 | 2.436 | 0.707 | 2.449 | 5.745 | 375750 |
|  | Elastix B | 2.922 | 2.414 | 0.707 | 2.345 | 5.745 | 397916 |
| JRC2018 | ANTs A | 2.814 | 2.398 | 0.707 | 2.236 | 5.568 | 397460 |
|  | ANTs B | 3.253 | 2.697 | 0.707 | 2.646 | 6.364 | 381326 |
|  | ANTs C | 3.358 | 2.782 | 0.707 | 2.646 | 6.557 | 385464 |
|  | CMTK A | 2.841 | 2.378 | 0.707 | 2.236 | 5.568 | 441218 |
|  | CMTK B | 2.820 | 2.371 | 0.707 | 2.236 | 5.568 | 421384 |
|  | CMTK C | 2.869 | 2.410 | 0.707 | 2.345 | 5.701 | 403114 |
|  | Elastix A | 2.966 | 2.398 | 0.707 | 2.449 | 5.745 | 391286 |
|  | Elastix B | 2.990 | 2.569 | 0.707 | 2.345 | 5.831 | 386058 |
| FCWB | ANTs A | 2.879 | 2.536 | 0.707 | 2.236 | 5.701 | 408720 |
|  | ANTs B | 3.688 | 2.858 | 1.000 | 3.000 | 7.071 | 325596 |
|  | ANTs C | 3.547 | 3.059 | 0.707 | 2.915 | 6.928 | 361668 |
|  | CMTK A | 3.415 | 2.967 | 0.707 | 2.915 | 6.557 | 379116 |
|  | CMTK B | 3.280 | 3.081 | 0.707 | 2.646 | 6.245 | 397938 |
|  | CMTK C | 3.345 | 2.687 | 0.707 | 2.646 | 6.519 | 341304 |
|  | Elastix A | 4.480 | 4.266 | 1.000 | 3.808 | 8.307 | 305690 |
|  | Elastix B | 3.419 | 2.633 | 0.707 | 2.915 | 6.671 | 373924 |
| Tefor | ANTs A | 2.856 | 2.475 | 0.707 | 2.236 | 5.701 | 403824 |
|  | ANTs B | 3.583 | 2.955 | 0.707 | 2.915 | 6.856 | 346276 |
|  | ANTs C | 3.522 | 2.991 | 0.707 | 2.915 | 6.708 | 395204 |
|  | CMTK A | 3.302 | 2.756 | 0.707 | 2.646 | 6.671 | 431620 |
|  | CMTK B | 3.429 | 2.851 | 0.707 | 2.828 | 6.856 | 402012 |
|  | CMTK C | 3.736 | 3.057 | 1.000 | 3.000 | 7.106 | 375272 |
|  | Elastix A | 3.240 | 2.631 | 0.707 | 2.646 | 6.124 | 356548 |
|  | Elastix B | 3.346 | 2.919 | 0.707 | 2.828 | 6.403 | 327064 |

**Table S55:** LOP_L : lobula plate

| Template | Algorithm | Mean | Std dev | 10th perc | median | 90th perc | N |
| --- | --- | --- | --- | --- | --- | --- | --- |
| JFRC2010 | ANTs A | 8.631 | 6.875 | 2.236 | 6.856 | 16.778 | 362688 |
|  | ANTs B | 6.772 | 5.477 | 2.000 | 5.523 | 12.590 | 338210 |
|  | ANTs C | 7.036 | 5.551 | 2.000 | 5.701 | 13.285 | 401672 |
|  | CMTK A | 8.762 | 6.656 | 2.236 | 6.964 | 17.958 | 409008 |
|  | CMTK B | 7.881 | 5.869 | 2.121 | 6.364 | 15.906 | 383972 |
|  | CMTK C | 7.383 | 5.528 | 2.121 | 5.916 | 14.526 | 433212 |
|  | Elastix A | 5.820 | 5.269 | 1.581 | 4.359 | 11.597 | 335456 |
|  | Elastix B | 5.722 | 5.012 | 1.581 | 4.359 | 11.269 | 352972 |
| JFRC2013 | ANTs A | 5.921 | 5.503 | 1.581 | 4.528 | 11.619 | 398430 |
|  | ANTs B | 6.585 | 5.388 | 1.732 | 5.196 | 12.845 | 388774 |
|  | ANTs C | 6.848 | 5.833 | 1.581 | 5.292 | 13.711 | 393766 |
|  | CMTK A | 7.142 | 7.562 | 1.581 | 5.000 | 14.387 | 424038 |
|  | CMTK B | 7.345 | 7.940 | 1.581 | 5.099 | 14.213 | 422190 |
|  | CMTK C | 6.931 | 5.242 | 2.000 | 5.701 | 13.285 | 434874 |
|  | Elastix A | 5.742 | 5.123 | 1.581 | 4.359 | 11.358 | 427686 |
|  | Elastix B | 5.650 | 5.276 | 1.414 | 4.183 | 11.402 | 438740 |
| JRC2018 | ANTs A | 5.663 | 5.194 | 1.414 | 4.243 | 11.336 | 405342 |
|  | ANTs B | 5.857 | 5.177 | 1.581 | 4.528 | 11.402 | 389376 |
|  | ANTs C | 6.158 | 5.304 | 1.581 | 4.743 | 12.166 | 399826 |
|  | CMTK A | 5.602 | 5.111 | 1.414 | 4.243 | 11.068 | 426888 |
|  | CMTK B | 5.636 | 5.213 | 1.414 | 4.243 | 11.180 | 414846 |
|  | CMTK C | 5.794 | 5.173 | 1.581 | 4.472 | 11.336 | 401586 |
|  | Elastix A | 5.619 | 5.051 | 1.414 | 4.301 | 11.091 | 403720 |
|  | Elastix B | 5.529 | 5.317 | 1.414 | 4.123 | 10.724 | 436688 |
| FCWB | ANTs A | 6.237 | 5.717 | 1.581 | 4.690 | 12.268 | 398870 |
|  | ANTs B | 6.452 | 5.341 | 1.732 | 5.000 | 12.669 | 336656 |
|  | ANTs C | 6.778 | 5.600 | 1.732 | 5.339 | 13.058 | 359718 |
|  | CMTK A | 8.018 | 6.591 | 2.121 | 6.364 | 15.668 | 388196 |
|  | CMTK B | 7.788 | 6.416 | 2.121 | 6.083 | 15.395 | 365136 |
|  | CMTK C | 8.460 | 6.896 | 2.121 | 6.442 | 17.479 | 344492 |
|  | Elastix A | 9.498 | 8.798 | 2.345 | 7.416 | 17.903 | 384198 |
|  | Elastix B | 8.059 | 9.535 | 1.732 | 5.568 | 15.199 | 405130 |
| Tefor | ANTs A | 5.687 | 5.248 | 1.414 | 4.301 | 11.358 | 403146 |
|  | ANTs B | 6.159 | 5.413 | 1.581 | 4.796 | 11.874 | 354400 |
|  | ANTs C | 6.979 | 5.497 | 2.000 | 5.701 | 13.379 | 360948 |
|  | CMTK A | 7.749 | 6.901 | 1.581 | 5.568 | 16.643 | 400468 |
|  | CMTK B | 7.790 | 6.803 | 1.581 | 5.701 | 16.523 | 388860 |
|  | CMTK C | 7.053 | 5.804 | 1.732 | 5.523 | 14.053 | 371694 |
|  | Elastix A | 5.595 | 5.035 | 1.414 | 4.183 | 11.180 | 360282 |
|  | Elastix B | 5.576 | 5.172 | 1.414 | 4.183 | 11.068 | 382420 |

**Table S56:** PB : protocerebral bridge

| Template | Algorithm | Mean | Std dev | 10th perc | median | 90th perc | N |
| --- | --- | --- | --- | --- | --- | --- | --- |
| JFRC2010 | ANTs A | 5.083 | 4.229 | 1.000 | 3.808 | 10.977 | 272138 |
|  | ANTs B | 4.445 | 3.253 | 1.000 | 3.606 | 9.083 | 193424 |
|  | ANTs C | 4.761 | 3.300 | 1.000 | 4.062 | 9.539 | 229120 |
|  | CMTK A | 5.421 | 4.942 | 1.000 | 3.674 | 12.767 | 215634 |
|  | CMTK B | 4.547 | 4.221 | 1.000 | 3.240 | 9.874 | 236674 |
|  | CMTK C | 4.402 | 3.624 | 1.000 | 3.240 | 9.592 | 220518 |
|  | Elastix A | 3.227 | 2.491 | 0.707 | 2.646 | 6.481 | 226828 |
|  | Elastix B | 3.266 | 2.638 | 0.707 | 2.550 | 6.671 | 249262 |
| JFRC2013 | ANTs A | 3.837 | 3.072 | 0.707 | 3.000 | 8.185 | 219702 |
|  | ANTs B | 4.390 | 3.105 | 1.000 | 3.674 | 8.746 | 226656 |
|  | ANTs C | 4.608 | 3.175 | 1.000 | 3.873 | 9.110 | 228424 |
|  | CMTK A | 3.381 | 2.529 | 0.707 | 2.646 | 6.964 | 180538 |
|  | CMTK B | 3.580 | 2.653 | 1.000 | 2.915 | 7.246 | 166454 |
|  | CMTK C | 4.730 | 3.301 | 1.000 | 4.062 | 9.460 | 136042 |
|  | Elastix A | 3.323 | 2.621 | 0.707 | 2.646 | 6.819 | 220190 |
|  | Elastix B | 3.246 | 2.504 | 0.707 | 2.646 | 6.557 | 232012 |
| JRC2018 | ANTs A | 3.377 | 2.643 | 0.707 | 2.646 | 6.856 | 233940 |
|  | ANTs B | 3.677 | 2.885 | 0.707 | 2.915 | 7.416 | 234506 |
|  | ANTs C | 3.980 | 3.241 | 1.000 | 3.082 | 8.155 | 247940 |
|  | CMTK A | 3.578 | 2.779 | 0.707 | 2.915 | 7.416 | 231020 |
|  | CMTK B | 3.521 | 2.729 | 0.707 | 2.915 | 7.246 | 229874 |
|  | CMTK C | 3.531 | 2.737 | 0.707 | 2.915 | 7.246 | 228440 |
|  | Elastix A | 3.542 | 2.666 | 0.707 | 2.915 | 7.106 | 216916 |
|  | Elastix B | 3.319 | 2.578 | 0.707 | 2.646 | 6.708 | 235262 |
| FCWB | ANTs A | 4.420 | 3.391 | 1.000 | 3.536 | 9.220 | 189324 |
|  | ANTs B | 5.077 | 3.406 | 1.225 | 4.359 | 9.849 | 186316 |
|  | ANTs C | 5.080 | 3.429 | 1.225 | 4.359 | 9.874 | 205584 |
|  | CMTK A | 5.807 | 5.115 | 1.225 | 4.528 | 11.358 | 135822 |
|  | CMTK B | 5.475 | 5.004 | 1.000 | 4.243 | 10.724 | 167728 |
|  | CMTK C | 4.986 | 3.759 | 1.000 | 4.062 | 10.536 | 147642 |
|  | Elastix A | 6.782 | 6.050 | 1.581 | 5.568 | 11.874 | 162078 |
|  | Elastix B | 4.127 | 3.456 | 1.000 | 3.082 | 8.888 | 193698 |
| Tefor | ANTs A | 3.341 | 2.612 | 0.707 | 2.646 | 6.708 | 228940 |
|  | ANTs B | 4.523 | 3.262 | 1.000 | 3.808 | 9.165 | 176804 |
|  | ANTs C | 5.212 | 3.790 | 1.225 | 4.301 | 10.583 | 195598 |
|  | CMTK A | 4.028 | 3.049 | 1.000 | 3.240 | 8.426 | 215812 |
|  | CMTK B | 4.164 | 3.211 | 1.000 | 3.317 | 8.746 | 207496 |
|  | CMTK C | 4.652 | 3.359 | 1.000 | 3.808 | 9.301 | 169744 |
|  | Elastix A | 3.223 | 2.494 | 0.707 | 2.646 | 6.481 | 221016 |
|  | Elastix B | 3.029 | 2.442 | 0.707 | 2.345 | 6.325 | 256792 |

**Table S57:** AL_L : adult antennal lobe

| Template | Algorithm | Mean | Std dev | 10th perc | median | 90th perc | N |
| --- | --- | --- | --- | --- | --- | --- | --- |
| JFRC2010 | ANTs A | 5.077 | 5.521 | 1.225 | 3.536 | 10.320 | 1320934 |
|  | ANTs B | 3.626 | 2.491 | 1.225 | 3.162 | 6.364 | 1378504 |
|  | ANTs C | 3.687 | 2.552 | 1.225 | 3.240 | 6.442 | 1538688 |
|  | CMTK A | 4.100 | 3.519 | 1.225 | 3.240 | 7.280 | 1392898 |
|  | CMTK B | 3.798 | 2.808 | 1.225 | 3.240 | 6.708 | 1415636 |
|  | CMTK C | 4.126 | 3.226 | 1.225 | 3.464 | 7.550 | 1351974 |
|  | Elastix A | 3.282 | 2.046 | 1.000 | 3.000 | 5.745 | 1433024 |
|  | Elastix B | 3.365 | 2.045 | 1.000 | 3.000 | 5.916 | 1500192 |
| JFRC2013 | ANTs A | 3.265 | 1.967 | 1.000 | 3.000 | 5.745 | 1545570 |
|  | ANTs B | 4.382 | 3.871 | 1.225 | 3.317 | 8.396 | 1084472 |
|  | ANTs C | 4.133 | 3.487 | 1.225 | 3.240 | 7.649 | 1202770 |
|  | CMTK A | 3.513 | 2.322 | 1.000 | 3.082 | 6.205 | 1462514 |
|  | CMTK B | 3.561 | 2.364 | 1.000 | 3.082 | 6.364 | 1412544 |
|  | CMTK C | 3.782 | 2.740 | 1.225 | 3.162 | 6.856 | 1126876 |
|  | Elastix A | 3.266 | 1.984 | 1.000 | 3.000 | 5.745 | 1508478 |
|  | Elastix B | 3.241 | 1.938 | 1.000 | 2.915 | 5.701 | 1567854 |
| JRC2018 | ANTs A | 3.177 | 1.884 | 1.000 | 2.915 | 5.568 | 1575740 |
|  | ANTs B | 3.551 | 2.333 | 1.000 | 3.162 | 6.205 | 1454164 |
|  | ANTs C | 3.697 | 2.457 | 1.225 | 3.240 | 6.557 | 1474432 |
|  | CMTK A | 3.191 | 1.878 | 1.000 | 2.915 | 5.568 | 1604044 |
|  | CMTK B | 3.201 | 1.867 | 1.000 | 2.915 | 5.568 | 1605078 |
|  | CMTK C | 3.283 | 1.956 | 1.000 | 3.000 | 5.745 | 1577672 |
|  | Elastix A | 3.278 | 1.957 | 1.000 | 3.000 | 5.745 | 1571160 |
|  | Elastix B | 3.258 | 1.923 | 1.000 | 3.000 | 5.701 | 1577384 |
| FCWB | ANTs A | 3.610 | 2.495 | 1.000 | 3.082 | 6.403 | 1499740 |
|  | ANTs B | 4.067 | 3.255 | 1.225 | 3.240 | 7.517 | 1022826 |
|  | ANTs C | 4.352 | 3.775 | 1.225 | 3.317 | 8.307 | 1150918 |
|  | CMTK A | 3.710 | 2.592 | 1.225 | 3.240 | 6.557 | 1546656 |
|  | CMTK B | 3.731 | 2.659 | 1.225 | 3.240 | 6.557 | 1535038 |
|  | CMTK C | 3.763 | 2.616 | 1.225 | 3.240 | 6.671 | 1414018 |
|  | Elastix A | 3.745 | 2.440 | 1.225 | 3.317 | 6.633 | 1449056 |
|  | Elastix B | 3.669 | 2.483 | 1.000 | 3.162 | 6.557 | 1464562 |
| Tefor | ANTs A | 3.219 | 1.905 | 1.000 | 2.915 | 5.657 | 1573310 |
|  | ANTs B | 3.530 | 2.354 | 1.000 | 3.082 | 6.164 | 1479424 |
|  | ANTs C | 3.860 | 3.124 | 1.225 | 3.240 | 6.708 | 1554168 |
|  | CMTK A | 3.786 | 2.626 | 1.000 | 3.240 | 6.856 | 1546958 |
|  | CMTK B | 3.754 | 2.596 | 1.000 | 3.240 | 6.708 | 1583232 |
|  | CMTK C | 4.052 | 3.224 | 1.225 | 3.240 | 7.416 | 1560302 |
|  | Elastix A | 3.358 | 2.037 | 1.000 | 3.000 | 5.874 | 1516002 |
|  | Elastix B | 3.752 | 2.534 | 1.225 | 3.240 | 6.819 | 1508102 |

**Table S58:** ME_L : medulla

| Template | Algorithm | Mean | Std dev | 10th perc | median | 90th perc | N |
| --- | --- | --- | --- | --- | --- | --- | --- |
| JFRC2010 | ANTs A | 6.214 | 5.692 | 1.581 | 4.950 | 11.726 | 3208676 |
|  | ANTs B | 5.101 | 4.599 | 1.581 | 4.301 | 8.860 | 2887600 |
|  | ANTs C | 5.413 | 4.467 | 1.581 | 4.583 | 9.513 | 3165268 |
|  | CMTK A | 6.169 | 5.528 | 1.581 | 4.950 | 11.424 | 3100936 |
|  | CMTK B | 5.857 | 5.420 | 1.581 | 4.690 | 10.607 | 3023396 |
|  | CMTK C | 6.269 | 5.209 | 1.732 | 5.148 | 11.203 | 2840876 |
|  | Elastix A | 4.853 | 4.440 | 1.581 | 4.123 | 8.396 | 3003908 |
|  | Elastix B | 4.947 | 4.383 | 1.581 | 4.243 | 8.602 | 3015102 |
| JFRC2013 | ANTs A | 4.898 | 4.452 | 1.581 | 4.123 | 8.544 | 3117410 |
|  | ANTs B | 5.354 | 4.713 | 1.581 | 4.528 | 9.327 | 2892702 |
|  | ANTs C | 5.629 | 4.817 | 1.581 | 4.743 | 9.950 | 3018516 |
|  | CMTK A | 6.786 | 7.513 | 1.732 | 5.000 | 11.853 | 2609038 |
|  | CMTK B | 7.041 | 8.260 | 1.732 | 5.050 | 11.937 | 2576206 |
|  | CMTK C | 6.090 | 4.471 | 2.121 | 5.339 | 10.464 | 2663474 |
|  | Elastix A | 4.892 | 4.286 | 1.581 | 4.183 | 8.544 | 3069924 |
|  | Elastix B | 4.922 | 4.402 | 1.581 | 4.183 | 8.631 | 3050764 |
| JRC2018 | ANTs A | 4.778 | 4.232 | 1.581 | 4.062 | 8.337 | 3135938 |
|  | ANTs B | 5.037 | 4.500 | 1.581 | 4.243 | 8.775 | 3102894 |
|  | ANTs C | 5.550 | 4.647 | 1.581 | 4.583 | 10.000 | 3145260 |
|  | CMTK A | 4.826 | 4.367 | 1.581 | 4.123 | 8.337 | 3073602 |
|  | CMTK B | 4.821 | 4.426 | 1.581 | 4.123 | 8.307 | 3085844 |
|  | CMTK C | 4.912 | 4.370 | 1.581 | 4.183 | 8.515 | 3136598 |
|  | Elastix A | 4.898 | 4.860 | 1.581 | 4.123 | 8.337 | 3036246 |
|  | Elastix B | 4.960 | 4.595 | 1.581 | 4.183 | 8.544 | 3024912 |
| FCWB | ANTs A | 5.014 | 4.453 | 1.581 | 4.243 | 8.775 | 3183810 |
|  | ANTs B | 5.481 | 4.718 | 1.581 | 4.690 | 9.513 | 3063022 |
|  | ANTs C | 6.039 | 4.628 | 2.000 | 5.196 | 10.700 | 3139250 |
|  | CMTK A | 6.419 | 5.405 | 1.732 | 5.148 | 12.247 | 3092300 |
|  | CMTK B | 6.099 | 5.175 | 1.732 | 5.000 | 11.247 | 3118278 |
|  | CMTK C | 6.466 | 5.424 | 2.000 | 5.339 | 11.511 | 2966210 |
|  | Elastix A | 7.160 | 7.213 | 2.121 | 5.745 | 13.172 | 3312420 |
|  | Elastix B | 6.300 | 6.954 | 1.732 | 5.000 | 11.424 | 3294236 |
| Tefor | ANTs A | 4.747 | 4.246 | 1.414 | 4.062 | 8.307 | 3107898 |
|  | ANTs B | 5.063 | 4.669 | 1.581 | 4.183 | 8.860 | 2912902 |
|  | ANTs C | 5.747 | 5.048 | 1.581 | 4.743 | 10.124 | 3115598 |
|  | CMTK A | 6.855 | 7.161 | 1.581 | 4.743 | 14.248 | 2831230 |
|  | CMTK B | 6.860 | 7.124 | 1.581 | 4.743 | 14.300 | 2839816 |
|  | CMTK C | 6.573 | 7.246 | 1.581 | 4.743 | 11.874 | 2736262 |
|  | Elastix A | 4.830 | 4.531 | 1.581 | 4.123 | 8.367 | 2928176 |
|  | Elastix B | 4.871 | 4.529 | 1.581 | 4.123 | 8.515 | 2906158 |

**Table S59:** SLP_L : superior lateral protocerebrum

| Template | Algorithm | Mean | Std dev | 10th perc | median | 90th perc | N |
| --- | --- | --- | --- | --- | --- | --- | --- |
| JFRC2010 | ANTs A | 5.477 | 6.140 | 0.707 | 3.317 | 13.229 | 606516 |
|  | ANTs B | 4.670 | 4.987 | 0.707 | 2.915 | 11.068 | 542156 |
|  | ANTs C | 4.853 | 5.318 | 0.707 | 2.915 | 11.619 | 587638 |
|  | CMTK A | 5.890 | 6.810 | 0.707 | 3.162 | 15.395 | 642956 |
|  | CMTK B | 6.572 | 7.608 | 0.707 | 3.606 | 17.521 | 698112 |
|  | CMTK C | 6.180 | 6.991 | 0.707 | 3.240 | 16.263 | 566916 |
|  | Elastix A | 3.834 | 4.291 | 0.707 | 2.236 | 9.110 | 492408 |
|  | Elastix B | 4.177 | 4.764 | 0.707 | 2.236 | 10.512 | 510852 |
| JFRC2013 | ANTs A | 4.129 | 4.721 | 0.707 | 2.345 | 10.075 | 580284 |
|  | ANTs B | 5.679 | 5.772 | 0.707 | 3.808 | 13.416 | 500620 |
|  | ANTs C | 5.339 | 5.576 | 0.707 | 3.464 | 12.767 | 558630 |
|  | CMTK A | 5.027 | 4.664 | 1.000 | 3.606 | 10.909 | 493046 |
|  | CMTK B | 5.031 | 4.580 | 1.000 | 3.674 | 10.817 | 514176 |
|  | CMTK C | 5.827 | 5.227 | 1.000 | 4.243 | 12.649 | 486968 |
|  | Elastix A | 3.814 | 4.247 | 0.707 | 2.236 | 9.055 | 545258 |
|  | Elastix B | 3.862 | 4.464 | 0.707 | 2.236 | 9.083 | 539860 |
| JRC2018 | ANTs A | 3.767 | 4.286 | 0.707 | 2.121 | 9.083 | 568444 |
|  | ANTs B | 4.289 | 5.007 | 0.707 | 2.236 | 10.817 | 565658 |
|  | ANTs C | 4.532 | 5.175 | 0.707 | 2.646 | 11.000 | 601494 |
|  | CMTK A | 3.818 | 4.186 | 0.707 | 2.236 | 9.055 | 573080 |
|  | CMTK B | 3.792 | 4.268 | 0.707 | 2.236 | 9.083 | 560844 |
|  | CMTK C | 4.090 | 4.713 | 0.707 | 2.236 | 9.950 | 575064 |
|  | Elastix A | 3.712 | 4.114 | 0.707 | 2.236 | 8.803 | 533126 |
|  | Elastix B | 3.859 | 4.255 | 0.707 | 2.236 | 9.247 | 543210 |
| FCWB | ANTs A | 5.032 | 5.245 | 0.707 | 3.464 | 11.068 | 560066 |
|  | ANTs B | 5.343 | 5.448 | 0.707 | 3.606 | 12.390 | 493738 |
|  | ANTs C | 5.289 | 5.464 | 0.707 | 3.536 | 12.227 | 574654 |
|  | CMTK A | 5.321 | 5.640 | 0.707 | 3.536 | 12.450 | 535526 |
|  | CMTK B | 5.213 | 5.470 | 0.707 | 3.536 | 12.021 | 544466 |
|  | CMTK C | 4.920 | 5.019 | 0.707 | 3.240 | 11.446 | 526058 |
|  | Elastix A | 5.324 | 5.923 | 0.707 | 3.162 | 13.285 | 490822 |
|  | Elastix B | 4.630 | 5.219 | 0.707 | 2.550 | 11.769 | 492544 |
| Tefor | ANTs A | 3.820 | 4.392 | 0.707 | 2.121 | 9.220 | 579900 |
|  | ANTs B | 4.547 | 4.824 | 0.707 | 2.915 | 10.770 | 584556 |
|  | ANTs C | 4.987 | 5.100 | 0.707 | 3.317 | 11.424 | 631596 |
|  | CMTK A | 5.635 | 7.627 | 0.707 | 2.449 | 15.906 | 586094 |
|  | CMTK B | 5.655 | 7.379 | 0.707 | 2.646 | 15.182 | 581586 |
|  | CMTK C | 5.305 | 5.935 | 0.707 | 3.162 | 13.058 | 631288 |
|  | Elastix A | 3.724 | 4.277 | 0.707 | 2.121 | 8.972 | 539944 |
|  | Elastix B | 4.110 | 4.671 | 0.707 | 2.236 | 10.247 | 546230 |

**Table S60:**
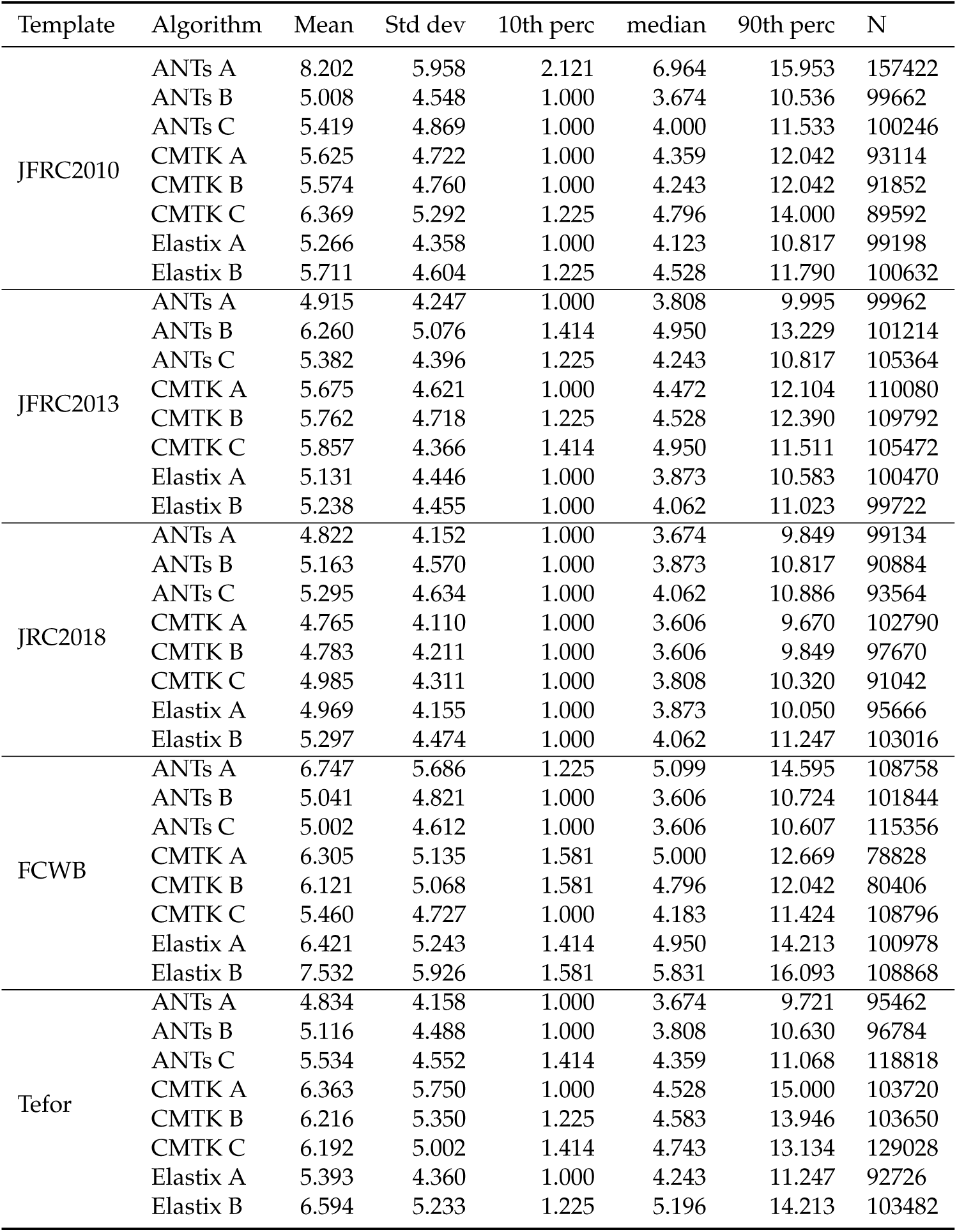
SIP_L : superior intermediate protocerebrum

**Table S61:** SMP_L : superior medial protocerebrum

| Template | Algorithm | Mean | Std dev | 10th perc | median | 90th perc | N |
| --- | --- | --- | --- | --- | --- | --- | --- |
| JFRC2010 | ANTs A | 4.487 | 4.759 | 0.707 | 2.915 | 10.223 | 2048810 |
|  | ANTs B | 2.871 | 2.689 | 0.707 | 2.121 | 6.083 | 1844162 |
|  | ANTs C | 2.884 | 2.818 | 0.707 | 2.121 | 6.124 | 1905962 |
|  | CMTK A | 3.977 | 4.464 | 0.707 | 2.550 | 8.803 | 1538652 |
|  | CMTK B | 3.601 | 3.655 | 0.707 | 2.449 | 8.031 | 1604384 |
|  | CMTK C | 4.044 | 4.257 | 0.707 | 2.646 | 9.220 | 1457268 |
|  | Elastix A | 2.834 | 2.623 | 0.707 | 2.121 | 5.874 | 1698384 |
|  | Elastix B | 3.509 | 3.241 | 0.707 | 2.550 | 7.583 | 1530020 |
| JFRC2013 | ANTs A | 2.554 | 2.523 | 0.707 | 1.732 | 5.431 | 1978286 |
|  | ANTs B | 2.862 | 2.798 | 0.707 | 2.121 | 6.083 | 1878714 |
|  | ANTs C | 2.794 | 2.794 | 0.707 | 2.000 | 6.000 | 1961782 |
|  | CMTK A | 2.860 | 2.840 | 0.707 | 2.121 | 6.083 | 1808334 |
|  | CMTK B | 2.973 | 2.895 | 0.707 | 2.121 | 6.403 | 1785804 |
|  | CMTK C | 3.238 | 2.861 | 0.707 | 2.449 | 6.856 | 1874888 |
|  | Elastix A | 2.726 | 2.593 | 0.707 | 2.121 | 5.568 | 1745092 |
|  | Elastix B | 2.828 | 2.758 | 0.707 | 2.121 | 5.874 | 1788762 |
| JRC2018 | ANTs A | 2.564 | 2.568 | 0.707 | 1.732 | 5.431 | 1956318 |
|  | ANTs B | 2.667 | 2.614 | 0.707 | 2.000 | 5.657 | 1819568 |
|  | ANTs C | 2.638 | 2.641 | 0.707 | 2.000 | 5.568 | 1914416 |
|  | CMTK A | 2.546 | 2.461 | 0.707 | 2.000 | 5.196 | 1955892 |
|  | CMTK B | 2.603 | 2.492 | 0.707 | 2.000 | 5.339 | 1883916 |
|  | CMTK C | 2.674 | 2.576 | 0.707 | 2.000 | 5.568 | 1853804 |
|  | Elastix A | 2.734 | 2.564 | 0.707 | 2.121 | 5.701 | 1798790 |
|  | Elastix B | 2.826 | 2.699 | 0.707 | 2.121 | 5.916 | 1893642 |
| FCWB | ANTs A | 2.785 | 2.838 | 0.707 | 2.000 | 6.000 | 1993800 |
|  | ANTs B | 3.344 | 2.972 | 0.707 | 2.550 | 6.964 | 1546890 |
|  | ANTs C | 3.265 | 2.969 | 0.707 | 2.449 | 6.856 | 1817190 |
|  | CMTK A | 3.279 | 3.135 | 0.707 | 2.345 | 7.000 | 1760952 |
|  | CMTK B | 3.161 | 3.019 | 0.707 | 2.236 | 6.671 | 1789962 |
|  | CMTK C | 3.238 | 2.987 | 0.707 | 2.345 | 7.000 | 1673578 |
|  | Elastix A | 4.141 | 4.058 | 0.707 | 2.915 | 9.301 | 1638518 |
|  | Elastix B | 3.809 | 3.587 | 0.707 | 2.646 | 8.396 | 1683704 |
| Tefor | ANTs A | 2.536 | 2.537 | 0.707 | 1.732 | 5.339 | 1953734 |
|  | ANTs B | 2.700 | 2.526 | 0.707 | 2.121 | 5.701 | 1875044 |
|  | ANTs C | 2.757 | 2.620 | 0.707 | 2.121 | 5.831 | 1996512 |
|  | CMTK A | 3.524 | 3.769 | 0.707 | 2.236 | 7.906 | 1744916 |
|  | CMTK B | 3.497 | 3.821 | 0.707 | 2.236 | 7.810 | 1736936 |
|  | CMTK C | 3.086 | 2.971 | 0.707 | 2.236 | 6.557 | 1797932 |
|  | Elastix A | 2.843 | 2.693 | 0.707 | 2.121 | 5.916 | 1789736 |
|  | Elastix B | 3.619 | 3.496 | 0.707 | 2.646 | 7.906 | 1671190 |

**Table S62:** AVLP_L : anterior ventrolateral protocerebrum

| Template | Algorithm | Mean | Std dev | 10th perc | median | 90th perc | N |
| --- | --- | --- | --- | --- | --- | --- | --- |
| JFRC2010 | ANTs A | 7.819 | 6.241 | 1.732 | 5.874 | 17.393 | 698640 |
|  | ANTs B | 5.936 | 4.431 | 1.581 | 5.000 | 11.511 | 648902 |
|  | ANTs C | 6.316 | 4.712 | 1.581 | 5.196 | 12.410 | 734882 |
|  | CMTK A | 6.250 | 4.859 | 1.581 | 5.000 | 12.669 | 858926 |
|  | CMTK B | 6.124 | 4.779 | 1.581 | 4.950 | 12.369 | 847098 |
|  | CMTK C | 6.577 | 4.771 | 1.732 | 5.523 | 12.903 | 798272 |
|  | Elastix A | 5.114 | 3.961 | 1.414 | 4.243 | 9.618 | 690276 |
|  | Elastix B | 5.902 | 4.761 | 1.414 | 4.743 | 11.790 | 745100 |
| JFRC2013 | ANTs A | 4.991 | 3.977 | 1.225 | 4.123 | 9.487 | 712030 |
|  | ANTs B | 6.906 | 4.970 | 1.732 | 5.745 | 13.583 | 560778 |
|  | ANTs C | 6.425 | 4.972 | 1.581 | 5.148 | 12.826 | 630090 |
|  | CMTK A | 5.867 | 4.372 | 1.581 | 4.950 | 11.424 | 667434 |
|  | CMTK B | 5.962 | 4.403 | 1.581 | 5.000 | 11.597 | 647920 |
|  | CMTK C | 6.650 | 4.636 | 1.732 | 5.568 | 12.903 | 526864 |
|  | Elastix A | 5.084 | 3.939 | 1.414 | 4.183 | 9.618 | 712258 |
|  | Elastix B | 5.025 | 3.841 | 1.414 | 4.183 | 9.426 | 734544 |
| JRC2018 | ANTs A | 4.947 | 3.929 | 1.225 | 4.123 | 9.327 | 718986 |
|  | ANTs B | 5.335 | 4.181 | 1.414 | 4.359 | 10.392 | 698242 |
|  | ANTs C | 5.566 | 4.273 | 1.581 | 4.528 | 10.724 | 736770 |
|  | CMTK A | 4.951 | 3.885 | 1.225 | 4.123 | 9.247 | 713194 |
|  | CMTK B | 4.979 | 3.921 | 1.414 | 4.123 | 9.301 | 698056 |
|  | CMTK C | 5.165 | 4.006 | 1.414 | 4.301 | 9.721 | 688070 |
|  | Elastix A | 5.034 | 3.953 | 1.414 | 4.123 | 9.487 | 695914 |
|  | Elastix B | 5.117 | 3.971 | 1.414 | 4.243 | 9.670 | 700946 |
| FCWB | ANTs A | 5.632 | 4.534 | 1.414 | 4.528 | 11.000 | 675074 |
|  | ANTs B | 6.545 | 4.684 | 1.732 | 5.523 | 12.826 | 548688 |
|  | ANTs C | 6.572 | 4.857 | 1.581 | 5.431 | 13.153 | 595164 |
|  | CMTK A | 6.046 | 4.456 | 1.581 | 5.000 | 11.705 | 656072 |
|  | CMTK B | 5.884 | 4.447 | 1.581 | 4.950 | 11.380 | 650172 |
|  | CMTK C | 6.348 | 4.550 | 1.581 | 5.339 | 12.268 | 568660 |
|  | Elastix A | 6.853 | 4.846 | 1.732 | 5.788 | 13.153 | 675556 |
|  | Elastix B | 6.550 | 5.625 | 1.581 | 5.000 | 13.675 | 685246 |
| Tefor | ANTs A | 4.977 | 3.939 | 1.225 | 4.123 | 9.407 | 732900 |
|  | ANTs B | 5.710 | 4.351 | 1.581 | 4.690 | 11.158 | 671192 |
|  | ANTs C | 6.294 | 4.676 | 1.581 | 5.196 | 12.369 | 718464 |
|  | CMTK A | 6.738 | 7.063 | 1.414 | 4.637 | 14.509 | 742970 |
|  | CMTK B | 6.745 | 6.995 | 1.414 | 4.637 | 14.646 | 728246 |
|  | CMTK C | 6.572 | 5.974 | 1.581 | 5.000 | 13.323 | 669718 |
|  | Elastix A | 5.089 | 3.930 | 1.414 | 4.243 | 9.539 | 714624 |
|  | Elastix B | 6.053 | 4.883 | 1.414 | 4.743 | 12.590 | 781204 |

**Table S63:** PVLP_L : posterior ventrolateral protocerebrum

| Template | Algorithm | Mean | Std dev | 10th perc | median | 90th perc | N |
| --- | --- | --- | --- | --- | --- | --- | --- |
| JFRC2010 | ANTs A | 9.112 | 6.664 | 2.000 | 7.416 | 19.118 | 187494 |
|  | ANTs B | 6.464 | 4.852 | 1.581 | 5.292 | 12.826 | 147296 |
|  | ANTs C | 6.652 | 5.096 | 1.581 | 5.523 | 13.000 | 152126 |
|  | CMTK A | 9.757 | 7.606 | 1.732 | 7.583 | 21.794 | 202130 |
|  | CMTK B | 9.154 | 7.118 | 1.732 | 7.141 | 19.925 | 197878 |
|  | CMTK C | 9.658 | 7.309 | 2.121 | 7.517 | 21.071 | 231152 |
|  | Elastix A | 5.831 | 4.685 | 1.414 | 4.528 | 11.874 | 158130 |
|  | Elastix B | 6.175 | 5.039 | 1.414 | 4.743 | 12.767 | 153506 |
| JFRC2013 | ANTs A | 5.290 | 4.309 | 1.225 | 4.123 | 11.180 | 160822 |
|  | ANTs B | 6.855 | 5.046 | 1.581 | 5.745 | 13.509 | 146820 |
|  | ANTs C | 6.684 | 5.161 | 1.581 | 5.385 | 13.583 | 147114 |
|  | CMTK A | 6.048 | 4.702 | 1.581 | 4.743 | 12.530 | 170840 |
|  | CMTK B | 6.044 | 4.733 | 1.581 | 4.743 | 12.550 | 167216 |
|  | CMTK C | 6.499 | 4.795 | 1.581 | 5.339 | 13.019 | 145374 |
|  | Elastix A | 5.297 | 4.297 | 1.225 | 4.123 | 11.136 | 160938 |
|  | Elastix B | 5.363 | 4.330 | 1.225 | 4.183 | 11.180 | 159022 |
| JRC2018 | ANTs A | 5.322 | 4.354 | 1.225 | 4.123 | 11.180 | 160018 |
|  | ANTs B | 5.638 | 4.674 | 1.225 | 4.359 | 11.874 | 154582 |
|  | ANTs C | 5.617 | 4.473 | 1.225 | 4.359 | 11.597 | 157128 |
|  | CMTK A | 5.481 | 4.422 | 1.225 | 4.183 | 11.597 | 151870 |
|  | CMTK B | 5.455 | 4.455 | 1.225 | 4.183 | 11.597 | 152622 |
|  | CMTK C | 5.399 | 4.402 | 1.225 | 4.183 | 11.269 | 155220 |
|  | Elastix A | 5.330 | 4.344 | 1.225 | 4.123 | 11.068 | 159874 |
|  | Elastix B | 5.490 | 4.411 | 1.225 | 4.243 | 11.489 | 153876 |
| FCWB | ANTs A | 6.041 | 5.003 | 1.225 | 4.528 | 13.077 | 157390 |
|  | ANTs B | 7.107 | 5.366 | 1.732 | 5.745 | 14.387 | 156264 |
|  | ANTs C | 7.099 | 5.517 | 1.581 | 5.701 | 14.283 | 154378 |
|  | CMTK A | 6.887 | 5.178 | 1.581 | 5.745 | 13.323 | 152632 |
|  | CMTK B | 6.696 | 5.210 | 1.581 | 5.523 | 13.058 | 151664 |
|  | CMTK C | 6.804 | 5.184 | 1.581 | 5.431 | 13.894 | 152840 |
|  | Elastix A | 8.175 | 5.608 | 2.121 | 7.000 | 15.764 | 177056 |
|  | Elastix B | 7.828 | 7.120 | 1.581 | 5.701 | 17.088 | 152194 |
| Tefor | ANTs A | 5.352 | 4.317 | 1.225 | 4.183 | 11.203 | 158444 |
|  | ANTs B | 6.163 | 4.734 | 1.581 | 5.000 | 12.288 | 146406 |
|  | ANTs C | 6.597 | 5.075 | 1.581 | 5.385 | 13.000 | 156388 |
|  | CMTK A | 7.418 | 6.978 | 1.414 | 5.050 | 17.219 | 148962 |
|  | CMTK B | 7.391 | 6.916 | 1.414 | 5.050 | 17.176 | 148222 |
|  | CMTK C | 7.187 | 6.039 | 1.581 | 5.292 | 16.016 | 157190 |
|  | Elastix A | 5.668 | 4.473 | 1.414 | 4.528 | 11.576 | 153402 |
|  | Elastix B | 6.148 | 4.801 | 1.581 | 4.899 | 12.530 | 150086 |

**Table S64:** IVLP_L : wedge

| Template | Algorithm | Mean | Std dev | 10th perc | median | 90th perc | N |
| --- | --- | --- | --- | --- | --- | --- | --- |
| JFRC2010 | ANTs A | 6.175 | 4.565 | 1.732 | 5.148 | 11.937 | 195518 |
|  | ANTs B | 5.508 | 3.674 | 1.732 | 4.796 | 9.644 | 186276 |
|  | ANTs C | 5.561 | 3.738 | 1.732 | 4.796 | 9.899 | 193810 |
|  | CMTK A | 5.668 | 3.873 | 1.581 | 4.950 | 10.223 | 191244 |
|  | CMTK B | 5.393 | 3.709 | 1.581 | 4.743 | 9.327 | 195038 |
|  | CMTK C | 5.692 | 3.859 | 1.732 | 5.000 | 10.025 | 204724 |
|  | Elastix A | 5.105 | 3.601 | 1.581 | 4.359 | 9.000 | 160224 |
|  | Elastix B | 5.062 | 3.395 | 1.581 | 4.472 | 8.972 | 178084 |
| JFRC2013 | ANTs A | 5.067 | 3.535 | 1.581 | 4.472 | 8.972 | 170302 |
|  | ANTs B | 5.353 | 3.430 | 1.732 | 4.743 | 9.274 | 201282 |
|  | ANTs C | 5.182 | 3.440 | 1.581 | 4.528 | 9.083 | 205812 |
|  | CMTK A | 5.150 | 3.632 | 1.581 | 4.472 | 9.220 | 173202 |
|  | CMTK B | 5.095 | 3.597 | 1.581 | 4.359 | 9.110 | 175770 |
|  | CMTK C | 5.395 | 3.656 | 1.732 | 4.690 | 9.487 | 196884 |
|  | Elastix A | 5.217 | 3.580 | 1.581 | 4.528 | 9.301 | 159198 |
|  | Elastix B | 5.129 | 3.495 | 1.581 | 4.528 | 9.083 | 161826 |
| JRC2018 | ANTs A | 5.046 | 3.502 | 1.581 | 4.359 | 8.860 | 168292 |
|  | ANTs B | 5.154 | 3.506 | 1.581 | 4.528 | 9.110 | 176482 |
|  | ANTs C | 5.174 | 3.407 | 1.581 | 4.528 | 8.972 | 189348 |
|  | CMTK A | 5.116 | 3.483 | 1.581 | 4.528 | 8.972 | 164240 |
|  | CMTK B | 5.130 | 3.505 | 1.581 | 4.528 | 9.055 | 161266 |
|  | CMTK C | 5.164 | 3.544 | 1.581 | 4.528 | 9.192 | 164222 |
|  | Elastix A | 5.095 | 3.511 | 1.581 | 4.472 | 9.000 | 157964 |
|  | Elastix B | 5.048 | 3.458 | 1.581 | 4.359 | 9.000 | 168066 |
| FCWB | ANTs A | 5.395 | 3.504 | 1.732 | 4.743 | 9.644 | 175478 |
|  | ANTs B | 5.711 | 3.834 | 1.732 | 5.000 | 10.223 | 172314 |
|  | ANTs C | 5.590 | 3.655 | 1.732 | 4.950 | 9.950 | 178622 |
|  | CMTK A | 5.998 | 4.269 | 1.732 | 5.050 | 11.091 | 199722 |
|  | CMTK B | 5.737 | 4.156 | 1.732 | 4.950 | 10.198 | 184238 |
|  | CMTK C | 5.385 | 3.441 | 1.732 | 4.796 | 9.513 | 176718 |
|  | Elastix A | 6.131 | 4.711 | 2.000 | 5.196 | 10.770 | 211960 |
|  | Elastix B | 6.655 | 6.056 | 2.000 | 5.196 | 12.104 | 189138 |
| Tefor | ANTs A | 5.009 | 3.438 | 1.581 | 4.472 | 8.718 | 172432 |
|  | ANTs B | 5.351 | 3.486 | 1.732 | 4.743 | 9.301 | 193554 |
|  | ANTs C | 5.773 | 3.943 | 1.732 | 5.000 | 10.223 | 199872 |
|  | CMTK A | 5.981 | 4.548 | 1.581 | 5.000 | 11.269 | 200804 |
|  | CMTK B | 6.250 | 4.823 | 1.732 | 5.148 | 11.832 | 198428 |
|  | CMTK C | 5.844 | 3.845 | 2.000 | 5.099 | 10.512 | 201972 |
|  | Elastix A | 5.109 | 3.341 | 1.581 | 4.528 | 8.972 | 163504 |
|  | Elastix B | 5.102 | 3.337 | 1.581 | 4.528 | 9.110 | 196782 |

**Table S65:** PLP_L : posterior lateral protocerebrum

| Template | Algorithm | Mean | Std dev | 10th perc | median | 90th perc | N |
| --- | --- | --- | --- | --- | --- | --- | --- |
| JFRC2010 | ANTs A | 6.063 | 6.078 | 0.707 | 3.808 | 15.067 | 425196 |
|  | ANTs B | 4.337 | 4.460 | 0.707 | 2.915 | 10.075 | 412812 |
|  | ANTs C | 4.621 | 4.692 | 0.707 | 3.000 | 10.440 | 373646 |
|  | CMTK A | 6.856 | 7.376 | 0.707 | 3.606 | 18.111 | 583892 |
|  | CMTK B | 5.412 | 6.127 | 0.707 | 2.828 | 14.595 | 487112 |
|  | CMTK C | 7.067 | 6.968 | 0.707 | 4.359 | 17.734 | 485036 |
|  | Elastix A | 3.563 | 4.048 | 0.707 | 2.121 | 8.746 | 427644 |
|  | Elastix B | 3.517 | 4.245 | 0.707 | 2.000 | 9.000 | 437410 |
| JFRC2013 | ANTs A | 3.314 | 4.099 | 0.707 | 1.732 | 8.093 | 445348 |
|  | ANTs B | 4.336 | 4.214 | 0.707 | 3.000 | 9.539 | 434152 |
|  | ANTs C | 4.099 | 4.118 | 0.707 | 2.828 | 9.110 | 446134 |
|  | CMTK A | 3.938 | 4.224 | 0.707 | 2.550 | 9.110 | 414136 |
|  | CMTK B | 4.019 | 4.284 | 0.707 | 2.550 | 9.247 | 404460 |
|  | CMTK C | 4.389 | 4.287 | 0.707 | 3.000 | 9.849 | 405546 |
|  | Elastix A | 3.376 | 4.025 | 0.707 | 2.000 | 8.155 | 428920 |
|  | Elastix B | 3.304 | 4.045 | 0.707 | 1.732 | 8.031 | 437122 |
| JRC2018 | ANTs A | 3.290 | 4.079 | 0.707 | 1.732 | 8.093 | 455796 |
|  | ANTs B | 3.428 | 4.028 | 0.707 | 2.121 | 8.124 | 465462 |
|  | ANTs C | 3.581 | 4.013 | 0.707 | 2.121 | 8.337 | 474688 |
|  | CMTK A | 3.277 | 4.041 | 0.707 | 1.732 | 8.031 | 446974 |
|  | CMTK B | 3.242 | 4.028 | 0.707 | 1.732 | 7.906 | 441626 |
|  | CMTK C | 3.317 | 4.074 | 0.707 | 2.000 | 7.937 | 434284 |
|  | Elastix A | 3.348 | 4.067 | 0.707 | 2.000 | 8.062 | 441954 |
|  | Elastix B | 3.332 | 4.099 | 0.707 | 1.732 | 8.185 | 444548 |
| FCWB | ANTs A | 3.598 | 4.510 | 0.707 | 2.000 | 9.055 | 475416 |
|  | ANTs B | 4.143 | 4.617 | 0.707 | 2.449 | 10.000 | 424340 |
|  | ANTs C | 4.230 | 4.600 | 0.707 | 2.646 | 9.925 | 420268 |
|  | CMTK A | 5.873 | 6.766 | 0.707 | 3.536 | 13.964 | 366682 |
|  | CMTK B | 5.520 | 6.419 | 0.707 | 3.317 | 12.903 | 352618 |
|  | CMTK C | 4.261 | 4.896 | 0.707 | 2.550 | 10.512 | 398244 |
|  | Elastix A | 6.091 | 5.778 | 1.000 | 4.243 | 13.946 | 338514 |
|  | Elastix B | 5.045 | 6.085 | 0.707 | 2.345 | 13.656 | 399758 |
| Tefor | ANTs A | 3.298 | 4.093 | 0.707 | 1.732 | 8.093 | 464996 |
|  | ANTs B | 3.830 | 4.210 | 0.707 | 2.345 | 8.972 | 452548 |
|  | ANTs C | 3.920 | 4.255 | 0.707 | 2.449 | 9.055 | 482426 |
|  | CMTK A | 4.351 | 6.384 | 0.707 | 2.000 | 11.705 | 443760 |
|  | CMTK B | 4.445 | 6.383 | 0.707 | 2.121 | 12.104 | 442604 |
|  | CMTK C | 4.642 | 5.547 | 0.707 | 2.646 | 11.402 | 446180 |
|  | Elastix A | 3.475 | 4.037 | 0.707 | 2.121 | 8.515 | 443568 |
|  | Elastix B | 3.436 | 4.098 | 0.707 | 2.000 | 8.718 | 442816 |

**Table S66:** AOTU_L : anterior optic tubercle

| Template | Algorithm | Mean | Std dev | 10th perc | median | 90th perc | N |
| --- | --- | --- | --- | --- | --- | --- | --- |
| JFRC2010 | ANTs A | 16.330 | 9.444 | 4.583 | 15.330 | 29.283 | 80554 |
|  | ANTs B | 8.677 | 7.334 | 1.581 | 6.245 | 19.609 | 14514 |
|  | ANTs C | 8.930 | 6.973 | 2.000 | 7.000 | 18.788 | 17056 |
|  | CMTK A | 12.808 | 10.303 | 1.732 | 10.223 | 29.034 | 17914 |
|  | CMTK B | 13.304 | 10.055 | 2.121 | 11.424 | 28.783 | 22412 |
|  | CMTK C | 13.049 | 9.556 | 2.550 | 10.817 | 27.295 | 16882 |
|  | Elastix A | 8.749 | 8.171 | 1.581 | 5.568 | 19.929 | 13058 |
|  | Elastix B | 8.253 | 7.852 | 1.414 | 5.196 | 19.925 | 14726 |
| JFRC2013 | ANTs A | 8.395 | 7.372 | 1.581 | 5.831 | 18.173 | 13596 |
|  | ANTs B | 8.361 | 6.597 | 1.581 | 6.481 | 18.014 | 31014 |
|  | ANTs C | 7.389 | 6.613 | 1.225 | 5.196 | 16.872 | 30028 |
|  | CMTK A | 12.449 | 8.885 | 2.121 | 11.380 | 25.159 | 19976 |
|  | CMTK B | 12.121 | 8.747 | 2.000 | 10.886 | 24.546 | 20068 |
|  | CMTK C | 9.563 | 7.412 | 2.121 | 7.550 | 20.591 | 16956 |
|  | Elastix A | 8.655 | 7.742 | 1.581 | 5.874 | 19.248 | 13438 |
|  | Elastix B | 8.887 | 8.003 | 1.581 | 6.083 | 19.975 | 13216 |
| JRC2018 | ANTs A | 8.895 | 8.129 | 1.581 | 5.788 | 20.906 | 13630 |
|  | ANTs B | 8.600 | 8.033 | 1.581 | 5.339 | 20.809 | 12338 |
|  | ANTs C | 8.763 | 7.562 | 1.581 | 6.364 | 19.717 | 14006 |
|  | CMTK A | 8.795 | 7.937 | 1.581 | 5.874 | 19.962 | 13928 |
|  | CMTK B | 9.002 | 8.064 | 1.581 | 5.916 | 20.444 | 13452 |
|  | CMTK C | 8.805 | 7.955 | 1.581 | 5.745 | 20.112 | 12312 |
|  | Elastix A | 8.529 | 7.943 | 1.581 | 5.523 | 19.187 | 13318 |
|  | Elastix B | 8.397 | 7.758 | 1.581 | 5.523 | 19.264 | 14250 |
| FCWB | ANTs A | 11.510 | 7.480 | 2.915 | 10.296 | 21.920 | 33530 |
|  | ANTs B | 6.655 | 7.305 | 1.000 | 3.808 | 17.960 | 15710 |
|  | ANTs C | 6.117 | 6.348 | 1.000 | 4.123 | 14.560 | 24774 |
|  | CMTK A | 9.340 | 7.220 | 2.121 | 7.416 | 19.352 | 14154 |
|  | CMTK B | 9.453 | 7.078 | 2.236 | 7.550 | 19.506 | 14176 |
|  | CMTK C | 7.192 | 6.623 | 1.225 | 5.050 | 16.643 | 24994 |
|  | Elastix A | 8.935 | 7.286 | 1.581 | 7.000 | 18.577 | 13950 |
|  | Elastix B | 8.916 | 7.954 | 1.581 | 6.364 | 20.106 | 15566 |
| Tefor | ANTs A | 8.627 | 7.949 | 1.581 | 5.523 | 20.012 | 14782 |
|  | ANTs B | 8.462 | 7.386 | 1.581 | 6.042 | 19.153 | 16114 |
|  | ANTs C | 7.989 | 6.187 | 1.732 | 6.364 | 16.462 | 31686 |
|  | CMTK A | 11.916 | 8.926 | 1.732 | 10.817 | 24.030 | 18692 |
|  | CMTK B | 11.288 | 8.859 | 1.732 | 8.860 | 23.675 | 19294 |
|  | CMTK C | 7.860 | 7.183 | 1.225 | 5.523 | 18.865 | 30052 |
|  | Elastix A | 8.494 | 7.825 | 1.414 | 5.568 | 19.261 | 13916 |
|  | Elastix B | 8.488 | 7.832 | 1.414 | 6.000 | 19.365 | 16528 |

**Table S67:** LH_R : lateral horn

| Template | Algorithm | Mean | Std dev | 10th perc | median | 90th perc | N |
| --- | --- | --- | --- | --- | --- | --- | --- |
| JFRC2010 | ANTs A | 6.263 | 4.807 | 1.581 | 5.000 | 12.669 | 306006 |
|  | ANTs B | 5.442 | 3.679 | 1.581 | 4.690 | 10.223 | 269522 |
|  | ANTs C | 5.904 | 4.146 | 1.581 | 5.000 | 11.358 | 304562 |
|  | CMTK A | 5.675 | 4.171 | 1.414 | 4.690 | 11.336 | 295072 |
|  | CMTK B | 5.607 | 4.386 | 1.414 | 4.528 | 11.203 | 282530 |
|  | CMTK C | 6.292 | 4.895 | 1.581 | 5.000 | 12.590 | 258870 |
|  | Elastix A | 4.858 | 3.502 | 1.414 | 4.123 | 9.247 | 286844 |
|  | Elastix B | 4.880 | 3.421 | 1.414 | 4.183 | 9.192 | 305128 |
| JFRC2013 | ANTs A | 5.114 | 3.581 | 1.581 | 4.359 | 9.618 | 318536 |
|  | ANTs B | 5.922 | 4.020 | 1.581 | 5.099 | 11.269 | 259806 |
|  | ANTs C | 6.058 | 4.162 | 1.581 | 5.148 | 11.597 | 314784 |
|  | CMTK A | 5.545 | 3.978 | 1.581 | 4.528 | 11.045 | 351580 |
|  | CMTK B | 5.460 | 3.869 | 1.581 | 4.528 | 10.840 | 350916 |
|  | CMTK C | 5.422 | 4.163 | 1.225 | 4.472 | 11.068 | 340114 |
|  | Elastix A | 5.051 | 3.591 | 1.581 | 4.243 | 9.644 | 283216 |
|  | Elastix B | 4.972 | 3.532 | 1.414 | 4.183 | 9.460 | 322730 |
| JRC2018 | ANTs A | 4.858 | 3.416 | 1.414 | 4.183 | 9.165 | 327132 |
|  | ANTs B | 5.205 | 3.477 | 1.581 | 4.528 | 9.644 | 315620 |
|  | ANTs C | 5.356 | 3.547 | 1.581 | 4.637 | 10.050 | 338732 |
|  | CMTK A | 4.935 | 3.424 | 1.414 | 4.243 | 9.247 | 326348 |
|  | CMTK B | 4.890 | 3.416 | 1.414 | 4.183 | 9.192 | 316248 |
|  | CMTK C | 4.953 | 3.447 | 1.414 | 4.243 | 9.301 | 311878 |
|  | Elastix A | 4.979 | 3.523 | 1.414 | 4.243 | 9.460 | 307772 |
|  | Elastix B | 5.027 | 3.514 | 1.581 | 4.301 | 9.434 | 313066 |
| FCWB | ANTs A | 5.352 | 3.653 | 1.581 | 4.583 | 10.050 | 346104 |
|  | ANTs B | 6.272 | 4.078 | 2.000 | 5.431 | 11.683 | 267224 |
|  | ANTs C | 6.367 | 4.302 | 1.732 | 5.431 | 12.104 | 293326 |
|  | CMTK A | 6.098 | 4.049 | 1.732 | 5.196 | 11.597 | 289148 |
|  | CMTK B | 5.860 | 3.918 | 1.581 | 5.000 | 11.269 | 297822 |
|  | CMTK C | 6.310 | 4.455 | 1.732 | 5.292 | 12.186 | 278118 |
|  | Elastix A | 6.531 | 4.648 | 1.732 | 5.431 | 12.689 | 276434 |
|  | Elastix B | 6.760 | 5.661 | 1.581 | 5.148 | 13.946 | 303830 |
| Tefor | ANTs A | 4.832 | 3.462 | 1.225 | 4.123 | 9.220 | 326606 |
|  | ANTs B | 5.288 | 3.638 | 1.581 | 4.528 | 9.925 | 301662 |
|  | ANTs C | 5.553 | 3.919 | 1.581 | 4.743 | 10.512 | 332776 |
|  | CMTK A | 7.508 | 7.925 | 1.414 | 4.743 | 19.313 | 276804 |
|  | CMTK B | 7.826 | 8.477 | 1.414 | 4.743 | 21.237 | 266390 |
|  | CMTK C | 6.622 | 6.748 | 1.581 | 4.690 | 13.342 | 282262 |
|  | Elastix A | 4.994 | 3.577 | 1.414 | 4.243 | 9.513 | 307694 |
|  | Elastix B | 5.162 | 3.549 | 1.581 | 4.472 | 9.747 | 306252 |

**Table S68:** GOR_L : gorget

| Template | Algorithm | Mean | Std dev | 10th perc | median | 90th perc | N |
| --- | --- | --- | --- | --- | --- | --- | --- |
| JFRC2010 | ANTs A | 5.602 | 4.184 | 1.414 | 4.528 | 11.424 | 87290 |
|  | ANTs B | 4.035 | 3.398 | 0.707 | 3.000 | 8.746 | 88176 |
|  | ANTs C | 4.587 | 3.902 | 0.707 | 3.464 | 10.223 | 72964 |
|  | CMTK A | 5.297 | 5.320 | 0.707 | 3.240 | 12.903 | 82058 |
|  | CMTK B | 4.871 | 5.090 | 0.707 | 3.000 | 11.489 | 76474 |
|  | CMTK C | 4.541 | 4.274 | 0.707 | 3.162 | 10.247 | 72248 |
|  | Elastix A | 3.501 | 2.745 | 0.707 | 2.828 | 7.106 | 81146 |
|  | Elastix B | 3.509 | 2.892 | 0.707 | 2.646 | 7.106 | 82774 |
| JFRC2013 | ANTs A | 3.289 | 2.510 | 0.707 | 2.646 | 6.708 | 86924 |
|  | ANTs B | 4.300 | 3.435 | 0.707 | 3.464 | 9.110 | 89722 |
|  | ANTs C | 4.520 | 3.518 | 1.000 | 3.606 | 9.618 | 86652 |
|  | CMTK A | 3.375 | 2.802 | 0.707 | 2.646 | 7.106 | 75820 |
|  | CMTK B | 3.466 | 2.862 | 0.707 | 2.646 | 7.211 | 77116 |
|  | CMTK C | 4.012 | 3.141 | 1.000 | 3.240 | 7.906 | 89236 |
|  | Elastix A | 3.411 | 2.688 | 0.707 | 2.646 | 7.106 | 78798 |
|  | Elastix B | 3.237 | 2.525 | 0.707 | 2.550 | 6.708 | 78848 |
| JRC2018 | ANTs A | 3.295 | 2.500 | 0.707 | 2.646 | 6.708 | 87562 |
|  | ANTs B | 3.520 | 2.700 | 0.707 | 2.915 | 7.141 | 87832 |
|  | ANTs C | 3.407 | 2.715 | 0.707 | 2.646 | 7.071 | 80968 |
|  | CMTK A | 3.346 | 2.707 | 0.707 | 2.646 | 7.141 | 71502 |
|  | CMTK B | 3.344 | 2.685 | 0.707 | 2.646 | 7.106 | 73202 |
|  | CMTK C | 3.420 | 2.605 | 0.707 | 2.828 | 7.000 | 84368 |
|  | Elastix A | 3.347 | 2.504 | 0.707 | 2.646 | 6.708 | 93686 |
|  | Elastix B | 3.283 | 2.490 | 0.707 | 2.646 | 6.671 | 86832 |
| FCWB | ANTs A | 3.230 | 2.593 | 0.707 | 2.550 | 6.708 | 82426 |
|  | ANTs B | 3.866 | 3.098 | 1.000 | 3.082 | 7.649 | 90258 |
|  | ANTs C | 3.637 | 2.769 | 1.000 | 2.915 | 7.348 | 90864 |
|  | CMTK A | 7.472 | 7.195 | 1.225 | 4.690 | 19.609 | 56578 |
|  | CMTK B | 6.593 | 6.545 | 1.225 | 4.272 | 16.823 | 67432 |
|  | CMTK C | 3.822 | 3.245 | 0.707 | 2.915 | 8.062 | 88790 |
|  | Elastix A | 9.001 | 8.786 | 1.000 | 5.568 | 24.525 | 75406 |
|  | Elastix B | 4.288 | 3.913 | 0.707 | 3.000 | 9.618 | 92154 |
| Tefor | ANTs A | 3.322 | 2.569 | 0.707 | 2.646 | 6.856 | 85508 |
|  | ANTs B | 3.771 | 2.933 | 0.707 | 3.000 | 7.810 | 87320 |
|  | ANTs C | 3.747 | 2.940 | 0.707 | 3.000 | 7.842 | 83340 |
|  | CMTK A | 4.081 | 4.368 | 0.707 | 2.646 | 8.860 | 68066 |
|  | CMTK B | 4.000 | 4.067 | 0.707 | 2.646 | 8.631 | 69704 |
|  | CMTK C | 3.973 | 3.401 | 0.707 | 3.000 | 8.093 | 88896 |
|  | Elastix A | 3.337 | 2.629 | 0.707 | 2.646 | 6.745 | 83424 |
|  | Elastix B | 3.740 | 2.994 | 1.000 | 2.915 | 7.906 | 80286 |

**Table S69:** MB_CA_L : calyx of adult mushroom body

| Template | Algorithm | Mean | Std dev | 10th perc | median | 90th perc | N |
| --- | --- | --- | --- | --- | --- | --- | --- |
| JFRC2010 | ANTs A | 4.603 | 3.060 | 1.414 | 4.000 | 8.660 | 386392 |
|  | ANTs B | 5.159 | 3.797 | 1.414 | 4.359 | 9.849 | 282626 |
|  | ANTs C | 4.697 | 3.345 | 1.225 | 4.062 | 8.972 | 359548 |
|  | CMTK A | 4.702 | 3.431 | 1.225 | 3.873 | 9.110 | 366014 |
|  | CMTK B | 4.523 | 3.054 | 1.225 | 3.873 | 8.515 | 386536 |
|  | CMTK C | 5.126 | 3.609 | 1.414 | 4.359 | 9.874 | 275910 |
|  | Elastix A | 4.364 | 2.937 | 1.225 | 3.808 | 8.155 | 316702 |
|  | Elastix B | 4.167 | 2.732 | 1.225 | 3.606 | 7.810 | 350304 |
| JFRC2013 | ANTs A | 4.462 | 2.962 | 1.225 | 3.808 | 8.515 | 319094 |
|  | ANTs B | 4.945 | 3.472 | 1.414 | 4.183 | 9.460 | 322084 |
|  | ANTs C | 4.926 | 3.366 | 1.414 | 4.183 | 9.434 | 320916 |
|  | CMTK A | 3.856 | 2.362 | 1.225 | 3.536 | 7.036 | 255508 |
|  | CMTK B | 4.038 | 2.488 | 1.225 | 3.606 | 7.280 | 250828 |
|  | CMTK C | 5.119 | 3.194 | 1.581 | 4.528 | 9.539 | 196572 |
|  | Elastix A | 3.841 | 2.367 | 1.000 | 3.464 | 7.106 | 322206 |
|  | Elastix B | 3.630 | 2.278 | 1.000 | 3.162 | 6.708 | 320008 |
| JRC2018 | ANTs A | 3.620 | 2.362 | 1.000 | 3.082 | 6.856 | 393546 |
|  | ANTs B | 4.268 | 3.009 | 1.000 | 3.606 | 8.062 | 369874 |
|  | ANTs C | 4.068 | 2.739 | 1.000 | 3.536 | 7.649 | 394530 |
|  | CMTK A | 3.851 | 2.473 | 1.000 | 3.317 | 7.280 | 413950 |
|  | CMTK B | 3.913 | 2.500 | 1.000 | 3.464 | 7.416 | 401238 |
|  | CMTK C | 4.106 | 2.579 | 1.225 | 3.606 | 7.649 | 382438 |
|  | Elastix A | 4.150 | 2.688 | 1.225 | 3.606 | 7.810 | 376294 |
|  | Elastix B | 3.766 | 2.355 | 1.000 | 3.317 | 7.036 | 382706 |
| FCWB | ANTs A | 4.460 | 3.043 | 1.225 | 3.808 | 8.544 | 310260 |
|  | ANTs B | 5.154 | 3.706 | 1.414 | 4.359 | 9.849 | 250986 |
|  | ANTs C | 4.784 | 3.468 | 1.225 | 4.062 | 9.220 | 304398 |
|  | CMTK A | 5.401 | 4.312 | 1.225 | 4.243 | 11.180 | 354154 |
|  | CMTK B | 5.091 | 3.900 | 1.225 | 4.183 | 10.025 | 353338 |
|  | CMTK C | 5.454 | 4.031 | 1.414 | 4.528 | 10.700 | 226302 |
|  | Elastix A | 6.715 | 5.527 | 1.225 | 5.000 | 15.116 | 243188 |
|  | Elastix B | 5.876 | 5.514 | 1.225 | 4.183 | 13.058 | 279418 |
| Tefor | ANTs A | 3.567 | 2.322 | 1.000 | 3.082 | 6.745 | 390314 |
|  | ANTs B | 4.780 | 3.532 | 1.225 | 4.062 | 9.083 | 266852 |
|  | ANTs C | 4.939 | 3.731 | 1.225 | 4.062 | 9.721 | 328196 |
|  | CMTK A | 5.220 | 5.363 | 1.000 | 3.606 | 10.416 | 352240 |
|  | CMTK B | 5.390 | 5.499 | 1.225 | 3.808 | 10.863 | 337534 |
|  | CMTK C | 5.253 | 4.117 | 1.414 | 4.301 | 10.000 | 261764 |
|  | Elastix A | 3.979 | 2.581 | 1.225 | 3.536 | 7.382 | 338726 |
|  | Elastix B | 3.839 | 2.465 | 1.000 | 3.317 | 7.106 | 357760 |

**Table S70:** SPS_L : superior posterior slope

| Template | Algorithm | Mean | Std dev | 10th perc | median | 90th perc | N |
| --- | --- | --- | --- | --- | --- | --- | --- |
| JFRC2010 | ANTs A | 7.310 | 6.162 | 1.581 | 5.431 | 16.279 | 494082 |
|  | ANTs B | 5.325 | 4.211 | 1.225 | 4.243 | 10.817 | 451332 |
|  | ANTs C | 5.226 | 3.977 | 1.225 | 4.243 | 10.440 | 481422 |
|  | CMTK A | 6.758 | 6.100 | 1.225 | 4.690 | 15.556 | 310332 |
|  | CMTK B | 6.426 | 5.861 | 1.225 | 4.472 | 14.748 | 335610 |
|  | CMTK C | 6.591 | 6.309 | 1.225 | 4.528 | 15.524 | 465200 |
|  | Elastix A | 4.177 | 3.341 | 1.000 | 3.317 | 8.276 | 430482 |
|  | Elastix B | 4.552 | 3.815 | 1.000 | 3.606 | 9.083 | 358604 |
| JFRC2013 | ANTs A | 4.277 | 3.570 | 1.000 | 3.240 | 8.775 | 386086 |
|  | ANTs B | 5.284 | 4.153 | 1.225 | 4.183 | 10.794 | 423082 |
|  | ANTs C | 5.146 | 4.120 | 1.225 | 4.062 | 10.512 | 413470 |
|  | CMTK A | 4.507 | 3.717 | 1.000 | 3.536 | 9.055 | 356370 |
|  | CMTK B | 4.563 | 3.702 | 1.000 | 3.606 | 9.192 | 363436 |
|  | CMTK C | 5.221 | 4.367 | 1.225 | 4.062 | 10.654 | 425034 |
|  | Elastix A | 4.205 | 3.479 | 1.000 | 3.240 | 8.515 | 390606 |
|  | Elastix B | 4.310 | 3.698 | 1.000 | 3.240 | 8.660 | 360798 |
| JRC2018 | ANTs A | 4.200 | 3.541 | 1.000 | 3.240 | 8.602 | 390572 |
|  | ANTs B | 4.684 | 3.780 | 1.000 | 3.606 | 9.539 | 376408 |
|  | ANTs C | 4.657 | 3.676 | 1.000 | 3.674 | 9.301 | 357000 |
|  | CMTK A | 4.666 | 3.860 | 1.000 | 3.606 | 9.327 | 330466 |
|  | CMTK B | 4.576 | 3.824 | 1.000 | 3.606 | 9.247 | 321912 |
|  | CMTK C | 4.444 | 3.730 | 1.000 | 3.536 | 8.944 | 327862 |
|  | Elastix A | 4.273 | 3.554 | 1.000 | 3.317 | 8.602 | 377022 |
|  | Elastix B | 4.377 | 3.704 | 1.000 | 3.464 | 8.820 | 358098 |
| FCWB | ANTs A | 4.690 | 3.962 | 1.000 | 3.606 | 9.618 | 405374 |
|  | ANTs B | 5.129 | 4.091 | 1.225 | 4.062 | 10.392 | 423360 |
|  | ANTs C | 4.792 | 3.673 | 1.225 | 3.808 | 9.592 | 415376 |
|  | CMTK A | 6.758 | 6.919 | 1.414 | 4.637 | 14.646 | 293282 |
|  | CMTK B | 6.170 | 6.604 | 1.225 | 4.183 | 12.903 | 327190 |
|  | CMTK C | 5.473 | 5.267 | 1.225 | 3.808 | 11.511 | 374896 |
|  | Elastix A | 6.416 | 5.949 | 1.225 | 4.528 | 14.526 | 435286 |
|  | Elastix B | 5.420 | 5.013 | 1.000 | 3.808 | 12.042 | 451186 |
| Tefor | ANTs A | 4.215 | 3.524 | 1.000 | 3.240 | 8.602 | 397556 |
|  | ANTs B | 4.961 | 4.020 | 1.225 | 3.873 | 10.149 | 424108 |
|  | ANTs C | 5.222 | 3.927 | 1.414 | 4.301 | 10.271 | 393994 |
|  | CMTK A | 5.359 | 5.153 | 1.000 | 3.808 | 11.402 | 340272 |
|  | CMTK B | 5.429 | 5.183 | 1.000 | 3.808 | 11.533 | 340788 |
|  | CMTK C | 5.132 | 4.455 | 1.225 | 3.873 | 10.607 | 447014 |
|  | Elastix A | 4.159 | 3.403 | 1.000 | 3.240 | 8.155 | 417846 |
|  | Elastix B | 4.672 | 3.709 | 1.225 | 3.674 | 9.220 | 359298 |

**Table S71:** IPS_L : inferior posterior slope

| Template | Algorithm | Mean | Std dev | 10th perc | median | 90th perc | N |
| --- | --- | --- | --- | --- | --- | --- | --- |
| JFRC2010 | ANTs A | 10.205 | 7.498 | 2.550 | 8.485 | 19.875 | 209044 |
|  | ANTs B | 7.295 | 4.887 | 2.121 | 6.205 | 14.160 | 183132 |
|  | ANTs C | 7.084 | 4.813 | 2.121 | 6.000 | 13.620 | 193510 |
|  | CMTK A | 9.097 | 5.863 | 2.345 | 7.937 | 17.734 | 167668 |
|  | CMTK B | 8.030 | 5.571 | 2.121 | 6.856 | 15.906 | 166860 |
|  | CMTK C | 8.026 | 5.762 | 2.000 | 6.442 | 16.568 | 213410 |
|  | Elastix A | 6.012 | 4.463 | 1.581 | 5.000 | 12.042 | 192168 |
|  | Elastix B | 6.456 | 4.616 | 1.732 | 5.339 | 12.767 | 192986 |
| JFRC2013 | ANTs A | 6.601 | 5.144 | 1.581 | 5.196 | 13.874 | 188088 |
|  | ANTs B | 7.082 | 4.713 | 2.121 | 6.083 | 13.675 | 176902 |
|  | ANTs C | 7.015 | 4.674 | 2.121 | 5.916 | 13.583 | 186466 |
|  | CMTK A | 6.739 | 4.972 | 1.581 | 5.568 | 13.342 | 169248 |
|  | CMTK B | 6.843 | 5.034 | 1.581 | 5.701 | 13.656 | 165164 |
|  | CMTK C | 7.357 | 4.957 | 2.121 | 6.164 | 14.474 | 189470 |
|  | Elastix A | 6.205 | 4.904 | 1.414 | 4.796 | 13.058 | 181192 |
|  | Elastix B | 6.383 | 4.985 | 1.581 | 5.050 | 13.058 | 180334 |
| JRC2018 | ANTs A | 6.002 | 4.824 | 1.414 | 4.583 | 12.369 | 185274 |
|  | ANTs B | 7.113 | 5.055 | 1.732 | 5.916 | 14.071 | 172736 |
|  | ANTs C | 7.175 | 4.979 | 2.000 | 6.042 | 14.089 | 171534 |
|  | CMTK A | 6.512 | 4.920 | 1.581 | 5.196 | 13.285 | 167674 |
|  | CMTK B | 6.583 | 4.995 | 1.581 | 5.339 | 13.416 | 161066 |
|  | CMTK C | 6.903 | 5.237 | 1.581 | 5.568 | 14.053 | 158340 |
|  | Elastix A | 6.178 | 4.929 | 1.414 | 4.796 | 12.806 | 184982 |
|  | Elastix B | 6.339 | 4.904 | 1.581 | 5.050 | 13.019 | 176462 |
| FCWB | ANTs A | 6.598 | 4.791 | 1.581 | 5.431 | 13.435 | 197364 |
|  | ANTs B | 6.999 | 4.749 | 2.000 | 6.042 | 13.304 | 162610 |
|  | ANTs C | 6.932 | 4.802 | 2.000 | 5.874 | 13.435 | 157050 |
|  | CMTK A | 9.012 | 6.139 | 2.236 | 7.778 | 17.607 | 247140 |
|  | CMTK B | 8.812 | 6.140 | 2.236 | 7.416 | 17.364 | 216198 |
|  | CMTK C | 7.849 | 5.540 | 2.000 | 6.442 | 16.016 | 189734 |
|  | Elastix A | 8.489 | 6.017 | 2.121 | 7.106 | 16.643 | 272580 |
|  | Elastix B | 8.059 | 6.710 | 1.581 | 6.205 | 16.778 | 286162 |
| Tefor | ANTs A | 6.097 | 4.774 | 1.414 | 4.796 | 12.669 | 184588 |
|  | ANTs B | 7.181 | 4.730 | 2.121 | 6.205 | 13.820 | 185350 |
|  | ANTs C | 7.416 | 4.873 | 2.236 | 6.364 | 14.248 | 173904 |
|  | CMTK A | 7.824 | 5.968 | 1.732 | 6.364 | 16.000 | 207344 |
|  | CMTK B | 8.036 | 6.168 | 1.732 | 6.481 | 16.583 | 199956 |
|  | CMTK C | 7.281 | 5.378 | 1.732 | 6.083 | 14.248 | 219074 |
|  | Elastix A | 5.726 | 4.444 | 1.414 | 4.528 | 11.597 | 200776 |
|  | Elastix B | 6.505 | 4.685 | 1.581 | 5.385 | 12.767 | 197976 |

**Table S72:** SCL_L : superior clamp

| Template | Algorithm | Mean | Std dev | 10th perc | median | 90th perc | N |
| --- | --- | --- | --- | --- | --- | --- | --- |
| JFRC2010 | ANTs A | 5.455 | 6.304 | 0.707 | 3.162 | 13.285 | 365908 |
|  | ANTs B | 3.064 | 3.233 | 0.707 | 2.000 | 7.314 | 310344 |
|  | ANTs C | 3.262 | 3.507 | 0.707 | 2.000 | 8.062 | 287574 |
|  | CMTK A | 3.271 | 4.042 | 0.707 | 1.732 | 8.660 | 316734 |
|  | CMTK B | 4.493 | 5.100 | 0.707 | 2.236 | 12.288 | 378216 |
|  | CMTK C | 4.372 | 4.872 | 0.707 | 2.345 | 11.597 | 354038 |
|  | Elastix A | 2.718 | 2.975 | 0.707 | 1.581 | 6.403 | 313594 |
|  | Elastix B | 2.957 | 3.658 | 0.707 | 1.581 | 7.036 | 302568 |
| JFRC2013 | ANTs A | 2.501 | 2.654 | 0.707 | 1.581 | 5.831 | 330144 |
|  | ANTs B | 3.351 | 3.483 | 0.707 | 2.121 | 8.124 | 306500 |
|  | ANTs C | 3.029 | 3.191 | 0.707 | 1.732 | 7.246 | 325802 |
|  | CMTK A | 3.161 | 2.930 | 0.707 | 2.236 | 6.964 | 308818 |
|  | CMTK B | 3.173 | 2.976 | 0.707 | 2.236 | 7.000 | 309956 |
|  | CMTK C | 3.734 | 3.275 | 0.707 | 2.828 | 7.937 | 296110 |
|  | Elastix A | 2.585 | 2.750 | 0.707 | 1.581 | 6.083 | 327732 |
|  | Elastix B | 2.587 | 2.758 | 0.707 | 1.581 | 6.083 | 324458 |
| JRC2018 | ANTs A | 2.486 | 2.641 | 0.707 | 1.581 | 5.788 | 333308 |
|  | ANTs B | 2.673 | 2.819 | 0.707 | 1.581 | 6.364 | 328600 |
|  | ANTs C | 2.700 | 2.774 | 0.707 | 1.732 | 6.403 | 328444 |
|  | CMTK A | 2.446 | 2.604 | 0.707 | 1.581 | 5.701 | 326512 |
|  | CMTK B | 2.434 | 2.604 | 0.707 | 1.581 | 5.701 | 326858 |
|  | CMTK C | 2.493 | 2.660 | 0.707 | 1.581 | 5.745 | 323692 |
|  | Elastix A | 2.561 | 2.688 | 0.707 | 1.581 | 5.916 | 328392 |
|  | Elastix B | 2.569 | 2.845 | 0.707 | 1.581 | 6.083 | 332010 |
| FCWB | ANTs A | 3.331 | 3.599 | 0.707 | 2.000 | 8.185 | 319986 |
|  | ANTs B | 3.417 | 3.764 | 0.707 | 2.121 | 8.337 | 291926 |
|  | ANTs C | 3.354 | 3.612 | 0.707 | 2.121 | 8.246 | 304426 |
|  | CMTK A | 3.989 | 4.814 | 0.707 | 2.236 | 9.849 | 252984 |
|  | CMTK B | 3.966 | 4.561 | 0.707 | 2.345 | 9.618 | 258242 |
|  | CMTK C | 3.019 | 3.575 | 0.707 | 1.732 | 7.106 | 311560 |
|  | Elastix A | 4.944 | 5.453 | 0.707 | 2.915 | 12.530 | 261840 |
|  | Elastix B | 3.853 | 5.500 | 0.707 | 1.732 | 9.721 | 278840 |
| Tefor | ANTs A | 2.467 | 2.608 | 0.707 | 1.581 | 5.745 | 332778 |
|  | ANTs B | 2.821 | 2.956 | 0.707 | 1.732 | 6.671 | 318136 |
|  | ANTs C | 3.077 | 3.168 | 0.707 | 2.000 | 7.246 | 337976 |
|  | CMTK A | 2.942 | 4.012 | 0.707 | 1.581 | 6.964 | 320210 |
|  | CMTK B | 2.960 | 4.025 | 0.707 | 1.581 | 7.000 | 317748 |
|  | CMTK C | 3.010 | 3.521 | 0.707 | 1.732 | 6.856 | 323594 |
|  | Elastix A | 2.625 | 2.881 | 0.707 | 1.581 | 6.124 | 316456 |
|  | Elastix B | 3.013 | 3.655 | 0.707 | 1.732 | 7.246 | 297784 |

**Table S73:**
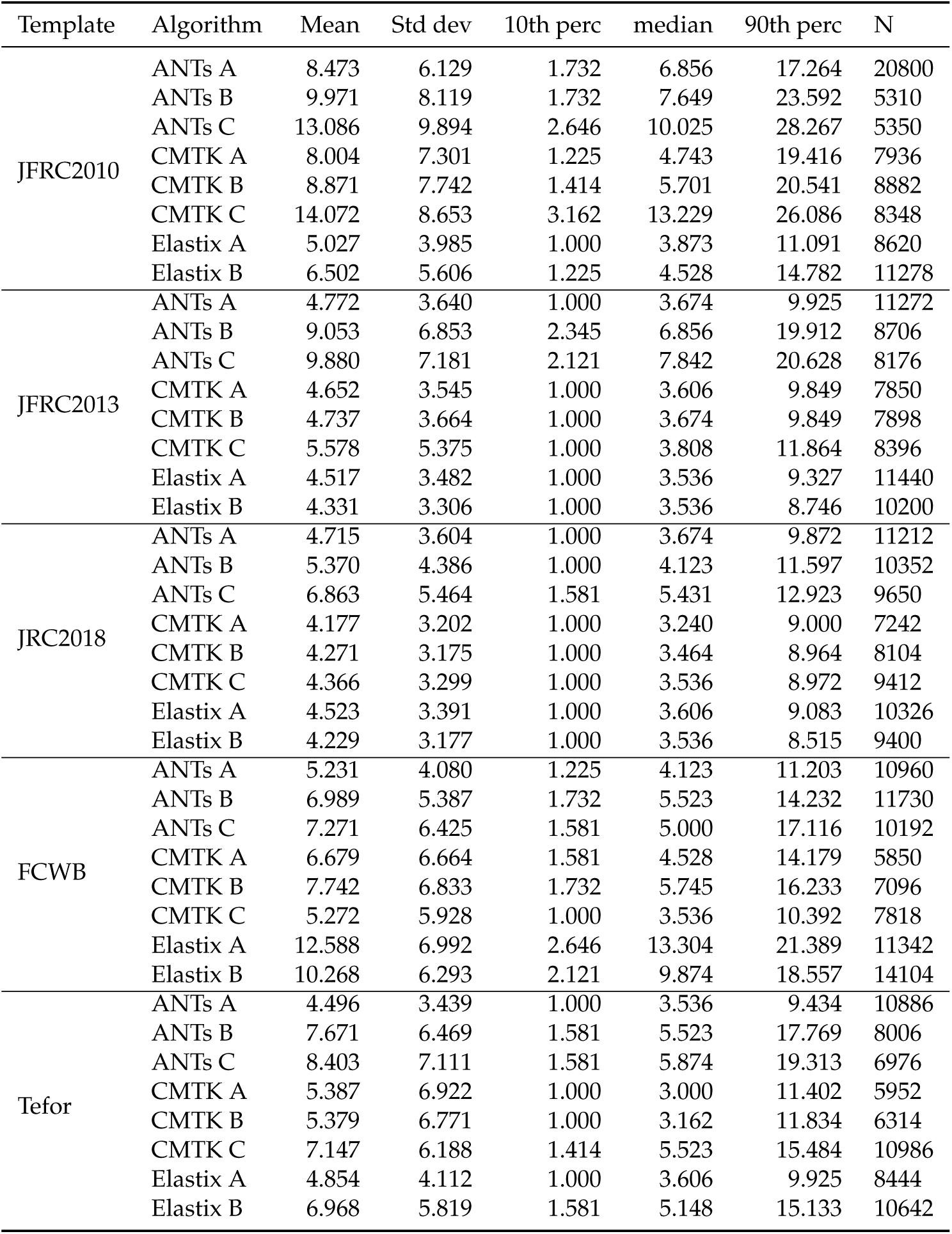
EPA_L : epaulette

**Table S74:** GA_L : gall

| Template | Algorithm | Mean | Std dev | 10th perc | median | 90th perc | N |
| --- | --- | --- | --- | --- | --- | --- | --- |
| JFRC2010 | ANTs A | 9.792 | 6.880 | 2.915 | 8.276 | 19.300 | 6906 |
|  | ANTs B | 7.664 | 3.981 | 2.646 | 7.141 | 13.304 | 5612 |
|  | ANTs C | 8.216 | 4.275 | 3.000 | 7.665 | 14.027 | 6166 |
|  | CMTK A | 8.411 | 4.059 | 3.082 | 8.276 | 13.675 | 4720 |
|  | CMTK B | 8.171 | 4.245 | 2.540 | 7.842 | 13.946 | 7860 |
|  | CMTK C | 8.291 | 4.460 | 2.646 | 8.031 | 14.370 | 7384 |
|  | Elastix A | 7.822 | 4.291 | 2.236 | 7.348 | 14.053 | 5158 |
|  | Elastix B | 7.306 | 3.983 | 2.121 | 7.036 | 12.669 | 4932 |
| JFRC2013 | ANTs A | 7.327 | 3.693 | 2.646 | 7.000 | 12.510 | 4904 |
|  | ANTs B | 8.966 | 4.455 | 3.464 | 8.544 | 15.149 | 3702 |
|  | ANTs C | 7.549 | 4.321 | 2.121 | 7.071 | 13.952 | 3488 |
|  | CMTK A | 8.723 | 4.345 | 3.162 | 8.515 | 14.474 | 5628 |
|  | CMTK B | 8.507 | 4.346 | 3.000 | 8.155 | 14.474 | 6310 |
|  | CMTK C | 8.175 | 4.181 | 3.162 | 7.416 | 13.964 | 4600 |
|  | Elastix A | 8.221 | 3.821 | 3.240 | 8.062 | 13.435 | 4282 |
|  | Elastix B | 7.716 | 4.024 | 2.550 | 7.314 | 13.388 | 5236 |
| JRC2018 | ANTs A | 7.185 | 3.763 | 2.345 | 6.708 | 12.530 | 5298 |
|  | ANTs B | 7.432 | 3.864 | 2.449 | 7.106 | 12.826 | 5168 |
|  | ANTs C | 7.357 | 3.788 | 2.646 | 6.856 | 12.708 | 6022 |
|  | CMTK A | 7.320 | 3.819 | 2.550 | 6.856 | 12.767 | 4962 |
|  | CMTK B | 7.285 | 3.863 | 2.345 | 6.856 | 12.927 | 4570 |
|  | CMTK C | 7.396 | 4.024 | 2.236 | 7.106 | 13.077 | 5154 |
|  | Elastix A | 7.352 | 3.767 | 2.345 | 7.141 | 12.530 | 5522 |
|  | Elastix B | 7.662 | 4.076 | 2.345 | 7.246 | 13.416 | 4790 |
| FCWB | ANTs A | 7.724 | 3.923 | 2.646 | 7.382 | 13.153 | 4658 |
|  | ANTs B | 7.151 | 4.437 | 2.000 | 6.481 | 13.058 | 5524 |
|  | ANTs C | 7.915 | 3.768 | 3.240 | 7.649 | 13.058 | 5910 |
|  | CMTK A | 7.525 | 3.861 | 2.646 | 7.211 | 13.088 | 5108 |
|  | CMTK B | 7.678 | 3.944 | 2.915 | 7.246 | 13.416 | 4854 |
|  | CMTK C | 8.755 | 3.762 | 4.183 | 8.276 | 14.230 | 3826 |
|  | Elastix A | 8.976 | 4.336 | 3.606 | 8.544 | 14.612 | 5108 |
|  | Elastix B | 7.824 | 3.881 | 2.646 | 7.616 | 13.077 | 5912 |
| Tefor | ANTs A | 7.285 | 3.799 | 2.449 | 6.856 | 12.765 | 5312 |
|  | ANTs B | 7.383 | 3.776 | 2.737 | 7.000 | 12.610 | 5956 |
|  | ANTs C | 8.357 | 4.225 | 3.041 | 8.031 | 13.964 | 6816 |
|  | CMTK A | 8.646 | 4.419 | 3.082 | 8.185 | 14.577 | 4492 |
|  | CMTK B | 9.250 | 4.824 | 3.162 | 9.301 | 15.182 | 3766 |
|  | CMTK C | 9.734 | 4.802 | 3.536 | 9.301 | 16.553 | 4354 |
|  | Elastix A | 7.722 | 4.249 | 2.000 | 7.416 | 13.583 | 4586 |
|  | Elastix B | 7.232 | 3.787 | 2.345 | 7.036 | 12.300 | 5808 |

**Table S75:** LAL_R : lateral accessory lobe

| Template | Algorithm | Mean | Std dev | 10th perc | median | 90th perc | N |
| --- | --- | --- | --- | --- | --- | --- | --- |
| JFRC2010 | ANTs A | 11.570 | 7.574 | 2.345 | 10.464 | 22.113 | 176308 |
|  | ANTs B | 5.994 | 5.228 | 1.414 | 4.528 | 12.767 | 53780 |
|  | ANTs C | 6.006 | 5.241 | 1.414 | 4.528 | 12.669 | 48400 |
|  | CMTK A | 9.958 | 6.654 | 2.345 | 8.660 | 19.672 | 95702 |
|  | CMTK B | 10.801 | 7.191 | 2.345 | 9.539 | 21.378 | 84010 |
|  | CMTK C | 8.467 | 5.685 | 2.121 | 7.246 | 16.748 | 102936 |
|  | Elastix A | 6.023 | 5.398 | 1.414 | 4.528 | 12.689 | 48958 |
|  | Elastix B | 6.049 | 4.992 | 1.414 | 4.743 | 12.042 | 56630 |
| JFRC2013 | ANTs A | 4.988 | 5.337 | 1.000 | 3.162 | 11.790 | 47072 |
|  | ANTs B | 5.080 | 3.931 | 1.414 | 4.000 | 10.320 | 158968 |
|  | ANTs C | 5.849 | 4.347 | 1.581 | 4.690 | 11.705 | 130506 |
|  | CMTK A | 6.402 | 5.851 | 1.414 | 4.528 | 14.018 | 37398 |
|  | CMTK B | 6.525 | 5.589 | 1.581 | 4.950 | 13.435 | 41744 |
|  | CMTK C | 6.529 | 4.821 | 1.732 | 5.292 | 12.923 | 82804 |
|  | Elastix A | 5.209 | 5.190 | 1.000 | 3.606 | 11.597 | 51690 |
|  | Elastix B | 5.227 | 5.188 | 1.000 | 3.606 | 11.597 | 50770 |
| JRC2018 | ANTs A | 5.152 | 5.552 | 1.000 | 3.240 | 12.186 | 46336 |
|  | ANTs B | 5.764 | 4.816 | 1.225 | 4.528 | 11.604 | 53168 |
|  | ANTs C | 5.795 | 4.116 | 1.581 | 4.950 | 10.724 | 75474 |
|  | CMTK A | 5.810 | 5.894 | 1.000 | 3.808 | 13.730 | 33196 |
|  | CMTK B | 5.847 | 5.896 | 1.000 | 3.873 | 13.739 | 32616 |
|  | CMTK C | 5.934 | 5.925 | 1.225 | 4.062 | 13.647 | 35616 |
|  | Elastix A | 5.765 | 5.560 | 1.000 | 4.062 | 13.058 | 41278 |
|  | Elastix B | 5.981 | 5.689 | 1.225 | 4.123 | 13.675 | 37368 |
| FCWB | ANTs A | 6.636 | 5.688 | 1.225 | 4.796 | 14.916 | 50220 |
|  | ANTs B | 5.839 | 4.106 | 1.581 | 4.899 | 11.402 | 133492 |
|  | ANTs C | 6.722 | 4.678 | 1.732 | 5.701 | 13.285 | 105324 |
|  | CMTK A | 10.190 | 6.568 | 2.345 | 9.192 | 19.975 | 113524 |
|  | CMTK B | 9.805 | 6.403 | 2.345 | 8.602 | 18.947 | 71940 |
|  | CMTK C | 7.424 | 5.566 | 1.732 | 6.000 | 15.572 | 74432 |
|  | Elastix A | 9.369 | 6.192 | 2.236 | 8.246 | 18.453 | 186822 |
|  | Elastix B | 11.057 | 7.018 | 2.449 | 10.149 | 21.225 | 128624 |
| Tefor | ANTs A | 5.094 | 5.423 | 1.000 | 3.317 | 12.042 | 46480 |
|  | ANTs B | 5.715 | 4.603 | 1.414 | 4.528 | 11.790 | 46570 |
|  | ANTs C | 7.075 | 5.221 | 1.732 | 5.701 | 13.874 | 61358 |
|  | CMTK A | 10.621 | 6.530 | 2.550 | 9.849 | 19.698 | 80892 |
|  | CMTK B | 10.684 | 6.664 | 2.550 | 9.823 | 19.975 | 77374 |
|  | CMTK C | 10.544 | 7.218 | 2.236 | 9.301 | 20.881 | 81748 |
|  | Elastix A | 6.861 | 5.579 | 1.581 | 5.431 | 14.177 | 41516 |
|  | Elastix B | 8.750 | 5.938 | 2.121 | 7.416 | 17.507 | 103946 |

**Table S76:** SAD : saddle

| Template | Algorithm | Mean | Std dev | 10th perc | median | 90th perc | N |
| --- | --- | --- | --- | --- | --- | --- | --- |
| JFRC2010 | ANTs A | 5.974 | 5.347 | 1.581 | 4.528 | 11.619 | 815870 |
|  | ANTs B | 4.852 | 3.709 | 1.581 | 4.123 | 8.631 | 856040 |
|  | ANTs C | 4.952 | 3.894 | 1.581 | 4.183 | 8.860 | 901644 |
|  | CMTK A | 5.157 | 4.126 | 1.581 | 4.183 | 9.618 | 867552 |
|  | CMTK B | 5.005 | 3.960 | 1.581 | 4.123 | 9.165 | 846594 |
|  | CMTK C | 5.125 | 4.054 | 1.581 | 4.243 | 9.192 | 846840 |
|  | Elastix A | 4.538 | 3.565 | 1.414 | 3.808 | 8.185 | 750648 |
|  | Elastix B | 4.755 | 3.790 | 1.414 | 4.000 | 8.573 | 723218 |
| JFRC2013 | ANTs A | 4.608 | 3.762 | 1.225 | 3.808 | 8.515 | 796100 |
|  | ANTs B | 5.149 | 4.085 | 1.581 | 4.359 | 9.192 | 790572 |
|  | ANTs C | 5.098 | 4.178 | 1.581 | 4.243 | 9.192 | 820998 |
|  | CMTK A | 4.904 | 3.861 | 1.414 | 4.123 | 9.055 | 746112 |
|  | CMTK B | 4.907 | 3.813 | 1.414 | 4.123 | 9.000 | 741396 |
|  | CMTK C | 5.122 | 3.787 | 1.581 | 4.359 | 9.247 | 766278 |
|  | Elastix A | 4.569 | 3.627 | 1.225 | 3.808 | 8.307 | 717052 |
|  | Elastix B | 4.614 | 3.717 | 1.225 | 3.808 | 8.396 | 734362 |
| JRC2018 | ANTs A | 4.567 | 3.849 | 1.225 | 3.808 | 8.276 | 757694 |
|  | ANTs B | 4.892 | 3.770 | 1.581 | 4.123 | 8.775 | 783234 |
|  | ANTs C | 5.052 | 3.894 | 1.581 | 4.243 | 9.083 | 815800 |
|  | CMTK A | 4.598 | 3.879 | 1.225 | 3.808 | 8.396 | 751242 |
|  | CMTK B | 4.568 | 3.770 | 1.225 | 3.808 | 8.367 | 739994 |
|  | CMTK C | 4.631 | 3.699 | 1.414 | 3.808 | 8.396 | 733568 |
|  | Elastix A | 4.553 | 3.665 | 1.225 | 3.808 | 8.276 | 720306 |
|  | Elastix B | 4.719 | 3.895 | 1.414 | 3.873 | 8.544 | 725992 |
| FCWB | ANTs A | 4.685 | 3.689 | 1.225 | 3.873 | 8.544 | 751334 |
|  | ANTs B | 4.765 | 3.932 | 1.414 | 4.000 | 8.544 | 846026 |
|  | ANTs C | 4.749 | 3.804 | 1.414 | 4.000 | 8.515 | 855024 |
|  | CMTK A | 5.086 | 3.868 | 1.581 | 4.301 | 9.165 | 831892 |
|  | CMTK B | 4.993 | 3.845 | 1.581 | 4.243 | 8.972 | 819294 |
|  | CMTK C | 4.922 | 3.642 | 1.581 | 4.183 | 8.746 | 781292 |
|  | Elastix A | 5.654 | 5.084 | 1.581 | 4.583 | 10.100 | 802456 |
|  | Elastix B | 5.328 | 4.575 | 1.581 | 4.472 | 9.618 | 872146 |
| Tefor | ANTs A | 4.574 | 3.664 | 1.225 | 3.808 | 8.276 | 756640 |
|  | ANTs B | 5.023 | 3.748 | 1.581 | 4.301 | 9.000 | 811812 |
|  | ANTs C | 5.192 | 3.922 | 1.581 | 4.359 | 9.407 | 828668 |
|  | CMTK A | 5.294 | 4.177 | 1.414 | 4.359 | 10.075 | 769982 |
|  | CMTK B | 5.270 | 4.061 | 1.414 | 4.359 | 10.075 | 761656 |
|  | CMTK C | 5.243 | 4.073 | 1.581 | 4.359 | 9.460 | 783740 |
|  | Elastix A | 4.763 | 3.676 | 1.414 | 4.062 | 8.544 | 710380 |
|  | Elastix B | 5.181 | 3.827 | 1.581 | 4.472 | 9.407 | 750662 |

## C. Qualitative registration results

Finally, we present a more thorough qualitative comparison of registration results. We first randomly selected two of our evaluation images, then display three z-slices of those nc82 images after registration to a particular template, using a particular algorithm. This is repeated for every template-algorithm pair, with the same two moving images. The z-slices were chosen to show similar anatomical structures, and therefore vary across templates.

Arrows were placed once on each template image at similar (but not identical) locations, and are displayed at the same location in each warped image. Similar anatomical structure around each arrow for both images indicates that registration is consistent from image to image. These anatomical structures include the: mushroom body, ellipsoid body, lateral horn, protocerebral bridge, and the anterior optic tubercle. Large differences in structure at the same location in the two aligned images is an indication of poor registration accuracy, but good apparent alignment could still result from inadequate registration. See the discussion limitations of using manually chosen landmark points in Section 3.2.1 and Section 4.1.

Despite the limitations of this qualitative assessment, we will briefly point out some observations given the examples below. For the CMTK registrations using JRC 2018F, the ellipsoid body appears overly deformed using CMTK A (Figure 18) and CMTK C (Figure 19), but not CMTK B (Figure 19). This is consistent with the JSD. Some registrations produced unusable results, with significant scaling issues, see Tefor-CMTK B (Figure 51). Over-warping is evident using ANTs C, for example, with JFRC 2013 (Figure 33). Surprisingly, some elastix transformations over-warp as well despite generally low-values of JSD, for example JFRC 2010 and Elastix A (Figure 45). It could be that elastix does not over-warp on average, but happened to produce a poor result for one of the images randomly selected here.

In general, there exists an algorithm that works well for every template. Overall, registration quality seems most consistent when using the JRC 2018 template.

**Figure 15:**
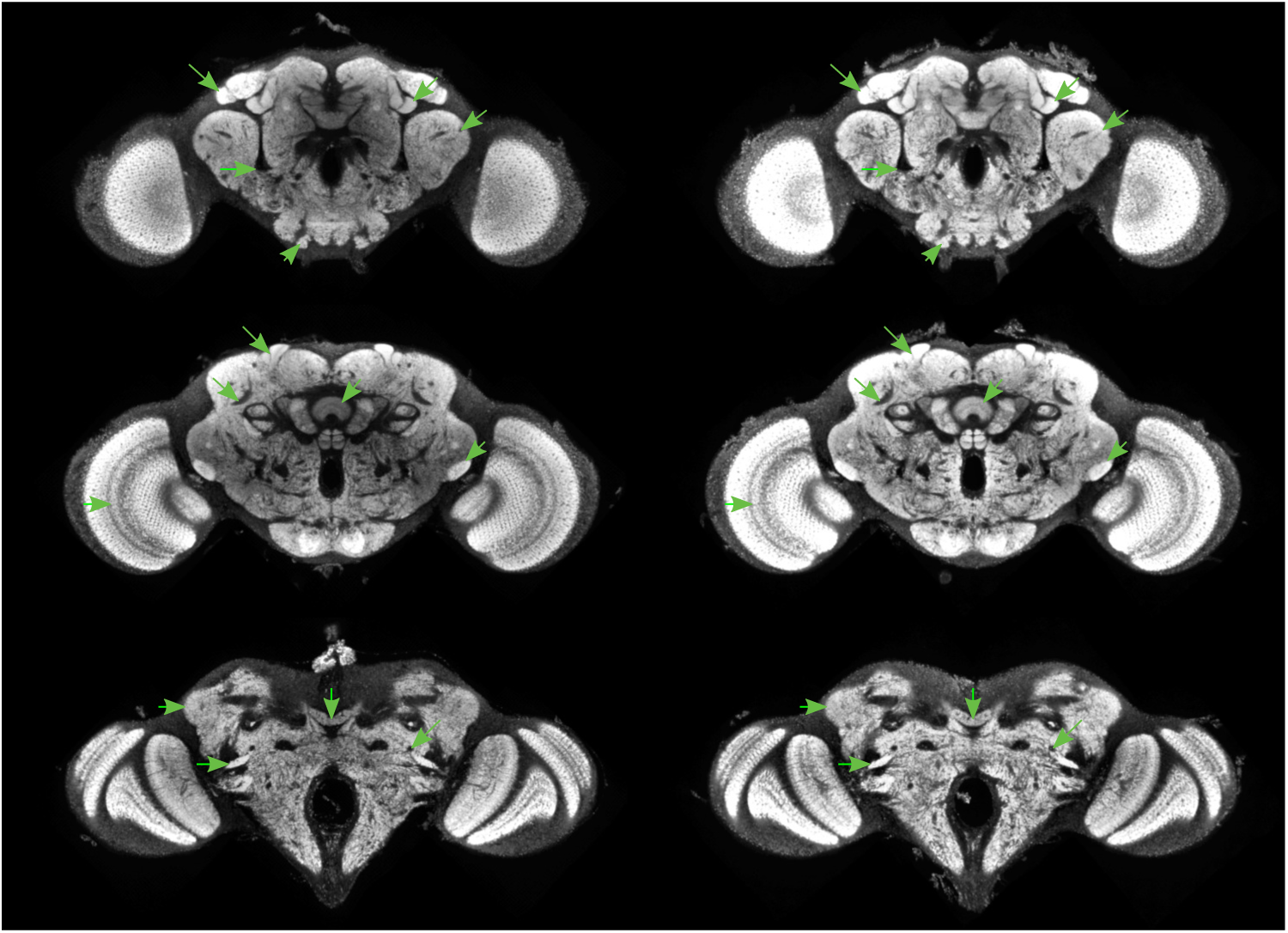
JRC2018 antsA

**Figure 16:**
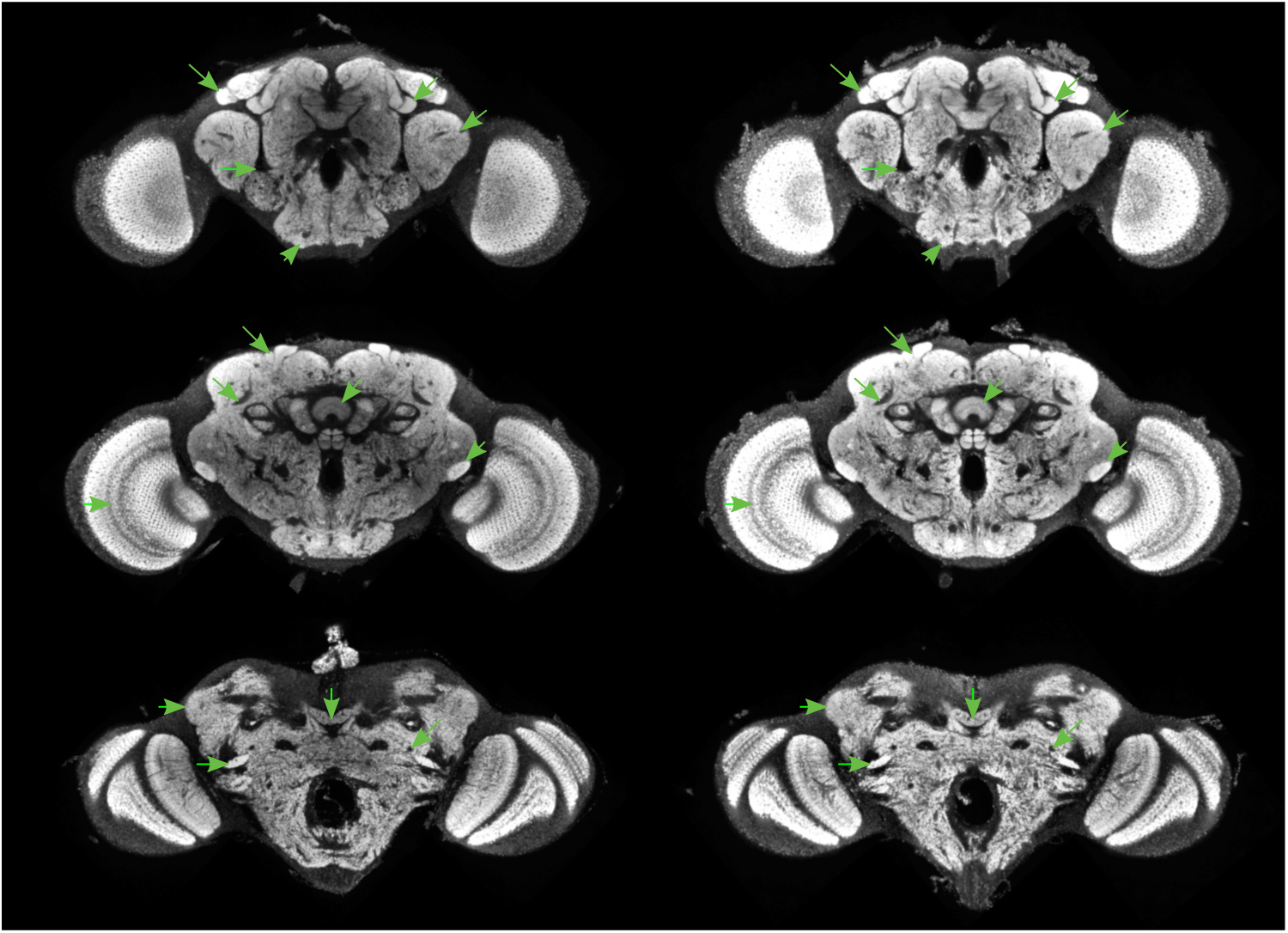
JRC2018 antsB

**Figure 17:**
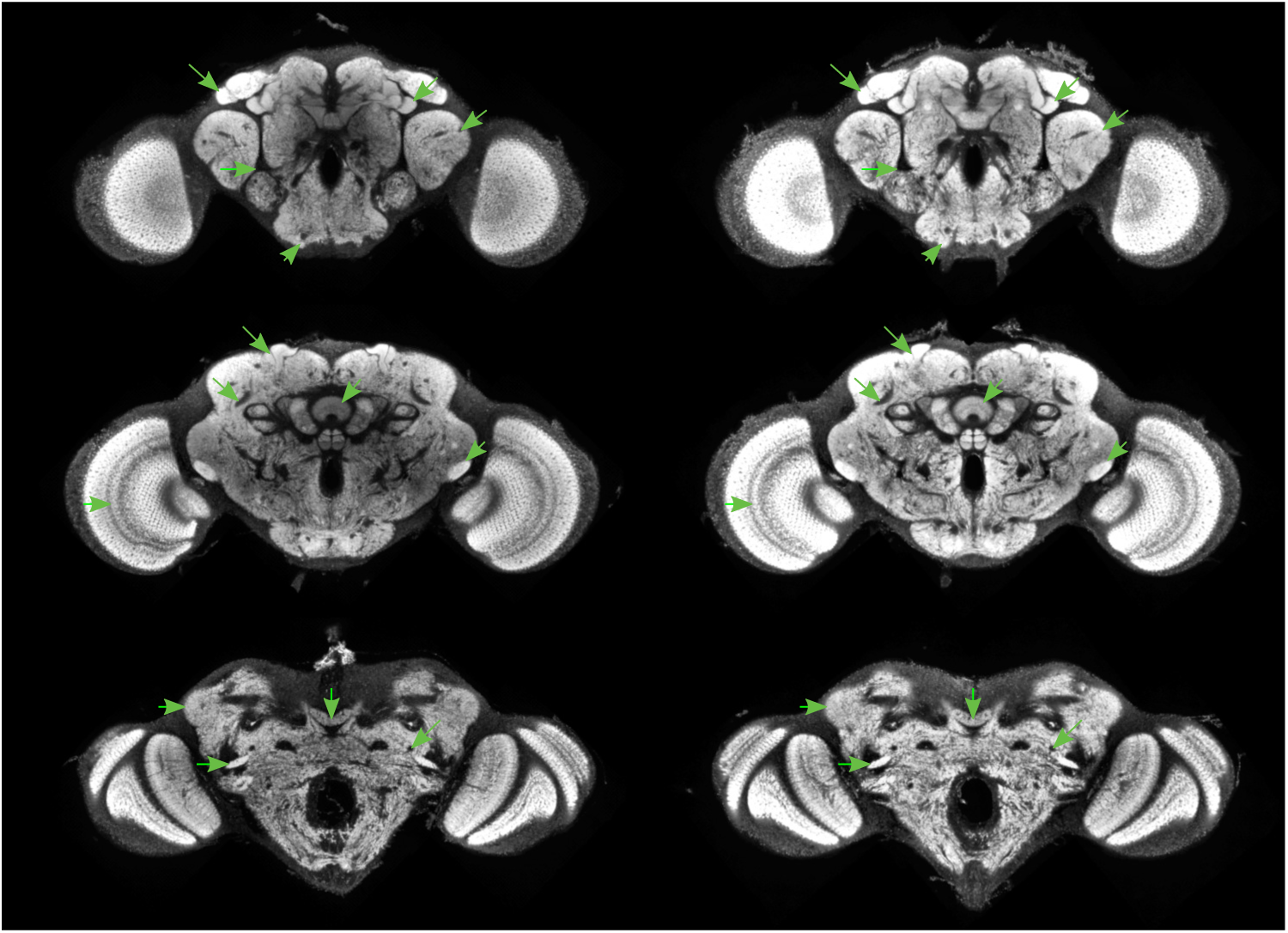
JRC2018 antsC

**Figure 18:**
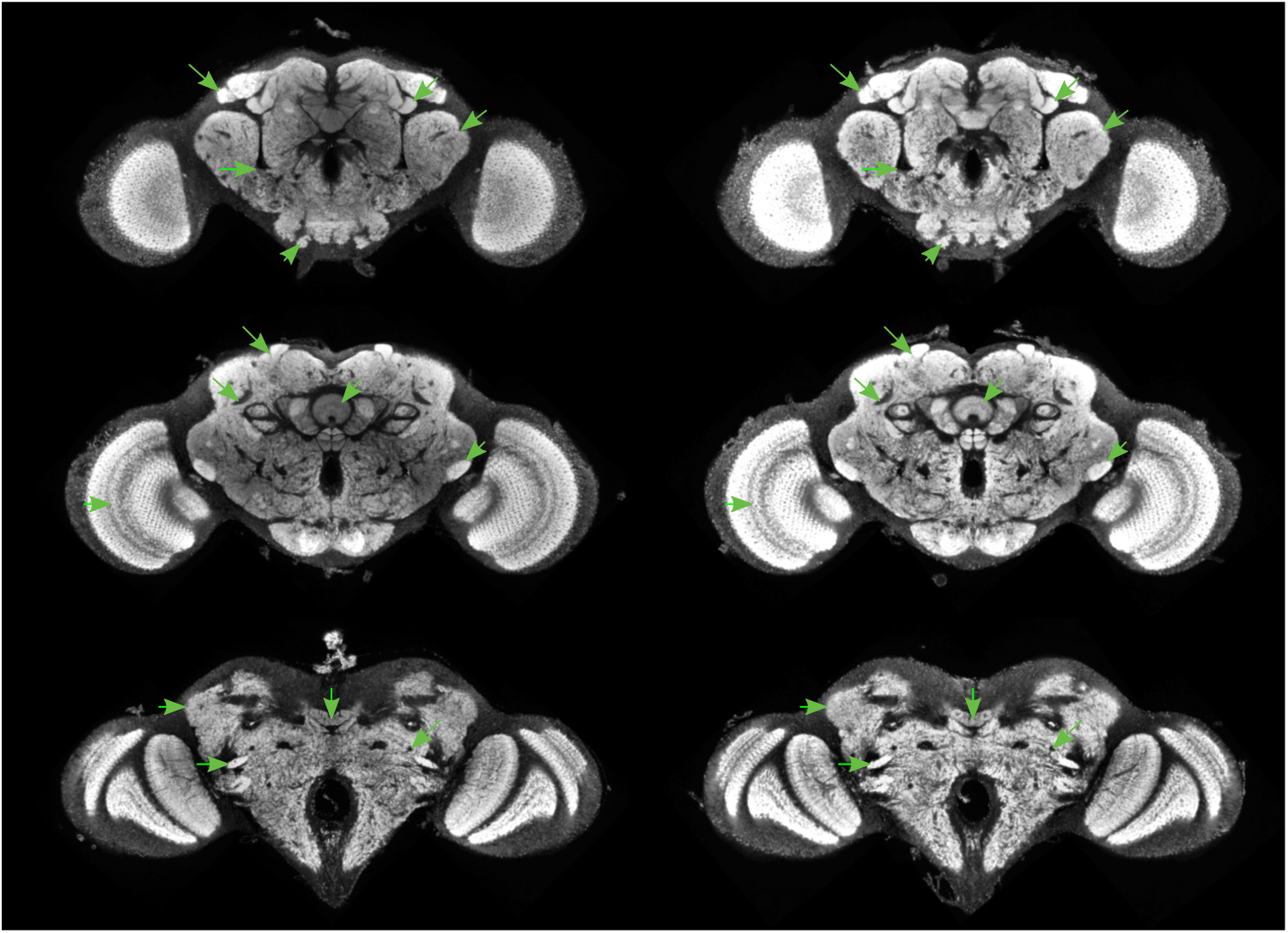
JRC2018 cmtkA

**Figure 19:**
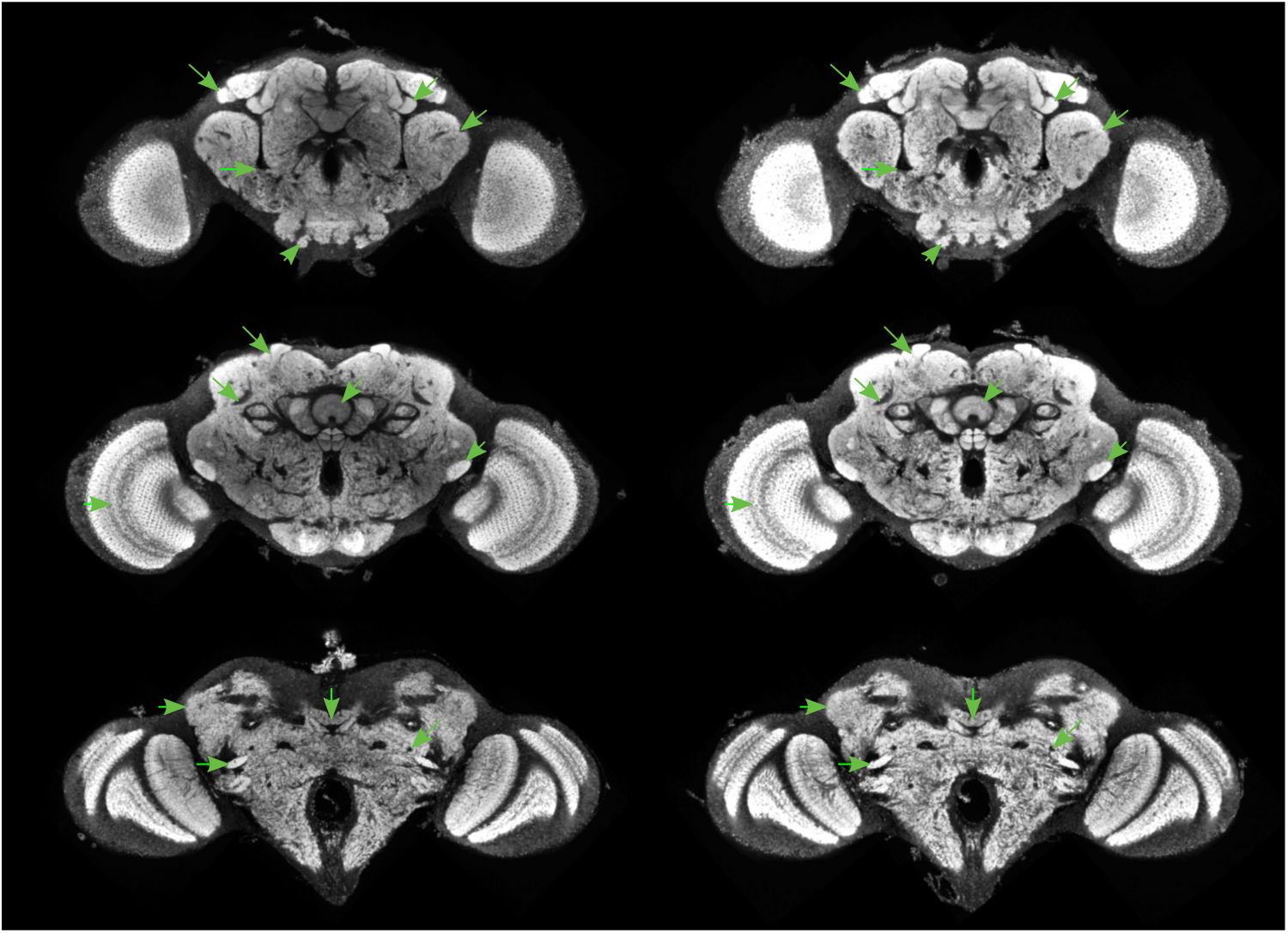
JRC2018 cmtkB

**Figure 20:**
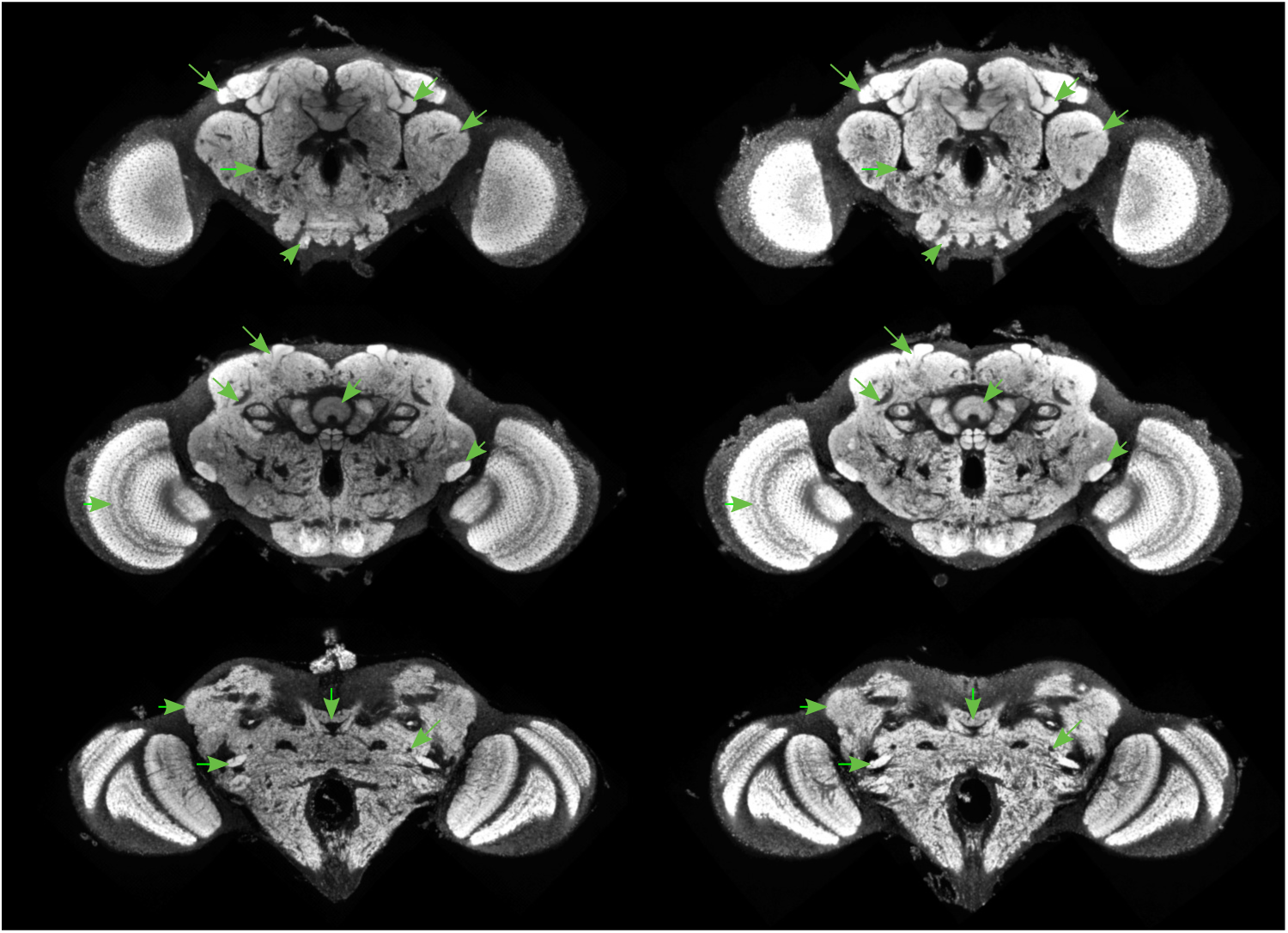
JRC2018 cmtkC

**Figure 21:**
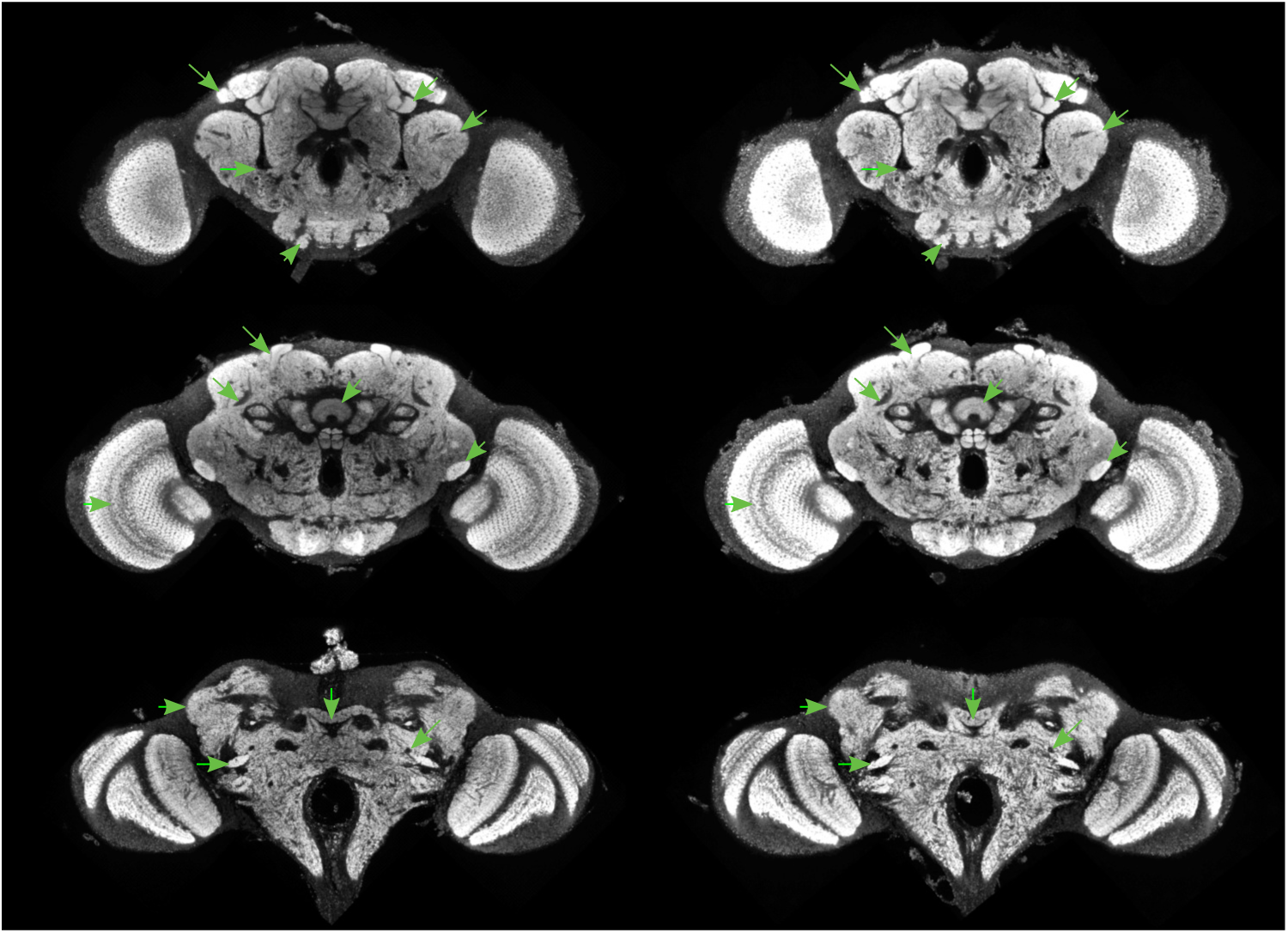
JRC2018 elastixA

**Figure 22:**
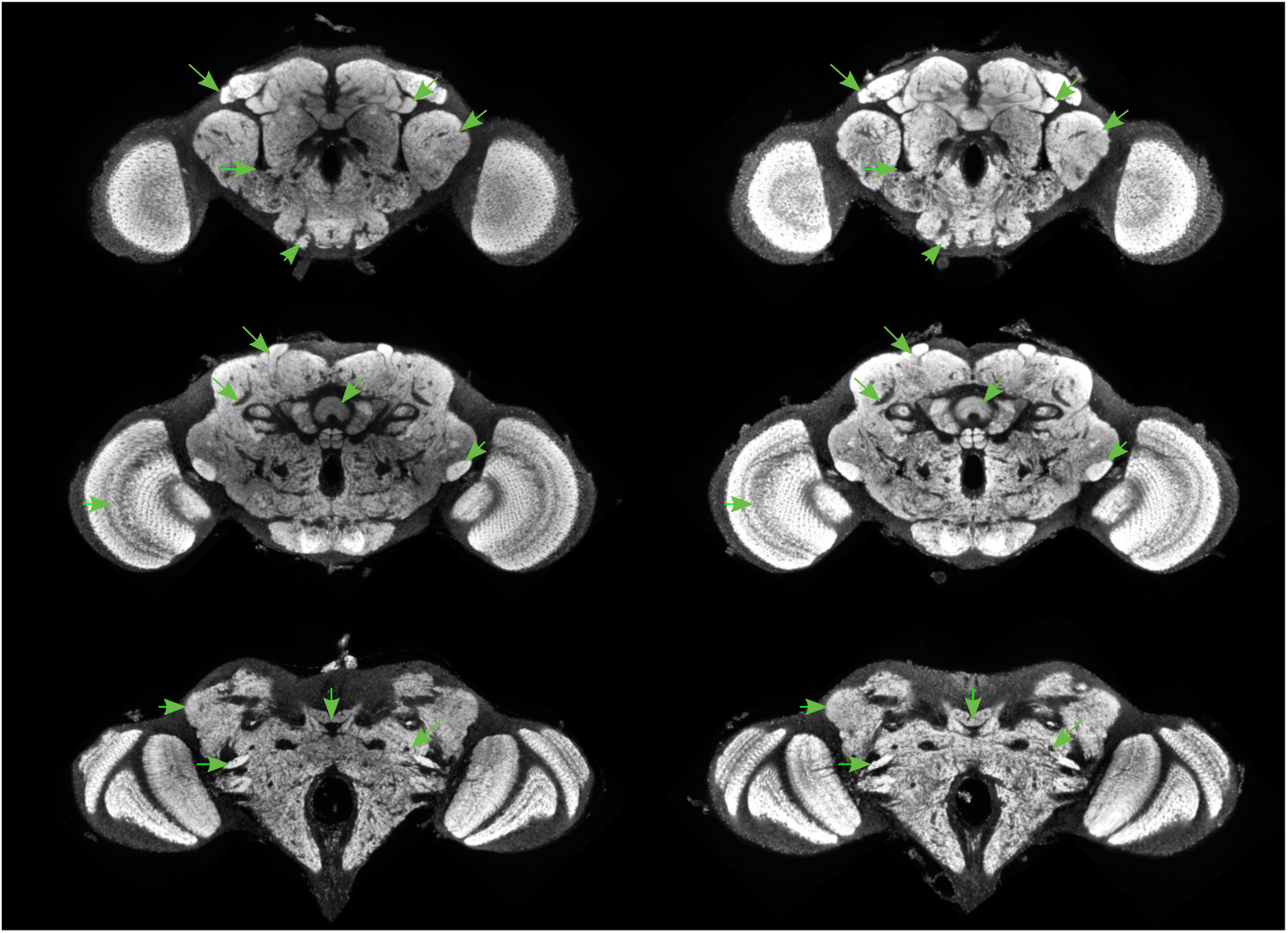
JRC2018 elastixB

**Figure 23:**
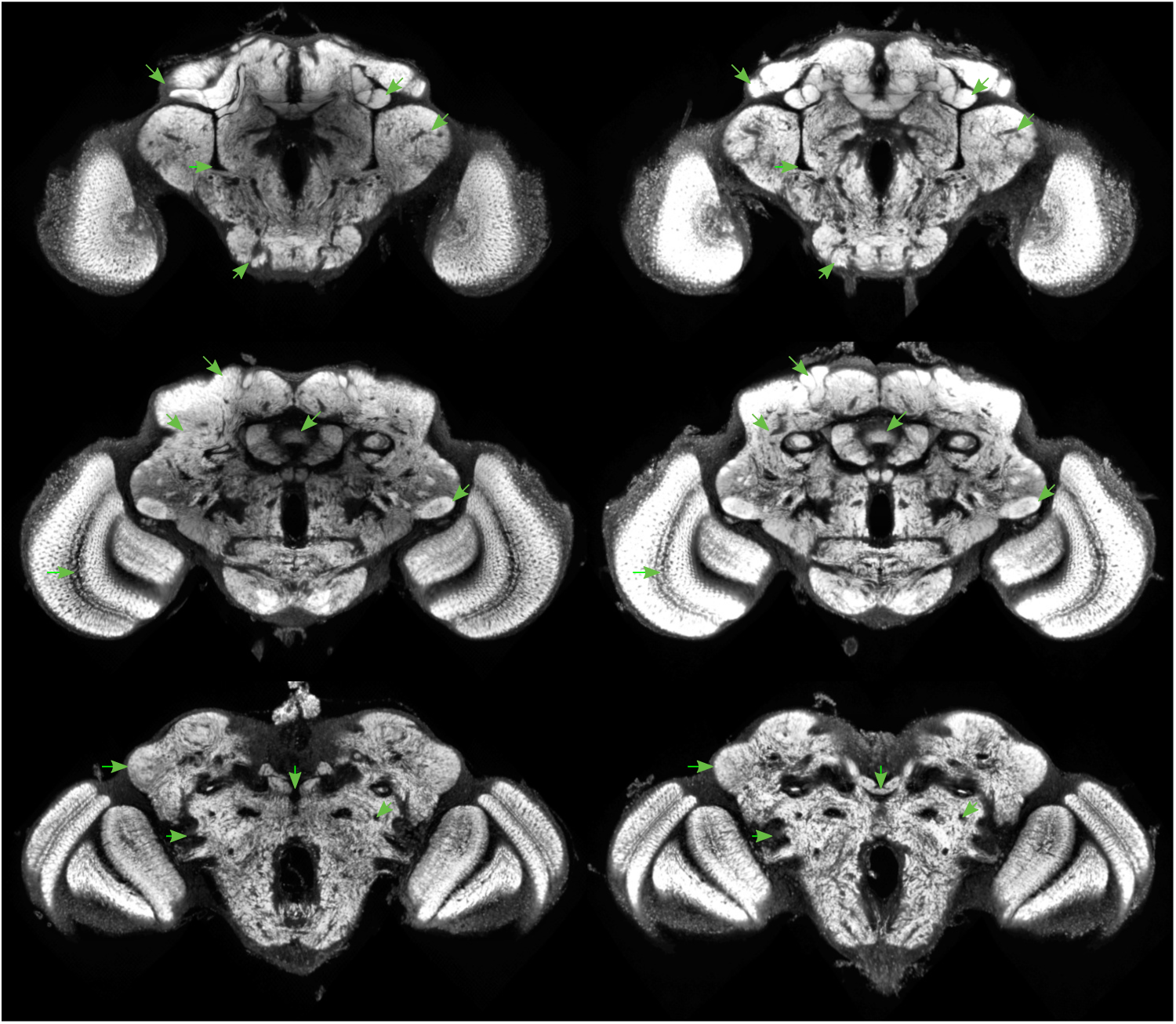
FCWB antsA

**Figure 24:**
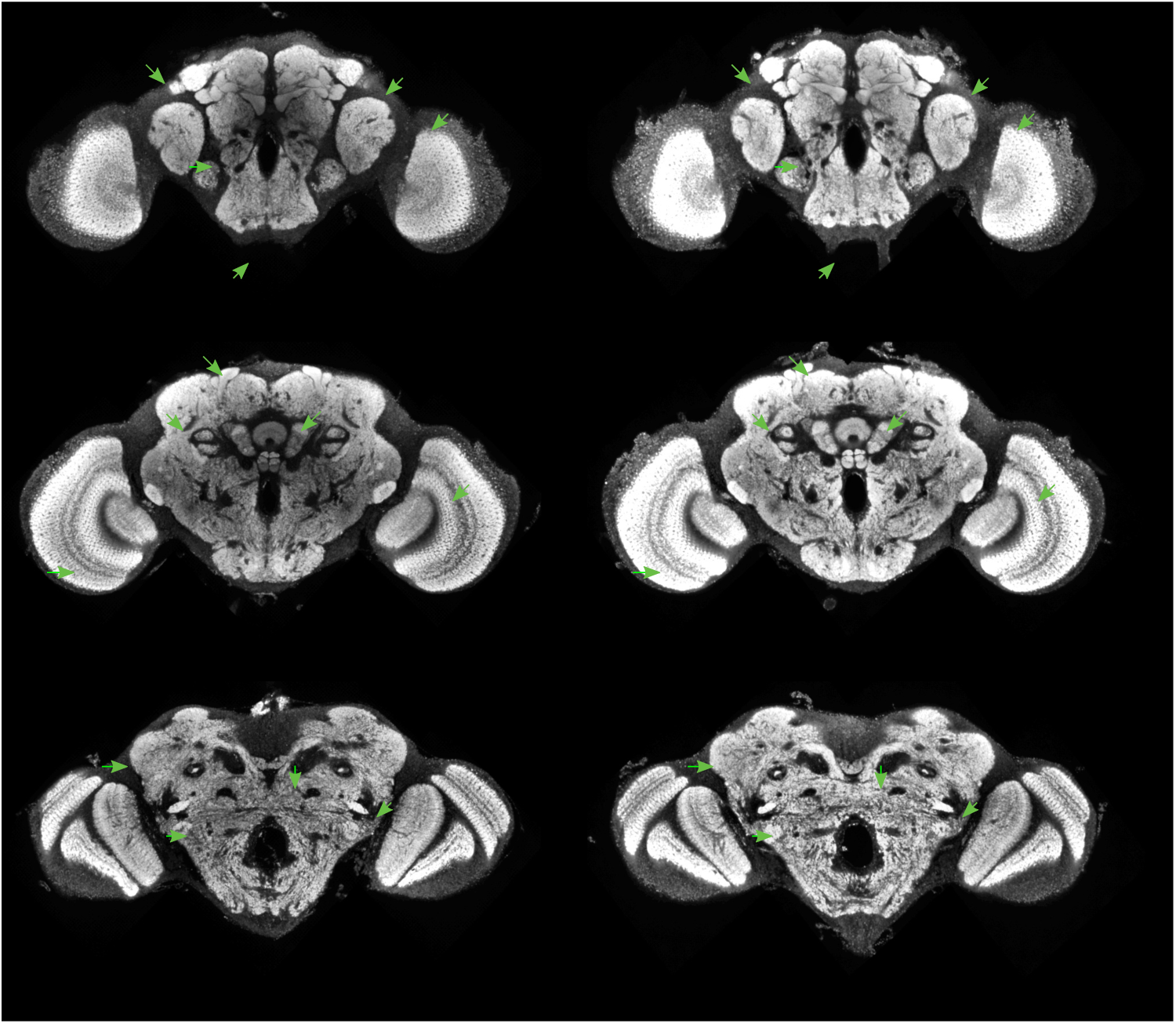
FCWB antsB

**Figure 25:**
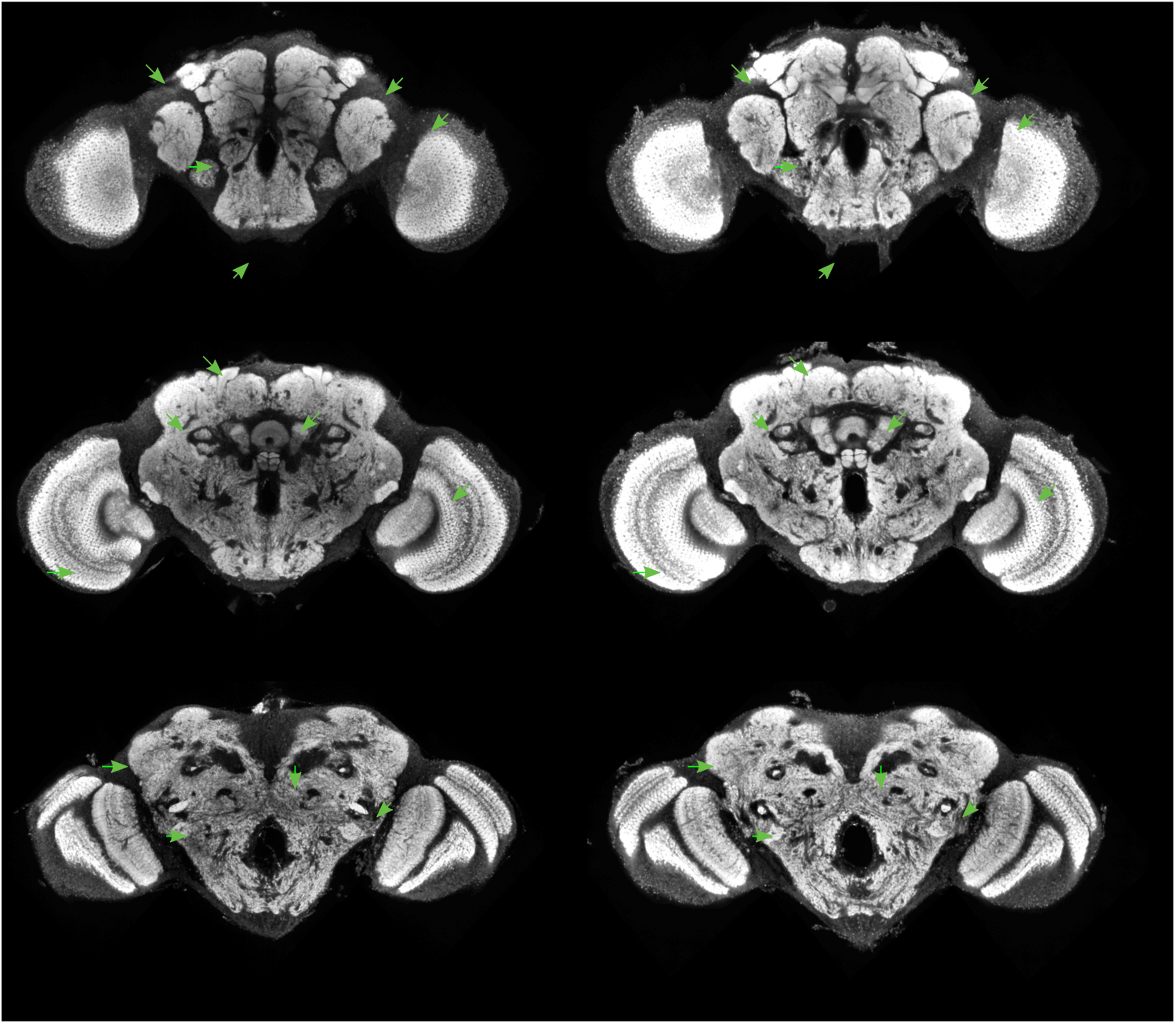
FCWB antsC

**Figure 26:**
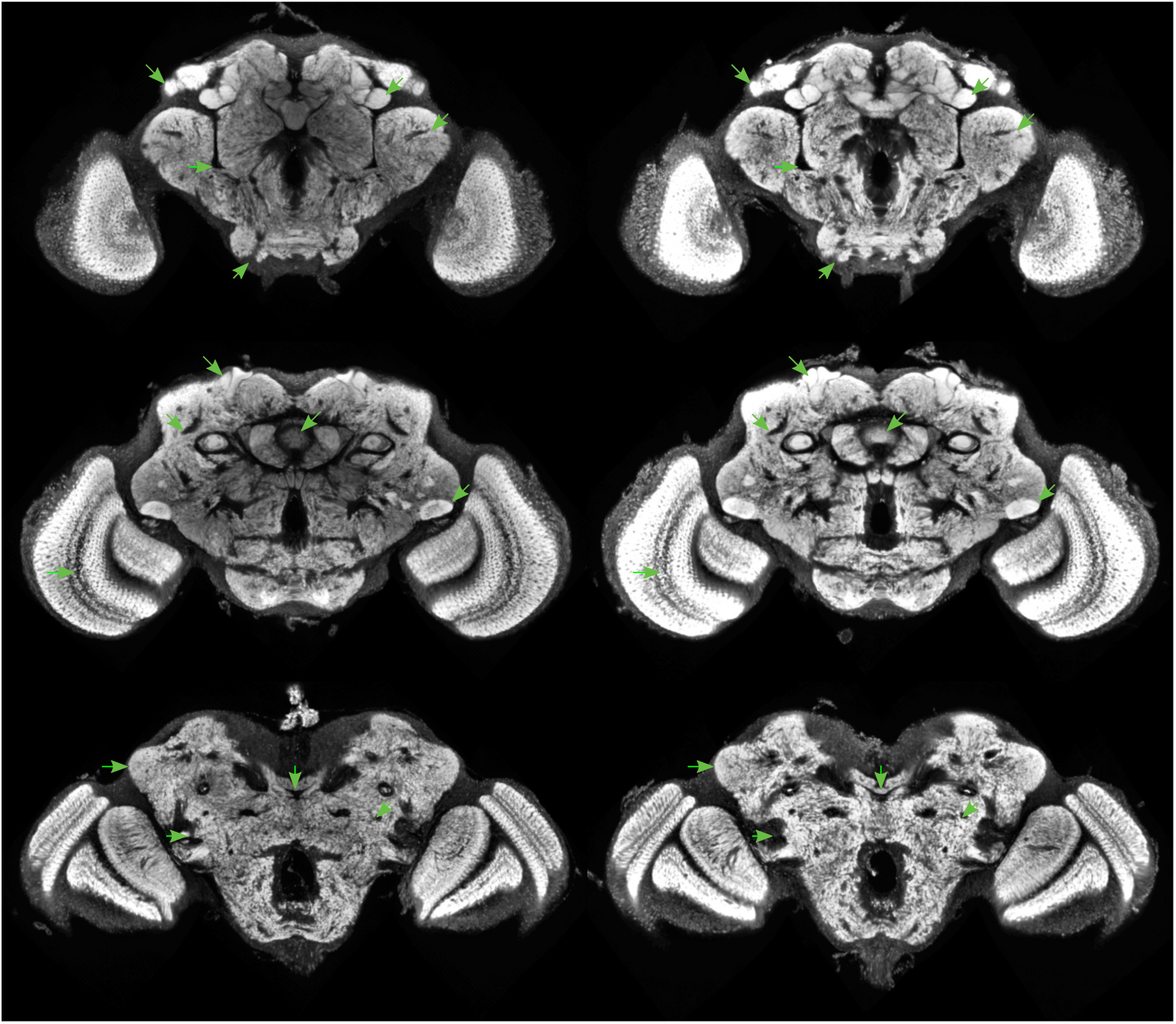
FCWB cmtkA

**Figure 27:**
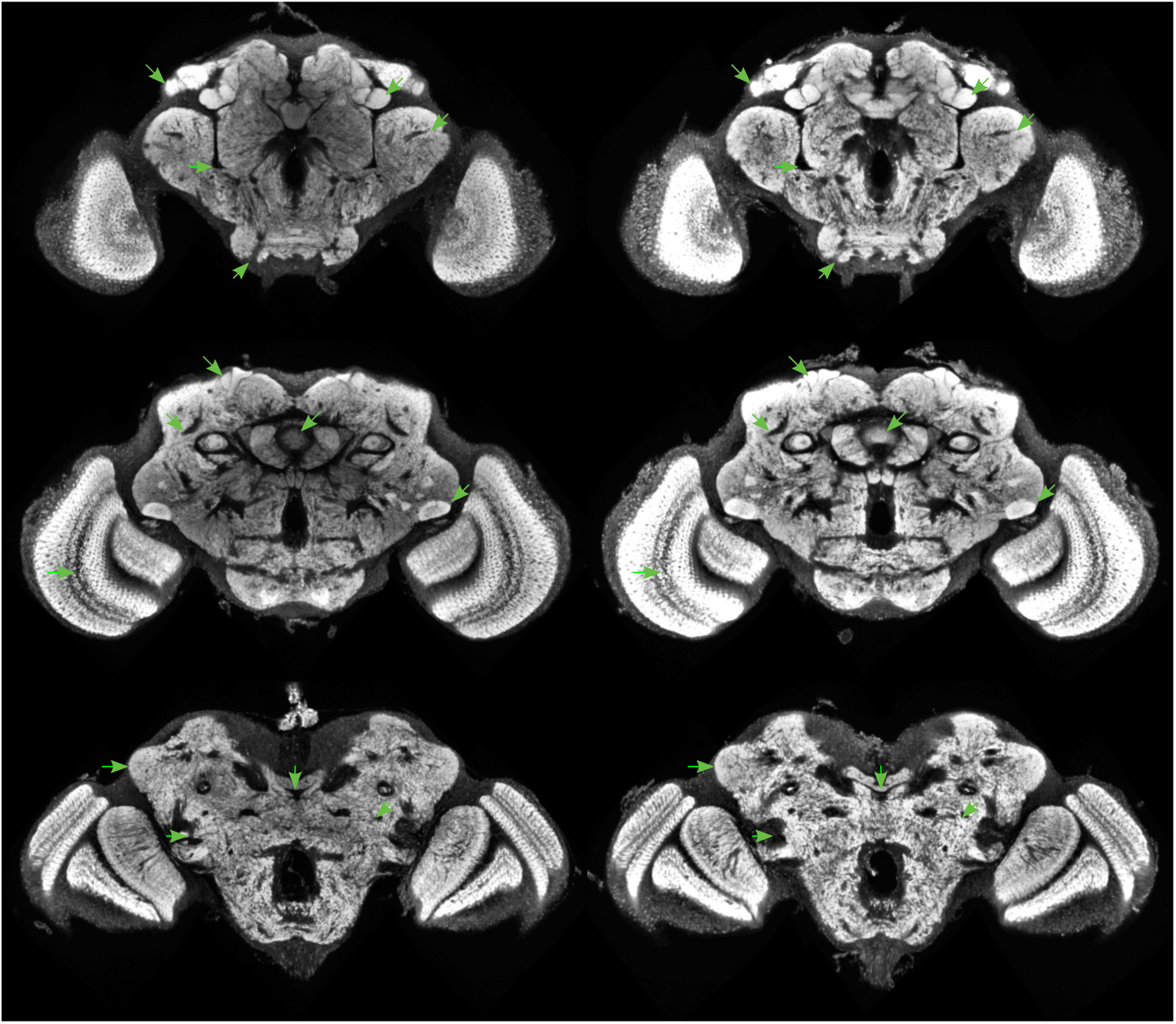
FCWB cmtkB

**Figure 28:**
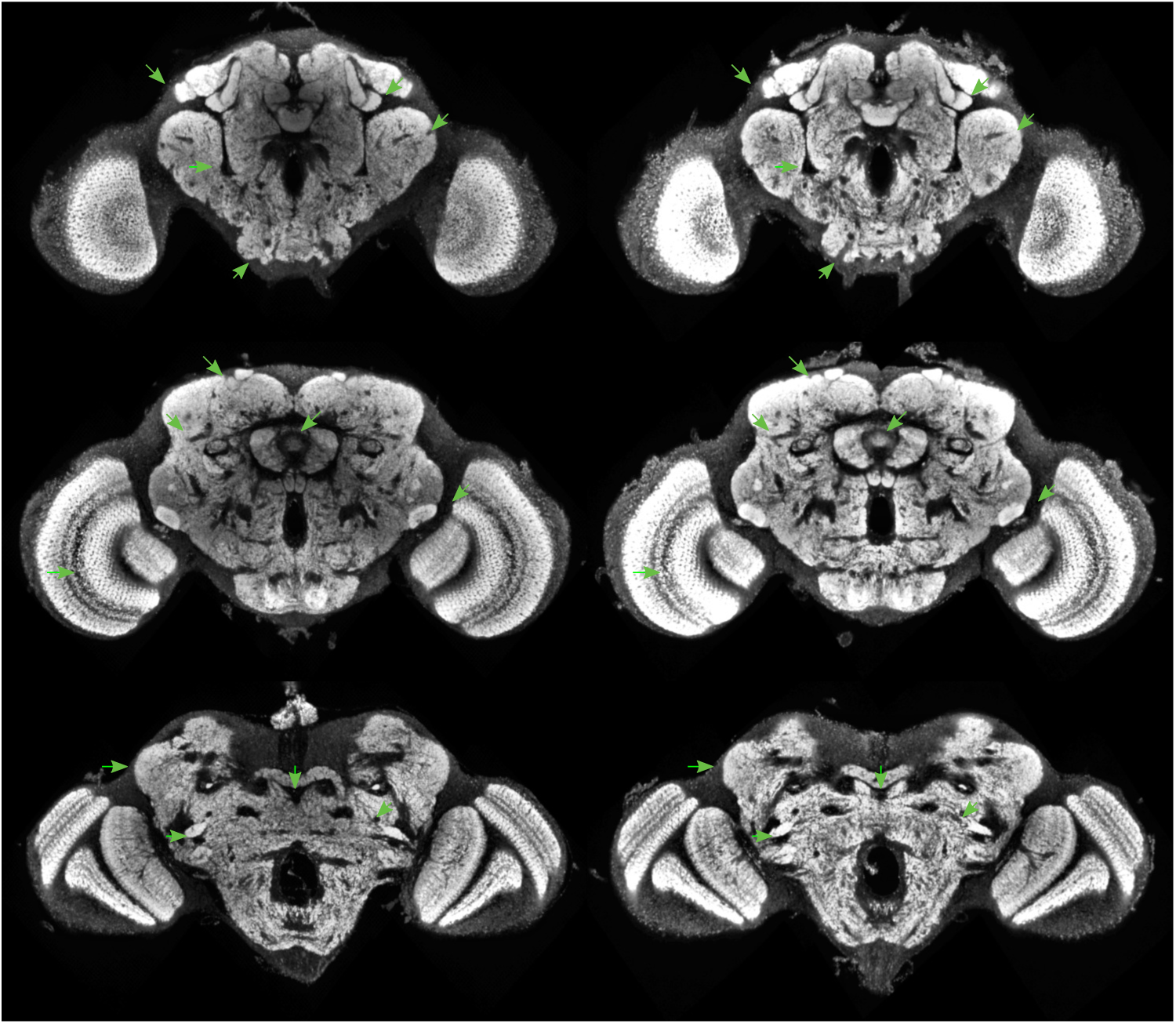
FCWB cmtkC

**Figure 29:**
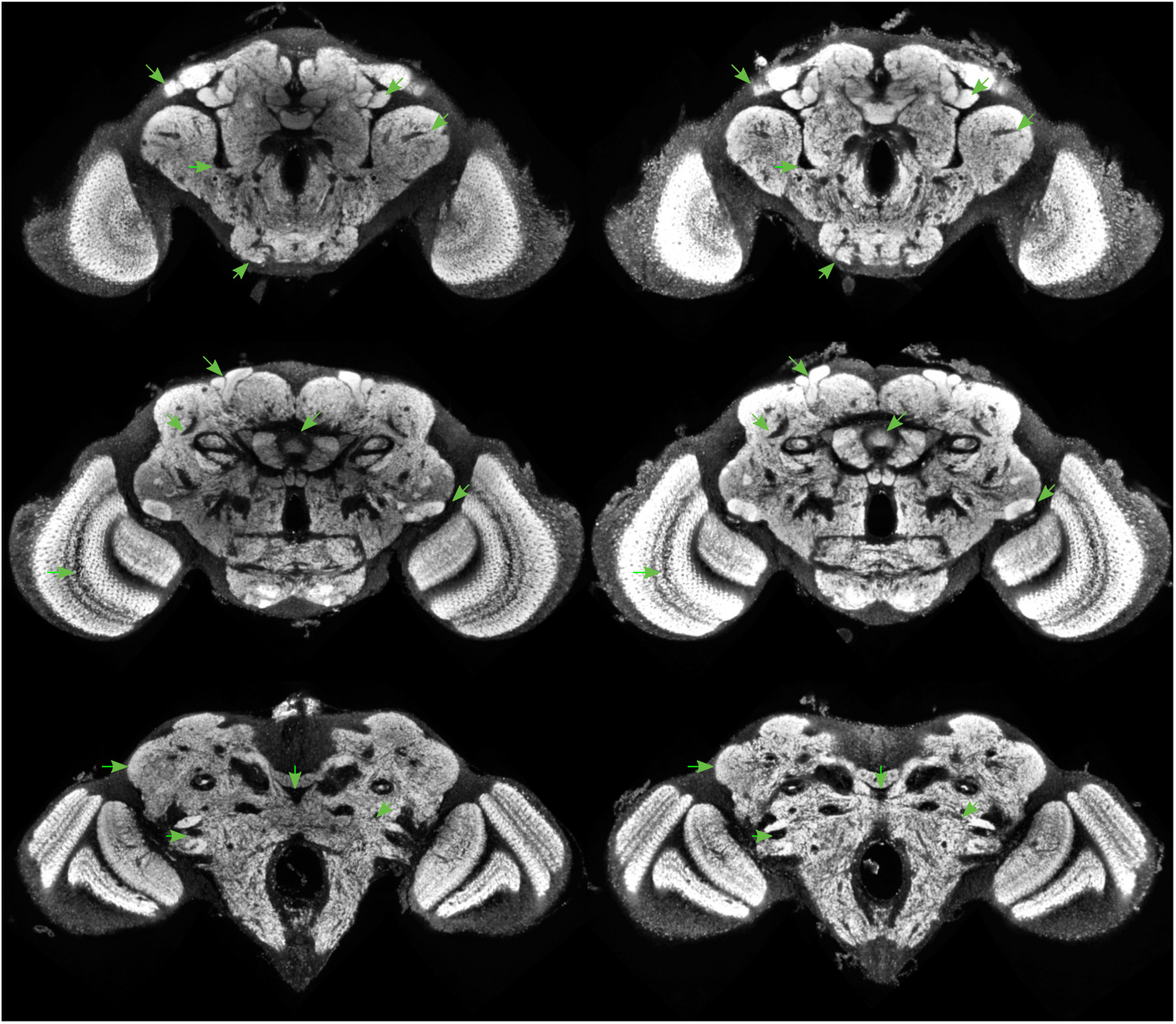
FCWB elastixA

**Figure 30:**
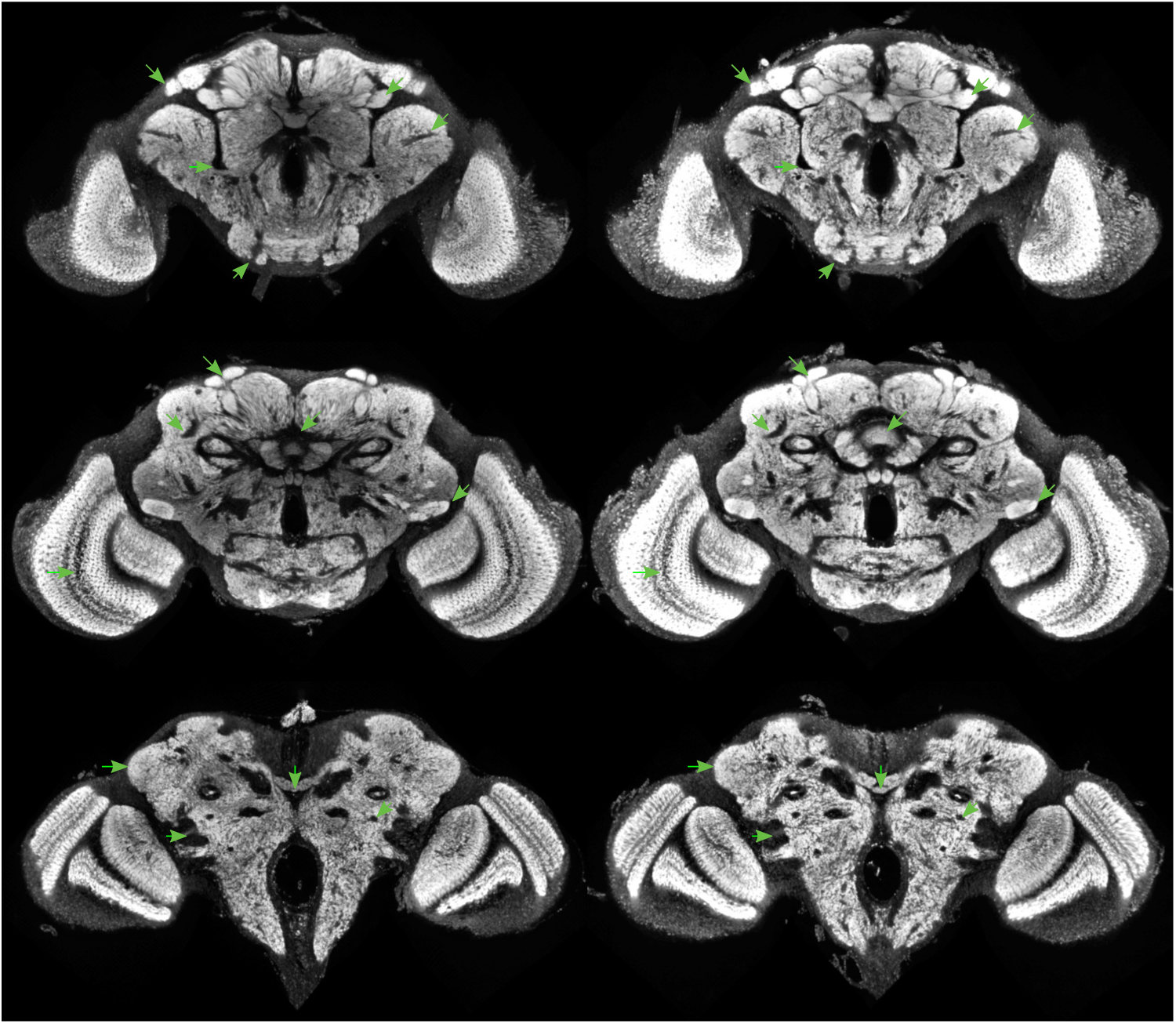
FCWB elastixB

**Figure 31:**
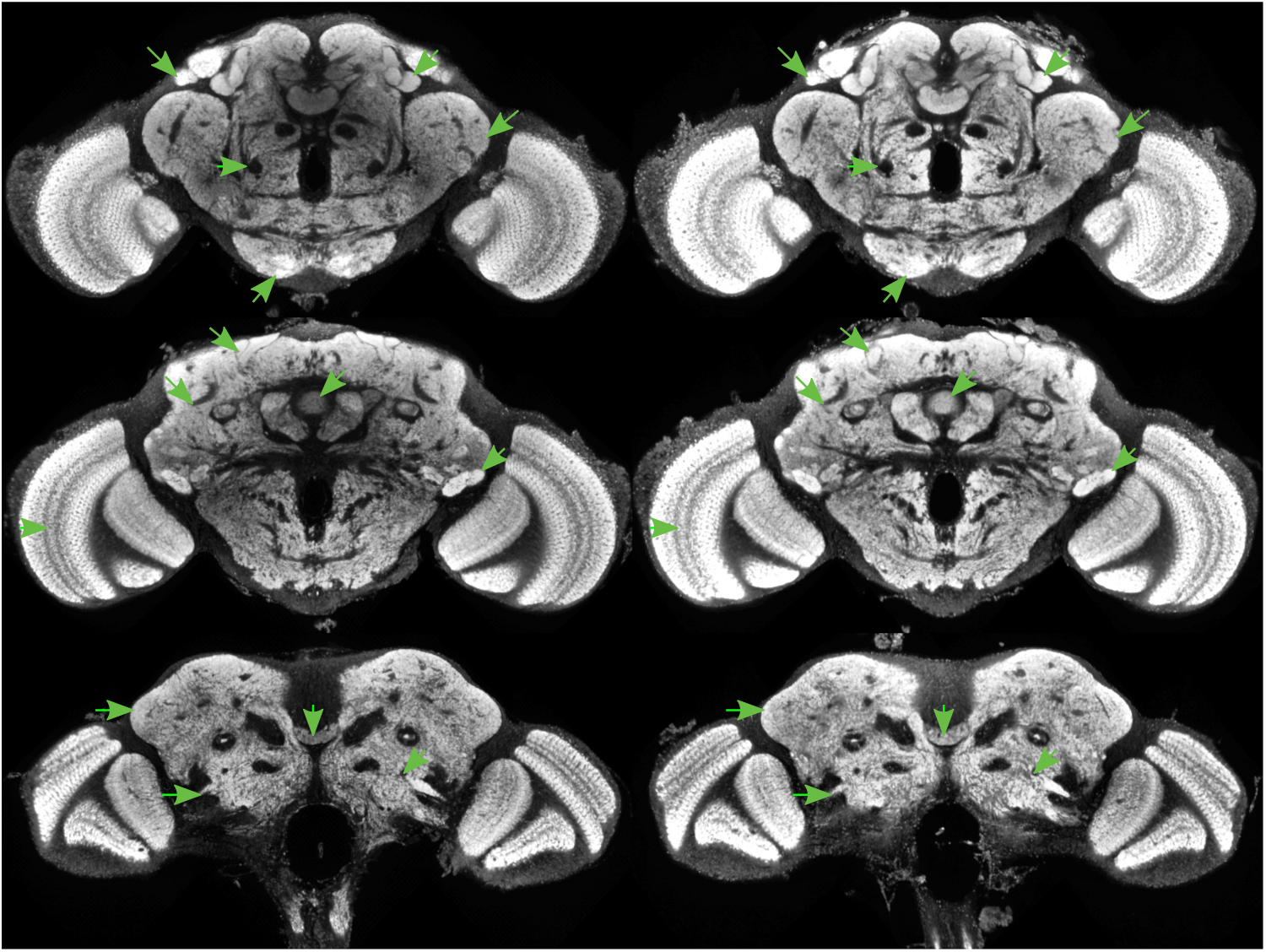
JFRC2013 antsA

**Figure 32:**
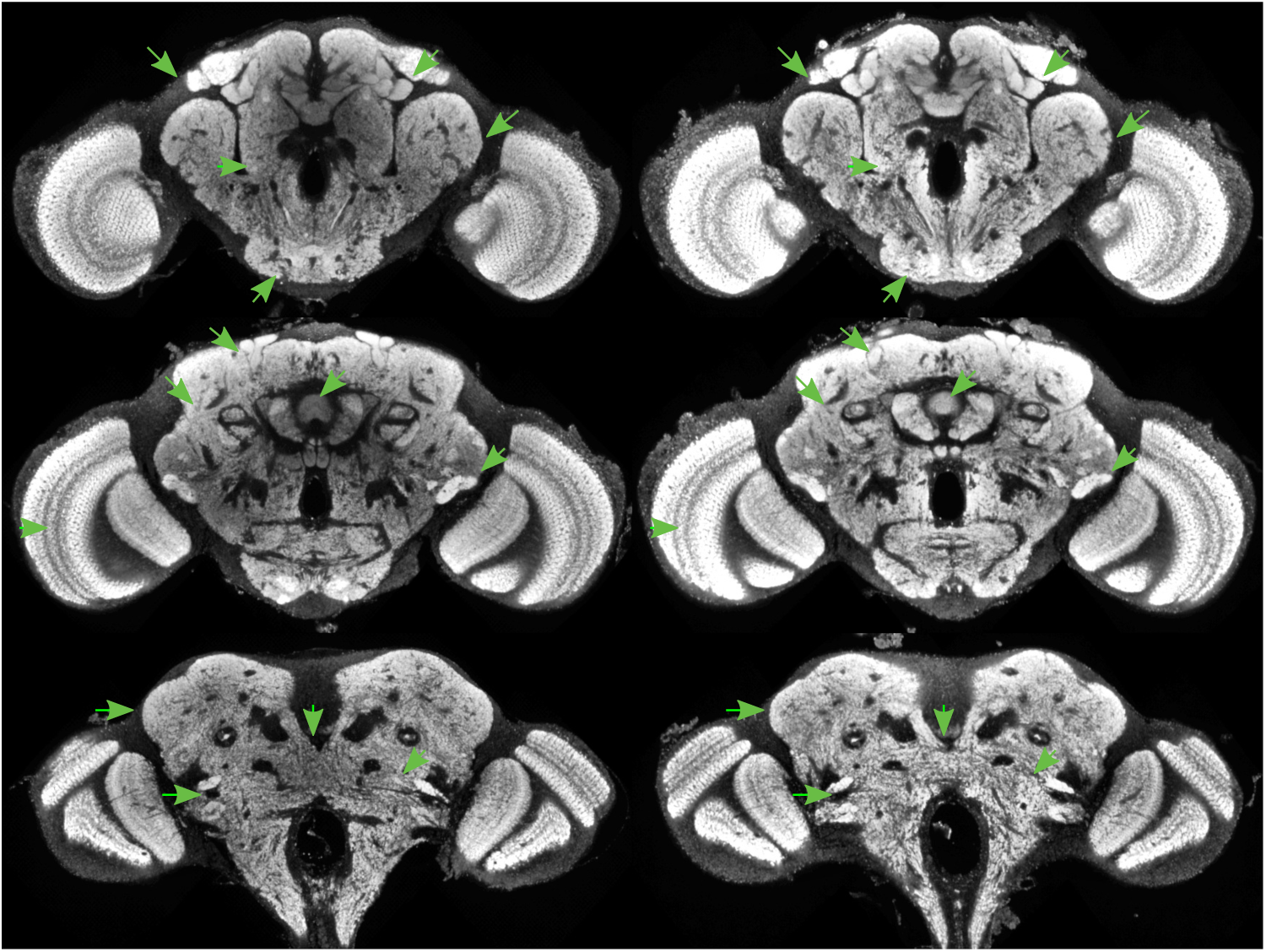
JFRC2013 antsB

**Figure 33:**
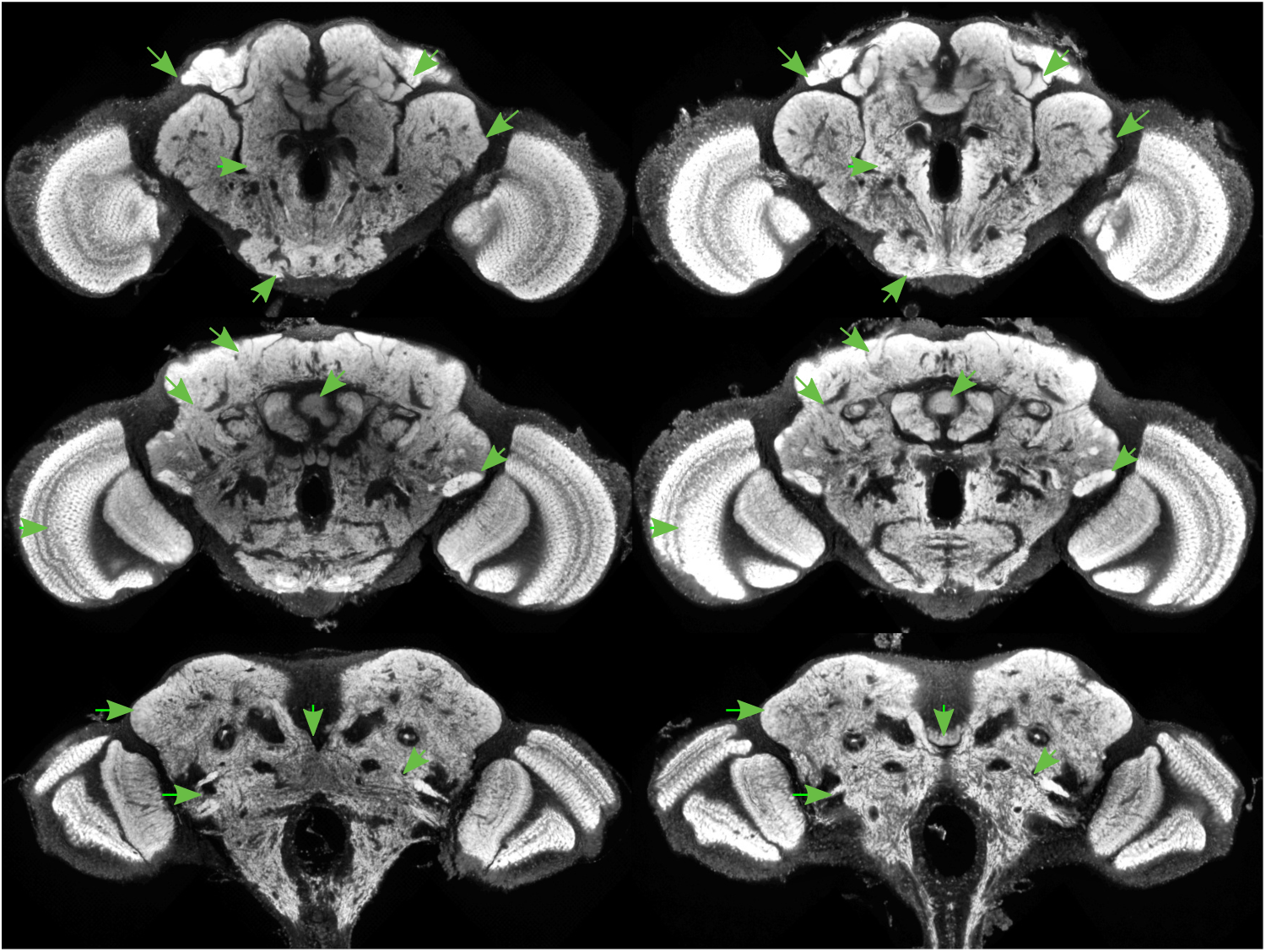
JFRC2013 antsC

**Figure 34:**
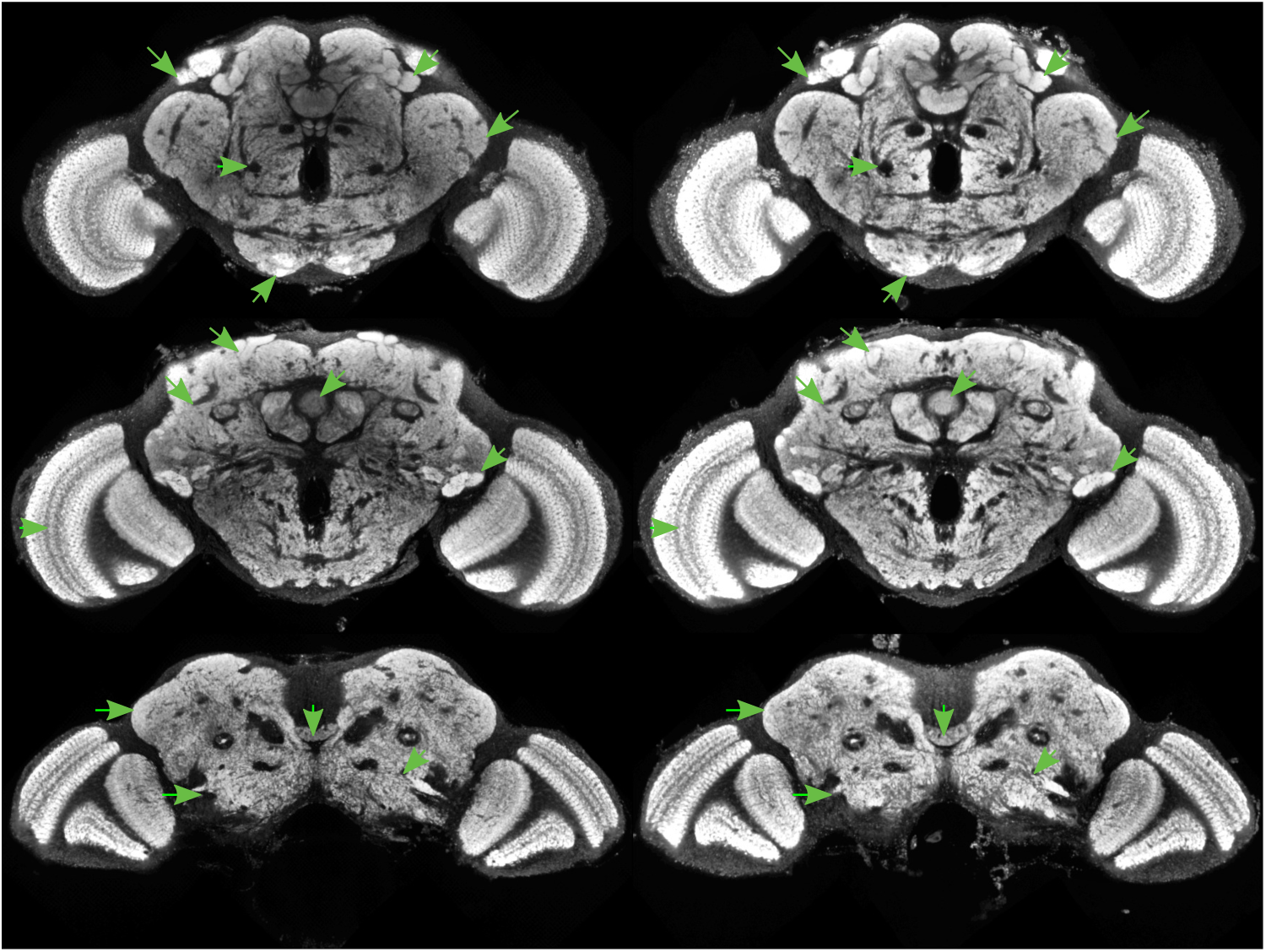
JFRC2013 cmtkA

**Figure 35:**
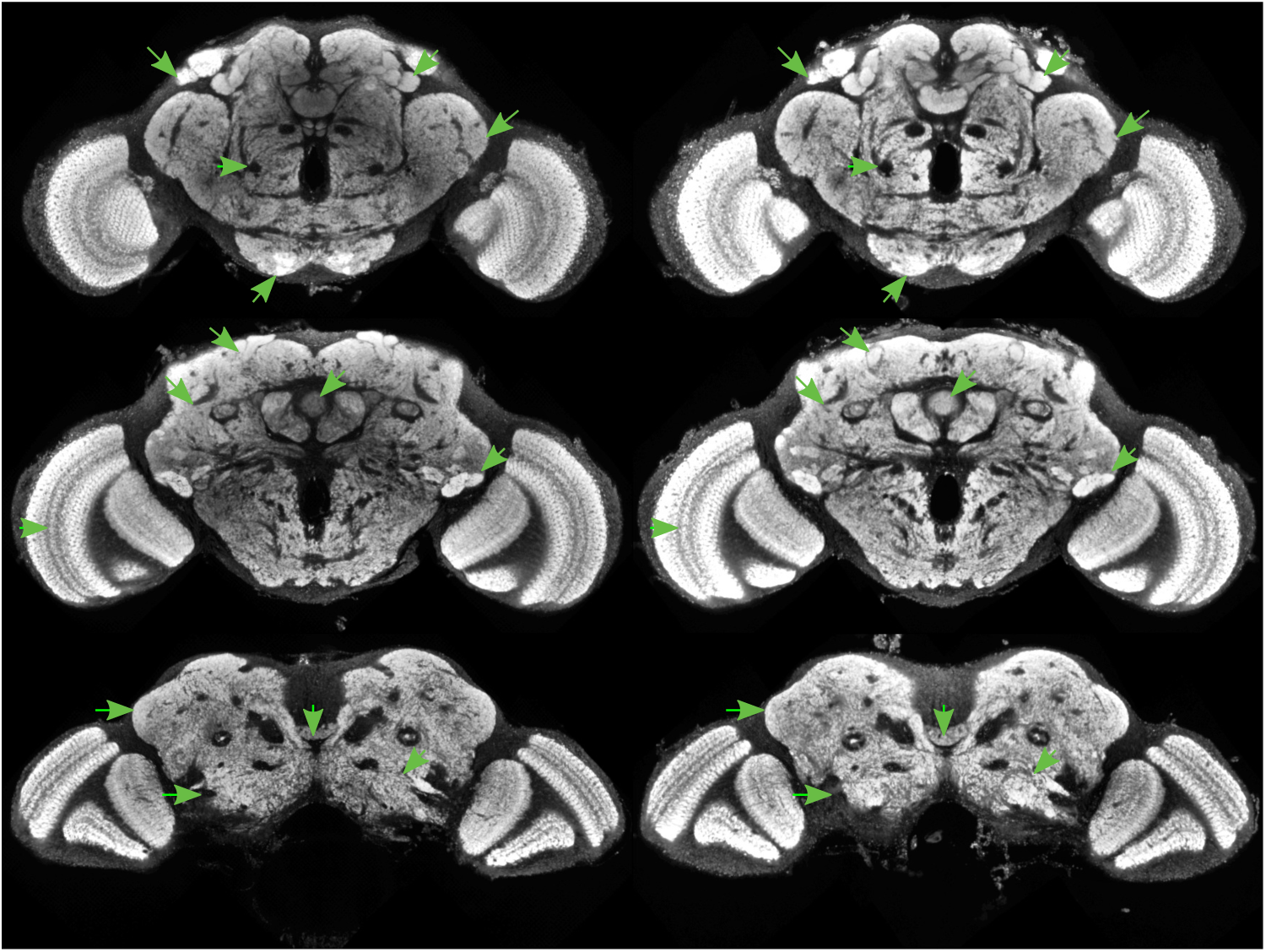
JFRC2013 cmtkB

**Figure 36:**
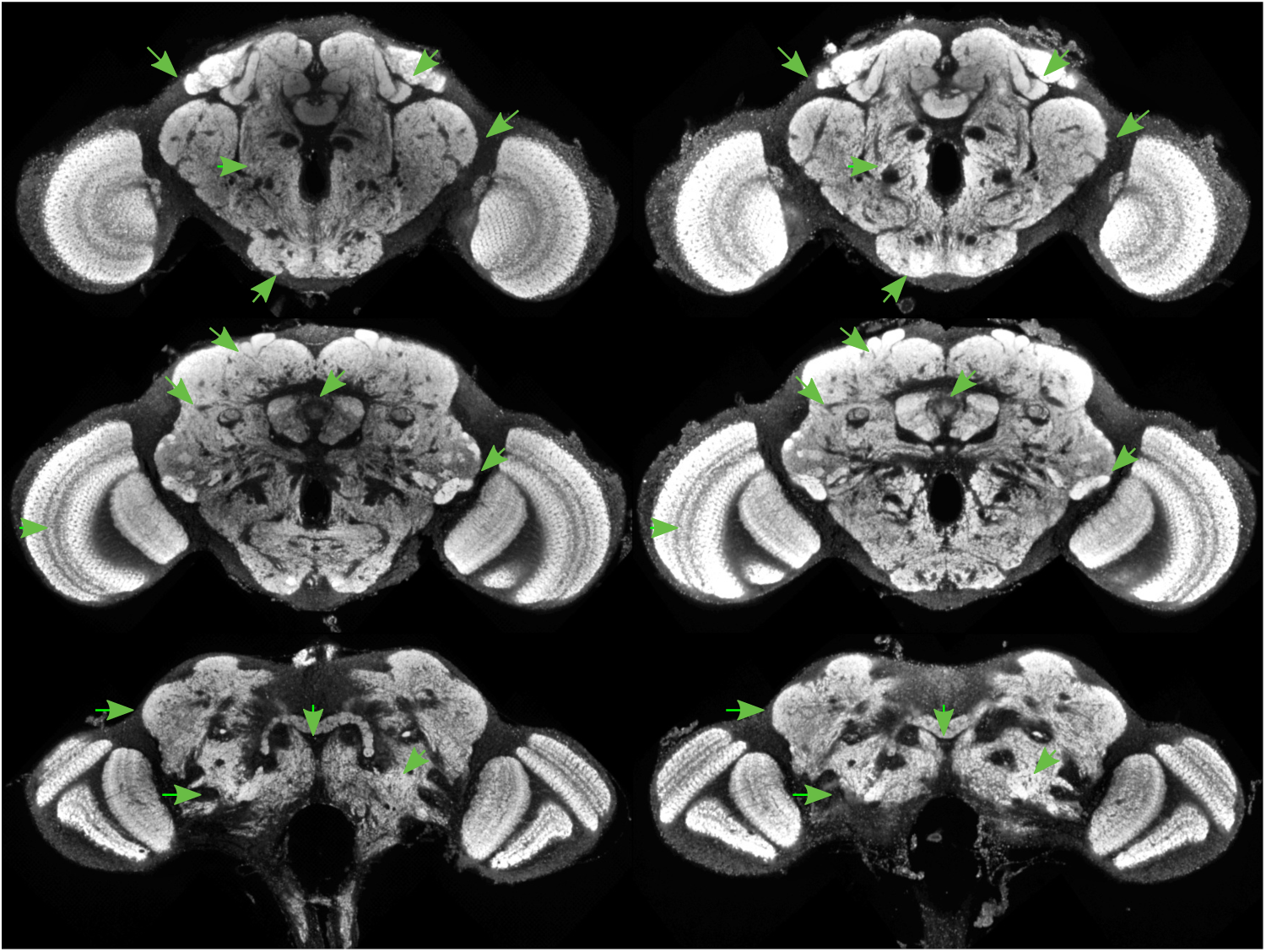
JFRC2013 cmtkC

**Figure 37:**
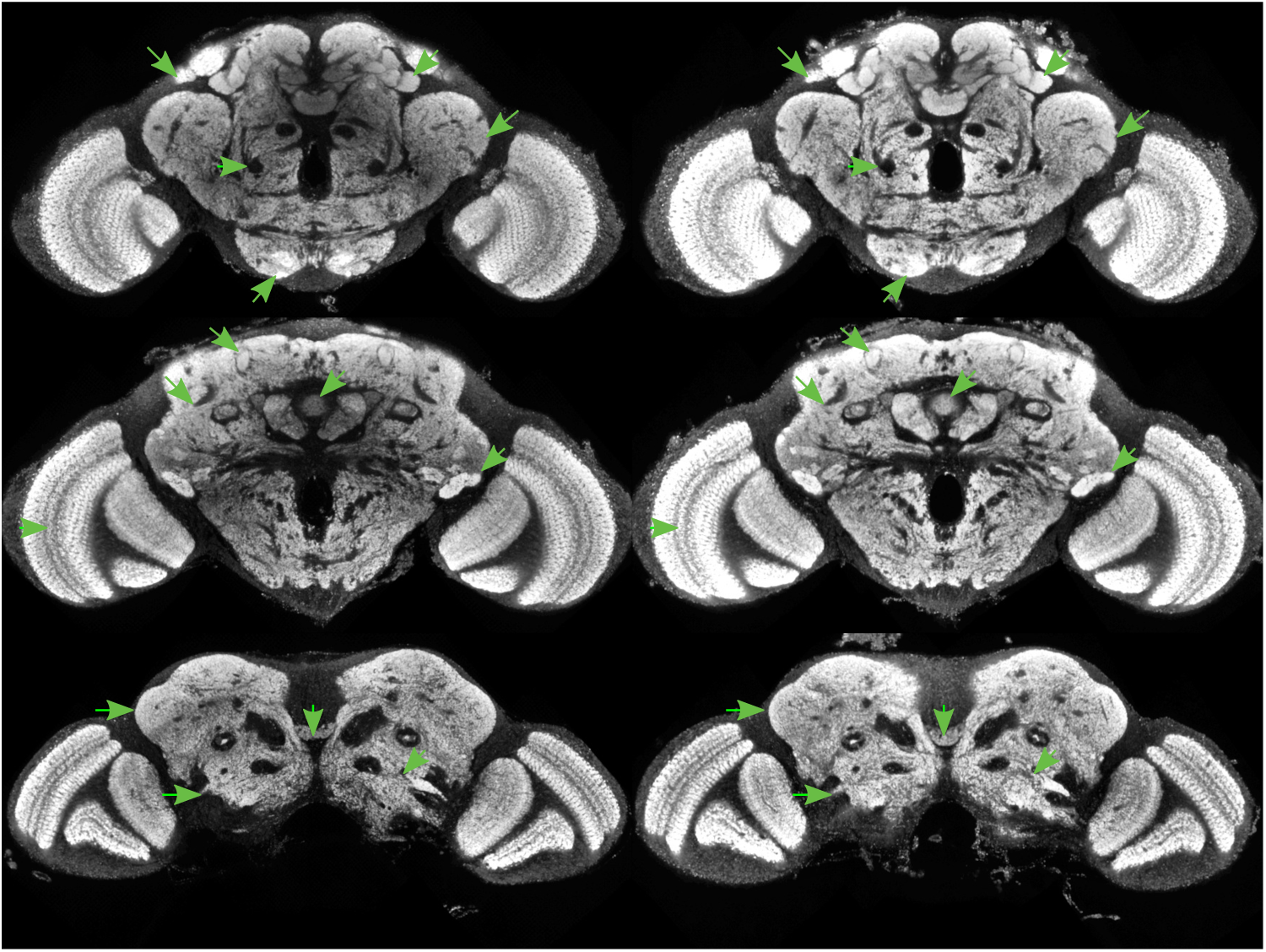
JFRC2013 elastixA

**Figure 38:**
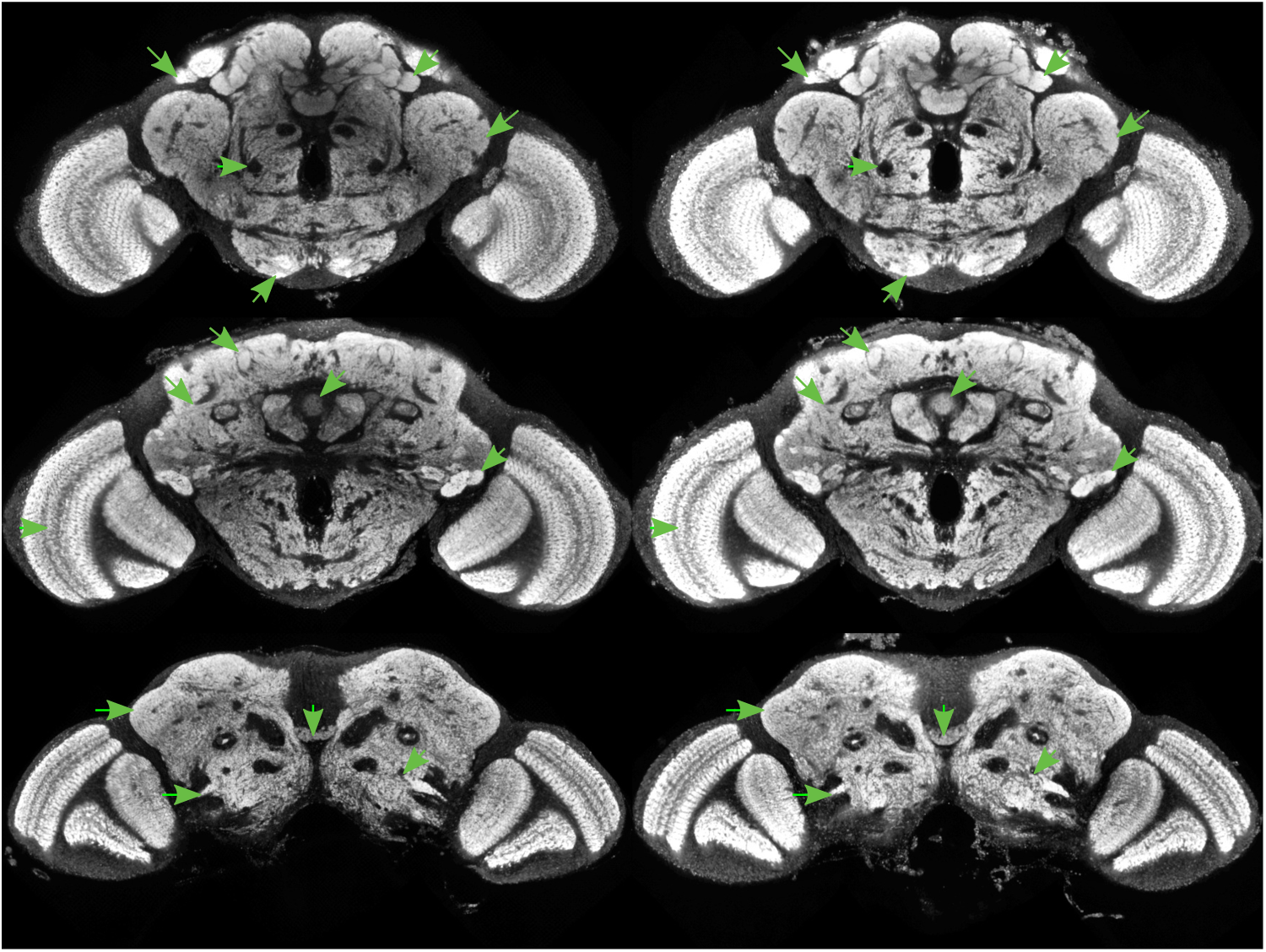
JFRC2013 elastixB

**Figure 39:**
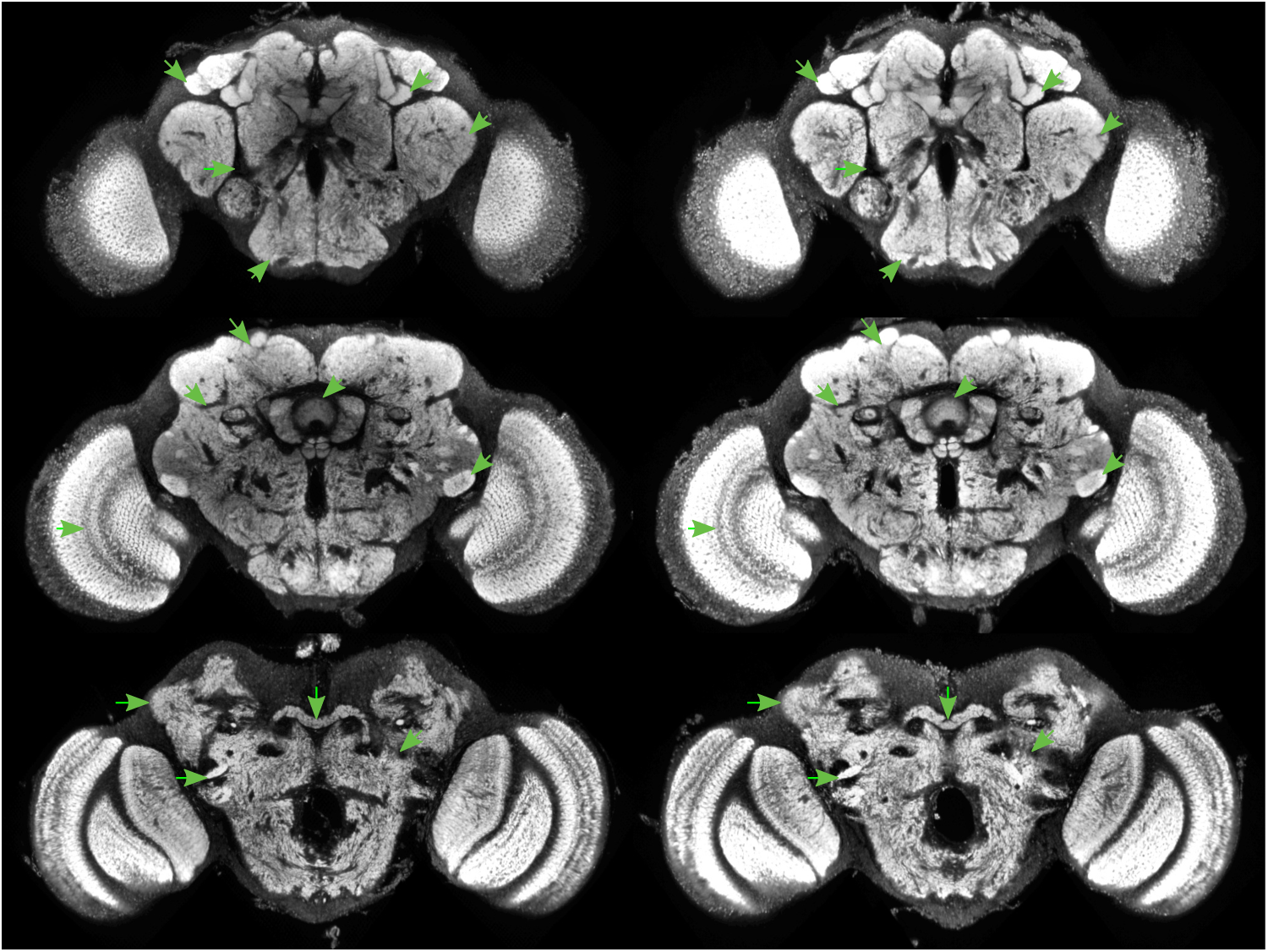
JFRC2010 antsA

**Figure 40:**
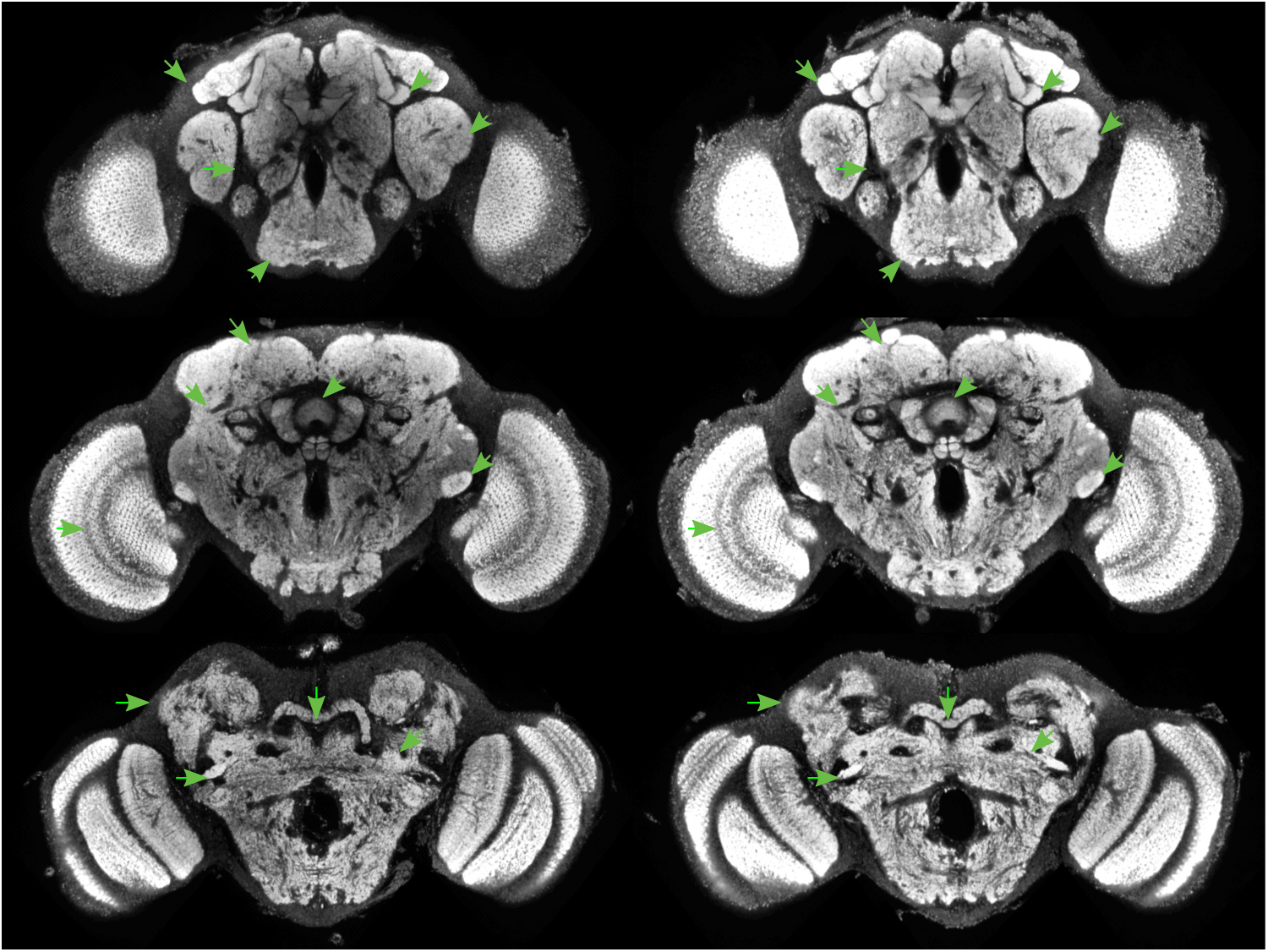
JFRC2010 antsB

**Figure 41:**
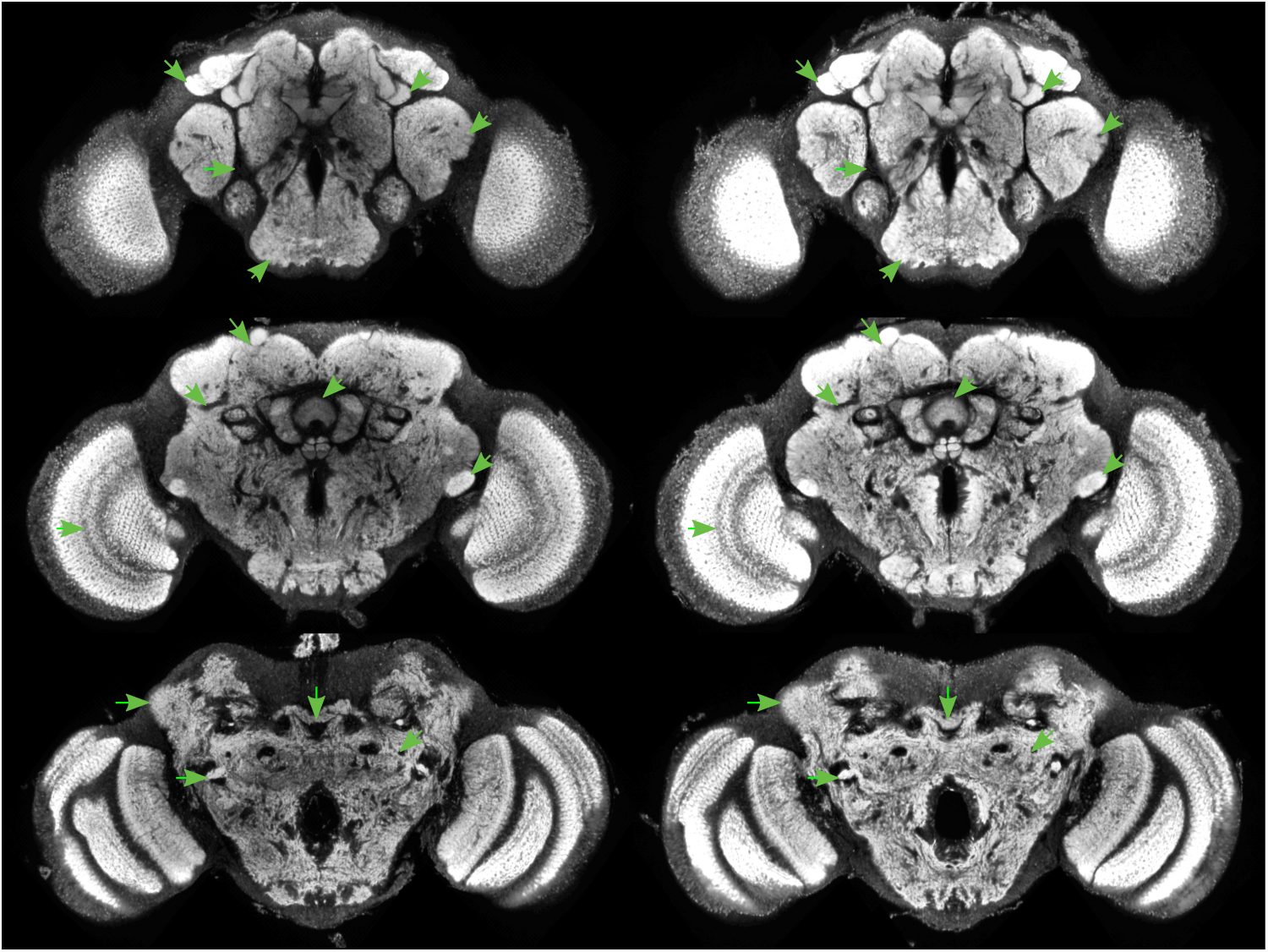
JFRC2010 antsC

**Figure 42:**
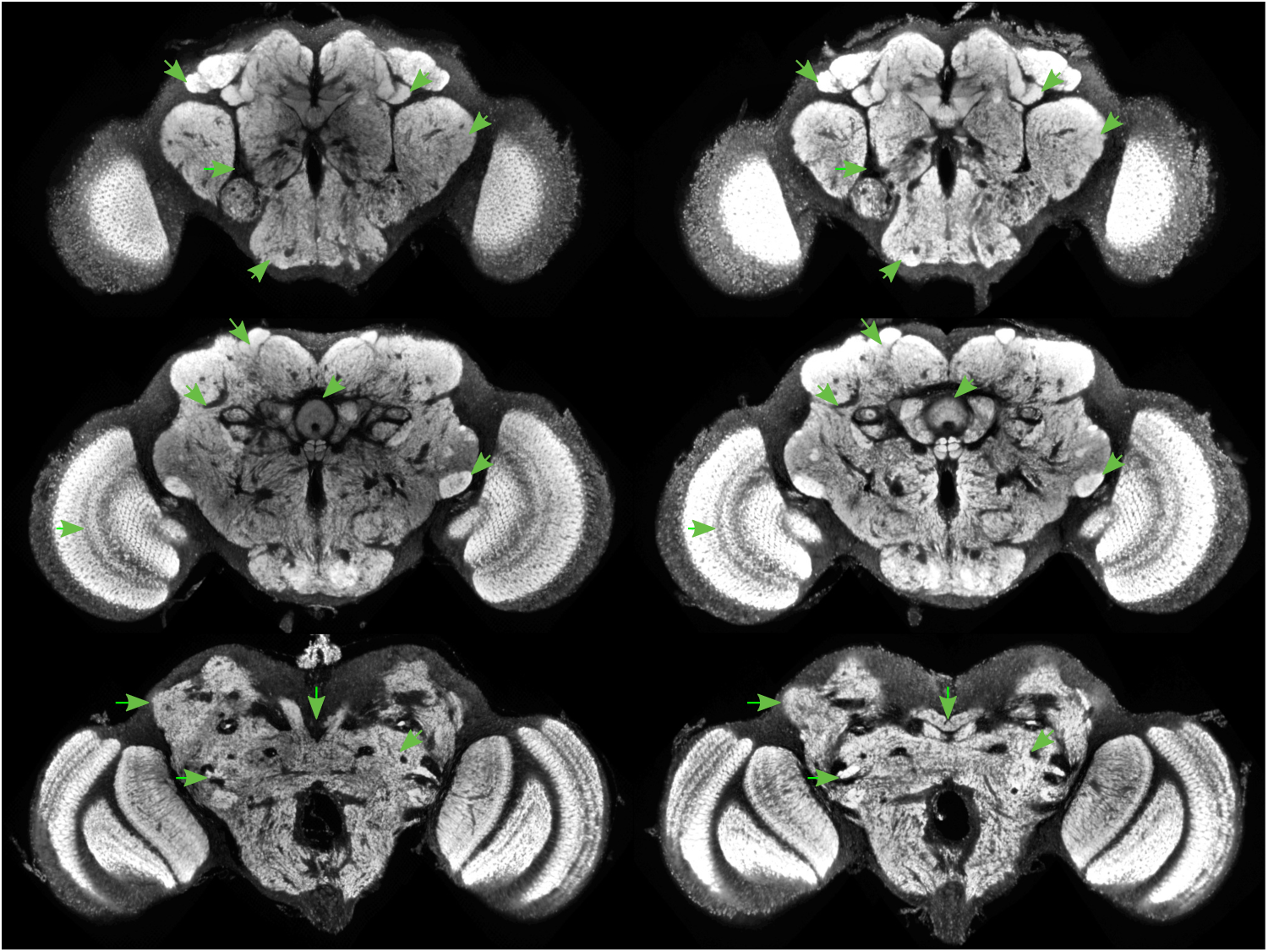
JFRC2010 cmtkA

**Figure 43:**
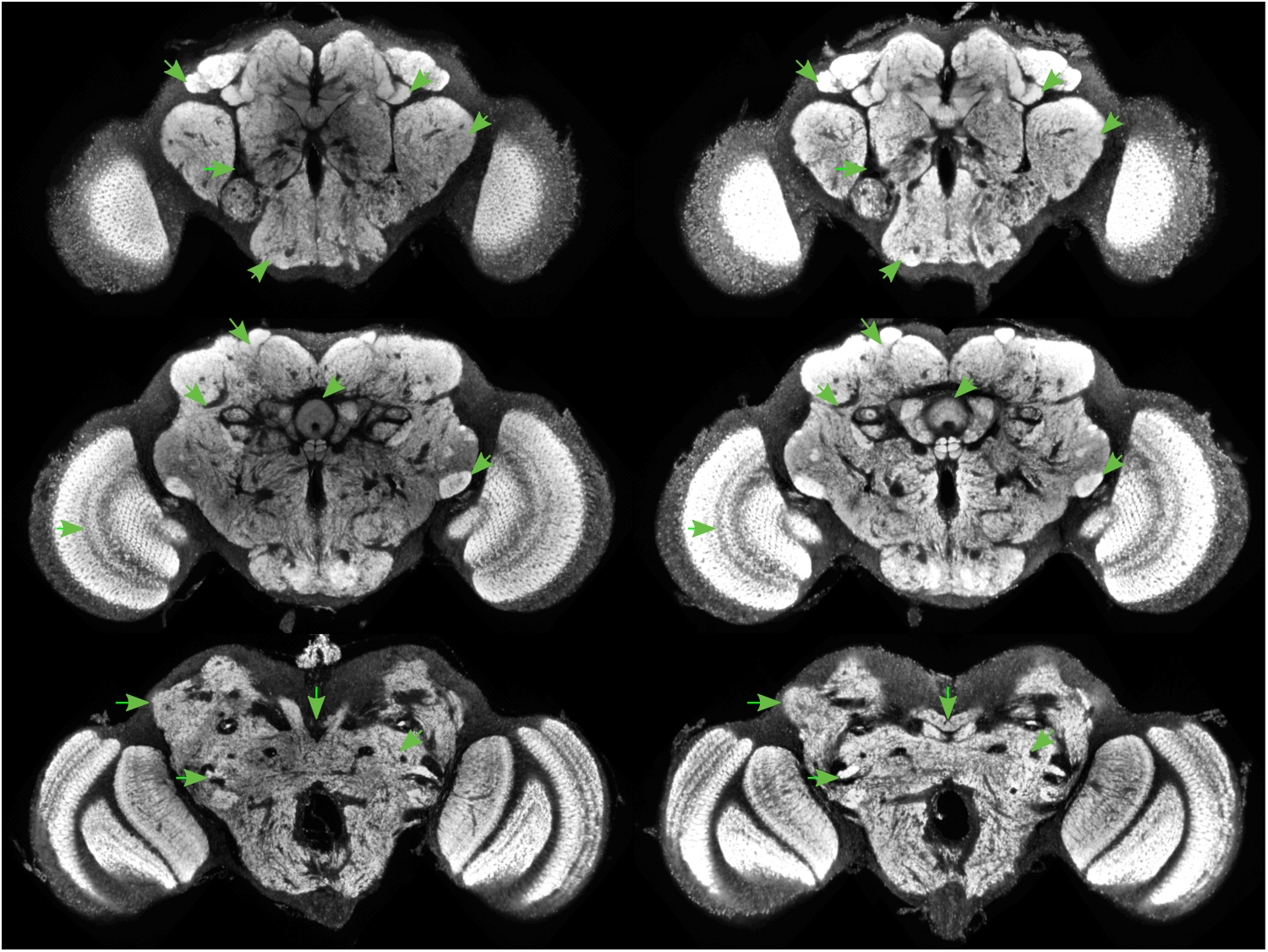
JFRC2010 cmtkB

**Figure 44:**
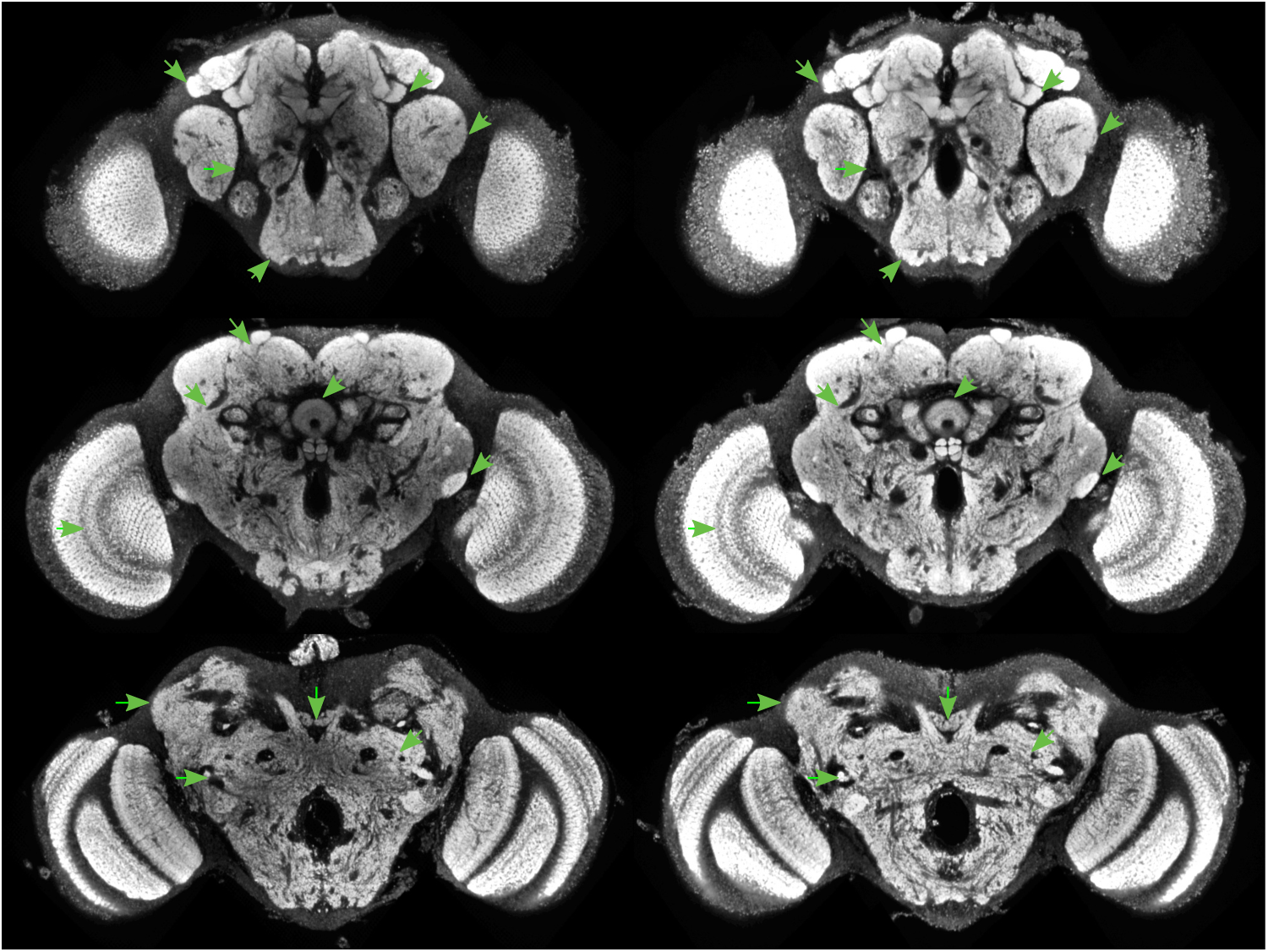
JFRC2010 cmtkC

**Figure 45:**
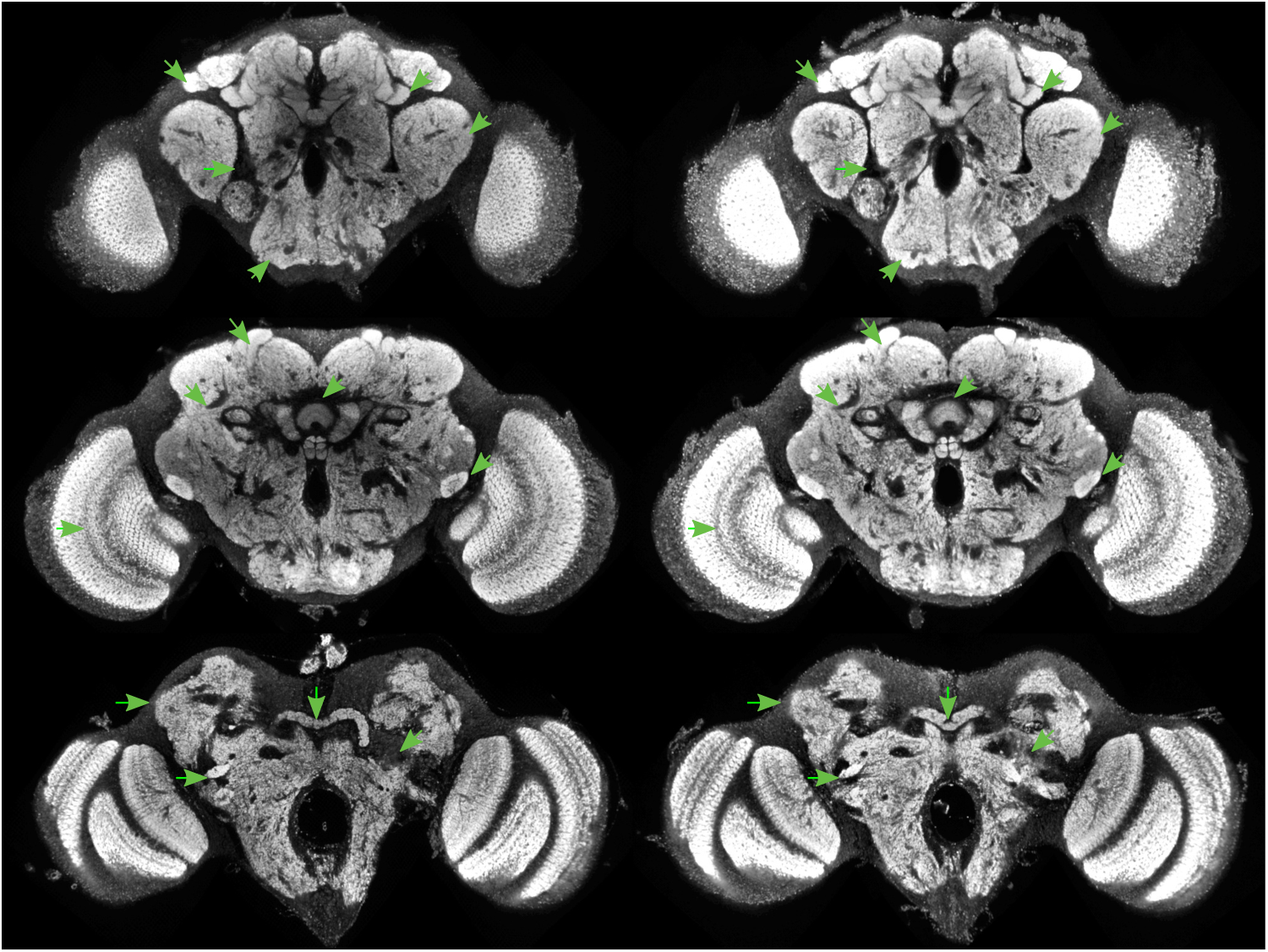
JFRC2010 elastixA

**Figure 46:**
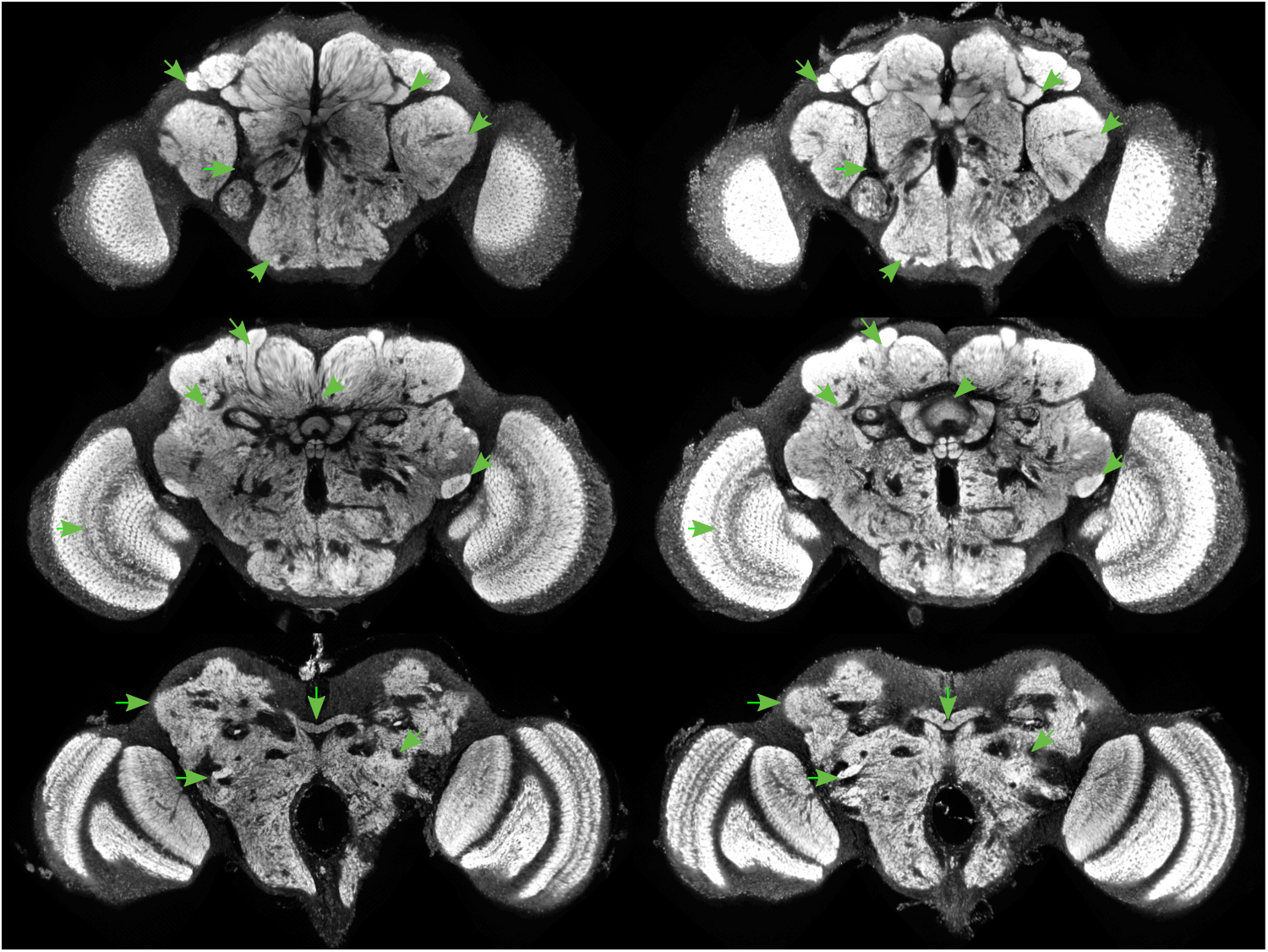
JFRC2010 elastixB

**Figure 47:**
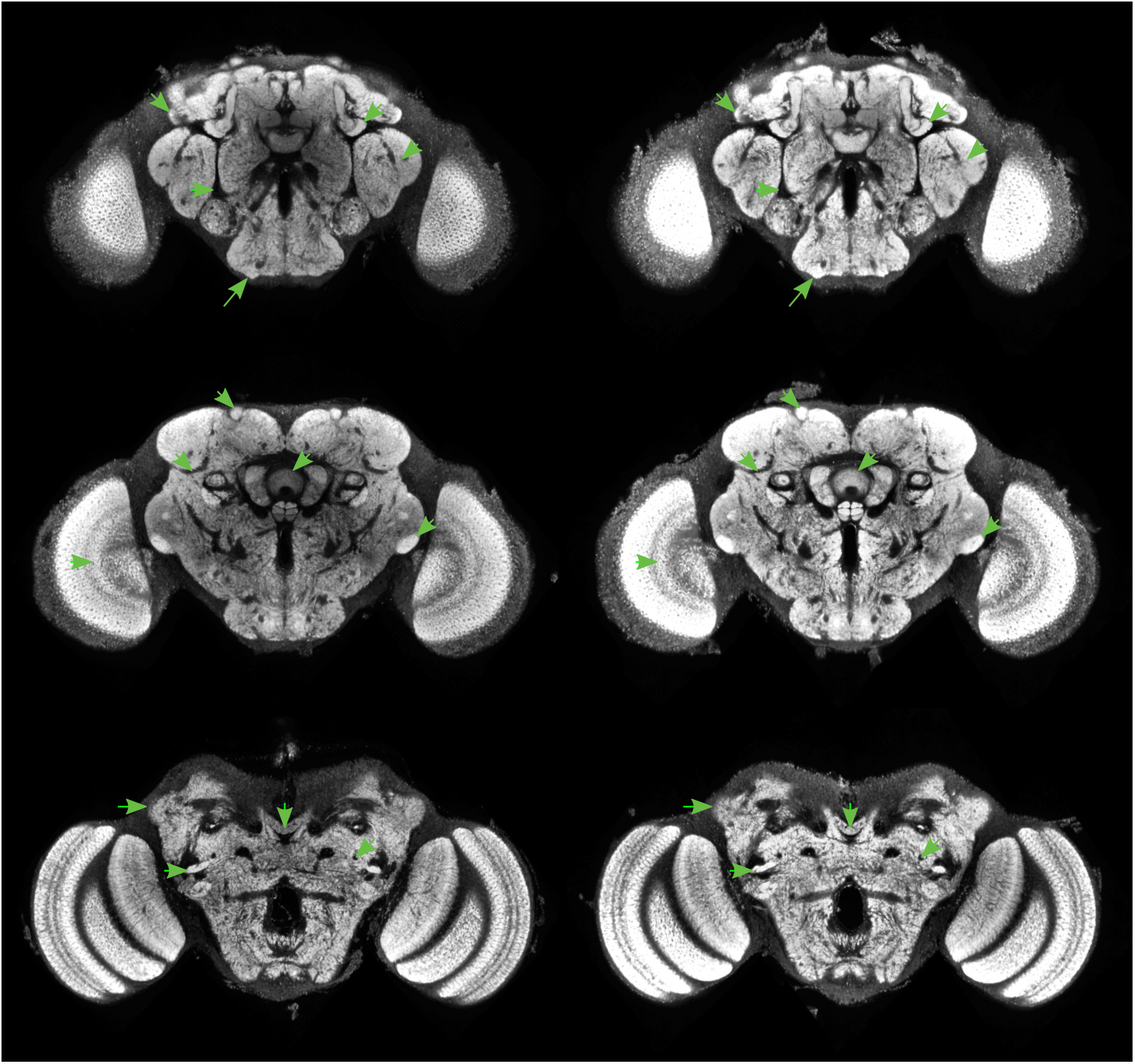
Tefor antsA

**Figure 48:**
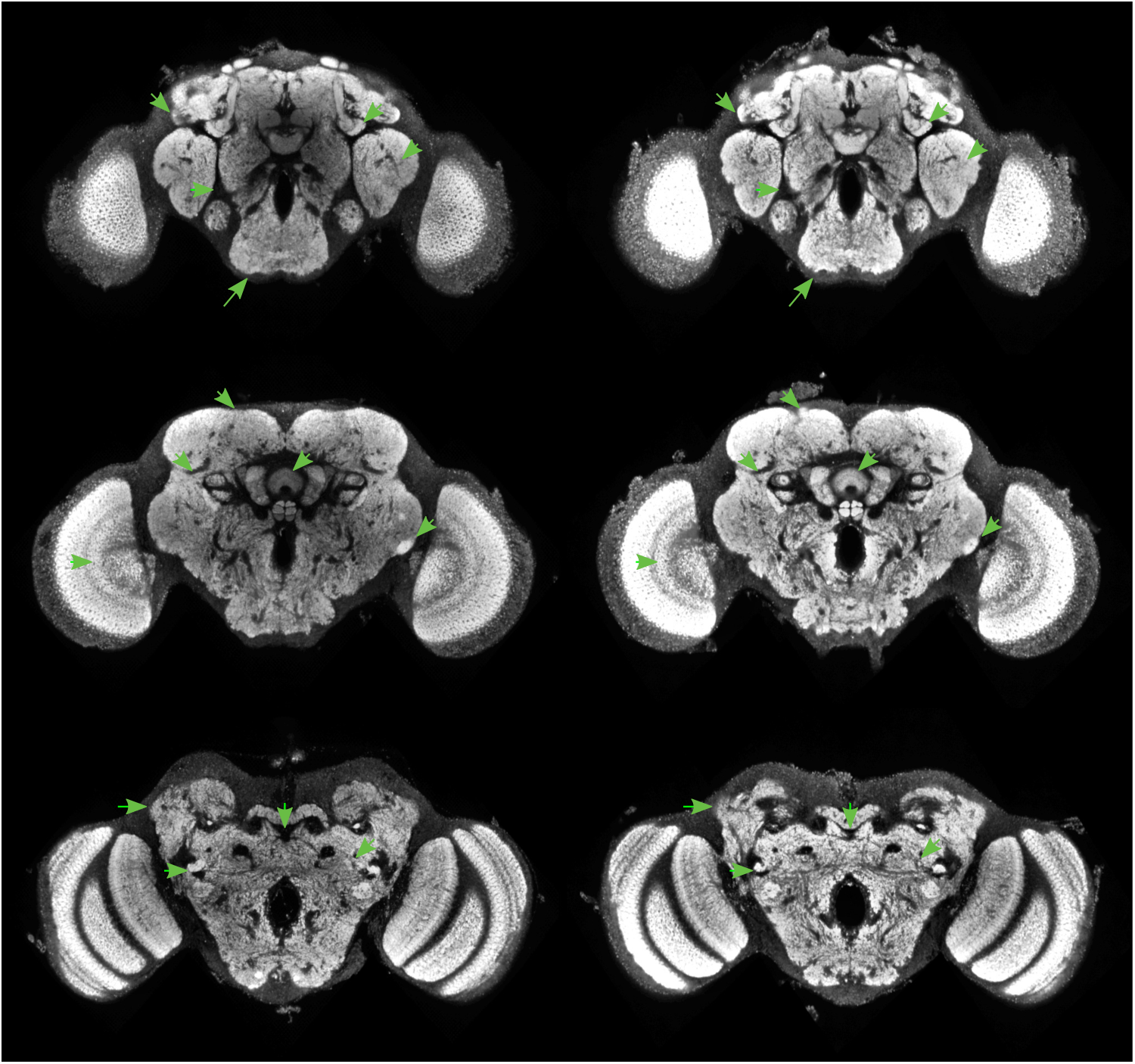
Tefor antsB

**Figure 49:**
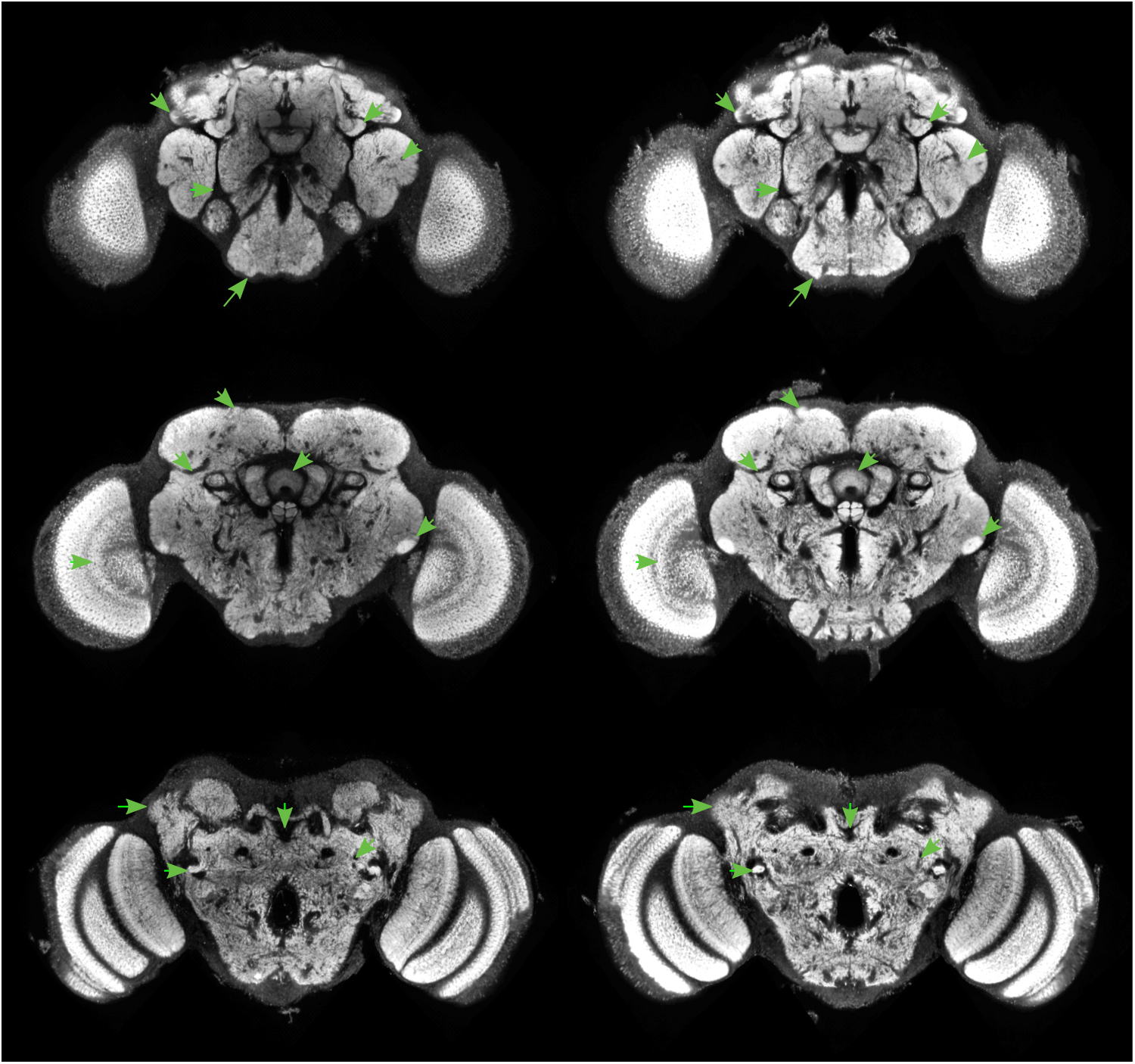
Tefor antsC

**Figure 50:**
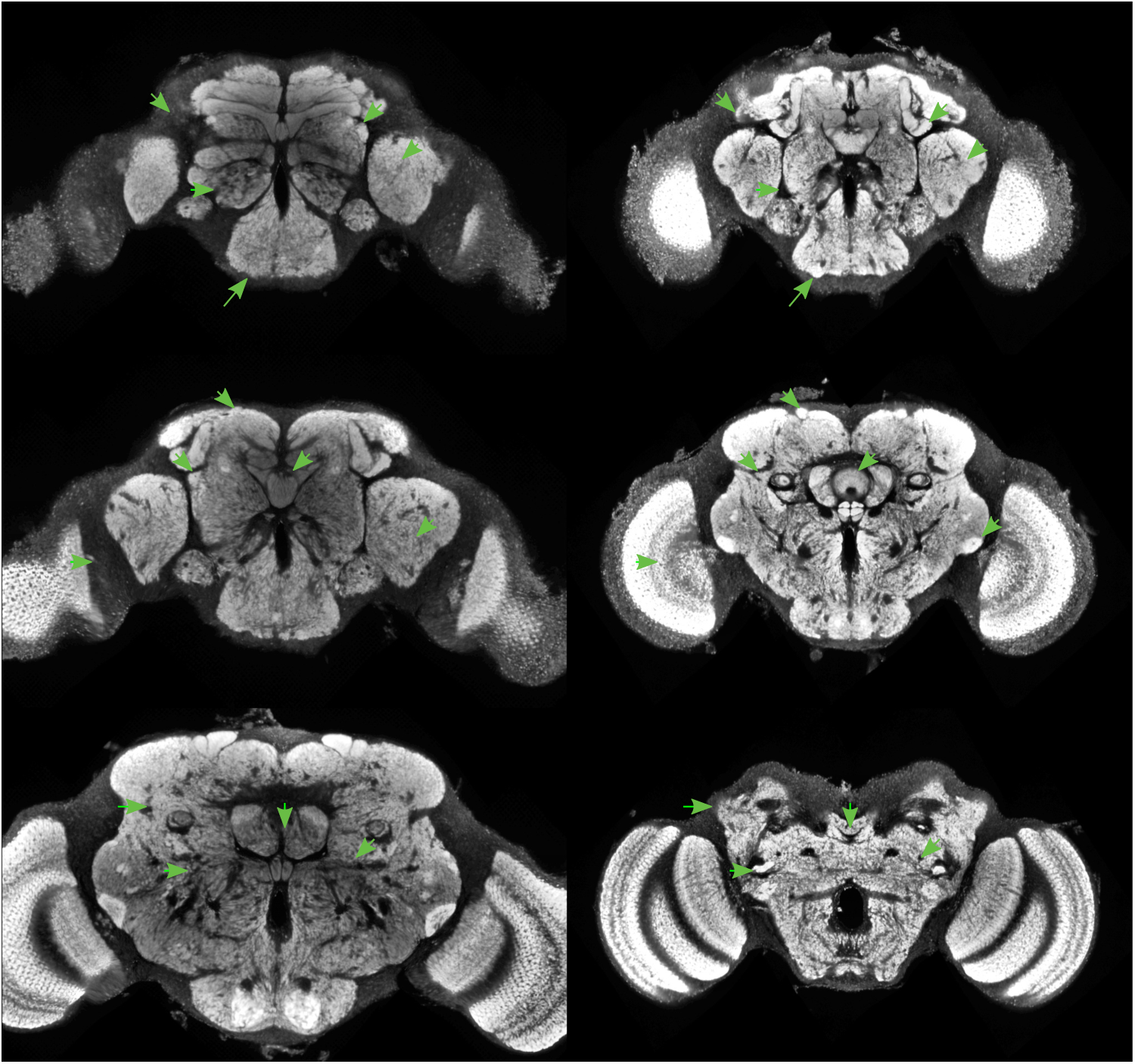
Tefor cmtkA

**Figure 51:**
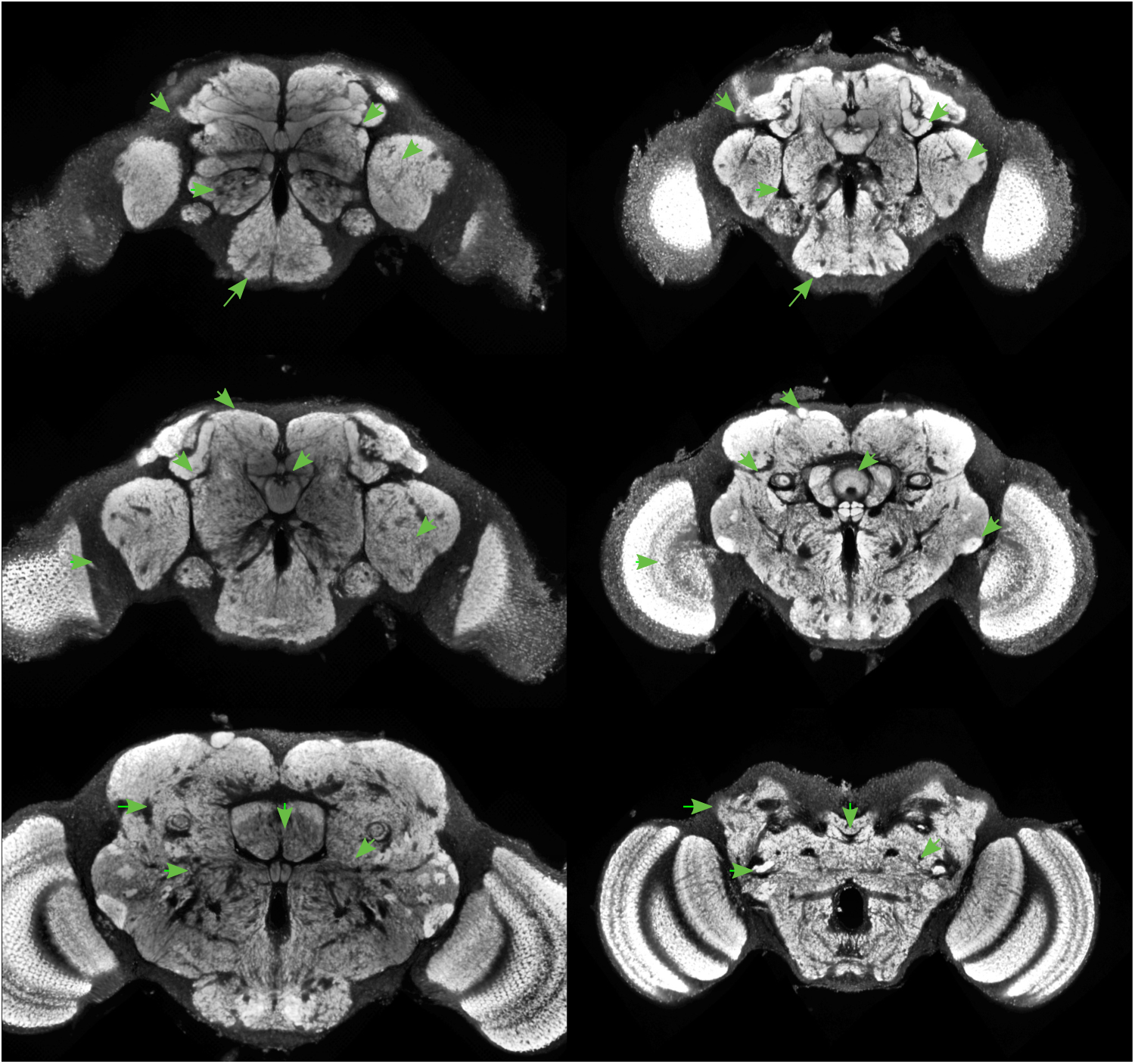
Tefor cmtkB

**Figure 52:**
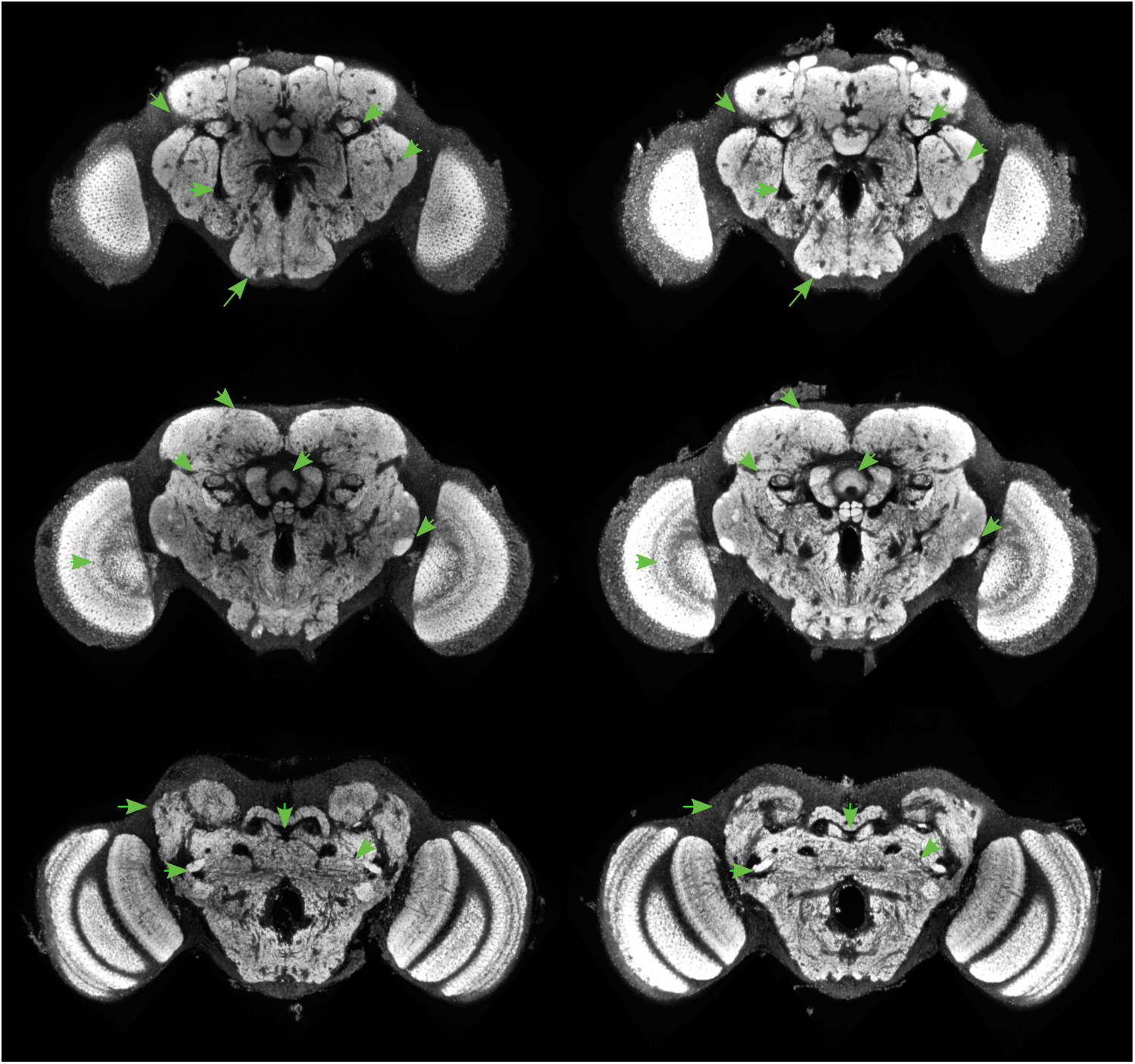
Tefor cmtkC

**Figure 53:**
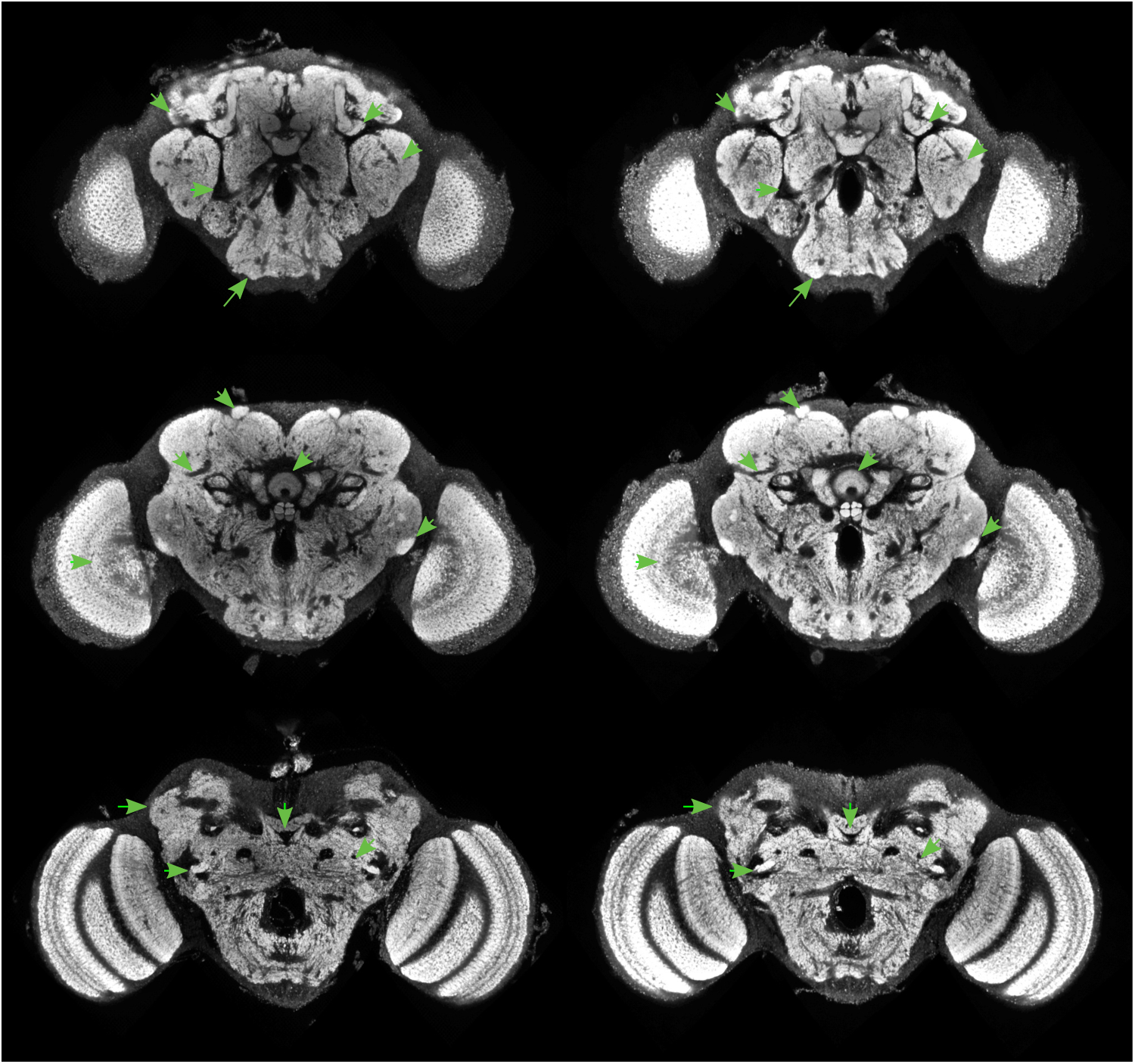
Tefor elastixA

**Figure 54:**
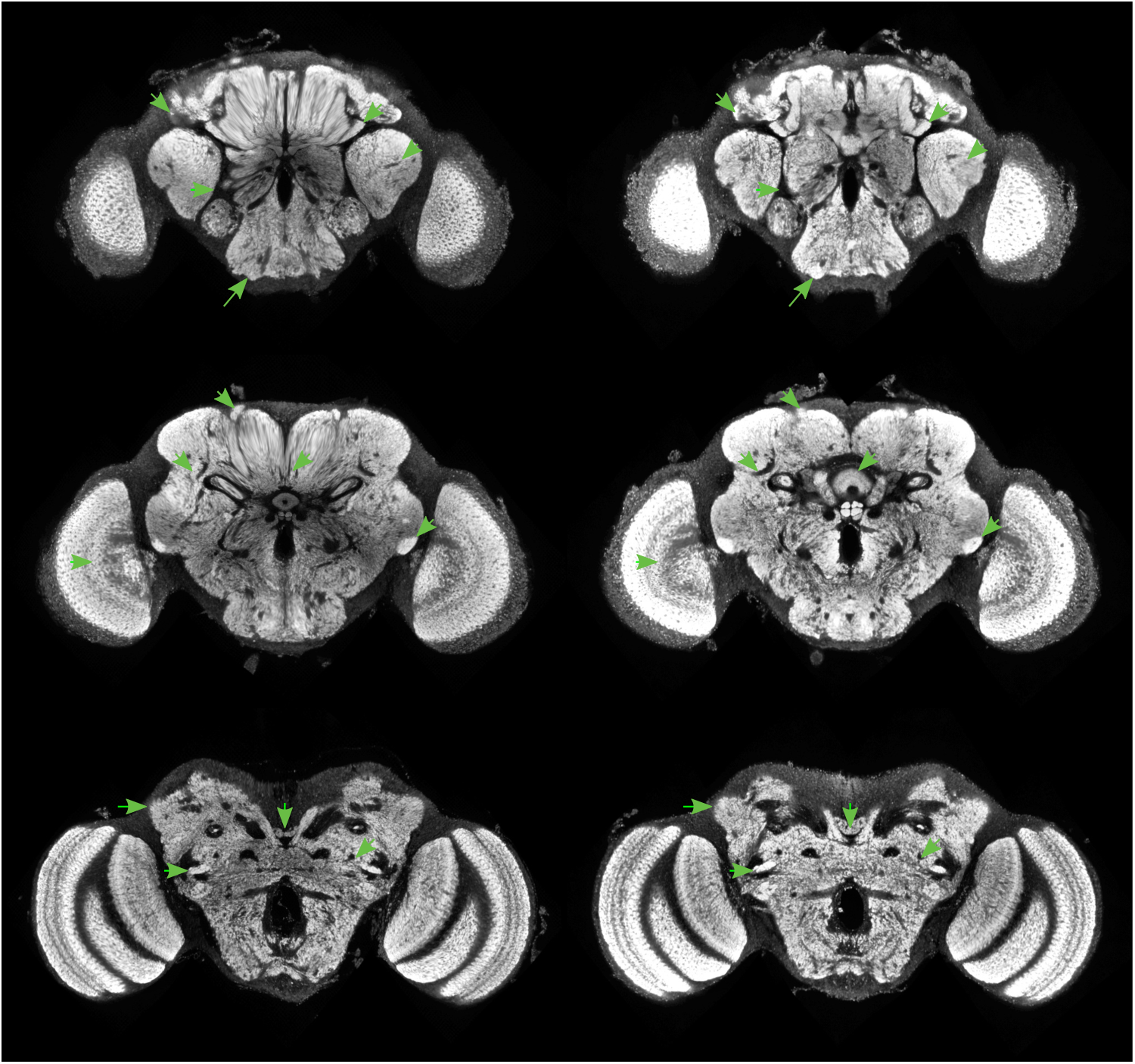
Tefor elastixB

